# The gut microbiota of environmentally enriched mice regulates visual cortical plasticity

**DOI:** 10.1101/2021.07.16.452307

**Authors:** Leonardo Lupori, Sara Cornuti, Raffaele Mazziotti, Elisa Borghi, Emerenziana Ottaviano, Giulia Sagona, Tommaso Pizzorusso, Paola Tognini

**Author notes:** These authors contributed equally to this work (Leonardo Lupori, Sara Cornuti). Co-last authors (Tommaso Pizzorusso, Paola Tognini).

## Abstract

Exposing animals to an enriched environment (EE) has dramatic effects on brain structure, function and plasticity. The poorly known “EE derived signals” mediating the EE effects are thought to be generated within the central nervous system. Here, we shift the focus to the body periphery, revealing that gut microbiota signals are crucial for EE-driven plasticity. Developmental analysis of intestinal bacteria composition in EE mice revealed striking differences from standard condition (ST) animals and enhanced levels of short-chain fatty acids (SCFA). Depleting the EE mice gut microbiota with an antibiotic cocktail decreased SCFA and prevented EE induction of adult ocular dominance (OD) plasticity, spine dynamics and microglia rearrangement. SCFA treatment in ST mice mimicked the EE induction of adult OD plasticity and morphological microglial rearrangement. Remarkably, transferring the microbiota of EE mice to ST recipients activated adult OD plasticity. Thus, taken together our data suggest that experience-dependent changes in gut microbiota regulate brain plasticity.

## INTRODUCTION

The complexity of brain circuits is sculpted both by innate genetic programs and environmental stimuli capable of activating brain plasticity and, in turn, reshaping brain circuits and behavior. This process is particularly evident in the development and refinement of neuronal circuits belonging to sensory modalities (Hübener and Bonhoeffer, 2014). Since the 1960s scientists noticed that raising rodents in an enriched housing condition characterized by elevated social interactions, cognitive, sensory and motor stimulations (Kempermann, 2019) could improve their learning and memory abilities (Rosenzweig and Bennett, 1996). Interestingly, the benefits of this enriched environment (EE) go beyond memory features and embrace several aspects of brain functionalities, including healthy development, emotional behavior, recovery from neural damage, and positive effects on a variety of preclinical models of neuropsychiatric diseases (Nithianantharajah and Hannan, 2006). Indeed, EE is known to enhance neuronal activation, signaling and plasticity throughout various brain regions, including sensory areas, and at different ages (Baroncelli et al., 2010). For example, ocular dominance (OD) plasticity, a change in eye preference of cortical cell responses induced by monocular deprivation (MD), is typically observed during a critical period of postnatal development (Hensch, 2005; Levelt and Hübener, 2012). However, EE mice showed strong OD plasticity (Greifzu et al., 2014), and recovery from amblyopia also during adulthood (Sale et al., 2007; Tognini et al., 2012). Moreover, EE was found to accelerate several molecular and functional aspects of visual cortical development (Cancedda et al., 2004), indicating that the visual system is highly influenced by EE both during development and adulthood.

A variety of mechanisms have been discovered to contribute to the EE plasticity enhancement, such as increased hippocampal neurogenesis, changes in the excitatory/inhibitory circuit ratio, promotion of structural plasticity of dendritic spines, alteration in BDNF levels, microglia rearrangement, chromatin remodeling and others (Baroncelli et al., 2010; Kempermann, 2019; Pizzorusso et al., 2007). So far, all the mechanisms explaining EE influence on neural plasticity have been searched inside the brain. However, the complexity of EE stimuli should bring benefits to the whole body. Thus, we seek to investigate if signals coming from the periphery could participate in EE-driven plasticity in the central nervous system (CNS), focusing on the gut microbiota.

The intestinal microbiota has emerged as a novel and complex regulator of system-wide physiology (Schroeder and Bäckhed, 2016). Classically, the gut microbes have been studied in the context of host metabolic homeostasis and immune system development and function (Nicholson et al., 2012; Rooks and Garrett, 2016; Tremaroli and Bäckhed, 2012). Nevertheless, the relationship between the intestinal flora and its host seems to be deeply intricate, going far beyond the influence on metabolism and immunity (Murakami and Tognini, 2019). In fact, recent reports have suggested the commensals to play an important role in brain-related processes: myelination (Gacias et al., 2016; Hoban et al., 2016), microglia maturation (Erny et al., 2015), neurogenesis (Möhle et al., 2016), blood-brain-barrier (BBB) permeability (Braniste et al., 2014), and finally influence on behavioral outcomes (Buffington et al., 2016, 2021; Forsythe and Bienenstock, 2016; Hsiao et al., 2013; Tognini, 2017). Despite this evidence, little is known about how intestinal bacteria impinge on neuronal function and especially, nobody has ever explored if there is a link between experience-dependent plasticity and the gut microbiota. In this study, we show that EE significantly alters the intestinal microbiota in C57BL/6J mice, and that microbiota manipulation through antibiotics interferes with EE-driven visual cortical plasticity, dendritic spine dynamics and microglial rearrangement. Strikingly, the fecal microbiota transplant (FT) from EE donors to adult standard (ST) mice was able to enhance OD plasticity in the ST recipients. Overall, our study introduces a new key concept: environment-dependent changes in the gut microbiota composition can regulate brain plasticity.

## RESULTS

### EE modulates the composition of the gut microbiota

C57BL/6J mice were reared from birth in ST (standard laboratory cage 26 × 42 × 18 cm, containing only bedding and nesting material), or EE condition (large cage 44 × 62 × 28 cm, containing also running wheels, and differently shaped objects: tunnels, shelters, toys, nesting material, Fig. 1a) from birth (EE birth). In order to investigate the intestinal bacteria composition, fresh faeces were collected in a longitudinal way from ST and EE birth mice before weaning: postnatal day (P) 20, a few days after weaning (P25) and during adulthood (P90) (Fig. 1b). The bacterial DNA was extracted from the faeces and analyzed through 16S rRNA sequencing (bacterial abundance are reported in Suppl. Table 1 and Suppl. Table 2).

**Fig. 1:**
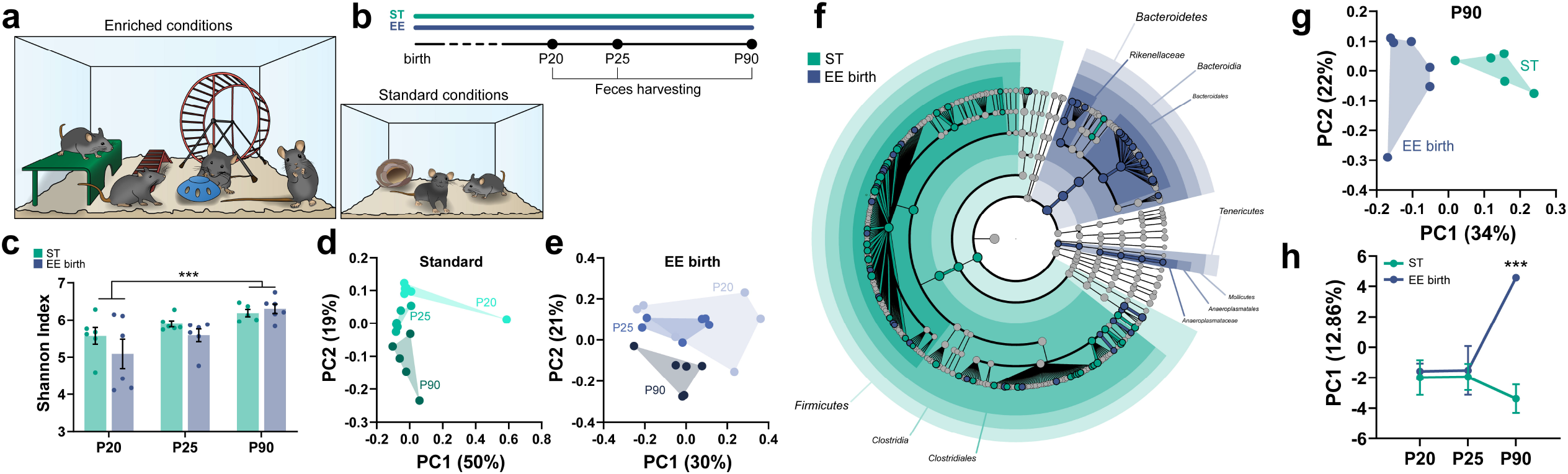
EE modulates the gut microbiota composition. (**a**) Schematic of enriched and standard housing conditions. (**b**) Experimental Timeline. (**c**) Shannon Index for all the conditions, showing alpha-diversity (N=6 per group, N=5 in P90 ST, two-way ANOVA, age*housing, age, F_2,29_=8.764 p=0.0011, post-hoc Tukey test P20 vs P90 p<0.001). (**d-e**) Principal Coordinates Analysis (PCoA), showing beta-diversity (unweighted UniFrac distance) at different ages in ST and EE birth animals (ANOSIM test, ST group: P20 vs P25 R=0.32 p=0.002, P25 vs P90 R=0.424 p=0.004, P20 vs P90 R=0.461 p=0.009; EE group: P20 vs P25 R=-0.007 p=0.46, P25 vs P90 R=0.483 p=0.0015, P20 vs P90 R=0.291 p=0.008). (**f**) Cladogram of differential bacterial composition (ST vs EE, P90) using LEfSe analysis. Shaded areas represent taxa significantly enriched in the one of the conditions. (**g**) PCoA between ST and EE birth animals at P90 (ANOSIM test, ST vs EE birth, P90: R= 0.445 p=0.0015). (**h**) First principal component from an integrated Principal Component Analysis (PCA) of OTU abundance of all the experimental groups (two-way ANOVA time*housing, interaction F_2,28_=8.482 p=0.0013, post-hoc Holm Sidak ST vs EE birth (P90) t_28_=5.18 p<0.001).

To analyze the results, we first calculated alpha-diversity, a parameter reflecting species richness in a microbial ecosystem (Thukral, 2017). We observed that there was a significant increase in alpha-diversity in both ST and EE mice between P20 and P90 (Fig. 1c, Suppl. Fig. 1). However, there were no inter-group differences at any age tested, indicating that housing animals in EE did not alter the alpha-diversity in the microbial ecosystem. Second, we computed the beta-diversity, which measures the degree of phylogenetic similarity between microbial communities, using the unweighted UniFrac algorithm (Lozupone and Knight, 2005). Principal coordinates analysis (PCoA) of unweighted UniFrac distance showed that the microbiota of ST animals clustered in different groups at P20, P25 and P90 (Fig. 1d). EE birth mice also displayed an age-dependent difference between P90 and P20 or P25 mice, although P20 and P25 were not significantly different (Fig. 1e).

Third, we sought to explore differences in the bacterial composition between EE birth and ST conditions by using Linear discriminant analysis (LDA) effect size (LEfSe) method (Segata et al., 2011). LEfSe revealed a relatively small, although significant difference in some taxa in P20 ST mice vs P20 EE, as shown by the cladogram in Suppl. Fig. 2a, and by the LDA score in Suppl. Fig. 2b. In particular, at P20 species in Mollicutes class, *Anearoplasmataceae* family and Anearoplasmatales order were significantly more abundant in EE with respect to ST. PCoA of the unweighted UniFrac distance matrix confirmed the phylogenetic similarities between ST and EE at P20 (Suppl. Fig. 2c). At P25 the cladogram displayed a situation similar to P20, in addition taxa in Erysipelotrichia class, Erysipelotrichales order, and *Erysipelotrichaceae* family were more abundant in EE than in ST mice (Suppl. Fig. 3a-b). Also, beta-diversity showed clustering in more separate groups (Suppl. Fig. 3c). Intriguingly, at P90 the composition of the microbiota in EE birth mice dramatically diverged from that of age-matched ST rodents, as displayed by the cladogram in Fig. 1f. The enrichment in taxa belonging to Tenericutes already present at P20 and P25 was maintained. Furthermore, species present in the family *Rikenellaceae*, the order Bacteroidales and the class Bacteroidia, all included in the phylum Bacteroidetes, were more abundant in EE. On the other hand, several taxa belonging to the phylum Firmicutes were significantly enriched in ST (Fig. 1f, Suppl. Fig. 4). As expected, PCoA of beta diversity calculated through the unweighted UniFrac distance demonstrated that EE and ST microbes clustered in separate groups at P90 (Fig. 1g) indicating robust differences in the membership of gut bacteria between EE and ST mice.

Principal component analysis (PCA) of OTU abundance comparing ST and EE birth mice at all ages and considering the first component (PC1) indicated that the differences in the microbiota composition were significant only at P90, while higher similarity was present in microbes composition at P20 and P25 (Fig. 1h).

Thus, our 16S rRNA-seq data demonstrated a progressive developmental divergence of the microbiota composition between mice living in EE and ST condition. Considering that EE is able to promote different forms of plasticity in a variety of brain areas during adulthood, we further explored the possible link between gut microbiota and EE-driven plasticity in the adult visual cortex.

### Depletion of the gut microbiota in EE mice prevented OD plasticity

Short periods of MD do not induce OD plasticity in adult ST mice (Lehmann and Löwel, 2008), whereas they induce strong plasticity in adult EE mice, demonstrating that EE potently activates visual cortical plasticity (Greifzu et al., 2014; Sale et al., 2007). To investigate the contribution of signals coming from the intestinal microbes on EE-induced visual cortical plasticity, the microbiota of the animals living in EE was depleted using a wide spectrum antibiotic cocktail (ABX) in drinking water. The ABX treatment started 1 week before birth (ABX in dams’ water) and continued until P120 (Fig. 2a). The efficacy of the treatment was demonstrated by 16S rRNA sequencing of fecal bacteria after ABX administration (Suppl. Fig. 5a-b). In order to test OD plasticity, mice were subjected to optical imaging of the intrinsic signal (IOS, Fig. 2b-d). IOS was performed before and after three days of MD (3dMD) in the same subjects (Fig. 2a). Strikingly, ABX treatment completely prevented OD plasticity in adult EE mice (Fig. 2e-f). As a control, we also confirmed that 3dMD elicited an OD shift in ST mice during the critical period, but not in adult mice (Suppl. Fig. 6a-b). Overall, these data indicate that an intact intestinal microbiota is necessary for the enhancement of adult plasticity observed in EE rodents.

**Fig. 2:**
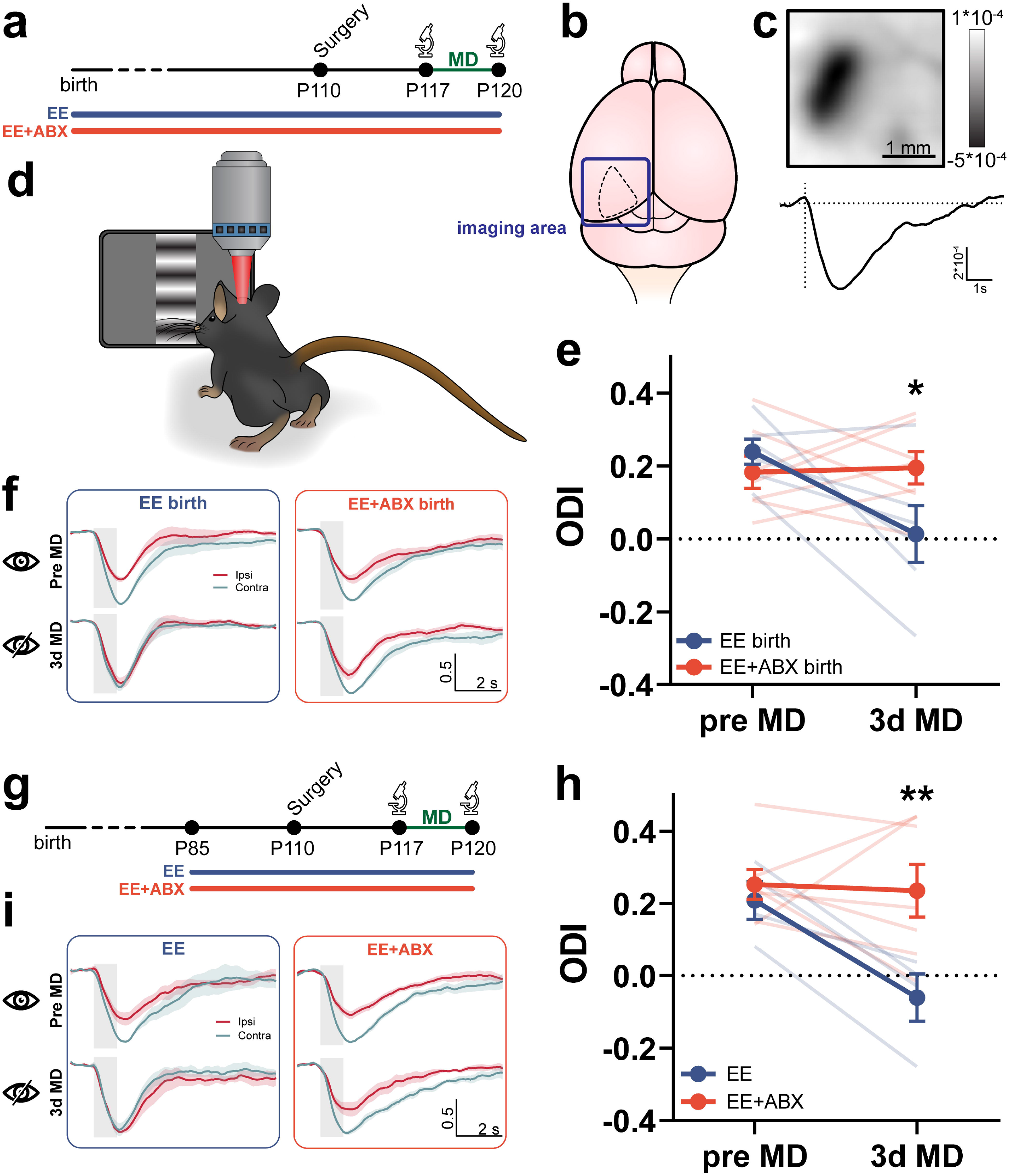
ABX treatment prevents the plasticity enhancing effect of EE. (**a**) Experimental Timeline. (**b,d**) Schematic of the imaging setup and imaging area. (**c**) Representative intrinsic signal response (top: response image, bottom: signal timeline) from the binocular primary visual cortex to visual stimulation. (**e**) Ocular Dominance Index (ODI) before and after monocular deprivation. Thin lines represent single animals (N=6 EE-birth, N=7 EE+ABX-birth, two-way RM ANOVA time*housing, interaction F_1,11_=6.716 p=0.025, post-hoc Holm-Sidak, EE-birth vs EE+ABX-birth (3dMD) t_11_=3.3 p=0.013). (**f**) Normalized response timeline (mean±SEM) to contralateral (contra) and ipsilateral (ipsi) eye stimulation before and after MD for both groups. Shaded area represents visual stimulus duration. (**g**) Experimental Timeline. (**h**) ODI before and after monocular deprivation. (N=4 EE, N=7 EE+ABX, two-way RM ANOVA time*housing, interaction F_1,9_=6.18 p=0.035, post-hoc Holm-Sidak, EE vs EE+ABX (3dMD) t18=3.25 p=0.009). (**i**) Normalized response timeline (mean±SEM) to contralateral (contra) and ipsilateral (ipsi) eye stimulation before and after MD for both groups. Shaded area represents visual stimulus duration.

To rule out any developmental effects of the ABX treatment from birth, we performed the 3dMD experiment in mice reared in EE and treated with ABX for only 5 weeks from P85 until P120 (Fig. 2g). This short EE protocol during adulthood was sufficient to induce OD plasticity in response to 3dMD (Fig. 2h-i, EE group). The induction of OD plasticity by brief EE was also sensitive to microbiota depletion. Indeed, if ABX was administered simultaneously to EE, the EE-driven plasticity was blocked (Fig. 2h-i, EE+ABX group).

Importantly, before 3dMD, visual responses and baseline OD of both EE+ABX birth and EE+ABX mice were not different from their controls (Suppl. Fig. 6c,d), thus excluding a possible direct effect of ABX treatment on cortical responses.

In conclusion, an unperturbed intestinal microbiota is necessary for EE promotion of OD plasticity.

### EE effects on dendritic spines and microglia morphology are blocked by ABX administration

Dendritic spines are protrusions harboring the postsynaptic machinery of excitatory synapses. Spines are dynamic compartments and their structural remodeling is thought to be one of the signatures of the ongoing rewiring of neuronal circuits (Holtmaat and Svoboda, 2009). Since EE is known to increase the density and dynamics of dendritic spines in the cortex (Ali et al., 2019; Xu et al., 2016), we asked whether manipulating the gut microbiota could interfere with EE-driven plasticity through mechanisms involving spines remodelling. We performed repeated two-photon in vivo imaging to analyze dendritic spines in a cohort of transgenic Thy1-GFP-M mice expressing GFP in a sparse population of layer V pyramidal neurons in the visual cortex (Feng et al., 2000; Jung and Herms, 2014). We acquired images of the same apical dendritic segments for 40 days in the visual cortex (Fig. 3a). After two baseline imaging time points (1-day interval), animals were put in EE cages for 35 days (5 weeks) and received ABX in drinking water (EE+ABX group) or regular water (EE group) (Fig. 3b). The same dendritic segments were imaged in the two groups 9,10,19,20,34, and 35 days after the beginning of EE (Fig. 3b-d). The analysis of spine density revealed a significant interaction between time and housing conditions. Post-hoc tests showed that spine density in EE mice was significantly larger than in EE+ABX mice at all times after EE start, but it did not differ at baseline. Comparing spine density within each group at different time points confirmed that spine density did not change over time in the EE+ABX group. By contrast, the EE group displayed a dramatic increase in spine density occurring between the baseline and the first imaging time point after the beginning of EE (10 days after EE start, Fig. 3e-f). This observation was confirmed analyzing the data from single animals. Indeed, animals in the EE group showed a significant 25% increase in spine density after 10 days of EE with respect to baseline levels, while this effect was prevented in animals treated with ABX (Fig. 3h). To further dissect this effect, we calculated spine formation and elimination rates during this critical time window (Fig. 3g, Suppl. Fig. 7a). Across the first 10 days of EE, spine elimination rates were not different between the two groups, however, animals treated with ABX had a remarkably lower spine formation rate compared to controls, thus explaining the lack of increase in spine density in the ABX treated group (Fig. 3i). Moreover, EE+ABX group had a significantly lower spine formation rate with respect to EE controls also analyzing the short-term 24h spine dynamics. Indeed, short-term formation rate was comparable between the two groups at baseline, but significantly differed after 10 days of EE. Conversely, elimination rates were similar between the two groups (Fig. 3l). These data suggest that the ABX treatment prevents the increase in spine density typical of enriched animals by interfering with the process of formation of new spines and not by destabilizing existing spines or increasing the opposing process of spine elimination. This interpretation is further corroborated by the analysis of dendritic spines survival curves. Indeed, both for the subset of spines originally present at the first time point and for the subset of newborn spines that appeared after the start of EE, the treatment with ABX did not have any effect on survival curves (Suppl. Fig. 7b).

**Fig. 3:**
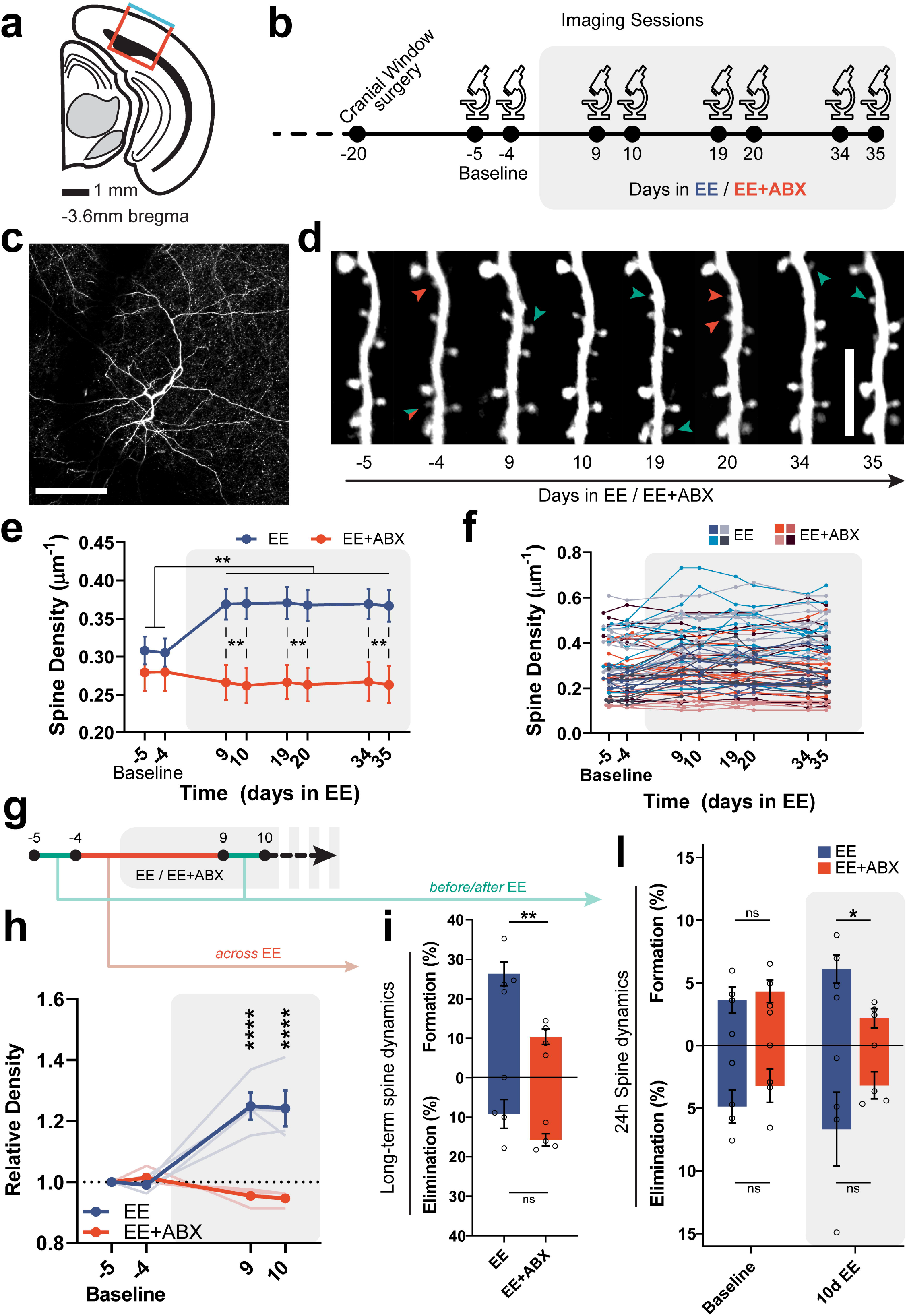
ABX treatment modulates the effect of EE on dendritic spines. (**a**) Schematic coronal section of the primary visual cortex showing the imaging site. (**b**) Experimental Timeline. (**c**) Representative low magnification image of a neuron. Scale bar=100 μm. (**d**) Representative high-magnification images of a dendritic segment at 8 consecutive time points. Green arrowheads: spine formation, red arrowheads: spine elimination. Scale bar=10 μm. (**e-f**) Spine density over time. Averages per dendrites (e) and single dendrites (f), color coded for the animals to which they belong. (N=39 EE, N=28 EE+ABX, two-way RM ANOVA time*treatment, interaction F_7,455_=20.97 p<0.0001, post-hoc Holm-Sidak, EE vs EE+ABX (9d- 35d) t_54-61_>3.15 p<0.01 for all comparisons; EE(9d-35d) vs EE(baseline) t_38_>5.73 p<0.0001 for all comparisons. (**g**) Schematic of the analysis for spine dynamics in (i) and (l). (**h**) Spine density relative to the first imaging time point. Thin lines represent single animals. (N=4 EE, N=4 EE+Abx, two-way RM ANOVA time*treatment, interaction F_3,18_=28.60 p<0.0001, post-hoc Holm-Sidak, EE vs EE+Abx (9d-10d) t_24_>7.52 p<0.0001 for al comparisons; EE(9d-10d) vs EE(-4) t_18_>7.56 p<0.0001 for al comparisons). (**i**) Long-term (10d) spine dynamics across the beginning of EE. (N=4 EE, N=4 EE+ABX, Formation: unpaired 2-tailed t-test t6=4.42 p=0.0045; Elimination: unpaired 2-tailed t-test t_6_=1.65 p=0.15) (**l**) Short-term 24h spine dynamics before and after EE. (N=4 EE, N=4 EE+ABX, Formation: two-way RM ANOVA time*treatment, interaction F_1,6_=6.620 p=0.042, post-hoc Holm-Sidak EE vs EE+ABX (10dEE) t_12_=2.87 p=0.028; Elimination: two-way RM ANOVA time*treatment, no significant differences).

OD plasticity and activity-dependent regulation of dendritic spines during the critical period of the visual cortex involve microglial cells (Cheadle et al., 2020; Sipe et al., 2016). Previous reports demonstrated that EE housing resulted in more numerous and longer microglial processes in different brain regions (Ali et al., 2019; Xu et al., 2016). Thus we asked whether prevention of EE effects by ABX treatment could also be associated with altered microglial morphology. To answer this question, we performed immunofluorescence for IBA-1, a specific microglial marker, in fixed sections of the visual cortex of three experimental groups: adult mice housed in EE for 5 weeks receiving ABX in drinking water or drinking regular water, and age-matched ST animals. Quantitative three-dimensional morphometric analysis of microglia highlighted striking differences between ST and EE housed mice (Fig. 4a). Using Sholl analysis, which reflects microglial process arborization, we found a robust and significant increase in microglia complexity in the EE mice compared to ST group (Fig. 4b). This difference was particularly compelling at a distance of 14-40 μm from the soma (See Suppl.Table 3 for Tukey’s multiple comparisons test). Importantly, the ABX treatment completely prevented the effect of EE on microglia, maintaining the complexity to the levels of the ST group (Fig. 4a-b).

**Fig. 4:**
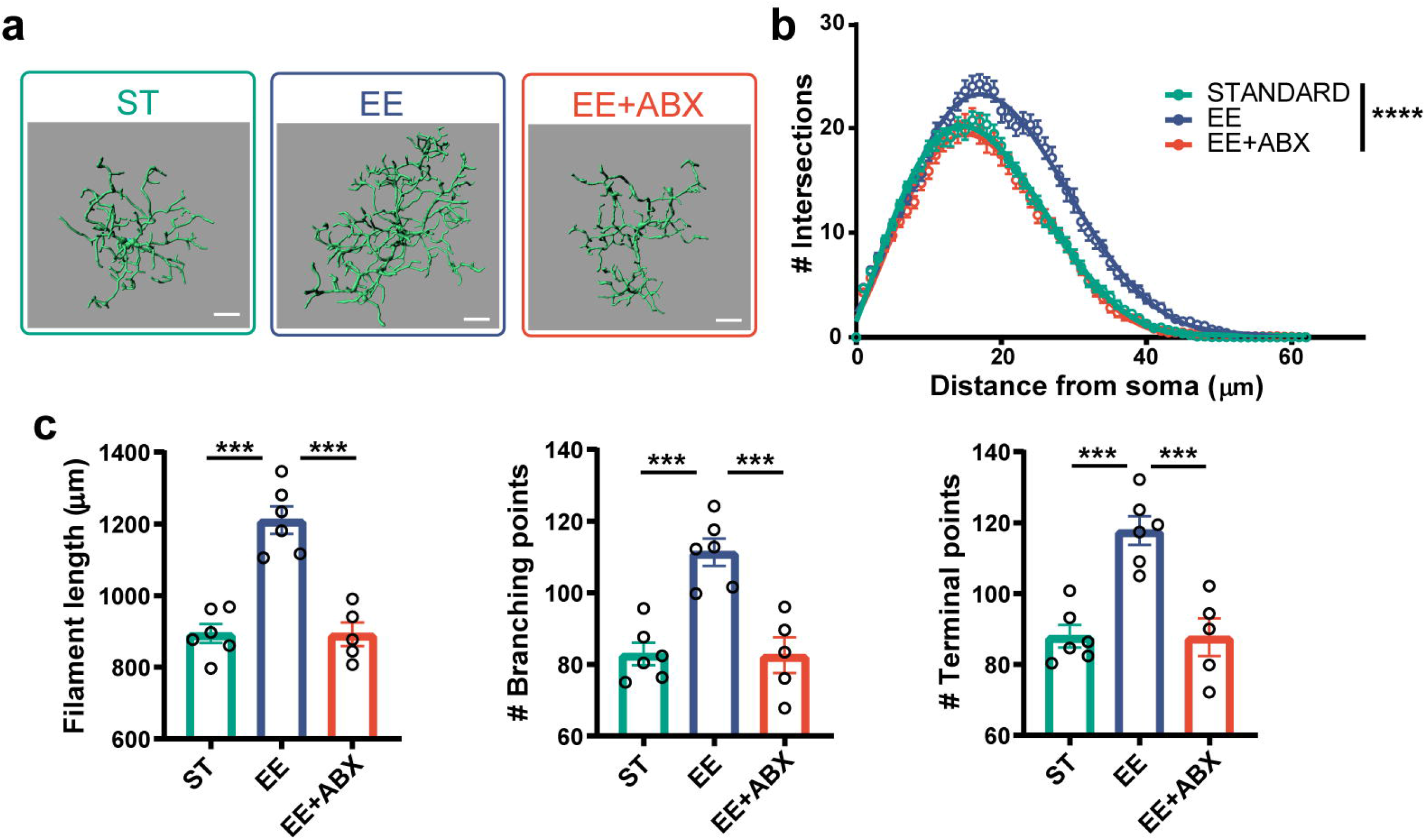
ABX treatment alters microglia morphology in EE mice. (**a**) IMARIS semi-automatic three-dimensional reconstruction (scale bar: 10μm) and quantitative morphometric analysis of IBA-1+ microglia. (**b**) Sholl analysis of microglial cells (N=30 EE, N=27 EE+ABX, N=32 ST two-way ANOVA distance*treatment, interaction F_124,5332_=5.886, p<0.0001, see Suppl. Table 3 for Tukey’s post-hoc comparisons. (**c**) Filaments length (Ordinary one-way ANOVA, F2,14=31.35 p<0.0001, Tukey’s post-hoc ST vs EE p<0.0001, EE vs EE+ABX p<0.0001). (**d**) Number of branching points (Ordinary one-way ANOVA, F_2,14_=18.22 p=0.0001, Tukey’s post-hoc ST vs EE p=0.0003, EE vs EE+ABX p=0.0005). (**e**) Number of terminals (Ordinary one-way ANOVA, F_2,14_=18.14 p=0.0001, Tukey’s post-hoc ST vs EE p=0.0003, EE vs EE+ABX p=0.0005). (**c-e**) N=6 EE, N=5 EE+ABX, N=6 ST each symbol represents one mouse. At least five cells measured per mouse.

To further explore the morphological changes, we examined single features of microglia ramification. We observed significantly shorter processes, and a significant decrease in the number of branching points and terminal points in mice housed in EE and subjected to ABX treatment compared to animals in EE condition alone, while ST animals displayed parameters very similar to EE+ABX (Fig. 4c-e, Suppl. Fig. 8).

Overall, these data indicate that ABX treatment dramatically affects the microglial morphology of EE mice.

### SCFA administration enhances OD plasticity in adult mice and affects microglia morphology

To dissect the possible mechanisms through which the gut microbiota of EE mice could promote cortical plasticity in adulthood, we focused on commensals derived metabolites. Short-chain fatty acids (SCFAs), the main metabolites produced in the colon by bacterial fermentation of dietary fibers and resistant starch, have been shown to play a key role in neuro-immuno-endocrine regulation, and in several aspects of the gut-microbiota-brain axis (Dalile et al., 2019). GC-MS analysis of fecal derived SCFA showed a significant increase in EE with respect to ST mice in acetate, propionate and butyrate concentration, whereas ABX administration completely blocked this effect (Suppl. Fig. 9). Based on those results, we explored the possibility that SCFA could be microbiota-derived metabolites mediating the pro-plasticity effect of EE. A SCFA mix, containing butyrate, propionate and acetate, was dissolved in the drinking water and administered to standard adult mice for 4 weeks. During the last week of treatment, mice were subjected to IOS imaging before and after 3dMD (Fig. 5a). Strikingly, SCFA in drinking water were able to promote visual cortical plasticity in adult ST mice, while, as expected, control age-matched ST animals drinking regular water did not show the OD shift (Fig. 5b-c). Importantly, SCFA administration also altered the morphology of microglial cells in the visual cortex mimicking EE effects (Fig. 5d). Indeed, SCFA treatment significantly increased process arborization (Fig. 5e and Suppl. Table 4 for Sidak’s multiple comparisons test), filament length, the number of branching points and terminal points in SCFA treated mice with respect to ST controls (Fig. 5f-h, Suppl. Fig. 10).

**Fig. 5:**
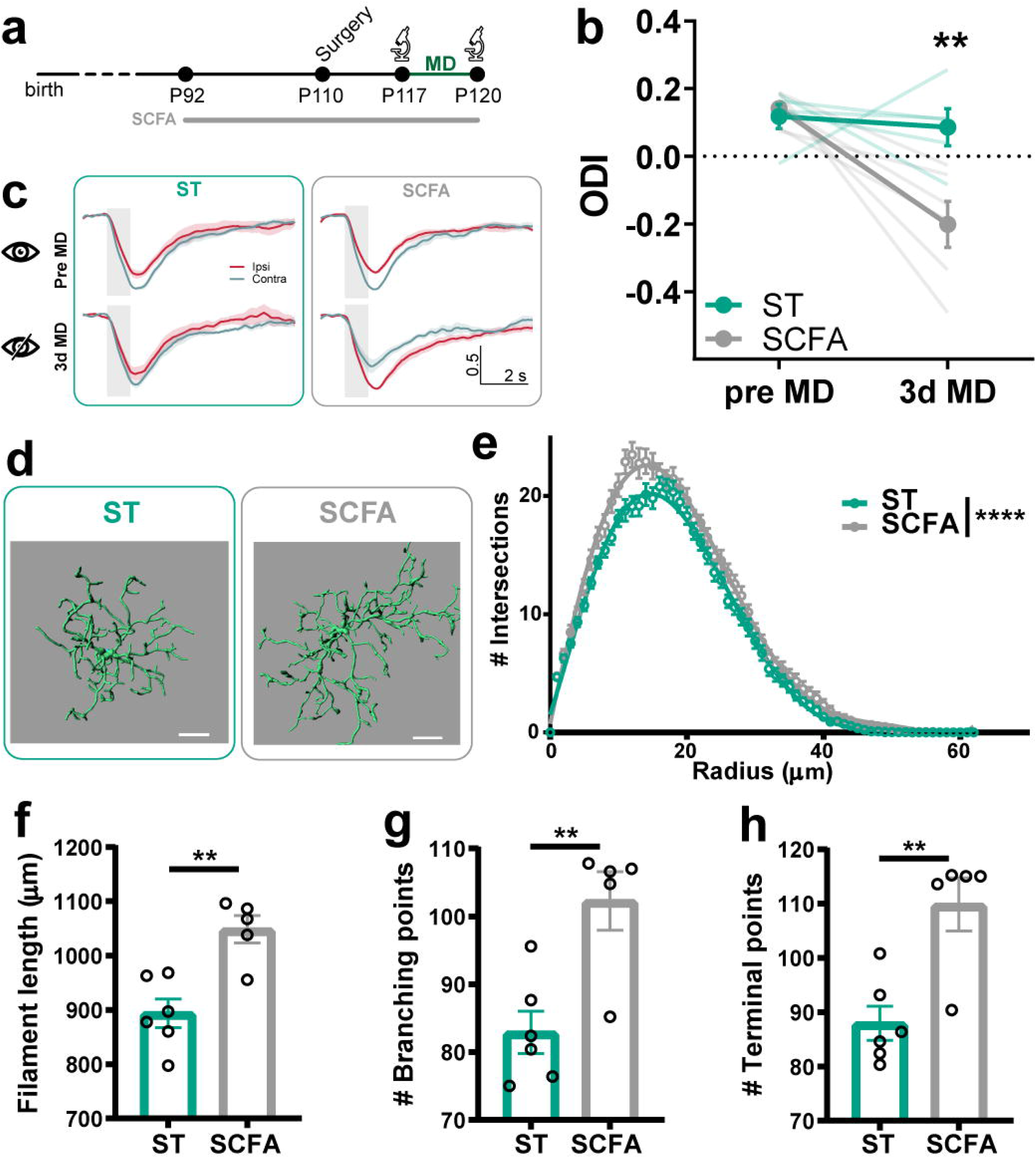
SCFAs administration mimics the effect of EE on plasticity and microglia morphology. (**a**) Experimental Timeline. (**b**) ODI before and after monocular deprivation (MD). Thin lines represent single animals (N=5 ST, N=6 SCFA, two-way RM ANOVA time*treatment, interaction F_1,9_=6.89 p=0.028, post-hoc Holm-Sidak, ST vs SCFA (3dMD) t_18_=4.10 p=0.014). (**c**) Normalized response timeline (mean±SEM) to contralateral (contra) and ipsilateral (ipsi) eye stimulation before and after MD for both groups. Shaded area represents visual stimulus duration. (**d**) IMARIS semi-automatic three-dimensional reconstruction (scale bar: 10μm) and quantitative morphometric analysis of IBA-1+ microglia. (**e**) Sholl analysis of microglial cells (N=32 ST, N=25 SCFA, two-way ANOVA distance*treatment, interaction F_62,3410_=1.903, p<0.0001, see Suppl. Fig. 11b for Sidak’s post-hoc comparisons). (**f**) Filaments length (unpaired 2-tailed t-test, t_9_=4.166 p=0.0024). (**g**) Number of branching points (unpaired 2-tailed t-test, t_9_=3.722 p=0.0048). (**h**) Number of terminals (unpaired 2-tailed t-test, t_9_=3.911 p=0.0036). (**f-h**) N=6 ST, N=5 SCFA, each symbol represents one mouse. At least five cells measured per mouse.

Our results suggest that SCFA could be candidate molecules through which the microbiota influence cortical plasticity, possibly involving microglia remodelling mechanisms.

### Faecal transplantation transfers the plastic phenotype from EE to ST adult mice

To provide further independent evidence to support the role of intestinal microbes in OD plasticity, adult ST mice were subjected to a FT experiment. Fresh faeces from EE mice or ST animals as a control were collected and a PBS suspension was freshly prepared. The FT was performed in conventionally raised ST recipient mice, previously treated with ABX to favor the engraftment of the new species (Fig. 6a). FT mice were subjected to oral gavage with the donor feces suspension for 5 days. To exclude a direct effect of the ABX pre-treatment on plasticity, a further control group of mice was administered with PBS (the vehicle of the donor feces suspension). After four weeks, OD plasticity was investigated through IOS imaging (Fig. 6a). Remarkably, transferring faeces from EE donors to ST recipients was able to induce visual cortical plasticity in ST mice, while, as expected, control animals inoculated with feces from ST donor mice or the PBS vehicle did not display any OD shift after 3dMD (Fig. 6b-c).

**Fig. 6:**
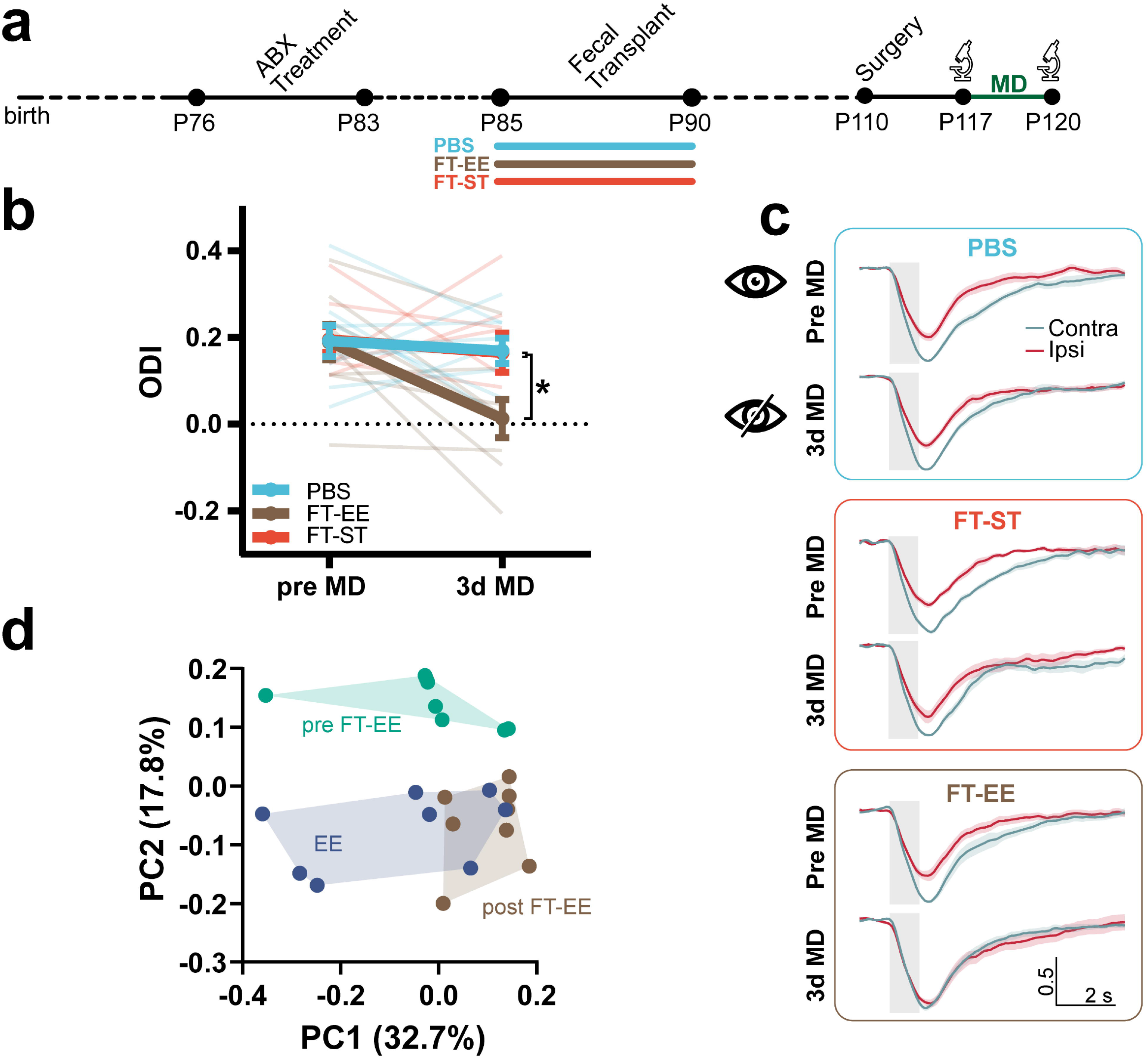
Faecal transplant ODI. (**a**) Experimental timeline. (**b**) ODI before and after monocular deprivation. Thin lines represent single animals (N=9 PBS, N=9 FT-EE, N=8 FT-ST, two-way RM ANOVA time*treatment, interaction F_2,23_=2.888 p=0.076, post-hoc Holm-Sidak, PBS vs FT-EE (3dMD) t_46_=2.92 p=0.029); PBS vs FT-ST (3dMD) t46=0.094 p>0.99; FT-EE vs FT-ST (3dMD) t46=2.773 p=0.039 (**c**) Normalized response timeline (mean±SEM) to contralateral (contra) and ipsilateral (ipsi) eye stimulation before and after MD for all groups. Shaded gray area represents visual stimulus duration. (**d**) PCoA (unweighted UniFrac distance) of EE, preFT-EE and postFT-EE animals. (N=8 per group, ANOSIM test, preFT-EE vs EE, R=0.32, p=0.003; preFT-EE vs postFT-EE, R=0.40, p=0.001; EE vs postFT-EE R=0.17, p=0.03).

Importantly, PCoA of unweighted UniFrac distance showed that the microbiota of ST recipients after the FT with EE feces (postFT-EE) was overlapping the microbiota of EE donor mice, while the microbiota of ST animals before FT (preFT-EE) clustered in a separate group with respect to EE donors and Post-FT (Fig. 6d). Although still significantly different, the ANOSIM R was only 0.17 in the EE donors vs postFT-EE comparison, indicating the presence of a substantial overlap in the phylogenetic composition of ST recipients and EE donors microbiota (Fig. 6d, Suppl. Table 5 for the relative bacteria abundance). No intergroup differences between PreFT-EE, postFT-EE and EE donors in microbial alpha-diversity were found (Suppl. Fig. 11a-b).

All together, those data demonstrate the efficacy of our FT protocol, and indicate that the “plastic phenotype” of the adult EE mouse can be transferred to a ST animal through the fecal microbiota.

## DISCUSSION

Since the pioneeristic studies of Rosenzweig and his collaborators, EE has been widely used as a model for studies on experience-dependent regulation of brain function. Those studies demonstrated that EE dramatically affects brain morphology, chemistry, and physiology; eliciting remarkable plastic responses ranging from molecular to anatomical and functional changes (Rosenzweig and Bennett, 1996). Several investigations continued this line of research trying to understand the specific mechanisms through which EE affects brain plasticity to realize the EE translational potential. However, the mechanisms to explain the EE impact on brain circuits and behaviour have been classically explored into the brain itself: among many others, studies identified changes in neurotrophin levels (Pham et al., 2002), altered expression of plasticity-related genes (Rampon et al., 2000) and synaptic proteins (Nithianantharajah et al., 2004; Song et al., 2018), inhibitory circuits remodelling, perineuronal nets modifications (Sale et al., 2007; Slaker et al., 2016), and epigenetic mark changes (Fischer et al., 2007; Wang et al., 2013). Our study shifts the focus from the CNS to the periphery revealing that signals coming from the gut microbiota contribute to EE-driven cortical plasticity.

### The gut microbiota is modulated by EE

Our study identifies two powerful regulators of the mouse gut microbiota: age and environment. Indeed, we found that fecal bacterial diversity (microbiota alpha-diversity) increases between P20 and P90 in both EE and ST mice. This increase in the complexity of the ecosystem was a general feature process of gut microbiota maturation, not linked to the rearing environment. Significant differences in the phylogenetic composition (beta-diversity) of the bacteria community at distinct ages were observed in ST and EE animals. Notably, EE dramatically altered the fecal microbiota composition. It has to be underscored that this regulation is specific to EE and it is independent from the dietary regimen or other rearing conditions since the ST and EE cages were housed one next to the other and mice received the same food. The number of species differing between ST and EE mice was relatively small, but still significant, before and a few days after weaning. In particular, the LEfSe analysis at P20 revealed that species in the Mollicutes class, *Anearoplasmataceae* family and Anearoplasmatales order were significantly more abundant in EE with respect to ST. At P25 species belonging to the same taxa were still significantly high in EE with respect to ST, in addition to species in the Erysipelotrichia class. However, the most striking difference was present in adulthood, when species belonging to the phylum Tenericutes, and Bacteroidetes were significantly enriched in EE. Why the effect of EE was so prominent in adulthood, while at juvenile ages the species distinction was lower, has not been explored yet. We could speculate that this developmental regulation of EE effect on microbiota might be caused by the fact that adult animals could fully experience the complexity of the stimulation offered by EE. The gut microbiota could be particularly sensitive to features of the EE including physical activity, exploring new toys, spatial patterns of objects, which are activities preferred by older than little mice. However, we cannot exclude other microbiota remodelers starting to impinge on its composition during early postnatal development. For instance, differences in maternal care from enriched dams might account for the distinct composition observed before weaning. Also, the immune system of EE mice could be influenced to send specific signals to the intestinal microbes by still unknown stimuli, thus shaping the ecosystem toward a precise composition. Finally, as metabolism and the microbiota are deeply intertwined, metabolic cues might also play a role. For instance, physical exercise is an important EE component capable of modulating brain plasticity (van Praag et al., 2000), however, it cannot explain the full OD plasticity enhancement observed in EE rodents (Baroncelli et al., 2012). Some studies demonstrated changes in the microbiota composition and diversity of rodents after physical exercise, and an increase in the Bacteroidetes phylum and the *Lachnospiraceae* family (Evans et al., 2014; Mika et al., 2015).

Changes in bacteria families after diet or physical exercise were also correlated to behavioural alterations. For instance, increases in taxa belonging to the *Lachnospiraceae* family were associated with reduced anxiety in exercised mice (Kang et al., 2014). Thus, enhancement in exercise might, indeed, contribute to the experience-dependent modifications observed in our microbiota data, however other still unidentified cues coming from EE could work as commensal remodelers. Since the gut-brain axis is a bidirectional communication system, we cannot exclude that messages coming from the enriched brain may also shape the intestinal microbes in order to shift their metabolic potential toward specific biochemical pathways to sustain more dynamic and plastic neural circuitries.

### Cellular underpinnings of gut microbiota action on OD plasticity

Strikingly, the depletion of the microbiota in EE mice completely blocked visual cortical plasticity. This effect was present both when animals were raised in EE from birth or only with a 5-week EE experience during adulthood. These experiments suggest that an intact microbiota contributes to the EE-dependent enhancement in cortical plasticity.

The microbiota depletion was performed using a classic approach consisting in the administration of a wide spectrum antibiotic mix used in several gut-microbiota-brain axis studies (Chu et al., 2019; Desbonnet et al., 2015; Gacias et al., 2016; Olson et al., 2018). Importantly, microbiome depletion induced by 5-week ABX administration also affected two important EE-induced cellular mechanisms which have been previously involved in plasticity. Indeed, ABX-treated animals did not show increased dendritic spine dynamics and the microglial morphology typical of EE mice (Ali et al., 2019; Xu et al., 2016).

Microglia colonizes the brain at the embryonic stage of development, and the intestinal microbiota has been shown to be a major determinant of microglia maturation, differentiation and function (Erny et al., 2015). Microglia cells are essential players in neuropathology, although recent studies have highlighted their possible roles in brain physiology and plasticity (Wu et al., 2015). Various reports demonstrated that EE housing resulted in more numerous and longer microglial processes in different brain regions (Ali et al., 2019; Xu et al., 2016), demonstrating that microglia morphology in EE is more ramified with respect to standard conditions. Moreover, EE promotion of hippocampal neurogenesis is accompanied by microglia proliferation, and it is impaired in immune deficient mice (Ziv et al., 2006). We found that ABX supplementation significantly impacted on microglia morphology in the visual cortex, strongly decreasing its arborization and filament length. During critical periods, short MD makes microglial cells hyper-ramified, suggesting this shape-signature could be directly connected to OD plasticity (Sipe et al., 2016).

Notably, microglia has been suggested to phagocytose pre- and postsynaptic elements participating in synapse remodelling during development and plasticity (Cheadle et al., 2020; Hong et al., 2016; Paolicelli et al., 2011; Schafer et al., 2012; Schecter et al., 2017). Here, we observed that ABX-depletion of a functional microbiota not only impacted microglia morphology, but also dendritic spine dynamics in the visual cortex of adult EE rodents. Recent studies demonstrated that EE is able to increase dendritic spine dynamics in the cerebral cortex (Jung and Herms, 2014). Furthermore, spine remodelling has been shown to be a structural feature of plasticity in the visual system (Bochner et al., 2014; Chakravarthy et al., 2006; El-Boustani et al., 2018; Hofer et al., 2009; Mataga et al., 2004; Oray et al., 2004; Sajo et al., 2016; Sun et al., 2019; Vidal et al., 2016; Villa et al., 2016) and other sensory modalities (Holtmaat and Svoboda, 2009), suggesting that experience actively sculpts the synaptic landscape to modulate neural circuit function. Our data demonstrated that EE significantly enhanced spine density, through an increase in the growth of new spines. Remarkably, ABX administration counteracted EE-dependent structural plasticity in the visual cortex, as ABX inhibited EE enhancement of both spine density and growth, while spine removal was virtually unaltered. Interfering with the microbiota in EE was essentially translated into hampering the biological processes underlying new spine formation.

### Searching a causal link between microbiota and cortical plasticity: SCFA administration and the faecal transplant

The gut microbiota could communicate with the brain influencing neural function through a variety of communication routes. Among them, our commensals contribute and complement host metabolism by modulating metabolic reactions and producing specific substrates such as SCFA. SCFA act locally in the intestine and also participate in the maintenance of the host metabolic homeostasis by enhancing nutrient absorptions, and curbing the glycemic response (Alexander et al., 2019). Intriguingly, SCFA can be released in the systemic circulation, cross the BBB and reach the brain, where they have been demonstrated to be fundamental for BBB integrity (Braniste et al., 2014), microglia maturation and homeostasis (Erny et al., 2015). Moreover, sodium butyrate decreases microglial activation and pro-inflammatory cytokine secretion (Patnala et al., 2017; Yamawaki et al., 2018) and promotes visual cortical plasticity (Silingardi et al., 2010). Interestingly, we found that living in EE increases the production of SCFA. Furthermore, treating ST adult mice with oral SCFAs for 4 weeks mimicked the effects of FT and EE promoting OD plasticity. As SCFA are highly versatile molecules, it is extremely difficult to envision a molecular mechanism for their effects on neural circuits, and the complexity and crosstalk among SCFA pathways acting on the brain further complicate the picture. SCFA might affect microglia surveillance activity, neural tissue epigenetic landscape and thus gene expression, neurotransmitter levels, or SCFA might act through indirect modulation of endocrine and immune pathways (Dalile et al., 2019). Here, we observed that SCFA treatment was associated with changes in microglia morphology in the visual cortex, rendering it hyper-ramified with respect to ST control. This shape has been observed in conditions of high plasticity such as EE and could favour OD plasticity through still unknown signals. Therefore, we are tempted to speculate that the microbiota could activate cortical plasticity mechanisms through SCFA driven microglia remodelling. Future studies will disentangle this issue, hopefully helping in the identification of specific mechanisms which could be targeted by new microbiota-based strategies for neurological diseases.

Transferring a specific phenotype by transplantation of the intestinal microbes is an important proof to causally link the microbiota to physiology and behaviour. Strikingly, the transplantation of the fecal microbiota of EE mice to adult ST animals clearly induced OD plasticity. This is the first indication that the intestinal microbiota is involved in sensory system plasticity, a developmentally regulated process of plasticity. This observation suggests that EE-derived microbiota might be a powerful tool to trigger juvenile forms of plasticity facilitating the recovery from amblyopia or other serious conditions such as traumatic brain injury, stroke, and post-traumatic stress disorder, leading to new therapeutic strategies for neurodegenerative and neurodevelopmental diseases, or possibly enhancing cognitive performance in healthy subjects. Also, our data prove that signals coming from the intestine overran the stability of the adult sensory circuits and the plasticity brakes.

In summary these findings introduce a novel and paramount concept: experience-dependent changes in the gut microbiota composition can modulate brain circuit function and plasticity. This is a new idea of an “experience-gut microbiota-brain” link: experience does not only impact the brain directly, but also through a previously ignored pathway involving signals coming from the body periphery.

## Author contribution

LL and SC acquired, analyzed, interpreted the data, and critically revised the manuscript, GS acquired and analyzed the data, RM analyzed and interpreted the data, EB and EO performed the SCFA analysis and interpreted the data. PT performed the experiment and conceived the project. TP and PT drafted the manuscript and supervised the project.

## Acknowledgments

We thank Manuel Tongiani and Martina Nasisi for their help with the experiments, and Alexia Tiberi for help with the Imaris Software. Special thanks to Elena Putignano (Institute of Neuroscience, CNR), Vania Liverani and Antonella Calvello (Scuola Normale Superiore) for technical assistance in the lab; Dr. Silvia Burchielli, Cecilia Ciampi and Sara Ciampi (Centre for Experimental Biomedicine, CNR) for assistance in the vivarium; Dr. Maria Cristina Casiraghi, Elisa Adele Colombo and Giovanni Fiorillo (Università degli Studi di Milano) for technical assistance for SCFA measurements. We thank Prof. Concetta Morrone and Prof. Paola Binda (University of Pisa) for insightful comments.

## Funding

This research was supported by H2020-MSCA-IF-2016 749697 GaMePLAY and PRIN2017 2017HMH8FA.

## Conflicts of Interest

The authors declare no conflict of interest.

## MATERIALS AND METHODS

### Animals and housing

All experiments were carried out in accordance with the European Directives (2010/63/EU), and were approved by the Italian Ministry of Health (authorization number 140/2018-PR).

Male and female C57BL/6J mice were used in this study. ST mice were housed in conventional cages (365 × 207 × 140 mm, 2-3 animals per cage) with nesting material. Enriched mice were housed in larger cages (480 × 375 × 210 mm), in larger groups (5-6 mice per cage). Enriched cages were equipped with a variety of sterile toys of different shapes: running wheels, tunnels, climbing devices, food dispensers in different locations and nesting materials. The toys were substituted once a week to ensure the novelty of the environment. The ST and EE cages used in all the experiments were individually ventilated cages to safely maintain the microbiota composition. Mice were kept under a 12 hour dark: 12 hour light cycle, with food (standard diet) and water ad libitum.

A group of mice were raised in EE from birth. The dams of EE-pups were transferred from ST cages to EE cages 6-7 days before delivery. The large spectrum antibiotic cocktail (ABX, vancomycin 0.5 g/l, metronidazole 1 g/l, ampicillin 1 g/l and neomycin 1 g/l) was added to the dam’s water when they were transferred in the EE cages. The litters continued to drink ABX after weaning (postnatal day (P) 21) until the day of ocular dominance plasticity assessment.

A second group of mice lived in EE only for 5 weeks during adulthood, from P85 to P120, and was exposed to the ABX in drinking water only during those 5 weeks.

To study dendritic spine dynamics, Thy-1 GFP transgenic mice (line M (Feng et al., 2000)) were housed in EE cages from P90 for 5 weeks. The control group had ad libitum access to water, while the treated group had ad libitum access to the ABX in drinking water for 5 weeks. For all the experiments, the ABX was freshly prepared and changed every 2 days.

For the short chain fatty acids (SCFAs) treatment a mix of 25 mM sodium propionate, 40 mM sodium butyrate and 67.5 mM sodium acetate (Sigma-Aldrich) was added to drinking water as previously described (Erny et al., 2015) for 4 weeks from P90. The solution was freshly prepared and changed every 2 days. Regular water was administered to control mice.

To avoid cage-effects on our experiments, the animals used in all the experimental groups came from different EE or ST cages.

### Faecal DNA extraction and 16S rRNA sequencing

To analyze the composition of the microbiota of ST and EE mice at different ages, fresh faeces were collected longitudinally in the same subject at P20, P25 and P90, snap-frozen in liquid nitrogen and stored at -80 °C. To avoid the cage-effect on microbiota composition, the animals used for the analysis belonged to different cages. Bacterial DNA was extracted using a specific kit (QIAamp Powerfecal DNA kit, Qiagen), and its concentration was quantified by Nanodrop 2000 C Spectrophotometer (ThermoFisher Scientific).

The 16s rRNA sequencing and analysis was performed by a service offered by Zymo Research (Irvine, CA, USA).

#### Targeted Library Preparation

The DNA samples were prepared for targeted sequencing with the Quick-16S^™^ NGS Library Prep Kit (Zymo Research). The primer sets used were Quick-16S^™^ Primer Set V3-V4 (Zymo Research). The sequencing library was prepared using an innovative library preparation process in which PCR reactions were performed in real-time PCR machines to control cycles and therefore limit PCR chimera formation. The final PCR products were quantified with qPCR fluorescence readings and pooled together based on equal molarity. The final pooled library was cleaned up with the Select-a-Size DNA Clean & Concentrator^™^, then quantified with TapeStation^®^ (Agilent Technologies, Santa Clara, CA) and Qubit^®^ (Thermo Fisher Scientific, Waltham, WA).

#### Control Samples

The ZymoBIOMICS^®^ Microbial Community Standard (Zymo Research) was used as a positive control for each DNA extraction, if performed. The ZymoBIOMICS^®^ Microbial Community DNA Standard (Zymo Research) was used as a positive control for each targeted library preparation. Negative controls (i.e. blank extraction control, blank library preparation control) were included to assess the level of bioburden carried by the wet-lab process.

#### Sequencing

The final library was sequenced on Illumina^®^ MiSeq^™^ with a v3 reagent kit (600 cycles). The sequencing was performed with >10% PhiX spike-in.

#### Bioinformatics Analysis

Unique amplicon sequences were inferred from raw reads using the DADA2 pipeline (Callahan et al., 2016). Chimeric sequences were also removed with the DADA2 pipeline. Taxonomy assignment was performed using Uclust from Qiime v.1.9.1 with the Zymo Research Database, a 16S database that is internally designed and curated, as reference. Composition visualization, alpha-diversity, and beta-diversity analyses were performed with Qiime v.1.9.1 (Caporaso et al., 2010). If applicable, taxonomy that have significant abundance among different groups were identified by LEfSe using default settings (Segata et al., 2011).

### SCFA quantification

Fecal short chain fatty acids (SCFAs) quantification was performed by gas chromatography.

Concentrations of acetic, propionic, butyric acids were assessed as previously described (Borgo et al., 2017). We analyzed female mice. For each animal, 1 to 3 fecal pellets were collected and stored at -80°C until testing. Before experiment, feces were weighted and suspended in 1 ml of double distilled water before extraction with ethyl ether-hexane (1:1 v/v). The aqueous phase was frozen at -80°C and the organic layer was collected for the analysis by a Varian 3400 CX gas liquid chromatograph equipped with a Varian 8200 CX autosampler and a HP-FFAP fused-silica capillary column (30 m, 0.53 mm i.d. with a 1-mm film). Quantification of the SCFAs was obtained through calibration curves of acetic, propionic, and butyric acid in concentrations between 0.25 and 10 mM (10 mM 2-ethylbutyric acid as internal standard).

SCFA concentrations ([SCFA]_sample_) are expressed as mg/g feces and was obtained by applying the following formula:

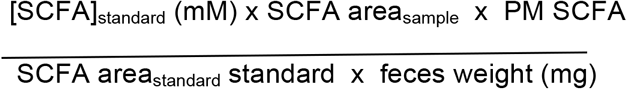

### Faecal transplantation

Four different groups of adult (~P120) EE C57BL/6J mice, living in different enrichment cages, or 4 different cages of age matched ST mice, were used as donors for this experiment. Recipient mice were adult C57BL/6J conventionally raised in ST condition, and housed 2/3 animals per cage. Before starting the faecal transplantation, the recipient mice were subjected to 1-week ABX in drinking water. After 2 days of washout, faecal transplantation was performed through oral gavage for 5 times, once/day (Murakami et al., 2016). To avoid stress for the oral gavage, the procedure was performed for three consecutive days, the animals rested for 2 days, and finally were subjected to the gavage for another 2 days. At the time of faecal transplantation, freshly harvested EE or ST donor faeces were suspended in sterile PBS and mixed with a vortex for 10 min. The suspension was filtered with a cell strainer to remove large debris and immediately used for the transplantation. Every recipient mouse received 200 μl of suspension volume. A group of mice were orally gavaged with sterile PBS (the vehicle of the faeces suspension). After the inoculation, mice were left in their home cage to wait for the engraftment of new bacteria species for 4 weeks. Fresh faecal pellets from EE recipient mice were collected before the ABX treatment and 4 weeks after the transplantation. The animals analysed were housed in different cages to avoid the cage-effect on microbiota composition. DNA was extracted and 16s rRNA-seq was performed as explained in the previous section “Faecal DNA extraction and 16S rRNA sequencing”.

### Monocular Deprivation

Mice were anesthetized with isoflurane (3% induction; 1% maintenance) and placed on a heated pad maintained at 37 °C. The area surrounding the right eye was cleaned in a centrifugal manner with Povidone-iodine diluted 1:1 in saline using cotton swabs. The eye was covered with a thin layer of a dexamethasone/tobramycin ointment (Tobradex, Alcon Novartis) to prevent inflammation and infection. Eyelids were sutured with 3 or 4 horizontal mattress stitches by using a 6-0 surgical suture. After surgery, the animals were monitored and allowed to recover in a heated box.

### Intrinsic Optical Signal Imaging (IOS)

#### Surgery

Surgery for IOS imaging was performed as described in (Mazziotti et al., 2017). Mice were anesthetized with isoflurane (3% induction; 1% maintenance) and head fixed on a stereotaxic frame using ear bars. Body temperature was monitored using a heating pad and a rectal probe to maintain the animals’ body at 37°C. A subcutaneous injection of lidocaine (2%) was provided to anesthetize the local area and the eyes were protected with a dexamethasone-based ointment (Tobradex, Alcon Novartis). The scalp was removed and the skull cleaned with saline. The skin was secured to the skull using cyanoacrylate and a thin layer of cyanoacrylate was poured over the exposed skull to attach a custom-made metal ring (9 mm internal diameter) centered over the binocular visual cortex. A thin layer of clear nail polish was applied over the area to ameliorate optical access. After surgery, the animals were allowed to recover fully in a heated box and monitored to ensure the absence of any sign of discomfort. Before any other experimental procedure, mice were left to recover for at least 48 hours.

#### Imaging and data analysis

Mice were anesthetized with isoflurane (3% induction; 1% maintenance) and chlorprothixene anesthesia (1.5 mg/kg, i.p.) at P120 and P123 (after 3 days of monocular deprivation) Images were visualized using a custom Leica microscope (Leica Microsystems). Red light illumination was provided by 8 individually addressable LEDs (WS2812) attached to the objective (Leica Z6 APO coupled with a Leica PlanApo 2.0X 10447178) by a custom 3D-printed conical holder. Visual stimuli were generated using Matlab Psychtoolbox and presented on a gamma-corrected 24” monitor (C24F390FHU).

Horizontal sine-wave gratings were presented in the binocular portion of the visual field enclosed in a Gaussian envelope spanning -10 to +10 degrees of azimuth and -5 to +60 (full monitor height) degrees of altitude, with a spatial frequency of 0.03 cycles per degree, mean luminance 20 cd/m^2^ and a contrast of 90%. The stimulus consisted of the abrupt contrast reversal of a grating with a temporal frequency of 4 Hz for 1 second, time-locked with a 12-bit depth acquisition camera (PCO edge 5.5) using a parallel port trigger. The interstimulus time was 13 seconds. Frames were acquired at 30 fps with a resolution of 540 x 640 pixels. The signal was averaged for at least 8 groups of 20 trials, stimulating each eye alternatively to prevent biases due to different time in anesthesia for the contralateral and ipsilateral eyes. The signal was then downsampled in time to 10 fps and in space to 270 x 320 pixels. Fluctuations of reflectance (R) for each pixel were computed as the normalized difference from the average baseline (ΔR/R). For each recording, an image representing the mean evoked response was computed by averaging frames between 0.5 to 2.5 seconds after stimulation. The mean image was then low-pass filtered with a 2D average square spatial filter (7 pixels). To select the binocular portion of the primary visual cortex for further analysis, a region of interest (ROI) was automatically calculated on the mean image of the response of the ipsilateral eye by selecting the pixels in the lowest 30% ΔR/R of the range between the maximal and minimal intensity pixel (Cang et al., 2005). To weaken background fluctuations a manually selected polygonal region of reference (ROR) was subtracted. The ROR was placed where no clear response, blood vessel artifact or irregularities of the skull were observed (Heimel et al., 2007). Mean evoked responses were quantitatively estimated as the average intensity inside the ROI. To measure ocular dominance we used the Ocular Dominance Index (ODI) calculated as 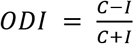 where C and I are the mean response amplitude evoked from contralateral and ipsilateral eye stimulation respectively. For the illustrations in the figures of the signal time course before and after MD, for each animal the contralateral and ipsilateral responses were normalized to the peak minimum value, and low-passed with a moving average (span: 3 samples). Signals were then averaged within each group.

### Two-Photon Imaging

#### Surgery

Cranial windows were implanted on mice around P70. Animals were anesthetized with isoflurane (3% induction; 1% maintenance). Dexamethasone (subcutaneous 0.2 mg/kg) and Lidocaine (2%, 15-20 μL, subcutaneous, scalp area) were administered. After a few minutes, the scalp was cleaned with three swabs of povidone-iodine and a large portion of skin, covering both the hemispheres, was removed. The skull was cleaned from the periosteum by initially using saline and then by carefully scraping it with a scalpel blade. We drew a circular area of 3 mm of diameter centered 3 mm lateral, 1 mm anterior to lambda to mark the craniotomy area and we applied a thin layer of a light-curing dental cement on the rest of the exposed skull (3M Vitrebond^™^ plus). A metal head-plate was fixed to the skull by using more dental cement. By carefully using a dental drill and a biopsy punch, a 3 mm circular groove was thinned until almost transparent on the previously marked area, then a few drops of cold sterile ACSF were applied to the area. A circular island of bone was removed with the tip of a sharp forceps without damaging the dura. We placed and held in place a circular 3 mm coverslip on the craniotomy and secured it to the skull by using dental resin (Lang Contemporary Ortho Jet^™^). We then covered thoroughly the skull and the head-plate with more dental resin to finish the surgery. The animals were allowed to recover in a heated box and monitored to ensure the absence of any sign of discomfort. Before any other experimental procedure, mice were left to recover for 12-16 days to reduce inflammation in the surgical area, thus increasing the optical access to the tissue. A more detailed explanation for surgical procedures can be found in (Holtmaat et al., 2009)

#### Imaging and data analysis

Imaging was performed using a Bruker Ultima Investigator^™^ microscope equipped with a GaSsP Photomultiplier tube and controlled by the scanning software Prairie View. Laser excitation was provided by a tunable Ti:Sapphire pulsating LASER (Chameleon Ultra^™^, Coherent) tuned at 920 nm and excitation power was controlled with a Pockels Cell. LASER power was maintained under a maximum of 40 mW on the sample.

Images were acquired using a 20x long WD water immersion objective (Olympus XLUMPlanFLN 20x N.A. = 1.00). For each mouse, 1-3 pyramidal neurons with soma position in layer V were imaged. For each neuron, in the first session, we acquired an epifluorescence image of the vasculature of the imaging area to relocate the same dendritic segments on subsequent days. We then acquired a low-magnification two-photon stack of the entire apical dendritic arborization of the cell (1024×1024 pixels, x-y resolution: 0.46 μm/px, z step: 3 μm). 1-5 dendritic segments (imaging depth: 0-200 μm from the brain surface) per cell were chosen randomly between those that had better optical clarity, and a high magnification stack of those segments was acquired (1024×1024 pixels, x-y resolution: 0.09 μm/px, z step: 1 μm). We ensured similar fluorescence across imaging sessions and, to prevent phototoxicity, the lowest laser power (< 40 mW) that could resolve all spines was used. Dendritic spines were counted manually with a custom-written MATLAB software by comparing simultaneously single, aligned z planes from the z-stacks of all 8 experimental time points. Images were first low-pass filtered with a 2D Gaussian filter (MATLAB function imgaussfilt, sigma: 1.1). We counted all clear spine protrusions emanating laterally from the dendrite, regardless of their shape (stubby, mushroom, thin). We considered the spines to be the same from one session to the other based on their relative position to structural landmarks and to very clear persistent spines present in all time points. Spines were considered different if they branched out from the dendritic segment more than 1 μm away from their previous position.

Spine Density was defined as the number of spines per μm of length of the dendritic segment. Similarly, the fraction of spines gained (F(gained)) and lost (F(lost)) between two time points t_1_ and t_2_ were defined as follows F_(gained)_=N_(gain)_/N_(t2)_*100 and F_(lost)_=N_(lost)_/N_(t2)_*100 (Holtmaat et al., 2005; Murmu et al., 2013). The time-dependent survival function was defined as SF = N_(t)_/N_(0)_ (Murmu et al., 2013) where N_(0)_ is the number of spines present at the first imaging time point, and N_(t)_ is the number of spines of the original set surviving after time t.

### Immunofluorescence analysis of microglial morphology

Mice were anesthetized with chloral hydrate (20ml/Kg BW) and perfused via intracardiac infusion with PBS and then 4% paraformaldehyde (PFA, w/vol, dissolved in 0.1 M phosphate buffer, pH 7.4). Brains were quickly removed and post-fixed overnight in PFA at 4 °C, then transferred to 30% sucrose (w/vol) solution. 45 μm coronal sections were cut on a freezing microtome (Leica) and free-floating sections were processed for immunofluorescence.

The cortical sections were incubated for 1 h in blocking solution containing 5% BSA (w/vol) and 0.3% Triton X-100 (vol/vol) in PBS, and incubated overnight at 4 °C with anti-Iba-1 (cat. no. 019-19741, Wako) diluted 1:500 in PBS with 1% BSA (w/vol) and 0.1% Triton X-100 (vol/vol).

Sections were then washed with PBS and incubated for 2h at 22-24 °C with Alexa Fluor 488– conjugated secondary antibody (cat. no A32731, Invitrogen), which was added at a dilution of 1:500 in the same solution as the primary antibody.

Sections were washed three times with PBS and mounted on slides, then they were air-dried and coverslipped with Vectashield mounting medium (cat. H-1000,Vector Laboratories).

Imaging was performed on an LSM 900 confocal microscope (Zeiss, Oberkochen, Germany) using a Plan-Apochromat 63x, NA:1.4 oil objective.

The area of the visual cortex was defined based on the mouse brain atlas (Paxinos and Franklin’s the Mouse Brain in Stereotaxic Coordinates).

Z-stacks of ~ 40 μm were acquired with a z-step of 0.50 μm (for a final voxel size of 0.1980717 × 0.1980717 × 0.5 μm). Images were then processed using the Filament Tracer Tool of IMARIS software (Bitplane). Between 5 and 10 cortical cells were reconstructed per analyzed mouse.

### Statistical analysis

The majority of statistical analyses were performed using GraphPad Prism version 7 (GraphPad Software, San Diego, CA, USA).

#### Gut microbiota analysis

to test whether two or more groups of samples were significantly different analysis of similarities (ANOSIM) and principal coordinate analysis were calculated using the Python library scikit-bio (http://scikit-bio.org/). The principal component analysis was performed on OTUs profiles using the Python package scikit-learn (https://scikit-learn.org/stable/).

#### IOS experiments

Differences between groups were tested for significance using two-way RM ANOVA, unless otherwise indicated. Holm–Sidak’s multiple comparisons *post hoc* tests were performed, when appropriate, to correct for multiple hypothesis testing.

#### Dendritic spines analysis

differences in spine density were evaluated using a two-way RM ANOVA time*housing. Holm–Sidak’s multiple comparisons *post hoc* tests were performed, when appropriate, to correct for multiple hypothesis testing. Spine formation and elimination were compared with two-tailed t-test when comparing two housing groups, or with two-way RM ANOVA when comparing more than two groups.

#### Microglia morphology

For the analysis of filament length, number of branching points and dendrite terminals, statistical differences between groups were assessed using unpaired two-tailed Student’s *t* test or Ordinary One-Way ANOVA followed by Tukey’s *post hoc* multiple comparisons test. For the Sholl analysis, a Two-Way ANOVA was applied for main and interaction effects between distance from the soma and treatment, followed by Sidak’s or Tukey’s *post hoc* multiple comparisons test.

All data are represented as the mean±SEM unless otherwise stated. N’s represent single animals unless otherwise stated. In the figures \**P* < 0.05, \*\**P* < 0.01 and \*\*\**P* < 0.001.

## Code availability

Custom MATLAB code for IOS and dendritic spine analysis is available upon request.

## Data availability

16S rRNA-seq data are available at BioProject under accession numbers PRJNA633066.

## Notes

### Competing Interest Statement

The authors have declared no competing interest.

