## Supplementary Figures for "The gut microbiota of environmentally enriched mice regulates visual cortical plasticity"

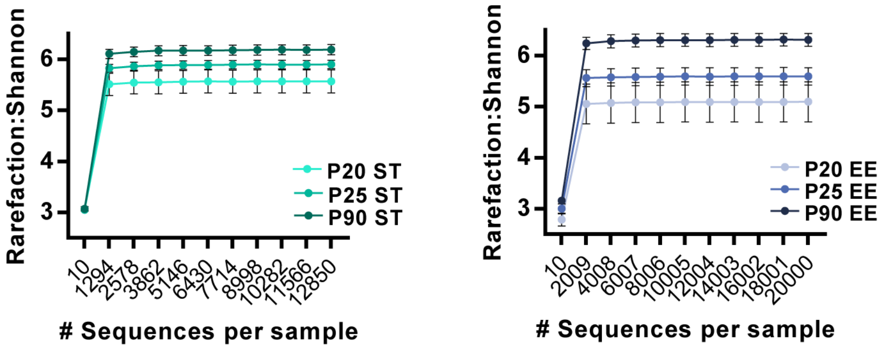


**Suppl. Fig 1**

Shannon alpha-diversity rarefaction curve of 16S rRNA sequencing data from ST (left) and EE birth mice (right) at P20, P25, P90 (P20 ST, N=6; P25 ST, N=6; P90 ST, N=5; P20 EE, N=6; P25 EE N=6; P90 EE N=6). The plots show the number of identified species as a function of sequencing depth. Data are presented as mean ± SEM.

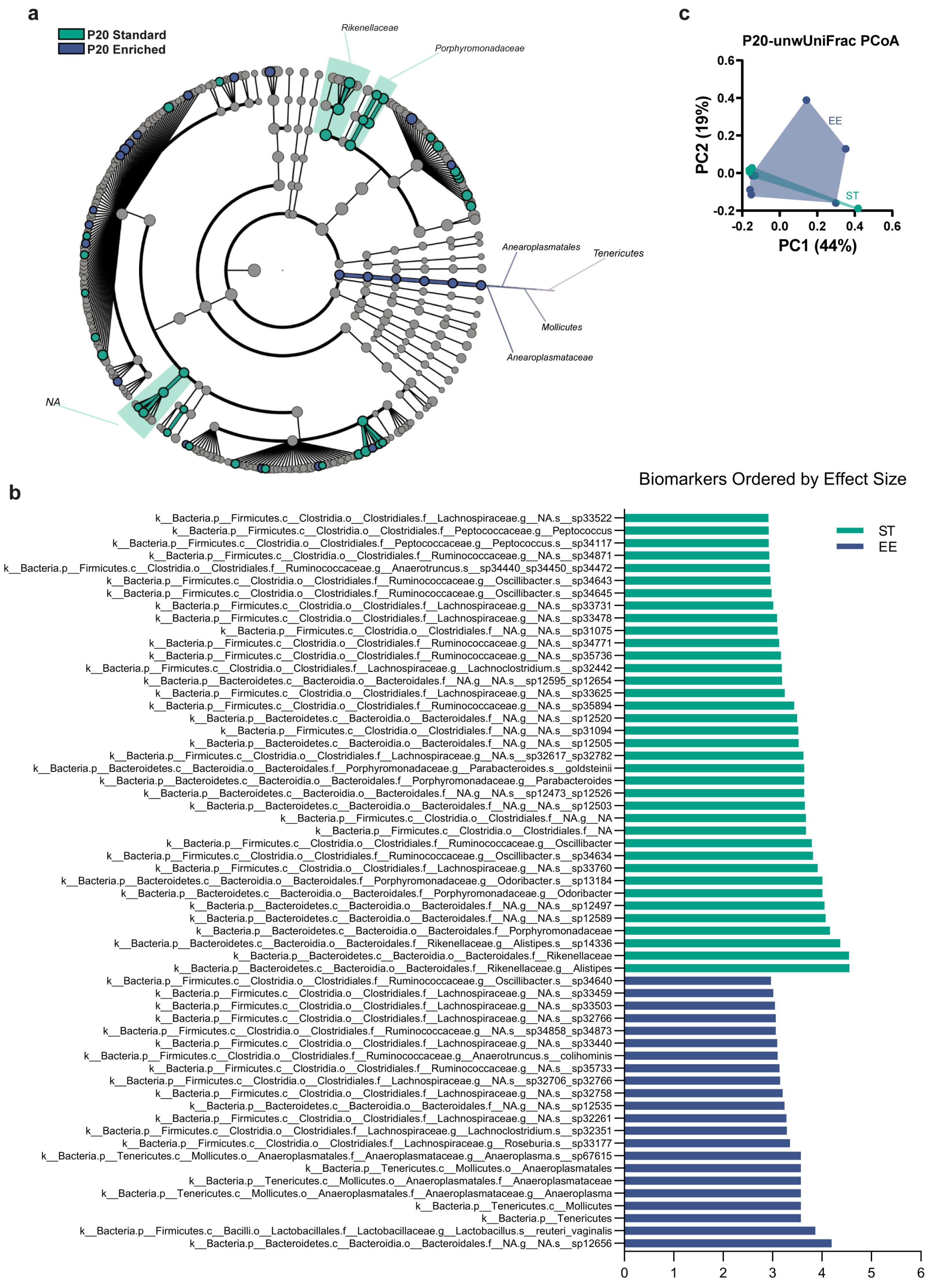


**Suppl. Fig 2**

**a**) Taxonomic representation (cladogram) of statistically different bacteria between ST and EE birth mice at P20. Bacterial classes enriched in each experimental group are represented in a different color (blue=EE, green=ST). Each circle's diameter is proportional to the taxon's abundance. Gray circles mean no differences between the groups (N=6 P20 ST, N=6 P20 EE). **b**) Histogram of the Linear Discriminant Analysis (LDA) scores ranked by effect size of bacteria differentially abundant between ST and EE birth mice at P20. **c**) PCoA between ST and EE birth animals at P20 (N=6 P20 ST, N=6 P20 EE; ANOSIM test, P20 ST vs EE birth, R=0.085 p=0.151).

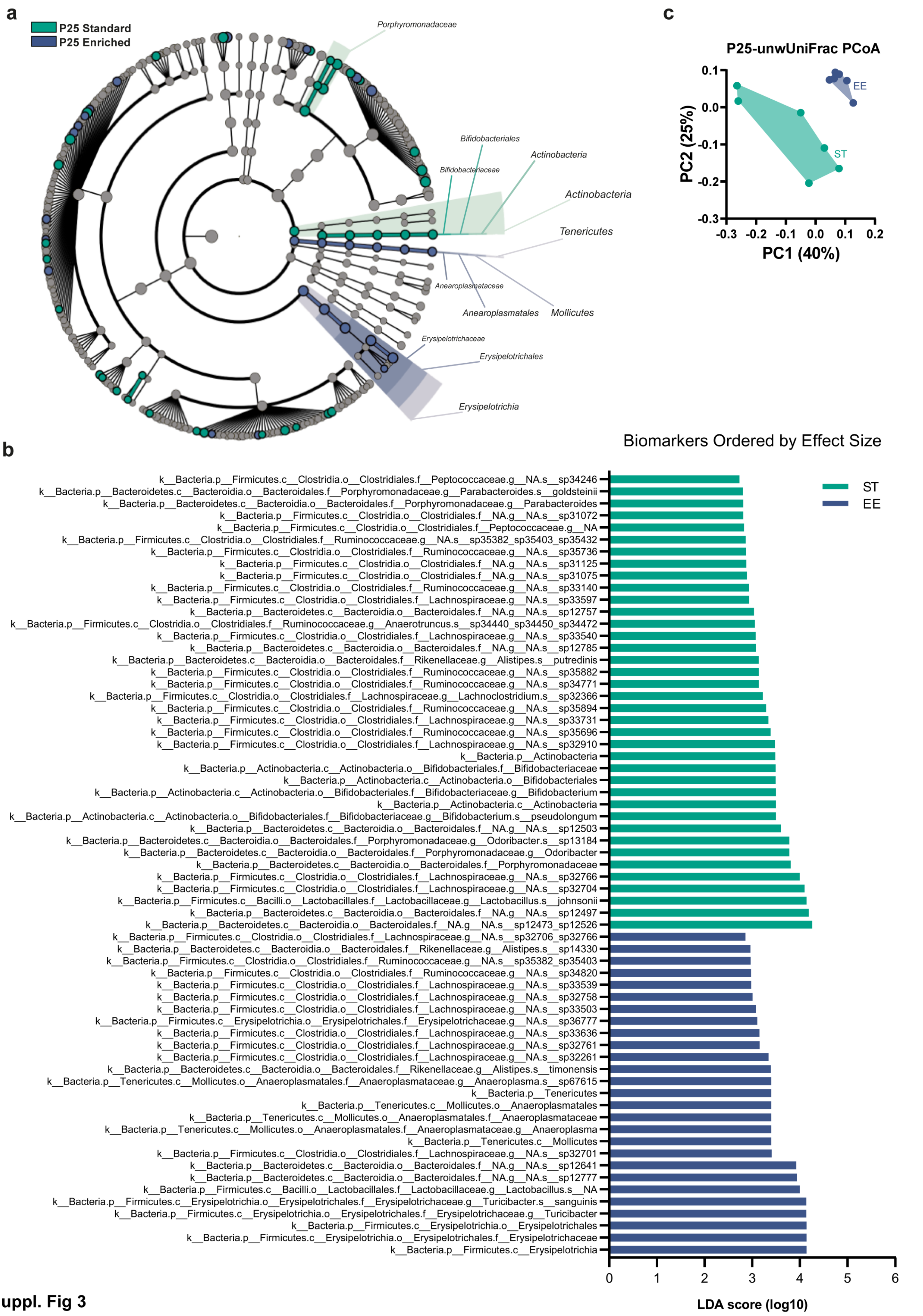


**Suppl. Fig 3**

**a**) Cladogram reporting the taxa (highlighted by small circles and by shading) showing different abundance values between EE birth and ST mice at P25 (blue=EE, green=ST). Each circle's diameter is proportional to the taxon's abundance. Gray circles mean no differences between groups (N=6 P25 ST, N=6 P25 EE). **b**) Histogram of the LDA scores ranked by effect size of bacteria differentially abundant between ST and EE birth mice at P25. **c**) PCoA between ST and EE birth animals at P25 (N=6 P25 ST, N=6 P25 EE; ANOSIM test, P25 ST vs EE birth, R=0.463 p=0.001).

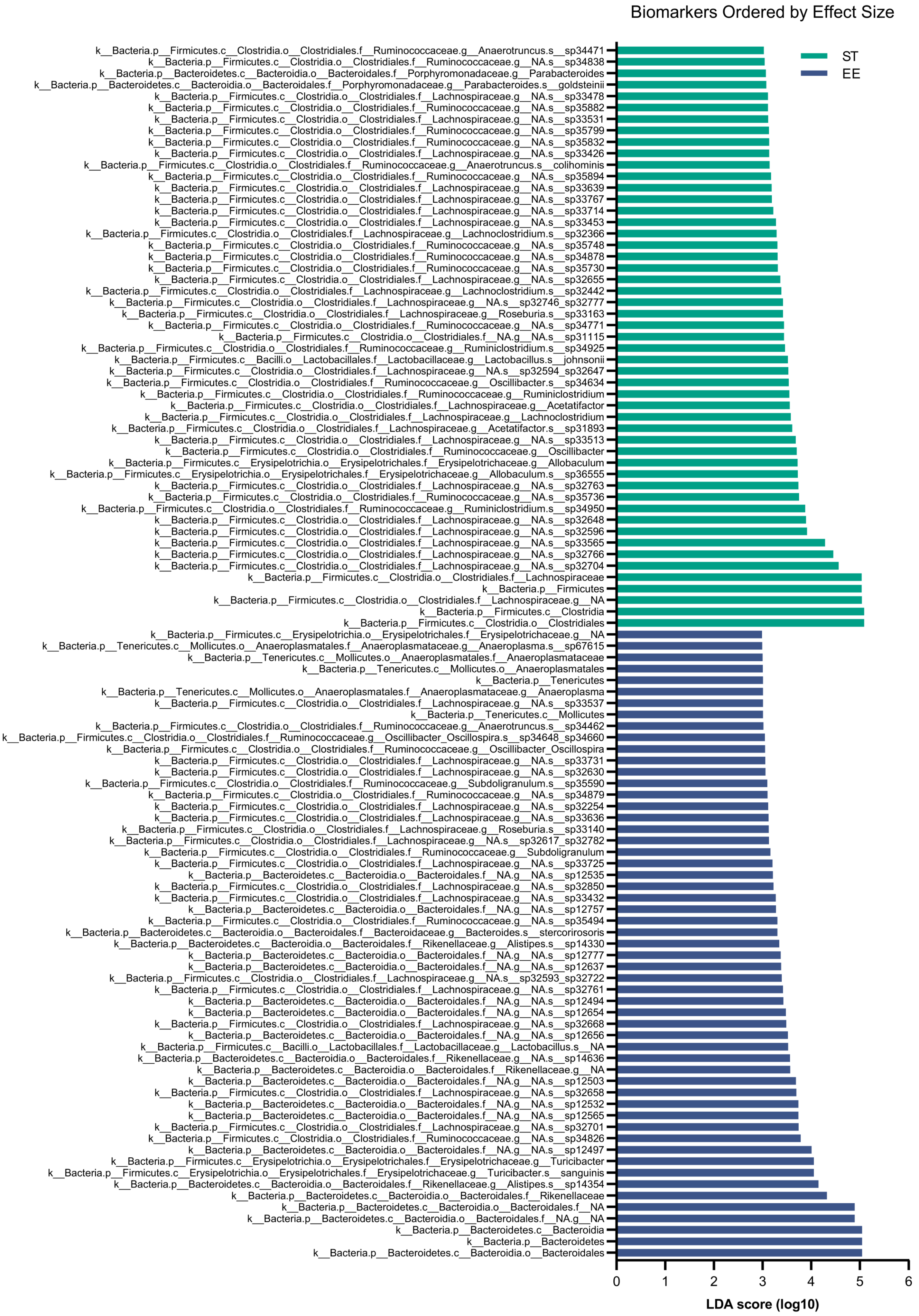


**Suppl. Fig 4**

Histogram of the LDA scores of bacteria differentially abundant between ST and EE birth mice at P90 (N=5 P90 ST, N=6 P90 EE).

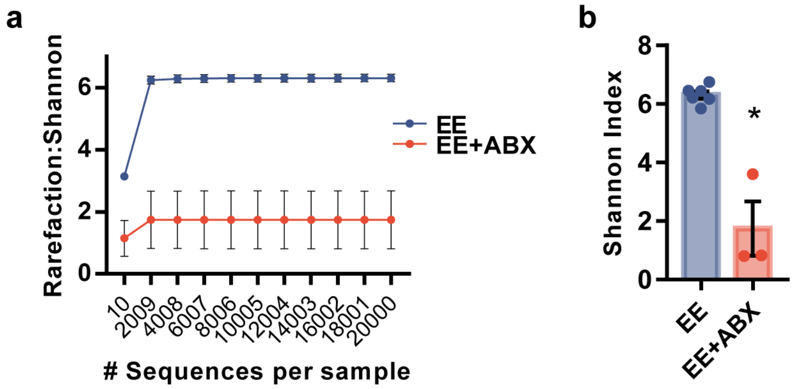


**Suppl. Fig 5**

**a**) Shannon alpha-diversity rarefaction plot of 16S rRNA sequencing data from EE and EE+ABX mice at P90 (N=6 EE, N=3 EE+ABX). **b**) Shannon alpha-diversity scatter plot of 16S rRNA in stool pellets collected from EE and EE+ABX mice at P90 (N=6 EE, N=3 EE+ABX; Mann-Whitney two-tailed test, U=0, p=0.0238). Single animals are represented as dots. Data are presented as mean ± SEM.

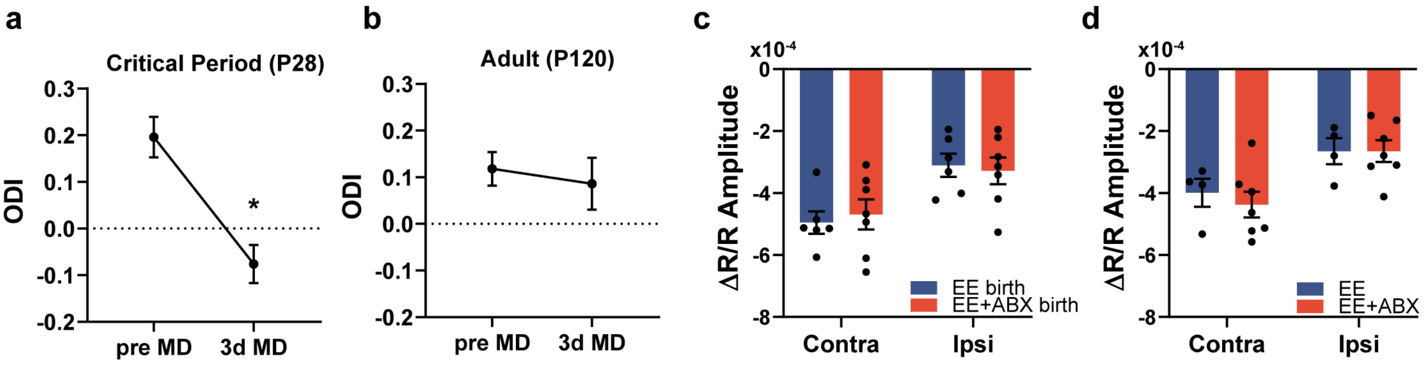


**Suppl. Fig 6**

**a**) Ocular Dominance Index (ODI) before and after monocular deprivation in young (P28, critical period) mice (N=4 two-tailed paired t-test $t_3$=3.25 p=0.047). **b**) Ocular Dominance Index (ODI) before and after monocular deprivation in adult (P120) mice (N=5 two-tailed paired t-test $t_4$=0.36 p=0.74). Data in (**b**) derive from the ST group shown in figure 6. **c**) Intrinsic signal response amplitude to contralateral and ipsilateral eye stimulation in EE birth and EE+ABXbirth animals before MD (N=6 EE-birth, N=7 EE+ABX-birth; two-way RM ANOVA eye*housing, interaction $F_{1,11}$=1.322 p=0.27, eye $F_{1,11}$=70.96 p<0.0001, housing $F_{1,11}$=0.005 p=0.95). (**c**) Intrinsic signal response amplitude to contralateral and ipsilateral eye stimulation in EE and EE+ABX animals before MD (N=6 EE-birth, N=7 EE+ABX-birth; two-way RM ANOVA eye*housing, interaction $F_{1,9}$=0.81 p=0.39, eye $F_{1,9}$=51.57 p<0.0001, housing $F_{1,9}$=0.11 p=0.75). Data are presented as mean ± SEM.

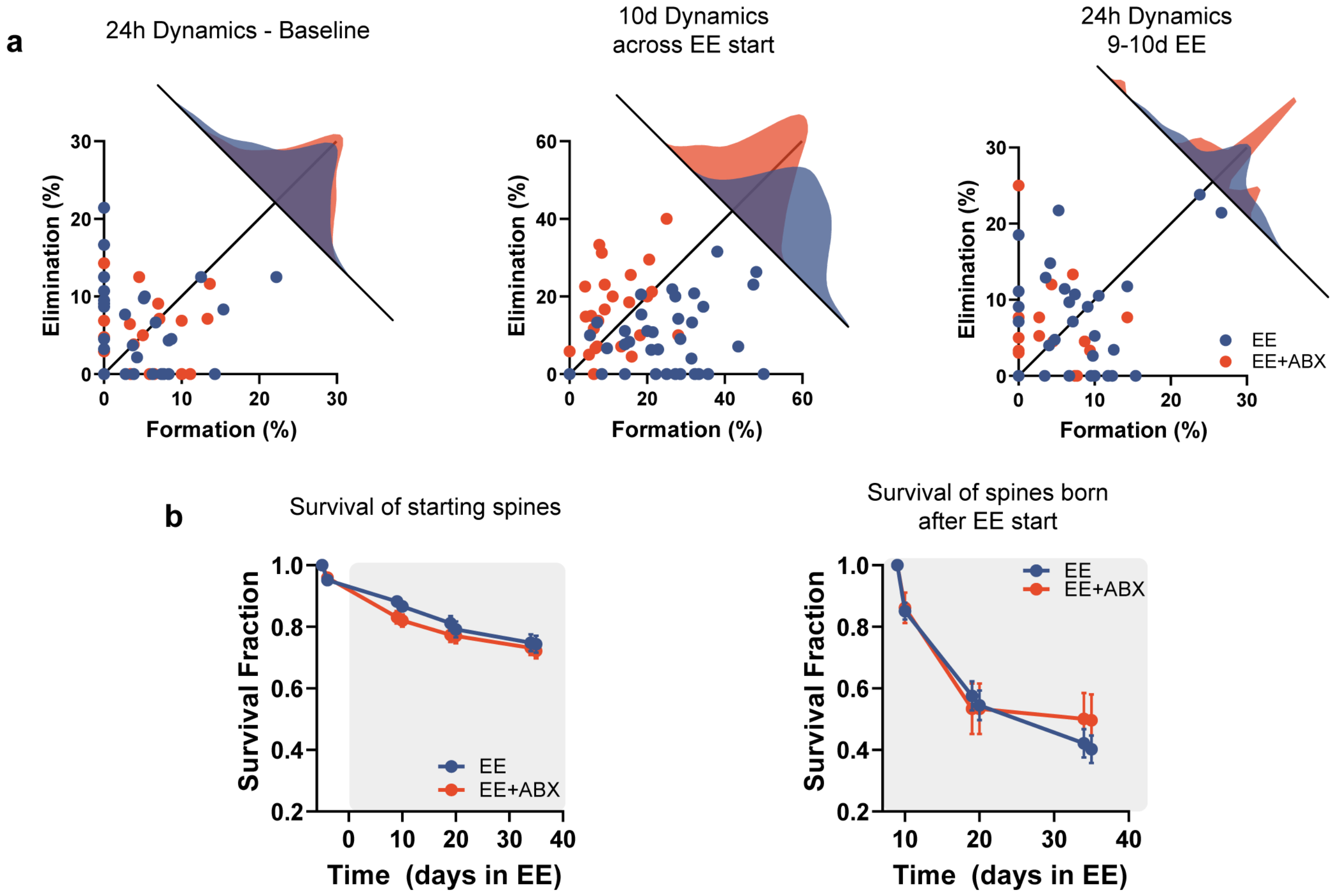


**Suppl. Fig 7**

**a**) Each dendrite spine elimination plotted as a function of spine formation for the time points analyzed in figure 3i-l. **b**) Left: spine survival curve of the spines present at the first time point (N=39 EE, N=28 EE+ABX; two-way RM ANOVA time*treatment, interaction: $F_{7,455}$=1.54 p=0.15, time: $F_{1.739,113.0}$=133.5 p<0.001, treatment: $F_{1,65}$=0.86 p=0.36). Right: spine survival curve of the spines born between -4d and 9d timepoint (N=39 EE, N=26 EE+ABX; two-way RM ANOVA time*treatment, interaction: $F_{5,310}$=1.27 p=0.28 time: $F_{1.989,123.3}$=93.57 p<0.001, treatment: $F_{1,62}$=0.12 p=0.73). Data are presented as mean ± SEM.

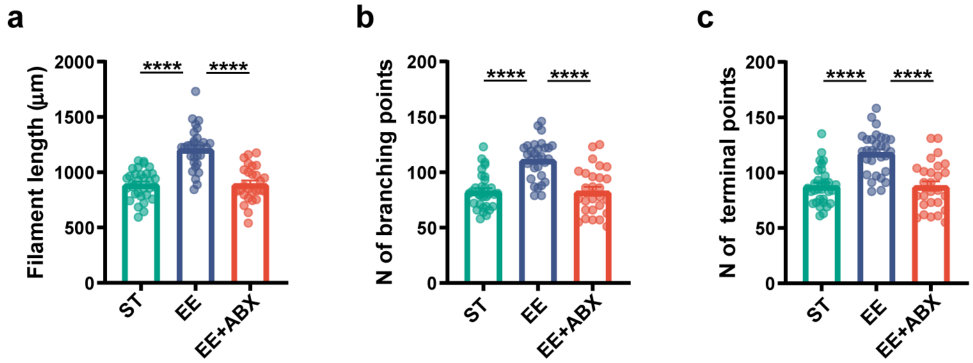


**Suppl. Fig 8**

IMARIS quantitative morphometric analysis of IBA-1+ microglia of ST mice, mice housed for 5 weeks in EE conditions with ABX treatment or in EE alone. **a**) Filament length (Ordinary one-way ANOVA, $F_{2,86}$=38.88 p<0.0001, Tukey's post-hoc ST vs EE p<0.0001, EE vs EE+ABX p<0.0001), **b**) number of branching points (Ordinary one-way ANOVA, $F_{2,86}$=24.75 p<0.0001, Tukey's post-hoc ST vs EE p<0.0001, EE vs EE+ABX p<0.0001) and **c**) number of terminal points (Ordinary one-way ANOVA, $F_{2,86}$=24.38 p<0.0001, Tukey's post-hoc ST vs EE p<0.0001, EE vs EE+ABX p<0.0001). Single cells are shown as dots (N=32 ST, N=30 EE, N=27 EE+ABX). Data are presented as mean ± SEM.

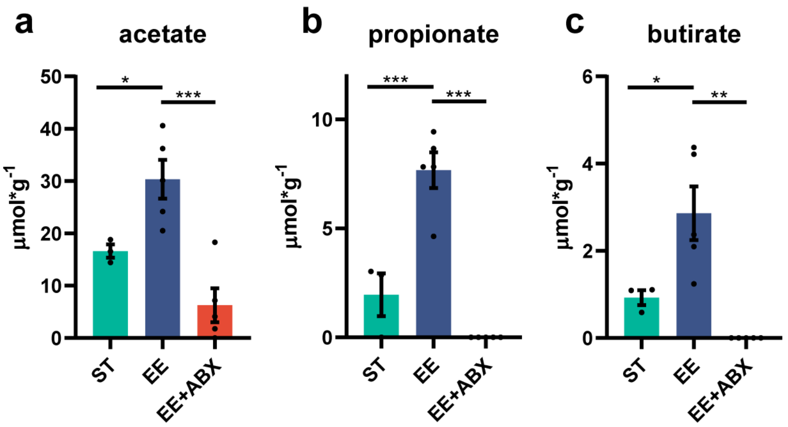


**Suppl. Fig 9**

Mass spectrometry analysis of SCFA components from feces samples collected from standard reared mice (ST), mice reared in Environmental Enrichment (EE) and mice raised in EE and treated with an antibiotics cocktail (EE+ABX) for five weeks. **a**) Acetate concentration in the different groups (one-way ANOVA p=0.001, $F_{2,10}$=14.70, post-hoc Holm-Sidak, ST vs EE $t_{3,5}$=2.67, p=0.046; ST vs EE+ABX $t_{3,5}$=2.01, p=0.072; EE vs EE+ABX $t_{3,5}$=5.41, p<0.001). **b**) Propionate concentration in the different groups (one-way ANOVA p<0.001, $F_{2,10}$=40.60, post-hoc Holm-Sidak, ST vs EE $t_{3,5}$=5.658, p<0.001; ST vs EE+ABX $t_{3,5}$=1.938, p=0.081; EE vs EE+ABX $t_{3,5}$=8.77, p<0.001). **c**) Butyrate concentration in the different groups (one-way ANOVA p<0.001, $F_{2,10}$=40.60, post-hoc Holm-Sidak, ST vs EE $t_{3,5}$=5.658, p<0.001; ST vs EE+ABX $t_{3,5}$=1.938, p=0.081; EE vs EE+ABX $t_{3,5}$=8.77, p<0.001).

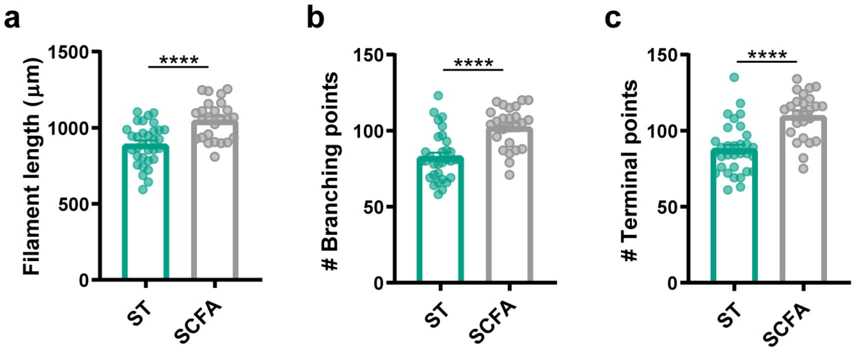


**Suppl. Fig 10**

IMARIS quantitative morphometric analysis of IBA-1+ microglia of mice housed in ST condition drinking regular water (ST) or housed in ST condition and drinking SCFA solution for 4 weeks (SCFA). **a**) Filament length (unpaired two-tailed t-test, $t_{55}$=4.550 p<0.0001), **b**) number of branching points (unpaired two-tailed t-test, $t_{55}$=4.938 p<0.0001) and **c**) number of terminal points (unpaired two-tailed t-test, $t_{55}$=5.149 p<0.0001). Single cells are shown as dots (N=32 ST, N=25 SCFA). Data are presented as mean ± SEM.

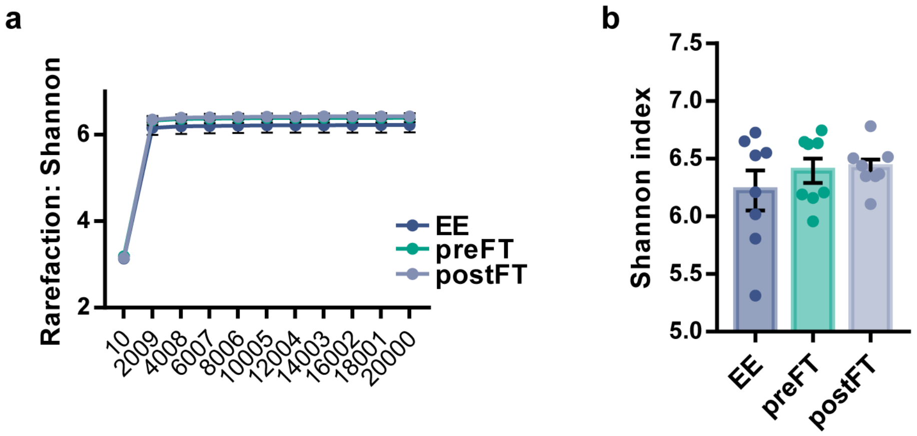


**Suppl. Fig 11**

**a**) Shannon alpha-diversity rarefaction plot and **b**) bar plot of 16S rRNA sequencing data from preFT, postFT and EE donors (N=8 mice per group, Kruskal-Wallis test, statistic=0.24, p=0.8869). Single animals are represented as dots. Data are presented as mean ± SEM.

### Suppl. Table 1

Relative abundance of bacterial species identified in P20, P25, P90 mice reared in standard (ST) condition.

### Suppl. Table 2

Relative abundance of bacterial species identified in P20, P25, P90 mice reared in enriched (EE) conditions from birth.

### Suppl. Table 3

Tukey's post-hoc comparisons table of microglial morphology (Sholl analysis; ST vs EE vs EE+ABX, Fig. 4b)

### Suppl. Table 4

Sidak's post-hoc comparisons table of microglial morphology (Sholl analysis; ST vs SCFA, Fig. 5e)

### Suppl. Table 5

Relative abundance of bacterial species identified in preFT, postFT, EE donors mice.
