## Supplementary material for "The gut microbiota of environmentally enriched mice regulates visual cortical plasticity": Suppl. Table 1

| SampleID | P20.ST1 | P20.ST2 | P20.ST3 | P20.ST4 | P20.ST5 | P20.ST6 | P25.ST1 | P25.ST2 | P25.ST3 | P25.ST4 | P25.ST5 | P25.ST6 | P90.ST1 | P90.ST2 | P90.ST3 | P90.ST4 | P90.ST5 |
|---|---|---|---|---|---|---|---|---|---|---|---|---|---|---|---|---|---|
| k__Bacteria;p__Actinobacteria;c__Actinobacteria;o__Bifidobacteriales;f__Bifidobacteriaceae;g__Bifidobacterium;s__pseudolongum | 0.000306 | 0.011567 | 0 | 0.001209 | 0 | 0.002804 | 0.003638 | 0.010006 | 0.00834 | 0.005105 | 0.00967 | 0.005101 | 0.002614 | 0.005974 | 0 | 0.011803 | 0.002946 |
| k__Bacteria;p__Actinobacteria;c__Actinobacteria;o__Corynebacteriales;f__Mycobacteriaceae;g__Mycobacterium;s__mucogenicum-phocaicum | 0 | 0 | 0 | 0 | 0 | 0.000407 | 0 | 0 | 0 | 0 | 0 | 0 | 0 | 0 | 0 | 0 | 0 |
| k__Bacteria;p__Actinobacteria;c__Coriobacteriia;o__Coriobacteriales;f__Coriobacteriaceae;g__Enterorhabdus;s__caecimuris | 0 | 0 | 0 | 0 | 0 | 0 | 0 | 0 | 0.000657 | 0 | 0 | 0.000887 | 0 | 0 | 0 | 0.003504 | 0 |
| k__Bacteria;p__Actinobacteria;c__Coriobacteriia;o__Coriobacteriales;f__Coriobacteriaceae;g__Enterorhabdus;s__mucosicola | 0 | 0 | 0 | 0 | 0 | 0 | 0 | 0 | 0 | 0 | 0 | 0 | 0 | 0.001021 | 0 | 0 | 0 |
| k__Bacteria;p__Actinobacteria;c__Coriobacteriia;o__Coriobacteriales;f__Coriobacteriaceae;g__Olsenella;s__profusa | 0 | 0 | 0 | 0 | 0 | 0 | 0 | 0 | 0 | 0 | 0 | 0 | 0 | 0.001842 | 0 | 0.001053 | 0 |
| k__Bacteria;p__Actinobacteria;c__Coriobacteriia;o__Coriobacteriales;f__Coriobacteriaceae;g__Paraeggerthella;s__hongkongensis | 0 | 0 | 0 | 0 | 0 | 0 | 0 | 0 | 0 | 0 | 0 | 0 | 0 | 0 | 0 | 0.000967 | 0 |
| k__Bacteria;p__Bacteroidetes;c__Bacteroidia;o__Bacteroidales;f__Bacteroidaceae;g__Bacteroides;s__acidifaciens | 0.041439 | 0.041442 | 0.005756 | 0.0551 | 0 | 0.046009 | 0.0151 | 0.017071 | 0.017063 | 0.01287 | 0.014535 | 0.013772 | 0.002646 | 0.004082 | 0.008869 | 0.000666 | 0.002127 |
| k__Bacteria;p__Bacteroidetes;c__Bacteroidia;o__Bacteroidales;f__Bacteroidaceae;g__Bacteroides;s__oleiciplenus-rodentium | 0.123143 | 0.054923 | 0.02792 | 0.124971 | 0.048541 | 0.124746 | 0.027236 | 0.018274 | 0.018231 | 0.025993 | 0.017708 | 0.039896 | 0.010777 | 0.015681 | 0.026641 | 0.003053 | 0.011521 |
| k__Bacteria;p__Bacteroidetes;c__Bacteroidia;o__Bacteroidales;f__NA;g__NA;s__NA | 0 | 0 | 0 | 0 | 0 | 0 | 0 | 0 | 0 | 0 | 0 | 0 | 0 | 0 | 0 | 0 | 0.0036 |
| k__Bacteria;p__Bacteroidetes;c__Bacteroidia;o__Bacteroidales;f__NA;g__NA;s__sp12473-sp12526 | 0.015702 | 0.002492 | 0.006863 | 0.011618 | 0.008713 | 0.009422 | 0.017439 | 0.03567 | 0.038013 | 0.039189 | 0.043514 | 0.042779 | 0.001742 | 0.004729 | 0.004892 | 0.004343 | 0 |
| k__Bacteria;p__Bacteroidetes;c__Bacteroidia;o__Bacteroidales;f__NA;g__NA;s__sp12475-sp12557 | 0.114181 | 0.12064 | 0.037317 | 0.122798 | 0.073668 | 0.125228 | 0.042336 | 0.102522 | 0.114313 | 0.108426 | 0.101472 | 0.080746 | 0.02349 | 0.06362 | 0.048742 | 0.036376 | 0.033678 |
| k__Bacteria;p__Bacteroidetes;c__Bacteroidia;o__Bacteroidales;f__NA;g__NA;s__sp12494 | 0.005362 | 0.005073 | 0.001304 | 0.005801 | 0 | 0.00707 | 0.001819 | 0.003437 | 0.004781 | 0.005557 | 0.004563 | 0.007429 | 0.002485 | 0.002987 | 0.004857 | 0.002107 | 0.002389 |
| k__Bacteria;p__Bacteroidetes;c__Bacteroidia;o__Bacteroidales;f__NA;g__NA;s__sp12497 | 0.029566 | 0.035474 | 0.025189 | 0.034429 | 0 | 0.040687 | 0.017153 | 0.035555 | 0.050824 | 0.047262 | 0.056417 | 0.020824 | 0.017101 | 0.018319 | 0.022453 | 0.015651 | 0.024154 |
| k__Bacteria;p__Bacteroidetes;c__Bacteroidia;o__Bacteroidales;f__NA;g__NA;s__sp12499 | 0 | 0 | 0.004649 | 0.004967 | 0 | 0.002698 | 0.001429 | 0.003265 | 0.001788 | 0 | 0 | 0.002905 | 0 | 0.001319 | 0.005701 | 0 | 0 |
| k__Bacteria;p__Bacteroidetes;c__Bacteroidia;o__Bacteroidales;f__NA;g__NA;s__sp12499-sp12616 | 0.00503 | 0.00237 | 0 | 0 | 0 | 0 | 0 | 0 | 0 | 0.00362 | 0 | 0 | 0 | 0 | 0 | 0.002515 | 0 |
| k__Bacteria;p__Bacteroidetes;c__Bacteroidia;o__Bacteroidales;f__NA;g__NA;s__sp12499-sp12616-sp12777 | 0 | 0 | 0 | 0.001373 | 0 | 0.001703 | 0 | 0 | 0.000456 | 0 | 0 | 0 | 0 | 0 | 0 | 0 | 0.002651 |
| k__Bacteria;p__Bacteroidetes;c__Bacteroidia;o__Bacteroidales;f__NA;g__NA;s__sp12502 | 0.014911 | 0.025996 | 0 | 0.015458 | 0.025515 | 0.017547 | 0 | 0.008994 | 0.010165 | 0.008707 | 0 | 0.008294 | 0 | 0.007965 | 0 | 0 | 0 |
| k__Bacteria;p__Bacteroidetes;c__Bacteroidia;o__Bacteroidales;f__NA;g__NA;s__sp12503 | 0.010136 | 0.013568 | 0 | 0.012059 | 0 | 0.014155 | 0.005562 | 0.014016 | 0.017884 | 0.016563 | 0 | 0.007939 | 0 | 0.006895 | 0.007144 | 0.004558 | 0 |
| k__Bacteria;p__Bacteroidetes;c__Bacteroidia;o__Bacteroidales;f__NA;g__NA;s__sp12505 | 0.013507 | 0.010479 | 0.008142 | 0.008383 | 0.015325 | 0.008849 | 0.007563 | 0.007963 | 0.009307 | 0.008851 | 0.013689 | 0.013639 | 0.004517 | 0.004804 | 0.009643 | 0.003182 | 0.003862 |
| k__Bacteria;p__Bacteroidetes;c__Bacteroidia;o__Bacteroidales;f__NA;g__NA;s__sp12520 | 0.011592 | 0.008373 | 0.006322 | 0.01121 | 0.016492 | 0.011397 | 0.003924 | 0.007123 | 0.010512 | 0.010372 | 0.00964 | 0.005234 | 0.008583 | 0.006023 | 0.011614 | 0.007525 | 0.007299 |
| k__Bacteria;p__Bacteroidetes;c__Bacteroidia;o__Bacteroidales;f__NA;g__NA;s__sp12532 | 0.008017 | 0.004125 | 0.004625 | 0.008448 | 0 | 0.008638 | 0.01354 | 0.02194 | 0.021443 | 0.020581 | 0.018796 | 0.026634 | 0.005453 | 0.005053 | 0.008622 | 0.004644 | 0.005302 |
| k__Bacteria;p__Bacteroidetes;c__Bacteroidia;o__Bacteroidales;f__NA;g__NA;s__sp12535 | 0 | 0 | 0 | 0 | 0 | 0 | 0.001221 | 0.001738 | 0.001478 | 0.001665 | 0 | 0.002062 | 0 | 0.000871 | 0.00366 | 0.001548 | 0 |
| k__Bacteria;p__Bacteroidetes;c__Bacteroidia;o__Bacteroidales;f__NA;g__NA;s__sp12536-sp12753 | 0.01657 | 0.012901 | 0.00674 | 0.019935 | 0.01478 | 0.020246 | 0.011019 | 0.021043 | 0.030604 | 0.025414 | 0.022573 | 0.017963 | 0.022909 | 0.010056 | 0.021855 | 0.011416 | 0.012208 |
| k__Bacteria;p__Bacteroidetes;c__Bacteroidia;o__Bacteroidales;f__NA;g__NA;s__sp12539 | 0.009728 | 0.001106 | 0.002657 | 0.009053 | 0 | 0.010522 | 0.003093 | 0.006989 | 0.006497 | 0.0103 | 0.006074 | 0.0108 | 0.006356 | 0.006571 | 0.012529 | 0.003999 | 0.005106 |
| k__Bacteria;p__Bacteroidetes;c__Bacteroidia;o__Bacteroidales;f__NA;g__NA;s__sp12549 | 0.004647 | 0.006846 | 0.001476 | 0.010164 | 0 | 0.014427 | 0.003457 | 0.006282 | 0.006789 | 0.009738 | 0.005409 | 0.008649 | 0 | 0.004928 | 0.006018 | 0.00273 | 0.003535 |
| k__Bacteria;p__Bacteroidetes;c__Bacteroidia;o__Bacteroidales;f__NA;g__NA;s__sp12565 | 0.013609 | 0.0076 | 0.005535 | 0.015916 | 0 | 0.016371 | 0.016555 | 0.026141 | 0.025713 | 0.029125 | 0.024507 | 0.030027 | 0.003743 | 0.00794 | 0.006827 | 0.003999 | 0.005466 |
| k__Bacteria;p__Bacteroidetes;c__Bacteroidia;o__Bacteroidales;f__NA;g__NA;s__sp12572-sp12578-sp12693 | 0.067431 | 0.039669 | 0.034783 | 0.07845 | 0.072112 | 0.07949 | 0.033915 | 0.059959 | 0.088563 | 0.065798 | 0.063639 | 0.122172 | 0.046528 | 0.068175 | 0.081612 | 0.057918 | 0.037638 |
| k__Bacteria;p__Bacteroidetes;c__Bacteroidia;o__Bacteroidales;f__NA;g__NA;s__sp12589 | 0.018409 | 0.0624 | 0.011808 | 0.029968 | 0.026916 | 0.036964 | 0.024767 | 0.03905 | 0.038196 | 0.036764 | 0.047442 | 0.021401 | 0.009777 | 0.035568 | 0.02284 | 0.012125 | 0.010801 |
| k__Bacteria;p__Bacteroidetes;c__Bacteroidia;o__Bacteroidales;f__NA;g__NA;s__sp12590 | 0.002655 | 0.003862 | 0 | 0.004363 | 0 | 0.005261 | 0 | 0 | 0 | 0 | 0 | 0 | 0 | 0.001294 | 0.001091 | 0.000559 | 0.001538 |
| k__Bacteria;p__Bacteroidetes;c__Bacteroidia;o__Bacteroidales;f__NA;g__NA;s__sp12595 | 0.005158 | 0.020467 | 0.009028 | 0.005948 | 0 | 0.005291 | 0.002131 | 0.004029 | 0.003705 | 0.003856 | 0.002629 | 0.006054 | 0 | 0.006023 | 0.009326 | 0.008019 | 0.006382 |
| k__Bacteria;p__Bacteroidetes;c__Bacteroidia;o__Bacteroidales;f__NA;g__NA;s__sp12595-sp12654 | 0.002757 | 0.004722 | 0.001058 | 0.004379 | 0 | 0.005352 | 0 | 0.000573 | 0.001734 | 0.000996 | 0 | 0.001353 | 0.002807 | 0.002763 | 0 | 0.003977 | 0 |
| k__Bacteria;p__Bacteroidetes;c__Bacteroidia;o__Bacteroidales;f__NA;g__NA;s__sp12597 | 0.005617 | 0.01743 | 0.005412 | 0.012974 | 0 | 0.018572 | 0.008836 | 0.012526 | 0.020585 | 0.024545 | 0.016167 | 0.039297 | 0.013487 | 0.009508 | 0.024177 | 0.013953 | 0.009688 |
| k__Bacteria;p__Bacteroidetes;c__Bacteroidia;o__Bacteroidales;f__NA;g__NA;s__sp12616-sp12777 | 0 | 0.000737 | 0 | 0 | 0 | 0 | 0 | 0 | 0 | 0 | 0 | 0 | 0 | 0 | 0 | 0 | 0 |
| k__Bacteria;p__Bacteroidetes;c__Bacteroidia;o__Bacteroidales;f__NA;g__NA;s__sp12629 | 0.002885 | 0.00072 | 0 | 0.00353 | 0 | 0.003256 | 0.002313 | 0.003838 | 0.003869 | 0.003783 | 0.002961 | 0.004857 | 0 | 0 | 0 | 0 | 0.000687 |
| k__Bacteria;p__Bacteroidetes;c__Bacteroidia;o__Bacteroidales;f__NA;g__NA;s__sp12637 | 0.002323 | 0.000474 | 0.003247 | 0.002451 | 0 | 0.002955 | 0.001689 | 0.001432 | 0.005748 | 0.004073 | 0.00417 | 0.001774 | 0.004324 | 0.002414 | 0.006968 | 0.002666 | 0.00252 |
| k__Bacteria;p__Bacteroidetes;c__Bacteroidia;o__Bacteroidales;f__NA;g__NA;s__sp12641 | 0.018383 | 0.013832 | 0.007134 | 0.020017 | 0 | 0.024572 | 0.012968 | 0.024537 | 0.031188 | 0.024636 | 0.022784 | 0.022199 | 0 | 0.011176 | 0.02474 | 0.013931 | 0.013124 |
| k__Bacteria;p__Bacteroidetes;c__Bacteroidia;o__Bacteroidales;f__NA;g__NA;s__sp12654 | 0.004749 | 0.002738 | 0.001845 | 0.004167 | 0 | 0.004583 | 0.001637 | 0.003704 | 0.0048 | 0.004254 | 0 | 0.005167 | 0 | 0.001867 | 0.002323 | 0.001333 | 0 |
| k__Bacteria;p__Bacteroidetes;c__Bacteroidia;o__Bacteroidales;f__NA;g__NA;s__sp12656 | 0.028443 | 0.03479 | 0.012767 | 0.02737 | 0.025749 | 0.027225 | 0.026249 | 0.058833 | 0.055404 | 0.050484 | 0.041429 | 0.034419 | 0.007002 | 0.016253 | 0.014359 | 0.0066 | 0.00684 |
| k__Bacteria;p__Bacteroidetes;c__Bacteroidia;o__Bacteroidales;f__NA;g__NA;s__sp12666 | 0.027754 | 0.027453 | 0.014981 | 0.028089 | 0 | 0.02923 | 0.015697 | 0.019554 | 0.025932 | 0.019839 | 0.028163 | 0.015613 | 0.014552 | 0.006397 | 0.024811 | 0.013609 | 0.014368 |
| k__Bacteria;p__Bacteroidetes;c__Bacteroidia;o__Bacteroidales;f__NA;g__NA;s__sp12757 | 0.001353 | 0.000667 | 0.000738 | 0.00201 | 0 | 0.002261 | 0.002235 | 0.002559 | 0.003522 | 0.003982 | 0.003354 | 0.003193 | 0.003646 | 0.002962 | 0.006546 | 0.003225 | 0.004026 |
| k__Bacteria;p__Bacteroidetes;c__Bacteroidia;o__Bacteroidales;f__NA;g__NA;s__sp12777 | 0.001557 | 0.001632 | 0 | 0 | 0 | 0.001779 | 0 | 0 | 0 | 0 | 0 | 0 | 0 | 0.003062 | 0 | 0 | 0 |
| k__Bacteria;p__Bacteroidetes;c__Bacteroidia;o__Bacteroidales;f__NA;g__NA;s__sp12778 | 0 | 0 | 0.002362 | 0 | 0 | 0 | 0.003431 | 0.004602 | 0.012391 | 0.007621 | 0.008401 | 0.000577 | 0.021973 | 0.098915 | 0.048847 | 0.072645 | 0.04857 |
| k__Bacteria;p__Bacteroidetes;c__Bacteroidia;o__Bacteroidales;f__NA;g__NA;s__sp12785 | 0.00166 | 0.001615 | 0.001378 | 0.002059 | 0 | 0.002744 | 0.004444 | 0.005595 | 0.005913 | 0.006191 | 0.005016 | 0.004169 | 0 | 0.00117 | 0.002921 | 0 | 0.001047 |
| k__Bacteria;p__Bacteroidetes;c__Bacteroidia;o__Bacteroidales;f__NA;g__NA;s__sp12790 | 0.005796 | 0 | 0 | 0.002811 | 0 | 0.001553 | 0.007095 | 0.007275 | 0.012866 | 0.010843 | 0.009851 | 0.012641 | 0.004227 | 0.008239 | 0.009009 | 0.000795 | 0.002356 |
| k__Bacteria;p__Bacteroidetes;c__Bacteroidia;o__Bacteroidales;f__NA;g__NA;s__sp12802 | 0.002451 | 0.002861 | 0.003345 | 0.003644 | 0 | 0.004975 | 0.005172 | 0.007409 | 0.007391 | 0.004887 | 0.006769 | 0.003127 | 0.005743 | 0.002414 | 0.009713 | 0.006321 | 0.003666 |
| k__Bacteria;p__Bacteroidetes;c__Bacteroidia;o__Bacteroidales;f__Porphyromonadaceae;g__Odoribacter;s__sp13184 | 0.03092 | 0.017711 | 0.029691 | 0.024854 | 0.038895 | 0.022401 | 0.037658 | 0.026676 | 0.014362 | 0.023912 | 0.021092 | 0.037035 | 0.015714 | 0.017 | 0.021538 | 0.002902 | 0.006513 |
| k__Bacteria;p__Bacteroidetes;c__Bacteroidia;o__Bacteroidales;f__Porphyromonadaceae;g__Parabacteroides;s__goldsteinii | 0.001966 | 0.031981 | 0.001845 | 0.000817 | 0 | 0.000724 | 0.000806 | 0.001738 | 0.001496 | 0.001575 | 0 | 0.001885 | 0.002872 | 0.002116 | 0.004821 | 0.000774 | 0.001146 |
| k__Bacteria;p__Bacteroidetes;c__Bacteroidia;o__Bacteroidales;f__Prevotellaceae;g__NA;s__sp14210 | 0.016902 | 0.017623 | 0.007208 | 0.017043 | 0 | 0.018889 | 0 | 0.004506 | 0.014472 | 0.008942 | 0.011725 | 0.011998 | 0.033009 | 0.029271 | 0.057434 | 0.020445 | 0.022288 |
| k__Bacteria;p__Bacteroidetes;c__Bacteroidia;o__Bacteroidales;f__Rikenellaceae;g__Alistipes;s__putredinis | 0 | 0.011304 | 0.011094 | 0 | 0.048697 | 0 | 0 | 0.004182 | 0.001551 | 0.003222 | 0.003294 | 0.004147 | 0.009777 | 0.003161 | 0.011614 | 0.001956 | 0.001669 |
| k__Bacteria;p__Bacteroidetes;c__Bacteroidia;o__Bacteroidales;f__Rikenellaceae;g__Alistipes;s__sp14330 | 0 | 0 | 0 | 0 | 0 | 0 | 0 | 0 | 0 | 0 | 0 | 0 | 0 | 0.002738 | 0 | 0 | 0 |
| k__Bacteria;p__Bacteroidetes;c__Bacteroidia;o__Bacteroidales;f__Rikenellaceae;g__Alistipes;s__sp14336 | 0.023975 | 0.003212 | 0.061399 | 0.031717 | 0.169273 | 0.037567 | 0.072717 | 0.054593 | 0.01449 | 0.038972 | 0.032152 | 0.028697 | 0.028556 | 0.007915 | 0.021714 | 0.012405 | 0.006742 |
| k__Bacteria;p__Bacteroidetes;c__Bacteroidia;o__Bacteroidales;f__Rikenellaceae;g__Alistipes;s__sp14338 | 0.014298 | 0.010356 | 0.007749 | 0.013726 | 0 | 0.014547 | 0.015671 | 0.026867 | 0.006971 | 0.013884 | 0.009337 | 0.021112 | 0.003904 | 0.008388 | 0.010065 | 0.003203 | 0.002389 |
| k__Bacteria;p__Bacteroidetes;c__Bacteroidia;o__Bacteroidales;f__Rikenellaceae;g__Alistipes;s__sp14343 | 0 | 0 | 0 | 0 | 0 | 0 | 0.000936 | 0 | 0.000894 | 0.000326 | 0 | 0 | 0 | 0.00117 | 0.003097 | 0 | 0 |
| k__Bacteria;p__Bacteroidetes;c__Bacteroidia;o__Bacteroidales;f__Rikenellaceae;g__Alistipes;s__sp14354 | 0.001226 | 0 | 0.021352 | 0.002304 | 0.050564 | 0.002668 | 0.047352 | 0.038668 | 0.01365 | 0.027514 | 0.022814 | 0.027943 | 0.023264 | 0.011972 | 0.026782 | 0.009503 | 0.009099 |
| k__Bacteria;p__Bacteroidetes;c__Bacteroidia;o__Bacteroidales;f__Rikenellaceae;g__Alistipes;s__timonensis | 0.00577 | 0.009759 | 0.004698 | 0.003382 | 0.013613 | 0.002186 | 0.000364 | 0.00191 | 0.002372 | 0.000833 | 0.001209 | 0.00031 | 0.006905 | 0.00346 | 0.010311 | 0.003848 | 0.002553 |
| k__Bacteria;p__Bacteroidetes;c__Bacteroidia;o__Bacteroidales;f__Rikenellaceae;g__NA;s__sp14636 | 0 | 0.082849 | 0.009544 | 0.010229 | 0 | 0.017608 | 0.010941 | 0.016155 | 0.009928 | 0.004453 | 0.006497 | 0.015213 | 0 | 0.003534 | 0 | 0 | 0.000458 |
| k__Bacteria;p__Cyanobacteria;c__Melainabacteria;o__Gastranaerophilales;f__NA;g__NA;s__sp23727 | 0.001787 | 0.000632 | 0 | 0.002075 | 0 | 0.001794 | 0.002079 | 0.000936 | 0.002482 | 0.003403 | 0 | 0.000843 | 0.001742 | 0.000946 | 0 | 0 | 0 |
| k__Bacteria;p__Cyanobacteria;c__Melainabacteria;o__Gastranaerophilales;f__NA;g__NA;s__sp23729 | 0 | 0 | 0 | 0 | 0 | 0 | 0 | 0 | 0 | 0 | 0 | 0 | 0 | 0.0056 | 0 | 0 | 0 |
| k__Bacteria;p__Deferribacteres;c__Deferribacteres;o__Deferribacterales;f__Deferribacteraceae;g__Mucispirillum;s__sp23890 | 0 | 0 | 0 | 0 | 0 | 0 | 0 | 0.000859 | 0.00073 | 0.001358 | 0 | 0.001353 | 0.007292 | 0.000821 | 0 | 0.000903 | 0.004484 |
| k__Bacteria;p__Firmicutes;c__Bacilli;o__Lactobacillales;f__Lactobacillaceae;g__Lactobacillus;s__NA | 0.011847 | 0.021959 | 0.010651 | 0.016863 | 0.051109 | 0.016778 | 0.000702 | 0.00401 | 0.005712 | 0.001864 | 0.003143 | 0.001619 | 0 | 0 | 0.004153 | 0.000365 | 0.002651 |
| k__Bacteria;p__Firmicutes;c__Bacilli;o__Lactobacillales;f__Lactobacillaceae;g__Lactobacillus;s__johnsonii | 0.017745 | 0.003072 | 0.002927 | 0.014396 | 0.025904 | 0.011035 | 0.00551 | 0.050717 | 0.045167 | 0.022228 | 0.058592 | 0.005611 | 0.0141 | 0 | 0 | 0.007675 | 0.008771 |
| k__Bacteria;p__Firmicutes;c__Bacilli;o__Lactobacillales;f__Lactobacillaceae;g__Lactobacillus;s__reuteri-vaginalis | 0.007532 | 0.001176 | 0.001919 | 0.004902 | 0.010735 | 0.003904 | 0.005042 | 0.011705 | 0.009891 | 0.006697 | 0.016831 | 0.003327 | 0.001775 | 0.00331 | 0.003062 | 0.000623 | 0 |
| k__Bacteria;p__Firmicutes;c__Bacilli;o__Lactobacillales;f__Lactobacillaceae;g__Lactobacillus;s__taiwanensis | 0 | 0 | 0 | 0 | 0 | 0 | 0.003846 | 0.004506 | 0.006716 | 0.007114 | 0.011483 | 0.003814 | 0 | 0.013092 | 0.010523 | 0.007955 | 0 |
| k__Bacteria;p__Firmicutes;c__Clostridia;o__Clostridiales;f__Family XIII;g__NA;s__sp31564 | 0 | 0 | 0 | 0 | 0 | 0 | 0 | 0 | 0 | 0 | 0 | 0.000266 | 0 | 0 | 0 | 0 | 0 |
| k__Bacteria;p__Firmicutes;c__Clostridia;o__Clostridiales;f__Family XIII;g__NA;s__sp31651 | 0 | 0 | 0 | 0 | 0 | 0 | 0 | 0 | 0.000858 | 0.000398 | 0 | 0 | 0 | 0.000672 | 0 | 0 | 0 |
| k__Bacteria;p__Firmicutes;c__Clostridia;o__Clostridiales;f__Lachnospiraceae;g__Acetatifactor;s__sp31893 | 0 | 0.000456 | 0 | 0 | 0 | 0 | 0.001351 | 0 | 0 | 0.000308 | 0 | 0.000288 | 0.000419 | 0 | 0 | 0.000559 | 0.000524 |
| k__Bacteria;p__Firmicutes;c__Clostridia;o__Clostridiales;f__Lachnospiraceae;g__Blautia;s__sp32015 | 0.002885 | 0.000544 | 0 | 0.001373 | 0 | 0.000618 | 0.001663 | 0.000611 | 0.001533 | 0.001448 | 0.001722 | 0.00122 | 0.001613 | 0.005551 | 0.001161 | 0.002924 | 0.005629 |
| k__Bacteria;p__Firmicutes;c__Clostridia;o__Clostridiales;f__Lachnospiraceae;g__Lachnoclostridium;s__sp32351 | 0 | 0 | 0 | 0 | 0 | 0 | 0 | 0 | 0 | 0 | 0 | 0 | 0 | 0.001817 | 0 | 0 | 0 |
| k__Bacteria;p__Firmicutes;c__Clostridia;o__Clostridiales;f__Lachnospiraceae;g__Lachnoclostridium;s__sp32364 | 0 | 0 | 0 | 0 | 0 | 0 | 0 | 0.000286 | 0 | 0.000344 | 0 | 0 | 0 | 0 | 0 | 0 | 0 |
| k__Bacteria;p__Firmicutes;c__Clostridia;o__Clostridiales;f__Lachnospiraceae;g__Lachnoclostridium;s__sp32366 | 0 | 0 | 0 | 0 | 0 | 0 | 0.003093 | 0.001318 | 0.005165 | 0.005123 | 0.006406 | 0.00428 | 0.003485 | 0.006272 | 0.003484 | 0.002988 | 0.005106 |
| k__Bacteria;p__Firmicutes;c__Clostridia;o__Clostridiales;f__Lachnospiraceae;g__Lachnoclostridium;s__sp32414 | 0 | 0 | 0 | 0 | 0 | 0 | 0 | 0 | 0 | 0 | 0 | 0 | 0.001839 | 0 | 0 | 0 | 0 |
| k__Bacteria;p__Firmicutes;c__Clostridia;o__Clostridiales;f__Lachnospiraceae;g__Lachnoclostridium;s__sp32442 | 0.002119 | 0.002721 | 0.003764 | 0.001373 | 0.004979 | 0.000724 | 0 | 0 | 0 | 0.000905 | 0.001209 | 0 | 0.005421 | 0 | 0.004399 | 0.009223 | 0.005335 |
| k__Bacteria;p__Firmicutes;c__Clostridia;o__Clostridiales;f__Lachnospiraceae;g__Marvinbryantia;s__sp32979 | 0.004672 | 0.002212 | 0.001697 | 0.002222 | 0 | 0.001507 | 0.004106 | 0.0021 | 0.002756 | 0.002118 | 0.00417 | 0.002595 | 0.002743 | 0.005526 | 0.009009 | 0.002515 | 0.003011 |
| k__Bacteria;p__Firmicutes;c__Clostridia;o__Clostridiales;f__Lachnospiraceae;g__NA;s__sp32146 | 0.003447 | 0.002352 | 0.009446 | 0.002255 | 0 | 0.001266 | 0.001897 | 0.000535 | 0 | 0.000579 | 0 | 0 | 0.001549 | 0.003435 | 0.00176 | 0.001376 | 0.001604 |
| k__Bacteria;p__Firmicutes;c__Clostridia;o__Clostridiales;f__Lachnospiraceae;g__NA;s__sp32156 | 0 | 0 | 0 | 0 | 0 | 0 | 0 | 0 | 0 | 0.000235 | 0 | 0 | 0 | 0 | 0 | 0 | 0 |
| k__Bacteria;p__Firmicutes;c__Clostridia;o__Clostridiales;f__Lachnospiraceae;g__NA;s__sp32165 | 0.000383 | 0 | 0.001156 | 0.000212 | 0 | 0 | 0.000676 | 0 | 0 | 0 | 0 | 0.000288 | 0 | 0.000523 | 0 | 0 | 0.000851 |
| k__Bacteria;p__Firmicutes;c__Clostridia;o__Clostridiales;f__Lachnospiraceae;g__NA;s__sp32166 | 0 | 0.000316 | 0.00246 | 0.000359 | 0 | 0 | 0 | 0 | 0.000347 | 0 | 0 | 0 | 0 | 0 | 0 | 0 | 0 |
| k__Bacteria;p__Firmicutes;c__Clostridia;o__Clostridiales;f__Lachnospiraceae;g__NA;s__sp32255 | 0 | 0 | 0 | 0 | 0 | 0 | 0 | 0 | 0 | 0 | 0 | 0 | 0.003162 | 0.003086 | 0.001548 | 0.005224 | 0.002978 |
| k__Bacteria;p__Firmicutes;c__Clostridia;o__Clostridiales;f__Lachnospiraceae;g__NA;s__sp32261 | 0 | 0 | 0 | 0 | 0 | 0 | 0 | 0 | 0 | 0 | 0 | 0 | 0.007712 | 0 | 0.008587 | 0.01019 | 0.01391 |

| | | | | | | | | | | | | | | | | | |
|---|---|---|---|---|---|---|---|---|---|---|---|---|---|---|---|---|---|
| k__Bacteria;p__Firmicutes;c__Clostridia;o__Clostridiales;f__Lachnospiraceae;g__NA;s__sp32486 | 0 | 0.000228 | 0 | 0.00018 | 0 | 0.000196 | 0.00065 | 0 | 0 | 0 | 0.000332 | 0 | 0 | 0.001095 | 0 | 0 | 0 |
| k__Bacteria;p__Firmicutes;c__Clostridia;o__Clostridiales;f__Lachnospiraceae;g__NA;s__sp32594 | 0.004468 | 0.005389 | 0.010086 | 0.001389 | 0 | 0.001176 | 0.009278 | 0.001489 | 0.002153 | 0.001702 | 0.002871 | 0.004613 | 0.006002 | 0.002016 | 0.000985 | 0 | 0 |
| k__Bacteria;p__Firmicutes;c__Clostridia;o__Clostridiales;f__Lachnospiraceae;g__NA;s__sp32594-sp32647 | 0 | 0 | 0 | 0 | 0 | 0 | 0 | 0 | 0 | 0 | 0 | 0 | 0.00868 | 0 | 0.010347 | 0.008342 | 0.009033 |
| k__Bacteria;p__Firmicutes;c__Clostridia;o__Clostridiales;f__Lachnospiraceae;g__NA;s__sp32596 | 0 | 0 | 0 | 0 | 0 | 0 | 0 | 0 | 0 | 0 | 0 | 0 | 0.030621 | 0 | 0.019391 | 0.012555 | 0.020259 |
| k__Bacteria;p__Firmicutes;c__Clostridia;o__Clostridiales;f__Lachnospiraceae;g__NA;s__sp32617-sp32782 | 0.005209 | 0.003265 | 0.031019 | 0.00299 | 0.013691 | 0.001281 | 0.010162 | 0.001241 | 0.003759 | 0.005394 | 0.003173 | 0.00397 | 0 | 0 | 0 | 0 | 0 |
| k__Bacteria;p__Firmicutes;c__Clostridia;o__Clostridiales;f__Lachnospiraceae;g__NA;s__sp32622 | 0.001481 | 0 | 0.002829 | 0.000654 | 0 | 0 | 0 | 0.000917 | 0.001423 | 0 | 0 | 0 | 0.001904 | 0.001045 | 0 | 0.004063 | 0.002847 |
| k__Bacteria;p__Firmicutes;c__Clostridia;o__Clostridiales;f__Lachnospiraceae;g__NA;s__sp32623 | 0 | 0.001176 | 0.001427 | 0 | 0 | 0 | 0 | 0 | 0 | 0 | 0 | 0 | 0 | 0.000722 | 0 | 0 | 0 |
| k__Bacteria;p__Firmicutes;c__Clostridia;o__Clostridiales;f__Lachnospiraceae;g__NA;s__sp32628-sp32767 | 0 | 0 | 0 | 0 | 0 | 0 | 0 | 0 | 0 | 0 | 0 | 0 | 0.004679 | 0 | 0 | 0 | 0 |
| k__Bacteria;p__Firmicutes;c__Clostridia;o__Clostridiales;f__Lachnospiraceae;g__NA;s__sp32648 | 0.001379 | 0 | 0.004329 | 0.000474 | 0.003345 | 0 | 0.004184 | 0.002291 | 0.001241 | 0.002534 | 0.002871 | 0.00377 | 0.016036 | 0.022899 | 0.006827 | 0.027884 | 0.022059 |
| k__Bacteria;p__Firmicutes;c__Clostridia;o__Clostridiales;f__Lachnospiraceae;g__NA;s__sp32655 | 0.002987 | 0.002019 | 0.009249 | 0 | 0 | 0 | 0.009044 | 0.003991 | 0.003029 | 0.004924 | 0.005318 | 0.004546 | 0.007389 | 0.007293 | 0.004012 | 0.010083 | 0.008215 |
| k__Bacteria;p__Firmicutes;c__Clostridia;o__Clostridiales;f__Lachnospiraceae;g__NA;s__sp32668 | 0.029949 | 0.016184 | 0.047279 | 0.01987 | 0.044807 | 0.010673 | 0 | 0 | 0 | 0.000706 | 0 | 0.000665 | 0 | 0 | 0 | 0 | 0 |
| k__Bacteria;p__Firmicutes;c__Clostridia;o__Clostridiales;f__Lachnospiraceae;g__NA;s__sp32693 | 0.012434 | 0 | 0 | 0.015327 | 0 | 0.017788 | 0 | 0 | 0 | 0 | 0 | 0 | 0 | 0 | 0 | 0 | 0 |
| k__Bacteria;p__Firmicutes;c__Clostridia;o__Clostridiales;f__Lachnospiraceae;g__NA;s__sp32704 | 0 | 0 | 0.012078 | 0 | 0 | 0 | 0.057955 | 0.018351 | 0.013304 | 0.017196 | 0.025171 | 0.033465 | 0.11148 | 0.025339 | 0.061974 | 0.04532 | 0.114453 |
| k__Bacteria;p__Firmicutes;c__Clostridia;o__Clostridiales;f__Lachnospiraceae;g__NA;s__sp32721 | 0 | 0 | 0 | 0 | 0 | 0 | 0 | 0 | 0 | 0 | 0 | 0 | 0.004195 | 0 | 0.001724 | 0.007073 | 0.002193 |
| k__Bacteria;p__Firmicutes;c__Clostridia;o__Clostridiales;f__Lachnospiraceae;g__NA;s__sp32735 | 0 | 0 | 0 | 0 | 0 | 0 | 0 | 0 | 0 | 0 | 0 | 0 | 0 | 0.037261 | 0 | 0 | 0 |
| k__Bacteria;p__Firmicutes;c__Clostridia;o__Clostridiales;f__Lachnospiraceae;g__NA;s__sp32746-sp32777 | 0.005336 | 0.003528 | 0.013382 | 0.002974 | 0 | 0.001613 | 0.008888 | 0.002597 | 0.002774 | 0.004706 | 0.005832 | 0.004369 | 0.003194 | 0.00351 | 0.004188 | 0.005654 | 0.005498 |
| k__Bacteria;p__Firmicutes;c__Clostridia;o__Clostridiales;f__Lachnospiraceae;g__NA;s__sp32758 | 0 | 0 | 0 | 0 | 0 | 0 | 0 | 0 | 0 | 0.001014 | 0 | 0 | 0.001162 | 0.002813 | 0 | 0.000559 | 0 |
| k__Bacteria;p__Firmicutes;c__Clostridia;o__Clostridiales;f__Lachnospiraceae;g__NA;s__sp32763 | 0.001532 | 0 | 0.002017 | 0.000523 | 0 | 0 | 0.032746 | 0.008039 | 0.009198 | 0.010155 | 0.010878 | 0.009713 | 0.014455 | 0.003435 | 0.01918 | 0.019908 | 0.015906 |
| k__Bacteria;p__Firmicutes;c__Clostridia;o__Clostridiales;f__Lachnospiraceae;g__NA;s__sp32766 | 0 | 0 | 0.010209 | 0 | 0 | 0 | 0.049041 | 0.018713 | 0.011606 | 0.014771 | 0.017436 | 0.025215 | 0.090023 | 0.0228 | 0.041598 | 0.036376 | 0.084539 |
| k__Bacteria;p__Firmicutes;c__Clostridia;o__Clostridiales;f__Lachnospiraceae;g__NA;s__sp32778 | 0 | 0 | 0 | 0 | 0 | 0 | 0 | 0 | 0 | 0 | 0 | 0 | 0 | 0.027205 | 0 | 0 | 0 |
| k__Bacteria;p__Firmicutes;c__Clostridia;o__Clostridiales;f__Lachnospiraceae;g__NA;s__sp32791 | 0.007353 | 0.010023 | 0.021057 | 0.00366 | 0.006068 | 0.00208 | 0.017932 | 0.002482 | 0.00323 | 0.00391 | 0.005379 | 0.009669 | 0.012713 | 0.003559 | 0.003906 | 0.004966 | 0.006873 |
| k__Bacteria;p__Firmicutes;c__Clostridia;o__Clostridiales;f__Lachnospiraceae;g__NA;s__sp32816 | 0 | 0 | 0 | 0 | 0 | 0 | 0 | 0 | 0 | 0 | 0 | 0 | 0 | 0.000597 | 0 | 0 | 0 |
| k__Bacteria;p__Firmicutes;c__Clostridia;o__Clostridiales;f__Lachnospiraceae;g__NA;s__sp32826 | 0 | 0 | 0 | 0 | 0 | 0 | 0 | 0 | 0 | 0 | 0 | 0 | 0 | 0.000821 | 0 | 0.001763 | 0.000556 |
| k__Bacteria;p__Firmicutes;c__Clostridia;o__Clostridiales;f__Lachnospiraceae;g__NA;s__sp32862 | 0 | 0 | 0.000467 | 0 | 0 | 0 | 0 | 0 | 0 | 0 | 0 | 0 | 0 | 0 | 0 | 0 | 0 |
| k__Bacteria;p__Firmicutes;c__Clostridia;o__Clostridiales;f__Lachnospiraceae;g__NA;s__sp32862-sp32872-sp32880 | 0 | 0 | 0 | 0 | 0 | 0 | 0 | 0 | 0 | 0 | 0 | 0 | 0 | 0.000523 | 0 | 0 | 0 |
| k__Bacteria;p__Firmicutes;c__Clostridia;o__Clostridiales;f__Lachnospiraceae;g__NA;s__sp32885 | 0 | 0.002141 | 0 | 0 | 0 | 0 | 0 | 0 | 0 | 0 | 0 | 0 | 0 | 0 | 0 | 0 | 0 |
| k__Bacteria;p__Firmicutes;c__Clostridia;o__Clostridiales;f__Lachnospiraceae;g__NA;s__sp32910 | 0.002783 | 0.002686 | 0.038448 | 0.002337 | 0.01797 | 0.001417 | 0.024118 | 0.001528 | 0.004161 | 0.003186 | 0.008552 | 0.003304 | 0.003453 | 0 | 0.002006 | 0 | 0 |
| k__Bacteria;p__Firmicutes;c__Clostridia;o__Clostridiales;f__Lachnospiraceae;g__NA;s__sp33416-sp33419 | 0 | 0 | 0 | 0 | 0 | 0 | 0 | 0 | 0 | 0 | 0 | 0 | 0 | 0.000846 | 0 | 0 | 0 |
| k__Bacteria;p__Firmicutes;c__Clostridia;o__Clostridiales;f__Lachnospiraceae;g__NA;s__sp33417 | 0 | 0 | 0 | 0 | 0 | 0 | 0 | 0 | 0 | 0 | 0 | 0 | 0 | 0.002016 | 0 | 0 | 0 |
| k__Bacteria;p__Firmicutes;c__Clostridia;o__Clostridiales;f__Lachnospiraceae;g__NA;s__sp33421-sp33679 | 0 | 0 | 0 | 0 | 0 | 0 | 0 | 0 | 0 | 0.000525 | 0 | 0 | 0 | 0.001294 | 0 | 0.001225 | 0.001211 |
| k__Bacteria;p__Firmicutes;c__Clostridia;o__Clostridiales;f__Lachnospiraceae;g__NA;s__sp33426 | 0 | 0 | 0 | 0 | 0 | 0 | 0 | 0 | 0.000493 | 0 | 0 | 0 | 0.001678 | 0.00112 | 0 | 0.002343 | 0.003469 |
| k__Bacteria;p__Firmicutes;c__Clostridia;o__Clostridiales;f__Lachnospiraceae;g__NA;s__sp33433 | 0.000332 | 0.000281 | 0.001747 | 0.000359 | 0 | 0.000271 | 0.002209 | 0.000496 | 0.000566 | 0.000869 | 0.001148 | 0.000754 | 0 | 0.000747 | 0 | 0.001677 | 0.001342 |
| k__Bacteria;p__Firmicutes;c__Clostridia;o__Clostridiales;f__Lachnospiraceae;g__NA;s__sp33436 | 0 | 0 | 0 | 0 | 0 | 0 | 0 | 0 | 0 | 0 | 0 | 0 | 0 | 0.000423 | 0 | 0 | 0 |
| k__Bacteria;p__Firmicutes;c__Clostridia;o__Clostridiales;f__Lachnospiraceae;g__NA;s__sp33451-sp33593 | 0 | 0 | 0.000541 | 0.000196 | 0 | 0 | 0.002521 | 0.000496 | 0.001058 | 0.000996 | 0.002055 | 0 | 0.000549 | 0 | 0.000985 | 0.000838 | 0.000982 |
| k__Bacteria;p__Firmicutes;c__Clostridia;o__Clostridiales;f__Lachnospiraceae;g__NA;s__sp33453 | 0.004238 | 0.003616 | 0 | 0.003399 | 0 | 0.002171 | 0.004028 | 0.001241 | 0.001971 | 0.002154 | 0.003566 | 0.002683 | 0.004646 | 0.004655 | 0.002956 | 0.007116 | 0.005237 |
| k__Bacteria;p__Firmicutes;c__Clostridia;o__Clostridiales;f__Lachnospiraceae;g__NA;s__sp33456 | 0 | 0 | 0 | 0 | 0 | 0 | 0.00104 | 0 | 0.000401 | 0.000597 | 0 | 0 | 0.000484 | 0.001444 | 0 | 0.000817 | 0.000916 |
| k__Bacteria;p__Firmicutes;c__Clostridia;o__Clostridiales;f__Lachnospiraceae;g__NA;s__sp33459 | 0.000306 | 0.000298 | 0 | 0 | 0 | 0.000241 | 0.003093 | 0.000745 | 0.000839 | 0.001738 | 0 | 0.001153 | 0 | 0.003236 | 0 | 0.00301 | 0.00396 |
| k__Bacteria;p__Firmicutes;c__Clostridia;o__Clostridiales;f__Lachnospiraceae;g__NA;s__sp33470 | 0 | 0 | 0 | 0.000441 | 0 | 0 | 0 | 0 | 0 | 0 | 0 | 0 | 0 | 0 | 0 | 0 | 0 |
| k__Bacteria;p__Firmicutes;c__Clostridia;o__Clostridiales;f__Lachnospiraceae;g__NA;s__sp33478 | 0 | 0.001422 | 0.001033 | 0.000654 | 0 | 0.000618 | 0.000624 | 0 | 0 | 0.000308 | 0.000514 | 0 | 0.002839 | 0.00336 | 0.004469 | 0.002472 | 0.002618 |
| k__Bacteria;p__Firmicutes;c__Clostridia;o__Clostridiales;f__Lachnospiraceae;g__NA;s__sp33492 | 0 | 0 | 0.005732 | 0 | 0 | 0 | 0 | 0 | 0 | 0 | 0 | 0 | 0 | 0.000647 | 0 | 0 | 0 |
| k__Bacteria;p__Firmicutes;c__Clostridia;o__Clostridiales;f__Lachnospiraceae;g__NA;s__sp33503 | 0 | 0 | 0 | 0 | 0 | 0 | 0 | 0 | 0 | 0 | 0 | 0 | 0.002388 | 0.008712 | 0 | 0.007955 | 0.006513 |
| k__Bacteria;p__Firmicutes;c__Clostridia;o__Clostridiales;f__Lachnospiraceae;g__NA;s__sp33513 | 0.008553 | 0.006477 | 0.017146 | 0.004396 | 0.009413 | 0.004734 | 0.004548 | 0.003514 | 0.003449 | 0.003638 | 0.006557 | 0.004613 | 0.02065 | 0.00112 | 0.008341 | 0.006794 | 0.010113 |
| k__Bacteria;p__Firmicutes;c__Clostridia;o__Clostridiales;f__Lachnospiraceae;g__NA;s__sp33518 | 0 | 0 | 0 | 0 | 0 | 0 | 0 | 0 | 0 | 0 | 0 | 0 | 0 | 0 | 0 | 0 | 0.001604 |
| k__Bacteria;p__Firmicutes;c__Clostridia;o__Clostridiales;f__Lachnospiraceae;g__NA;s__sp33522 | 0.000664 | 0.001545 | 0.003542 | 0.000507 | 0 | 0.000573 | 0.002261 | 0.000267 | 0.000237 | 0.000217 | 0 | 0.000399 | 0 | 0.000572 | 0 | 0.001849 | 0.001767 |
| k__Bacteria;p__Firmicutes;c__Clostridia;o__Clostridiales;f__Lachnospiraceae;g__NA;s__sp33525 | 0 | 0 | 0.000271 | 0 | 0 | 0 | 0 | 0 | 0 | 0 | 0 | 0 | 0 | 0 | 0 | 0 | 0 |
| k__Bacteria;p__Firmicutes;c__Clostridia;o__Clostridiales;f__Lachnospiraceae;g__NA;s__sp33529 | 0 | 0 | 0 | 0 | 0 | 0 | 0.000286 | 0 | 0 | 0 | 0 | 0 | 0 | 0 | 0 | 0.000623 | 0 |
| k__Bacteria;p__Firmicutes;c__Clostridia;o__Clostridiales;f__Lachnospiraceae;g__NA;s__sp33531 | 0 | 0 | 0 | 0 | 0 | 0 | 0 | 0 | 0 | 0 | 0 | 0 | 0.001742 | 0.001668 | 0 | 0.001075 | 0 |
| k__Bacteria;p__Firmicutes;c__Clostridia;o__Clostridiales;f__Lachnospiraceae;g__NA;s__sp33539 | 0 | 0 | 0 | 0 | 0 | 0 | 0 | 0 | 0 | 0 | 0 | 0 | 0 | 0.000921 | 0 | 0 | 0 |
| k__Bacteria;p__Firmicutes;c__Clostridia;o__Clostridiales;f__Lachnospiraceae;g__NA;s__sp33540 | 0 | 0 | 0 | 0 | 0 | 0 | 0.001429 | 0.001337 | 0.00115 | 0.0021 | 0.001753 | 0.000976 | 0.00213 | 0 | 0 | 0 | 0.001342 |
| k__Bacteria;p__Firmicutes;c__Clostridia;o__Clostridiales;f__Lachnospiraceae;g__NA;s__sp33565 | 0.003166 | 0.000807 | 0.014858 | 0.001683 | 0.007857 | 0.002563 | 0.006809 | 0.003895 | 0.003431 | 0.004181 | 0.007192 | 0.002728 | 0.046883 | 0.01817 | 0.009396 | 0.085372 | 0.048701 |
| k__Bacteria;p__Firmicutes;c__Clostridia;o__Clostridiales;f__Lachnospiraceae;g__NA;s__sp33577 | 0 | 0 | 0.00091 | 0 | 0 | 0 | 0.001273 | 0.000745 | 0.001004 | 0.00114 | 0.001813 | 0.001064 | 0.000839 | 0.001643 | 0 | 0.001483 | 0.001309 |
| k__Bacteria;p__Firmicutes;c__Clostridia;o__Clostridiales;f__Lachnospiraceae;g__NA;s__sp33579 | 0 | 0 | 0 | 0 | 0 | 0 | 0 | 0 | 0 | 0 | 0 | 0 | 0 | 0.000697 | 0 | 0 | 0 |
| k__Bacteria;p__Firmicutes;c__Clostridia;o__Clostridiales;f__Lachnospiraceae;g__NA;s__sp33582 | 0.001226 | 0 | 0.004354 | 0.000654 | 0 | 0 | 0.002105 | 0.00084 | 0.001533 | 0.001448 | 0 | 0 | 0.004001 | 0.005078 | 0 | 0.004708 | 0.003502 |
| k__Bacteria;p__Firmicutes;c__Clostridia;o__Clostridiales;f__Lachnospiraceae;g__NA;s__sp33597 | 0 | 0.001264 | 0.004575 | 0 | 0 | 0 | 0.001663 | 0.000458 | 0.001369 | 0.000869 | 0.00139 | 0.000554 | 0.00142 | 0 | 0 | 0.001376 | 0 |
| k__Bacteria;p__Firmicutes;c__Clostridia;o__Clostridiales;f__Lachnospiraceae;g__NA;s__sp33614 | 0 | 0 | 0 | 0 | 0 | 0 | 0 | 0 | 0 | 0 | 0 | 0 | 0 | 0.000871 | 0 | 0 | 0 |
| k__Bacteria;p__Firmicutes;c__Clostridia;o__Clostridiales;f__Lachnospiraceae;g__NA;s__sp33616-sp33639 | 0 | 0 | 0 | 0 | 0 | 0 | 0 | 0 | 0 | 0 | 0 | 0 | 0.001097 | 0.000747 | 0 | 0.002515 | 0.001866 |
| k__Bacteria;p__Firmicutes;c__Clostridia;o__Clostridiales;f__Lachnospiraceae;g__NA;s__sp33620 | 0 | 0 | 0 | 0 | 0 | 0 | 0 | 0 | 0.000474 | 0 | 0 | 0.000665 | 0 | 0 | 0 | 0 | 0.000949 |
| k__Bacteria;p__Firmicutes;c__Clostridia;o__Clostridiales;f__Lachnospiraceae;g__NA;s__sp33625 | 0.000409 | 0.000562 | 0.001033 | 0.000229 | 0 | 0.000437 | 0 | 0 | 0 | 0 | 0 | 0 | 0 | 0.000423 | 0 | 0.000279 | 0 |
| k__Bacteria;p__Firmicutes;c__Clostridia;o__Clostridiales;f__Lachnospiraceae;g__NA;s__sp33639 | 0 | 0 | 0.000713 | 0 | 0 | 0 | 0.000546 | 0.000325 | 0 | 0 | 0.000665 | 0 | 0 | 0.000523 | 0 | 0.001333 | 0.001342 |
| k__Bacteria;p__Firmicutes;c__Clostridia;o__Clostridiales;f__Lachnospiraceae;g__NA;s__sp33645-sp33744 | 0 | 0 | 0.000861 | 0 | 0 | 0 | 0.001897 | 0.001165 | 0.001807 | 0.001991 | 0.002297 | 0.00173 | 0.004066 | 0.009508 | 0.003343 | 0.006923 | 0.008739 |
| k__Bacteria;p__Firmicutes;c__Clostridia;o__Clostridiales;f__Lachnospiraceae;g__NA;s__sp33658 | 0.000792 | 0.000527 | 0.001205 | 0.000605 | 0 | 0 | 0.001819 | 0.001547 | 0.000803 | 0.001557 | 0 | 0.000931 | 0.005518 | 0.004754 | 0 | 0.004966 | 0.010244 |
| k__Bacteria;p__Firmicutes;c__Clostridia;o__Clostridiales;f__Lachnospiraceae;g__NA;s__sp33670 | 0 | 0 | 0.00273 | 0 | 0 | 0 | 0.005588 | 0.001509 | 0.001314 | 0.002118 | 0.00281 | 0.001464 | 0.003453 | 0.0056 | 0.002851 | 0.003547 | 0.006546 |
| k__Bacteria;p__Firmicutes;c__Clostridia;o__Clostridiales;f__Lachnospiraceae;g__NA;s__sp33673 | 0 | 0 | 0 | 0 | 0 | 0 | 0 | 0 | 0 | 0 | 0 | 0 | 0 | 0.001593 | 0 | 0.001612 | 0 |
| k__Bacteria;p__Firmicutes;c__Clostridia;o__Clostridiales;f__Lachnospiraceae;g__NA;s__sp33682 | 0 | 0 | 0 | 0 | 0 | 0 | 0 | 0 | 0.00042 | 0 | 0 | 0 | 0.001033 | 0.001444 | 0 | 0.001526 | 0.002193 |
| k__Bacteria;p__Firmicutes;c__Clostridia;o__Clostridiales;f__Lachnospiraceae;g__NA;s__sp33690 | 0 | 0 | 0.001033 | 0 | 0 | 0 | 0.000884 | 0 | 0 | 0 | 0 | 0 | 0 | 0.000846 | 0 | 0.00058 | 0 |
| k__Bacteria;p__Firmicutes;c__Clostridia;o__Clostridiales;f__Lachnospiraceae;g__NA;s__sp33694 | 0 | 0 | 0 | 0 | 0 | 0 | 0.001144 | 0.000477 | 0 | 0 | 0 | 0 | 0 | 0 | 0 | 0 | 0 |
| k__Bacteria;p__Firmicutes;c__Clostridia;o__Clostridiales;f__Lachnospiraceae;g__NA;s__sp33714 | 0 | 0 | 0.001624 | 0 | 0 | 0 | 0.001169 | 0 | 0 | 0 | 0 | 0 | 0.002678 | 0.001568 | 0.001021 | 0.001849 | 0.002913 |
| k__Bacteria;p__Firmicutes;c__Clostridia;o__Clostridiales;f__Lachnospiraceae;g__NA;s__sp33718 | 0 | 0 | 0 | 0 | 0 | 0 | 0 | 0 | 0 | 0 | 0 | 0 | 0 | 0 | 0 | 0.000494 | 0 |
| k__Bacteria;p__Firmicutes;c__Clostridia;o__Clostridiales;f__Lachnospiraceae;g__NA;s__sp33721 | 0 | 0.000737 | 0 | 0 | 0 | 0 | 0.000754 | 0 | 0 | 0.000471 | 0.000423 | 0 | 0 | 0.000597 | 0 | 0 | 0 |
| k__Bacteria;p__Firmicutes;c__Clostridia;o__Clostridiales;f__Lachnospiraceae;g__NA;s__sp33725 | 0 | 0 | 0 | 0 | 0 | 0 | 0 | 0 | 0 | 0 | 0 | 0 | 0 | 0 | 0 | 0.000967 | 0.001146 |
| k__Bacteria;p__Firmicutes;c__Clostridia;o__Clostridiales;f__Lachnospiraceae;g__NA;s__sp33731 | 0.004264 | 0.001387 | 0 | 0.002108 | 0 | 0.001583 | 0.000312 | 0.000668 | 0.000438 | 0.000561 | 0.000635 | 0.00051 | 0 | 0.000896 | 0 | 0.000559 | 0 |
| k__Bacteria;p__Firmicutes;c__Clostridia;o__Clostridiales;f__Lachnospiraceae;g__NA;s__sp33734 | 0 | 0 | 0.004649 | 0.00067 | 0 | 0 | 0 | 0.001184 | 0.001095 | 0.001068 | 0 | 0 | 0 | 0.005999 | 0 | 0.026315 | 0.015383 |
| k__Bacteria;p__Firmicutes;c__Clostridia;o__Clostridiales;f__Lachnospiraceae;g__NA;s__sp33738 | 0 | 0 | 0 | 0 | 0 | 0 | 0 | 0 | 0 | 0 | 0 | 0 | 0 | 0 | 0.001337 | 0.001483 | 0.00072 |
| k__Bacteria;p__Firmicutes;c__Clostridia;o__Clostridiales;f__Lachnospiraceae;g__NA;s__sp33744 | 0 | 0.000211 | 0.000738 | 0 | 0 | 0 | 0 | 0 | 0 | 0 | 0 | 0 | 0 | 0.000747 | 0 | 0 | 0 |
| k__Bacteria;p__Firmicutes;c__Clostridia;o__Clostridiales;f__Lachnospiraceae;g__NA;s__sp33755 | 0.00074 | 0.000491 | 0 | 0.000915 | 0 | 0.000573 | 0.001949 | 0.001165 | 0.00146 | 0.002172 | 0.00272 | 0.002062 | 0 | 0.002838 | 0 | 0.001526 | 0.005008 |
| k__Bacteria;p__Firmicutes;c__Clostridia;o__Clostridiales;f__Lachnospiraceae;g__NA;s__sp33756 | 0 | 0 | 0 | 0 | 0 | 0 | 0.001247 | 0 | 0 | 0 | 0 | 0 | 0 | 0 | 0 | 0 | 0 |
| k__Bacteria;p__Firmicutes;c__Clostridia;o__Clostridiales;f__Lachnospiraceae;g__NA;s__sp33760 | 0.012511 | 0.022748 | 0.039777 | 0.008693 | 0.023337 | 0.007703 | 0.00512 | 0.00189 | 0.000712 | 0.001539 | 0 | 0.001242 | 0.000774 | 0.000971 | 0.000669 | 0.001247 | 0.001309 |
| k__Bacteria;p__Firmicutes;c__Clostridia;o__Clostridiales;f__Lachnospiraceae;g__NA;s__sp33766 | 0.001149 | 0.000772 | 0.012841 | 0.000931 | 0.004901 | 0.000618 | 0.007875 | 0.002692 | 0.00407 | 0.003077 | 0.005953 | 0.003304 | 0.009293 | 0.00692 | 0.004082 | 0.002408 | 0.007397 |
| k__Bacteria;p__Firmicutes;c__Clostridia;o__Clostridiales;f__Lachnospiraceae;g__NA;s__sp33767 | 0.000332 | 0.000281 | 0.002681 | 0.000343 | 0 | 0.000181 | 0.001066 | 0.000229 | 0.000237 | 0.000869 | 0 | 0.00071 | 0.001387 | 0.000921 | 0.000985 | 0.002322 | 0.003044 |
| k__Bacteria;p__Firmicutes;c__Clostridia;o__Clostridiales;f__Lachnospiraceae;g__Roseburia;s__sp33112 | 0 | 0 | 0.001378 | 0 | 0 | 0.000211 | 0.000728 | 0.000401 | 0.000584 | 0.000543 | 0.000604 | 0.001175 | 0 | 0 | 0 | 0.000344 | 0 |

| | | | | | | | | | | | | | | | | | |
|---|---|---|---|---|---|---|---|---|---|---|---|---|---|---|---|---|---|
| k__Bacteria;p__Firmicutes;c__Clostridia;o__Clostridiales;f__Lachnospiraceae;g__Roseburia;s__sp33143 | 0.0048 | 0.00316 | 0.012988 | 0.002925 | 0.006379 | 0.001884 | 0.002859 | 0.001127 | 0.002153 | 0.001738 | 0.002055 | 0.001685 | 0 | 0.001867 | 0 | 0.00058 | 0.00072 |
| k__Bacteria;p__Firmicutes;c__Clostridia;o__Clostridiales;f__Lachnospiraceae;g__Roseburia;s__sp33163 | 0 | 0 | 0.000541 | 0 | 0 | 0 | 0.00052 | 0 | 0.000255 | 0 | 0 | 0 | 0 | 0.000548 | 0 | 0.000408 | 0.000622 |
| k__Bacteria;p__Firmicutes;c__Clostridia;o__Clostridiales;f__Lachnospiraceae;g__Roseburia;s__sp33177 | 0 | 0 | 0 | 0 | 0 | 0 | 0.001949 | 0.000439 | 0.000748 | 0.000815 | 0.000725 | 0 | 0 | 0 | 0 | 0.000989 | 0 |
| k__Bacteria;p__Firmicutes;c__Clostridia;o__Clostridiales;f__Lachnospiraceae;g__Tyzzerella;s__sp33258-sp33301 | 0 | 0 | 0 | 0 | 0 | 0 | 0 | 0 | 0 | 0 | 0 | 0 | 0 | 0 | 0 | 0.000602 | 0 |
| k__Bacteria;p__Firmicutes;c__Clostridia;o__Clostridiales;f__Lachnospiraceae;g__Tyzzerella;s__sp33281 | 0.000613 | 0 | 0 | 0 | 0 | 0 | 0 | 0 | 0 | 0 | 0 | 0 | 0 | 0.000747 | 0 | 0 | 0 |
| k__Bacteria;p__Firmicutes;c__Clostridia;o__Clostridiales;f__Lachnospiraceae;g__Tyzzerella;s__sp33287 | 0 | 0 | 0.001205 | 0 | 0 | 0 | 0.001144 | 0.000306 | 0 | 0.000217 | 0 | 0 | 0.000484 | 0 | 0 | 0.000387 | 0 |
| k__Bacteria;p__Firmicutes;c__Clostridia;o__Clostridiales;f__Lachnospiraceae;g__Tyzzerella;s__sp33289-sp33291 | 0 | 0 | 0.000443 | 0 | 0 | 0 | 0.00039 | 0 | 0 | 0 | 0 | 0 | 0 | 0 | 0 | 0 | 0 |
| k__Bacteria;p__Firmicutes;c__Clostridia;o__Clostridiales;f__NA;g__NA;s__sp31072 | 0.000383 | 0 | 0.001058 | 0 | 0 | 0.000166 | 0.004106 | 0.001108 | 0.000511 | 0.000869 | 0 | 0.000732 | 0 | 0.001444 | 0 | 0 | 0 |
| k__Bacteria;p__Firmicutes;c__Clostridia;o__Clostridiales;f__NA;g__NA;s__sp31075 | 0.001532 | 0.000755 | 0.001574 | 0.001193 | 0 | 0.001387 | 0.000416 | 0.000802 | 0.000274 | 0.000652 | 0 | 0.001375 | 0 | 0 | 0 | 0 | 0 |
| k__Bacteria;p__Firmicutes;c__Clostridia;o__Clostridiales;f__NA;g__NA;s__sp31082 | 0 | 0.001562 | 0.001082 | 0.000376 | 0 | 0.000513 | 0.00395 | 0.00296 | 0.00042 | 0.000543 | 0 | 0.003526 | 0 | 0.000846 | 0 | 0.000258 | 0 |
| k__Bacteria;p__Firmicutes;c__Clostridia;o__Clostridiales;f__NA;g__NA;s__sp31094 | 0.001328 | 0.016394 | 0 | 0.006487 | 0 | 0.007764 | 0 | 0 | 0 | 0 | 0 | 0 | 0 | 0 | 0 | 0 | 0 |
| k__Bacteria;p__Firmicutes;c__Clostridia;o__Clostridiales;f__NA;g__NA;s__sp31106 | 0 | 0 | 0 | 0 | 0 | 0 | 0 | 0 | 0 | 0 | 0 | 0 | 0.000516 | 0 | 0 | 0 | 0 |
| k__Bacteria;p__Firmicutes;c__Clostridia;o__Clostridiales;f__NA;g__NA;s__sp31115 | 0.002375 | 0 | 0 | 0.001209 | 0 | 0.000844 | 0.004756 | 0 | 0 | 0.002407 | 0 | 0.00224 | 0.016682 | 0.010753 | 0.008587 | 0.004106 | 0.00504 |
| k__Bacteria;p__Firmicutes;c__Clostridia;o__Clostridiales;f__NA;g__NA;s__sp31116 | 0 | 0 | 0 | 0 | 0 | 0 | 0 | 0.000573 | 0 | 0 | 0 | 0 | 0 | 0 | 0 | 0.000258 | 0 |
| k__Bacteria;p__Firmicutes;c__Clostridia;o__Clostridiales;f__NA;g__NA;s__sp31119 | 0.003677 | 0 | 0 | 0.001438 | 0 | 0.001236 | 0.003586 | 0.004946 | 0.000584 | 0.002353 | 0.001541 | 0.003504 | 0 | 0.00336 | 0.002182 | 0.00086 | 0.000818 |
| k__Bacteria;p__Firmicutes;c__Clostridia;o__Clostridiales;f__NA;g__NA;s__sp31125 | 0 | 0 | 0.005805 | 0 | 0 | 0 | 0.001637 | 0 | 0.00042 | 0.000235 | 0.000997 | 0.00173 | 0 | 0 | 0 | 0 | 0 |
| k__Bacteria;p__Firmicutes;c__Clostridia;o__Clostridiales;f__Peptococcaceae;g__NA;s__sp34196 | 0 | 0.000193 | 0 | 0.000196 | 0 | 0 | 0 | 0.000344 | 0 | 0.00038 | 0 | 0.000288 | 0 | 0 | 0 | 0 | 0 |
| k__Bacteria;p__Firmicutes;c__Clostridia;o__Clostridiales;f__Peptococcaceae;g__NA;s__sp34246 | 0.000536 | 0.000878 | 0 | 0.00018 | 0 | 0 | 0.001247 | 0.000344 | 0.00031 | 0.000235 | 0 | 0 | 0.001194 | 0.001593 | 0 | 0.001053 | 0.001571 |
| k__Bacteria;p__Firmicutes;c__Clostridia;o__Clostridiales;f__Peptococcaceae;g__Peptococcus;s__sp34117 | 0.001226 | 0.000527 | 0.001968 | 0.000605 | 0 | 0 | 0.000546 | 0 | 0 | 0 | 0 | 0.00031 | 0.000807 | 0.000348 | 0 | 0.000602 | 0 |
| k__Bacteria;p__Firmicutes;c__Clostridia;o__Clostridiales;f__Ruminococcaceae;g__Anaerotruncus;s__colihominis | 0 | 0 | 0 | 0 | 0 | 0 | 0.000598 | 0.000401 | 0.000274 | 0.000272 | 0 | 0 | 0.003227 | 0.000971 | 0 | 0.003203 | 0.003011 |
| k__Bacteria;p__Firmicutes;c__Clostridia;o__Clostridiales;f__Ruminococcaceae;g__Anaerotruncus;s__sp34440-sp34450-sp34472 | 0.001762 | 0.001931 | 0 | 0.001095 | 0 | 0.000889 | 0.001066 | 0.000344 | 0 | 0.001683 | 0.000755 | 0.000488 | 0 | 0 | 0 | 0 | 0 |
| k__Bacteria;p__Firmicutes;c__Clostridia;o__Clostridiales;f__Ruminococcaceae;g__Anaerotruncus;s__sp34443 | 0 | 0 | 0 | 0 | 0 | 0 | 0.00039 | 0 | 0 | 0 | 0 | 0 | 0 | 0 | 0 | 0 | 0 |
| k__Bacteria;p__Firmicutes;c__Clostridia;o__Clostridiales;f__Ruminococcaceae;g__Anaerotruncus;s__sp34452-sp34475 | 0 | 0.000579 | 0.001968 | 0.000359 | 0 | 0.000528 | 0.003379 | 0.002005 | 0.001734 | 0.001665 | 0.003112 | 0.002018 | 0 | 0.000946 | 0 | 0.000795 | 0 |
| k__Bacteria;p__Firmicutes;c__Clostridia;o__Clostridiales;f__Ruminococcaceae;g__Anaerotruncus;s__sp34453 | 0.002247 | 0.001703 | 0.009397 | 0.001291 | 0 | 0.001236 | 0.006133 | 0.003036 | 0.001022 | 0.002607 | 0.002629 | 0.003149 | 0.003001 | 0.004854 | 0.00183 | 0.006235 | 0.004157 |
| k__Bacteria;p__Firmicutes;c__Clostridia;o__Clostridiales;f__Ruminococcaceae;g__Anaerotruncus;s__sp34455 | 0 | 0.00072 | 0 | 0 | 0 | 0 | 0.001221 | 0 | 0 | 0 | 0 | 0 | 0 | 0 | 0 | 0 | 0 |
| k__Bacteria;p__Firmicutes;c__Clostridia;o__Clostridiales;f__Ruminococcaceae;g__Anaerotruncus;s__sp34462 | 0 | 0 | 0 | 0 | 0 | 0 | 0 | 0 | 0 | 0 | 0 | 0 | 0 | 0.001444 | 0 | 0.00058 | 0 |
| k__Bacteria;p__Firmicutes;c__Clostridia;o__Clostridiales;f__Ruminococcaceae;g__Anaerotruncus;s__sp34466 | 0 | 0 | 0 | 0 | 0 | 0.000392 | 0.000936 | 0 | 0 | 0 | 0 | 0.00031 | 0.001871 | 0.001369 | 0 | 0.001139 | 0 |
| k__Bacteria;p__Firmicutes;c__Clostridia;o__Clostridiales;f__Ruminococcaceae;g__Anaerotruncus;s__sp34466-sp34475 | 0 | 0 | 0 | 0 | 0 | 0 | 0 | 0.000382 | 0 | 0 | 0 | 0 | 0 | 0 | 0 | 0 | 0 |
| k__Bacteria;p__Firmicutes;c__Clostridia;o__Clostridiales;f__Ruminococcaceae;g__Anaerotruncus;s__sp34471 | 0 | 0 | 0.00305 | 0 | 0.001478 | 0 | 0.001351 | 0.000382 | 0.000237 | 0.000561 | 0 | 0 | 0.002549 | 0.002066 | 0.001126 | 0.000946 | 0.001309 |
| k__Bacteria;p__Firmicutes;c__Clostridia;o__Clostridiales;f__Ruminococcaceae;g__Anaerotruncus;s__sp34475 | 0 | 0.000825 | 0.003149 | 0.000768 | 0 | 0 | 0.001014 | 0.000458 | 0 | 0.000525 | 0 | 0 | 0 | 0.000921 | 0 | 0.000752 | 0.003535 |
| k__Bacteria;p__Firmicutes;c__Clostridia;o__Clostridiales;f__Ruminococcaceae;g__Anaerotruncus;s__sp34481 | 0 | 0 | 0.000738 | 0 | 0 | 0 | 0.000494 | 0 | 0 | 0.000217 | 0 | 0 | 0 | 0.000324 | 0 | 0.000451 | 0 |
| k__Bacteria;p__Firmicutes;c__Clostridia;o__Clostridiales;f__Ruminococcaceae;g__NA;s__sp33140 | 0 | 0 | 0.000787 | 0 | 0 | 0 | 0.001637 | 0.000363 | 0.000383 | 0.000579 | 0.000786 | 0.000355 | 0.002162 | 0.001693 | 0.001478 | 0.001032 | 0.001506 |
| k__Bacteria;p__Firmicutes;c__Clostridia;o__Clostridiales;f__Ruminococcaceae;g__NA;s__sp34771 | 0.006868 | 0.000614 | 0.005879 | 0.002092 | 0 | 0.001251 | 0.010292 | 0.001871 | 0.00104 | 0.002969 | 0 | 0.002484 | 0.005066 | 0.002713 | 0 | 0.005504 | 0.005269 |
| k__Bacteria;p__Firmicutes;c__Clostridia;o__Clostridiales;f__Ruminococcaceae;g__NA;s__sp34795 | 0 | 0 | 0.000935 | 0 | 0 | 0 | 0 | 0 | 0 | 0 | 0 | 0 | 0 | 0.001444 | 0 | 0 | 0 |
| k__Bacteria;p__Firmicutes;c__Clostridia;o__Clostridiales;f__Ruminococcaceae;g__NA;s__sp34795-sp34820 | 0 | 0.000439 | 0.000689 | 0 | 0 | 0 | 0 | 0 | 0.000639 | 0 | 0 | 0 | 0 | 0.000672 | 0 | 0.001075 | 0 |
| k__Bacteria;p__Firmicutes;c__Clostridia;o__Clostridiales;f__Ruminococcaceae;g__NA;s__sp34820 | 0 | 0 | 0 | 0.000359 | 0 | 0 | 0 | 0 | 0 | 0 | 0 | 0 | 0 | 0.001742 | 0 | 0.000731 | 0 |
| k__Bacteria;p__Firmicutes;c__Clostridia;o__Clostridiales;f__Ruminococcaceae;g__NA;s__sp34822 | 0 | 0 | 0 | 0 | 0 | 0 | 0 | 0 | 0 | 0 | 0 | 0 | 0 | 0 | 0 | 0.000946 | 0.000491 |
| k__Bacteria;p__Firmicutes;c__Clostridia;o__Clostridiales;f__Ruminococcaceae;g__NA;s__sp34838 | 0 | 0 | 0 | 0 | 0 | 0 | 0.001871 | 0.000726 | 0 | 0 | 0 | 0 | 0.001613 | 0.001245 | 0.00183 | 0.001376 | 0.001178 |
| k__Bacteria;p__Firmicutes;c__Clostridia;o__Clostridiales;f__Ruminococcaceae;g__NA;s__sp34858-sp34873 | 0 | 0 | 0.002509 | 0 | 0 | 0 | 0.002209 | 0 | 0 | 0 | 0 | 0.000976 | 0 | 0.003883 | 0 | 0.002408 | 0.004124 |
| k__Bacteria;p__Firmicutes;c__Clostridia;o__Clostridiales;f__Ruminococcaceae;g__NA;s__sp34863 | 0 | 0 | 0 | 0 | 0 | 0 | 0 | 0 | 0 | 0 | 0 | 0 | 0 | 0 | 0 | 0.000451 | 0 |
| k__Bacteria;p__Firmicutes;c__Clostridia;o__Clostridiales;f__Ruminococcaceae;g__NA;s__sp34871 | 0.0036 | 0.000632 | 0.002903 | 0.001797 | 0 | 0.000995 | 0.001663 | 0.003743 | 0 | 0 | 0 | 0 | 0 | 0.00122 | 0 | 0 | 0 |
| k__Bacteria;p__Firmicutes;c__Clostridia;o__Clostridiales;f__Ruminococcaceae;g__NA;s__sp34871-sp34878-sp34883 | 0 | 0 | 0 | 0 | 0 | 0 | 0 | 0 | 0 | 0 | 0.001239 | 0 | 0 | 0 | 0 | 0.001354 | 0 |
| k__Bacteria;p__Firmicutes;c__Clostridia;o__Clostridiales;f__Ruminococcaceae;g__NA;s__sp34878 | 0 | 0.001036 | 0.001796 | 0.000359 | 0 | 0.00104 | 0.004782 | 0 | 0.001095 | 0.000959 | 0.001148 | 0.000643 | 0.00997 | 0.003335 | 0.004329 | 0.00488 | 0.006906 |
| k__Bacteria;p__Firmicutes;c__Clostridia;o__Clostridiales;f__Ruminococcaceae;g__NA;s__sp34879 | 0.000306 | 0.000983 | 0.002263 | 0.000686 | 0 | 0.000663 | 0.002729 | 0.001394 | 0.001058 | 0.001611 | 0 | 0.002706 | 0 | 0.00224 | 0 | 0 | 0 |
| k__Bacteria;p__Firmicutes;c__Clostridia;o__Clostridiales;f__Ruminococcaceae;g__NA;s__sp34883 | 0 | 0.002773 | 0 | 0 | 0 | 0 | 0 | 0 | 0 | 0 | 0 | 0 | 0 | 0 | 0 | 0 | 0 |
| k__Bacteria;p__Firmicutes;c__Clostridia;o__Clostridiales;f__Ruminococcaceae;g__NA;s__sp34977 | 0 | 0.000807 | 0 | 0 | 0 | 0 | 0 | 0 | 0 | 0 | 0 | 0 | 0 | 0.000274 | 0.00088 | 0 | 0 |
| k__Bacteria;p__Firmicutes;c__Clostridia;o__Clostridiales;f__Ruminococcaceae;g__NA;s__sp34983 | 0 | 0 | 0.000394 | 0 | 0 | 0 | 0.000676 | 0 | 0 | 0 | 0 | 0 | 0 | 0.000398 | 0 | 0 | 0 |
| k__Bacteria;p__Firmicutes;c__Clostridia;o__Clostridiales;f__Ruminococcaceae;g__NA;s__sp35077 | 0 | 0 | 0 | 0 | 0 | 0 | 0 | 0 | 0 | 0.000344 | 0 | 0 | 0 | 0 | 0.001091 | 0 | 0 |
| k__Bacteria;p__Firmicutes;c__Clostridia;o__Clostridiales;f__Ruminococcaceae;g__NA;s__sp35181 | 0 | 0 | 0 | 0 | 0 | 0 | 0 | 0 | 0 | 0 | 0 | 0 | 0 | 0.001543 | 0 | 0 | 0 |
| k__Bacteria;p__Firmicutes;c__Clostridia;o__Clostridiales;f__Ruminococcaceae;g__NA;s__sp35382-sp35403-sp35432 | 0 | 0 | 0 | 0 | 0 | 0 | 0.000572 | 0.000229 | 0.000255 | 0 | 0 | 0.00122 | 0 | 0 | 0 | 0 | 0 |
| k__Bacteria;p__Firmicutes;c__Clostridia;o__Clostridiales;f__Ruminococcaceae;g__NA;s__sp35403 | 0 | 0 | 0 | 0 | 0 | 0 | 0 | 0 | 0 | 0 | 0 | 0 | 0 | 0.000473 | 0 | 0 | 0 |
| k__Bacteria;p__Firmicutes;c__Clostridia;o__Clostridiales;f__Ruminococcaceae;g__NA;s__sp35688-sp35748 | 0.000868 | 0.000211 | 0.003075 | 0.000343 | 0 | 0 | 0.001092 | 0 | 0.00031 | 0.000326 | 0 | 0 | 0.002485 | 0 | 0 | 0.00086 | 0.001309 |
| k__Bacteria;p__Firmicutes;c__Clostridia;o__Clostridiales;f__Ruminococcaceae;g__NA;s__sp35693 | 0.005387 | 0.00079 | 0.013062 | 0.002141 | 0.008246 | 0.001146 | 0.006705 | 0.002272 | 0.002445 | 0.001955 | 0.003807 | 0.002173 | 0.009486 | 0.005103 | 0.006124 | 0.008707 | 0.013943 |
| k__Bacteria;p__Firmicutes;c__Clostridia;o__Clostridiales;f__Ruminococcaceae;g__NA;s__sp35693-sp35877 | 0 | 0 | 0 | 0 | 0 | 0 | 0 | 0 | 0 | 0 | 0 | 0 | 0 | 0 | 0 | 0 | 0.001636 |
| k__Bacteria;p__Firmicutes;c__Clostridia;o__Clostridiales;f__Ruminococcaceae;g__NA;s__sp35696 | 0.003575 | 0.006688 | 0.014341 | 0.003088 | 0 | 0.001794 | 0.012241 | 0.003303 | 0.002372 | 0.00295 | 0.005379 | 0.002883 | 0 | 0.000996 | 0 | 0.00187 | 0 |
| k__Bacteria;p__Firmicutes;c__Clostridia;o__Clostridiales;f__Ruminococcaceae;g__NA;s__sp35730 | 0 | 0 | 0.000836 | 0 | 0 | 0 | 0.001897 | 0 | 0.000328 | 0.000869 | 0.000786 | 0 | 0.006356 | 0.003335 | 0.000845 | 0.003526 | 0.00792 |
| k__Bacteria;p__Firmicutes;c__Clostridia;o__Clostridiales;f__Ruminococcaceae;g__NA;s__sp35733 | 0.000587 | 0.001194 | 0.00182 | 0.000572 | 0 | 0.000317 | 0.001533 | 0.000745 | 0 | 0.000706 | 0 | 0.000244 | 0.001549 | 0.002713 | 0.001056 | 0.001204 | 0.002258 |
| k__Bacteria;p__Firmicutes;c__Clostridia;o__Clostridiales;f__Ruminococcaceae;g__NA;s__sp35736 | 0.000562 | 0.000474 | 0.001082 | 0.000327 | 0 | 0 | 0.001118 | 0.000439 | 0.00042 | 0 | 0 | 0.000488 | 0 | 0.00117 | 0 | 0.001118 | 0.001146 |
| k__Bacteria;p__Firmicutes;c__Clostridia;o__Clostridiales;f__Ruminococcaceae;g__NA;s__sp35748 | 0 | 0 | 0 | 0 | 0 | 0 | 0.001118 | 0 | 0 | 0 | 0 | 0 | 0.004227 | 0.004679 | 0.007531 | 0.003934 | 0.00432 |
| k__Bacteria;p__Firmicutes;c__Clostridia;o__Clostridiales;f__Ruminococcaceae;g__NA;s__sp35799 | 0.00097 | 0.000211 | 0.002165 | 0.000425 | 0 | 0.000663 | 0.002105 | 0.000229 | 0.000328 | 0.000344 | 0 | 0.000399 | 0.004162 | 0.002041 | 0.00088 | 0.003311 | 0.002651 |
| k__Bacteria;p__Firmicutes;c__Clostridia;o__Clostridiales;f__Ruminococcaceae;g__NA;s__sp35825 | 0 | 0 | 0 | 0 | 0 | 0 | 0.000312 | 0 | 0 | 0 | 0 | 0 | 0 | 0 | 0 | 0 | 0 |
| k__Bacteria;p__Firmicutes;c__Clostridia;o__Clostridiales;f__Ruminococcaceae;g__NA;s__sp35832 | 0 | 0 | 0 | 0 | 0 | 0 | 0 | 0.001222 | 0 | 0 | 0 | 0.000643 | 0.001549 | 0.001469 | 0.000985 | 0.001075 | 0.002356 |
| k__Bacteria;p__Firmicutes;c__Clostridia;o__Clostridiales;f__Ruminococcaceae;g__NA;s__sp35841 | 0.005668 | 0 | 0.001993 | 0.002043 | 0 | 0.000528 | 0.003379 | 0.001184 | 0.001277 | 0.001593 | 0.002448 | 0.00122 | 0.003549 | 0.004854 | 0 | 0.002623 | 0.003306 |
| k__Bacteria;p__Firmicutes;c__Clostridia;o__Clostridiales;f__Ruminococcaceae;g__NA;s__sp35849 | 0.000281 | 0 | 0.000615 | 0 | 0 | 0 | 0.001377 | 0.000325 | 0 | 0.000235 | 0 | 0 | 0 | 0 | 0 | 0 | 0.000884 |
| k__Bacteria;p__Firmicutes;c__Clostridia;o__Clostridiales;f__Ruminococcaceae;g__NA;s__sp35869 | 0.002783 | 0.001632 | 0.007281 | 0.001078 | 0 | 0.000814 | 0 | 0 | 0.00031 | 0.001068 | 0 | 0 | 0.001904 | 0.002514 | 0 | 0.001892 | 0.002487 |
| k__Bacteria;p__Firmicutes;c__Clostridia;o__Clostridiales;f__Ruminococcaceae;g__NA;s__sp35873 | 0 | 0 | 0 | 0 | 0 | 0 | 0 | 0 | 0 | 0 | 0 | 0 | 0 | 0.000473 | 0 | 0 | 0 |
| k__Bacteria;p__Firmicutes;c__Clostridia;o__Clostridiales;f__Ruminococcaceae;g__NA;s__sp35877 | 0.000638 | 0 | 0 | 0 | 0 | 0 | 0.000754 | 0.000707 | 0 | 0.000507 | 0 | 0 | 0.001484 | 0.001045 | 0 | 0.000731 | 0 |
| k__Bacteria;p__Firmicutes;c__Clostridia;o__Clostridiales;f__Ruminococcaceae;g__NA;s__sp35882 | 0.002145 | 0.002159 | 0.00615 | 0.001471 | 0 | 0.001146 | 0.008446 | 0.003514 | 0.000858 | 0.002588 | 0.002992 | 0.00051 | 0.002259 | 0.003136 | 0.000598 | 0.002429 | 0.003142 |
| k__Bacteria;p__Firmicutes;c__Clostridia;o__Clostridiales;f__Ruminococcaceae;g__NA;s__sp35894 | 0.001685 | 0.000562 | 0.017785 | 0.001177 | 0.014391 | 0.000739 | 0.008914 | 0.005538 | 0.00104 | 0.002607 | 0.003173 | 0.000798 | 0 | 0.00107 | 0 | 0.004708 | 0.00252 |
| k__Bacteria;p__Firmicutes;c__Clostridia;o__Clostridiales;f__Ruminococcaceae;g__Oscillibacter;s__sp34634 | 0.016673 | 0.008355 | 0.03867 | 0.008334 | 0.027071 | 0.003 | 0.003093 | 0.001127 | 0.00135 | 0.002263 | 0.002417 | 0.000444 | 0.012197 | 0.010927 | 0.005525 | 0.016683 | 0.016626 |
| k__Bacteria;p__Firmicutes;c__Clostridia;o__Clostridiales;f__Ruminococcaceae;g__Oscillibacter;s__sp34638 | 0 | 0 | 0 | 0 | 0 | 0 | 0 | 0 | 0 | 0 | 0 | 0 | 0 | 0 | 0 | 0.000989 | 0.001015 |
| k__Bacteria;p__Firmicutes;c__Clostridia;o__Clostridiales;f__Ruminococcaceae;g__Oscillibacter;s__sp34640 | 0 | 0.000369 | 0 | 0 | 0 | 0 | 0.002885 | 0 | 0.001131 | 0 | 0 | 0.001819 | 0.002065 | 0.001195 | 0.003273 | 0.00215 | 0.002586 |
| k__Bacteria;p__Firmicutes;c__Clostridia;o__Clostridiales;f__Ruminococcaceae;g__Oscillibacter;s__sp34642 | 0 | 0 | 0 | 0 | 0 | 0 | 0 | 0.000974 | 0 | 0.000796 | 0 | 0 | 0.002226 | 0.002987 | 0 | 0.003096 | 0.002193 |
| k__Bacteria;p__Firmicutes;c__Clostridia;o__Clostridiales;f__Ruminococcaceae;g__Oscillibacter;s__sp34643 | 0.001813 | 0.000439 | 0.004108 | 0.000588 | 0 | 0.000211 | 0.006731 | 0.000496 | 0.000401 | 0.001466 | 0.001964 | 0 | 0.000871 | 0.00112 | 0 | 0.000559 | 0.000524 |
| k__Bacteria;p__Firmicutes;c__Clostridia;o__Clostridiales;f__Ruminococcaceae;g__Oscillibacter;s__sp34645 | 0.001634 | 0.00079 | 0.00123 | 0.000833 | 0 | 0.000377 | 0.000832 | 0 | 0 | 0.000507 | 0 | 0 | 0 | 0.000373 | 0 | 0.000559 | 0.000622 |
| k__Bacteria;p__Firmicutes;c__Clostridia;o__Clostridiales;f__Ruminococcaceae;g__Oscillibacter;s__sp34648 | 0 | 0 | 0.001599 | 0 | 0 | 0 | 0.002287 | 0 | 0 | 0 | 0 | 0 | 0 | 0 | 0 | 0 | 0.001506 |
| k__Bacteria;p__Firmicutes;c__Clostridia;o__Clostridiales;f__Ruminococcaceae;g__Oscillibacter-Oscillospira;s__sp34648-sp34660 | 0 | 0 | 0 | 0 | 0 | 0 | 0 | 0 | 0 | 0 | 0 | 0 | 0 | 0 | 0 | 0 | 0.00036 |
| k__Bacteria;p__Firmicutes;c__Clostridia;o__Clostridiales;f__Ruminococcaceae;g__Ruminiclostridium;s__sp34857-sp34950 | 0.006204 | 0 | 0.009766 | 0.002386 | 0 | 0.002231 | 0.002131 | 0 | 0.001551 | 0.001466 | 0.002115 | 0 | 0 | 0 | 0 | 0 | 0 |
| k__Bacteria;p__Firmicutes;c__Clostridia;o__Clostridiales;f__Ruminococcaceae;g__Ruminiclostridium;s__sp34901 | 0.001506 | 0 | 0.00246 | 0.000686 | 0 | 0 | 0 | 0 | 0 | 0 | 0 | 0 | 0 | 0.000921 | 0 | 0 | 0 |
| k__Bacteria;p__Firmicutes;c__Clostridia;o__Clostridiales;f__Ruminococcaceae;g__Ruminiclostridium;s__sp34925 | 0 | 0 | 0.007995 | 0.000278 | 0 | 0.000482 | 0.001273 | 0 | 0.001478 | 0.000525 | 0.002055 | 0.000444 | 0.010454 | 0.003435 | 0.003027 | 0.013759 | 0.01103 |

| | | | | | | | | | | | | | | | | | |
|---|---|---|---|---|---|---|---|---|---|---|---|---|---|---|---|---|---|
| k__Bacteria;p__Firmicutes;c__Clostridia;o__Clostridiales;f__Ruminococcaceae;g__Ruminiclostridium;s__sp34950 | 0 | 0 | 0.001476 | 0 | 0 | 0 | 0 | 0 | 0 | 0 | 0 | 0 | 0.004001 | 0 | 0.001584 | 0.001462 | 0 |
| k__Bacteria;p__Firmicutes;c__Erysipelotrichia;o__Erysipelotrichales;f__Erysipelotrichaceae;g__Allobaculum;s__sp36555 | 0 | 0 | 0 | 0 | 0 | 0 | 0 | 0 | 0 | 0 | 0 | 0 | 0.007421 | 0 | 0.002112 | 0.033495 | 0.012142 |
| k__Bacteria;p__Firmicutes;c__Erysipelotrichia;o__Erysipelotrichales;f__Erysipelotrichaceae;g__NA;s__sp36777 | 0 | 0 | 0 | 0 | 0 | 0 | 0 | 0 | 0 | 0 | 0 | 0 | 0 | 0.000572 | 0 | 0 | 0 |
| k__Bacteria;p__Firmicutes;c__Erysipelotrichia;o__Erysipelotrichales;f__Erysipelotrichaceae;g__NA;s__sp36783 | 0 | 0 | 0 | 0 | 0 | 0 | 0 | 0 | 0.000365 | 0.000344 | 0 | 0 | 0 | 0 | 0 | 0 | 0 |
| k__Bacteria;p__Firmicutes;c__Erysipelotrichia;o__Erysipelotrichales;f__Erysipelotrichaceae;g__NA;s__sp36787 | 0 | 0 | 0 | 0 | 0 | 0.000166 | 0 | 0.00042 | 0.000292 | 0.000724 | 0.000574 | 0 | 0 | 0 | 0 | 0 | 0 |
| k__Bacteria;p__Firmicutes;c__Erysipelotrichia;o__Erysipelotrichales;f__Erysipelotrichaceae;g__Turicibacter;s__sanguinis | 0.005924 | 0.015446 | 0.007527 | 0.008317 | 0.045585 | 0.007839 | 0 | 0 | 0 | 0 | 0 | 0 | 0 | 0 | 0 | 0 | 0 |
| k__Bacteria;p__Proteobacteria;c__Alphaproteobacteria;o__Caulobacterales;f__Caulobacteraceae;g__Caulobacter;s__sp42762 | 0 | 0 | 0 | 0.000196 | 0 | 0 | 0 | 0 | 0 | 0 | 0 | 0 | 0 | 0 | 0 | 0 | 0 |
| k__Bacteria;p__Proteobacteria;c__Alphaproteobacteria;o__Rhodospirillales;f__Rhodospirillaceae;g__Thalassospira;s__sp46235 | 0 | 0 | 0 | 0 | 0 | 0 | 0 | 0 | 0.000292 | 0 | 0 | 0.003859 | 0.005195 | 0.002116 | 0.00644 | 0.00086 | 0.000589 |
| k__Bacteria;p__Proteobacteria;c__Alphaproteobacteria;o__Rickettsiales;f__Mitochondria;g__Arundo-Oryza meyeriana;s__NA | 0 | 0 | 0 | 0 | 0 | 0 | 0 | 0.000344 | 0 | 0.000398 | 0 | 0.00031 | 0 | 0 | 0 | 0 | 0 |
| k__Bacteria;p__Proteobacteria;c__Betaproteobacteria;o__Burkholderiales;f__Alcaligenaceae;g__Parasutterella;s__excrementihominis | 0 | 0 | 0 | 0 | 0 | 0 | 0 | 0 | 0 | 0.000471 | 0 | 0 | 0 | 0 | 0 | 0 | 0 |
| k__Bacteria;p__Proteobacteria;c__Betaproteobacteria;o__Burkholderiales;f__Alcaligenaceae;g__Parasutterella;s__sp48235 | 0.002247 | 0.005933 | 0 | 0.002811 | 0 | 0.001794 | 0.00065 | 0.001509 | 0.003814 | 0.001683 | 0.002236 | 0.002196 | 0.004227 | 0.005551 | 0.004188 | 0.007869 | 0.003502 |
| k__Bacteria;p__Proteobacteria;c__Deltaproteobacteria;o__Desulfovibrionales;f__Desulfovibrionaceae;g__Bilophila;s__sp52475 | 0.007915 | 0.005318 | 0.014735 | 0.00317 | 0 | 0.001055 | 0.007953 | 0.00147 | 0.001223 | 0.001991 | 0.002145 | 0.001375 | 0.004324 | 0.007168 | 0.001513 | 0.007611 | 0.004549 |
| k__Bacteria;p__Proteobacteria;c__Gammaproteobacteria;o__Enterobacteriales;f__Enterobacteriaceae;g__Escherichia-Shigella;s__coli | 0.001787 | 0.00158 | 0.00428 | 0.003938 | 0.023882 | 0.003 | 0 | 0.000592 | 0 | 0.000597 | 0 | 0.000998 | 0 | 0 | 0 | 0 | 0 |
| k__Bacteria;p__Tenericutes;c__Mollicutes;o__Anaeroplasmatales;f__Anaeroplasmataceae;g__Anaeroplasma;s__sp67615 | 0 | 0 | 0 | 0 | 0 | 0 | 0 | 0 | 0 | 0 | 0 | 0 | 0 | 0.002439 | 0 | 0 | 0 |
