## Supplementary material for "The gut microbiota of environmentally enriched mice regulates visual cortical plasticity": Suppl. Table 2

| SampleID | P20.EE1 | P20.EE2 | P20.EE8 | P20.EE12 | P20.EE13 | P20.EE15 | P25.EE1 | P25.EE2 | P25.EE8 | P25.EE12 | P25.EE13 | P25.EE15 | P90.EE1 | P90.EE2 | P90.EE8 | P90.EE12 | P90.EE13 | P90.EE15 |
|---|---|---|---|---|---|---|---|---|---|---|---|---|---|---|---|---|---|---|
| k__Bacteria;p__Actinobacteria;c__Actinobacteria;o__Bifidobacteriales;f__Bifidobacteriaceae;g__Bifidobacterium;s__pseudolongum | 0.000458 | 0.002215 | 0 | 0 | 0 | 0 | 0 | 0.008879 | 0 | 0 | 0 | 0 | 0.001288 | 0.008697 | 0.001359 | 0 | 0.000673 | 0.0008 |
| k__Bacteria;p__Actinobacteria;c__Actinobacteria;o__Corynebacteriales;f__Corynebacteriaceae;g__Corynebacterium;s__stationis | 0 | 0 | 0 | 0 | 0 | 0 | 0 | 0 | 0 | 0 | 0 | 0 | 0.000858 | 0 | 0 | 0 | 0 | 0 |
| k__Bacteria;p__Actinobacteria;c__Coriobacteriia;o__Coriobacteriales;f__Coriobacteriaceae;g__Collinsella;s__aerofaciens | 0 | 0 | 0 | 0 | 0 | 0 | 0 | 0 | 0 | 0 | 0 | 0 | 0 | 0 | 0.003471 | 0 | 0 | 0 |
| k__Bacteria;p__Actinobacteria;c__Coriobacteriia;o__Coriobacteriales;f__Coriobacteriaceae;g__Enterorhabdus;s__caecimuris | 0.000526 | 0 | 0 | 0 | 0 | 0 | 0 | 0.000608 | 0 | 0 | 0 | 0 | 0 | 0 | 0 | 0 | 0 | 0.000702 |
| k__Bacteria;p__Actinobacteria;c__Coriobacteriia;o__Coriobacteriales;f__Coriobacteriaceae;g__Olsenella;s__profusa | 0.000696 | 0.004118 | 0 | 0.002517 | 0 | 0 | 0 | 0.000314 | 0.000758 | 0.000267 | 0 | 0 | 0.000634 | 0.002209 | 0 | 0 | 0 | 0 |
| k__Bacteria;p__Bacteroidetes;c__Bacteroidia;o__Bacteroidales;f__Bacteroidaceae;g__Bacteroides;s__acidifaciens | 0.072743 | 0.042129 | 0.007922 | 0.013715 | 0.011055 | 0.005792 | 0.004759 | 0.006056 | 0.009128 | 0.009973 | 0.021722 | 0.050432 | 0.00607 | 0.001977 | 0 | 0.005508 | 0.003591 | 0.004797 |
| k__Bacteria;p__Bacteroidetes;c__Bacteroidia;o__Bacteroidales;f__Bacteroidaceae;g__Bacteroides;s__oleiciplenus-rodentium | 0.343041 | 0.319875 | 0.054308 | 0.284935 | 0.023212 | 0.051359 | 0.06348 | 0.104763 | 0.006365 | 0.006665 | 0.057413 | 0.07025 | 0.037808 | 0.013394 | 0.015581 | 0.017609 | 0.025363 | 0.016088 |
| k__Bacteria;p__Bacteroidetes;c__Bacteroidia;o__Bacteroidales;f__Bacteroidaceae;g__Bacteroides;s__stercorirosoris | 0 | 0 | 0 | 0 | 0 | 0 | 0 | 0 | 0 | 0 | 0 | 0 | 0.001492 | 0.000744 | 0.001796 | 0.005883 | 0.001543 | 0.002496 |
| k__Bacteria;p__Bacteroidetes;c__Bacteroidia;o__Bacteroidales;f__Bacteroidaceae;g__Bacteroides;s__vulgatus | 0 | 0 | 0 | 0 | 0 | 0 | 0 | 0 | 0 | 0 | 0 | 0 | 0 | 0 | 0.0025 | 0 | 0 | 0 |
| k__Bacteria;p__Bacteroidetes;c__Bacteroidia;o__Bacteroidales;f__NA;g__NA;s__sp12473-sp12526 | 0 | 0 | 0 | 0 | 0 | 0 | 0 | 0 | 0 | 0.001153 | 0 | 0 | 0.003924 | 0.001465 | 0.001408 | 0.006613 | 0.004517 | 0.008795 |
| k__Bacteria;p__Bacteroidetes;c__Bacteroidia;o__Bacteroidales;f__NA;g__NA;s__sp12475-sp12557 | 0.140343 | 0.143368 | 0.091901 | 0.04465 | 0.122457 | 0.08027 | 0.015339 | 0.039436 | 0.114386 | 0.116821 | 0.154162 | 0.226468 | 0.047843 | 0.040857 | 0.03689 | 0.036383 | 0.022529 | 0.062344 |
| k__Bacteria;p__Bacteroidetes;c__Bacteroidia;o__Bacteroidales;f__NA;g__NA;s__sp12494 | 0.003683 | 0.007087 | 0.002804 | 0 | 0.004383 | 0.002114 | 0 | 0.00588 | 0.003467 | 0.003007 | 0 | 0 | 0.012957 | 0.007674 | 0.006795 | 0.011529 | 0.004377 | 0.007352 |
| k__Bacteria;p__Bacteroidetes;c__Bacteroidia;o__Bacteroidales;f__NA;g__NA;s__sp12497 | 0 | 0.002461 | 0.014346 | 0 | 0.01111 | 0 | 0 | 0.024853 | 0.013489 | 0.007534 | 0.00177 | 0 | 0.040588 | 0.033299 | 0.044753 | 0.055689 | 0.020986 | 0.054661 |
| k__Bacteria;p__Bacteroidetes;c__Bacteroidia;o__Bacteroidales;f__NA;g__NA;s__sp12499 | 0 | 0 | 0.007023 | 0 | 0.008022 | 0.021107 | 0.007154 | 0.011878 | 0 | 0.002489 | 0.009439 | 0.017347 | 0.004496 | 0.002674 | 0 | 0.003001 | 0 | 0.002399 |
| k__Bacteria;p__Bacteroidetes;c__Bacteroidia;o__Bacteroidales;f__NA;g__NA;s__sp12499-sp12616 | 0 | 0 | 0 | 0 | 0.00408 | 0 | 0 | 0 | 0 | 0 | 0 | 0 | 0.002187 | 0 | 0.003325 | 0 | 0.001347 | 0 |
| k__Bacteria;p__Bacteroidetes;c__Bacteroidia;o__Bacteroidales;f__NA;g__NA;s__sp12502 | 0.01609 | 0.024411 | 0 | 0.024043 | 0 | 0.002561 | 0.00582 | 0.011388 | 0 | 0.011393 | 0.005668 | 0.003687 | 0.0084 | 0.003372 | 0 | 0.005192 | 0 | 0.007976 |
| k__Bacteria;p__Bacteroidetes;c__Bacteroidia;o__Bacteroidales;f__NA;g__NA;s__sp12503 | 0 | 0 | 0 | 0 | 0 | 0 | 0 | 0.00539 | 0.006419 | 0.003224 | 0 | 0 | 0.013345 | 0.011162 | 0.014149 | 0.019761 | 0.006313 | 0.01837 |
| k__Bacteria;p__Bacteroidetes;c__Bacteroidia;o__Bacteroidales;f__NA;g__NA;s__sp12505 | 0 | 0 | 0.008167 | 0 | 0.010834 | 0.005414 | 0 | 0.015014 | 0.014166 | 0.010174 | 0.009966 | 0 | 0.006703 | 0.006302 | 0.004927 | 0.00689 | 0.004545 | 0.006494 |
| k__Bacteria;p__Bacteroidetes;c__Bacteroidia;o__Bacteroidales;f__NA;g__NA;s__sp12520 | 0.001612 | 0.008022 | 0.003947 | 0.007222 | 0.00692 | 0.001495 | 0 | 0.002803 | 0.013977 | 0.01445 | 0.008723 | 0.012351 | 0.010832 | 0.008953 | 0.008737 | 0.012792 | 0.007996 | 0.01564 |
| k__Bacteria;p__Bacteroidetes;c__Bacteroidia;o__Bacteroidales;f__NA;g__NA;s__sp12532 | 0.007875 | 0.012353 | 0.02028 | 0 | 0.059161 | 0.019818 | 0 | 0.014602 | 0.046968 | 0.053825 | 0.055559 | 0.000262 | 0.01545 | 0.016417 | 0.017887 | 0.022761 | 0.009371 | 0.021275 |
| k__Bacteria;p__Bacteroidetes;c__Bacteroidia;o__Bacteroidales;f__NA;g__NA;s__sp12535 | 0 | 0 | 0.003866 | 0.002211 | 0.002674 | 0.002974 | 0 | 0.005723 | 0.001517 | 0.00152 | 0.001896 | 0.002975 | 0.00372 | 0.002046 | 0.002451 | 0.005291 | 0.003872 | 0.00546 |
| k__Bacteria;p__Bacteroidetes;c__Bacteroidia;o__Bacteroidales;f__NA;g__NA;s__sp12535-sp12543 | 0 | 0 | 0 | 0 | 0 | 0 | 0 | 0.001509 | 0 | 0 | 0 | 0 | 0 | 0 | 0 | 0 | 0 | 0 |
| k__Bacteria;p__Bacteroidetes;c__Bacteroidia;o__Bacteroidales;f__NA;g__NA;s__sp12536-sp12753 | 0.018177 | 0.006431 | 0.010426 | 0.011151 | 0.016017 | 0.009729 | 0 | 0.018875 | 0.034725 | 0.031272 | 0.036407 | 0.060126 | 0.018475 | 0.009441 | 0.010994 | 0.022288 | 0.007687 | 0.015464 |
| k__Bacteria;p__Bacteroidetes;c__Bacteroidia;o__Bacteroidales;f__NA;g__NA;s__sp12539 | 0.000594 | 0 | 0.010862 | 0 | 0.007636 | 0.005225 | 0.002243 | 0.012642 | 0.00325 | 0.005947 | 0.009776 | 0.012987 | 0.005068 | 0.005279 | 0.00381 | 0.004501 | 0.003058 | 0.004895 |
| k__Bacteria;p__Bacteroidetes;c__Bacteroidia;o__Bacteroidales;f__NA;g__NA;s__sp12549 | 0.004328 | 0.001247 | 0.003484 | 0.00901 | 0.002398 | 0.001031 | 0.002274 | 0.01178 | 0.003467 | 0.004043 | 0.002613 | 0.003387 | 0.005048 | 0.002163 | 0.002209 | 0.000888 | 0.000673 | 0.002165 |
| k__Bacteria;p__Bacteroidetes;c__Bacteroidia;o__Bacteroidales;f__NA;g__NA;s__sp12565 | 0.007926 | 0.008974 | 0.011188 | 0.003364 | 0.015466 | 0.01174 | 0.00961 | 0.017503 | 0.023159 | 0.019863 | 0.03192 | 0.030091 | 0.02197 | 0.014766 | 0.014853 | 0.017905 | 0.008978 | 0.024571 |
| k__Bacteria;p__Bacteroidetes;c__Bacteroidia;o__Bacteroidales;f__NA;g__NA;s__sp12572-sp12578-sp12693 | 0.102834 | 0.154589 | 0.02597 | 0 | 0.008298 | 0.009591 | 0.084215 | 0.102901 | 0.060917 | 0.050233 | 0.023471 | 0.00043 | 0.07157 | 0.078504 | 0.103922 | 0.051563 | 0.040401 | 0.070866 |
| k__Bacteria;p__Bacteroidetes;c__Bacteroidia;o__Bacteroidales;f__NA;g__NA;s__sp12589 | 0.01166 | 0.006234 | 0.011352 | 0 | 0.023598 | 0.018082 | 0.006791 | 0.034496 | 0.033072 | 0.036568 | 0.047953 | 0.045642 | 0.030676 | 0.01958 | 0.034827 | 0.021834 | 0.017816 | 0.038007 |
| k__Bacteria;p__Bacteroidetes;c__Bacteroidia;o__Bacteroidales;f__NA;g__NA;s__sp12590 | 0.000407 | 0.000295 | 0.000599 | 0.001435 | 0 | 0.00055 | 0 | 0.000549 | 0 | 0 | 0.000464 | 0.000655 | 0.001267 | 0.00086 | 0.001529 | 0.001935 | 0.000645 | 0.001814 |
| k__Bacteria;p__Bacteroidetes;c__Bacteroidia;o__Bacteroidales;f__NA;g__NA;s__sp12595 | 0.001239 | 0.00023 | 0.003185 | 0.013762 | 0.005762 | 0.002338 | 0 | 0 | 0.005526 | 0.006265 | 0.015949 | 0.004285 | 0.005988 | 0 | 0.007475 | 0.010463 | 0.004966 | 0.010608 |
| k__Bacteria;p__Bacteroidetes;c__Bacteroidia;o__Bacteroidales;f__NA;g__NA;s__sp12595-sp12654 | 0 | 0 | 0.000463 | 0 | 0.001902 | 0 | 0 | 0 | 0 | 0.001821 | 0 | 0 | 0.012364 | 0.013836 | 0.012766 | 0.00531 | 0.004685 | 0.01484 |
| k__Bacteria;p__Bacteroidetes;c__Bacteroidia;o__Bacteroidales;f__NA;g__NA;s__sp12597 | 0.005838 | 0.00146 | 0.012495 | 0 | 0.021503 | 0.006532 | 0 | 0.012505 | 0.015412 | 0.0433 | 0.010872 | 0 | 0.019844 | 0.020975 | 0.017256 | 0.020077 | 0.009146 | 0.019735 |
| k__Bacteria;p__Bacteroidetes;c__Bacteroidia;o__Bacteroidales;f__NA;g__NA;s__sp12616-sp12777 | 0 | 0 | 0 | 0 | 0 | 0 | 0 | 0 | 0.000921 | 0 | 0 | 0 | 0 | 0 | 0 | 0 | 0 | 0 |
| k__Bacteria;p__Bacteroidetes;c__Bacteroidia;o__Bacteroidales;f__NA;g__NA;s__sp12629 | 0 | 0 | 0.004219 | 0 | 0.002647 | 0.008749 | 0 | 0.001666 | 0.006772 | 0.007317 | 0.004867 | 0 | 0.000327 | 0 | 0 | 0 | 0 | 0 |
| k__Bacteria;p__Bacteroidetes;c__Bacteroidia;o__Bacteroidales;f__NA;g__NA;s__sp12637 | 0 | 0 | 0.003838 | 0 | 0.011275 | 0 | 0 | 0.001921 | 0.006744 | 0.007467 | 0 | 0 | 0.006949 | 0.008976 | 0.007062 | 0.012891 | 0.005471 | 0.010101 |
| k__Bacteria;p__Bacteroidetes;c__Bacteroidia;o__Bacteroidales;f__NA;g__NA;s__sp12641 | 0.020723 | 0.012091 | 0.009745 | 0.010233 | 0.018222 | 0.009161 | 0.018037 | 0.041141 | 0.036458 | 0.037838 | 0.034279 | 0.059826 | 0.018761 | 0.011325 | 0.013809 | 0.025091 | 0.007996 | 0.016069 |
| k__Bacteria;p__Bacteroidetes;c__Bacteroidia;o__Bacteroidales;f__NA;g__NA;s__sp12654 | 0.000339 | 0.005151 | 0.003866 | 0.030394 | 0.002812 | 0.001255 | 0 | 0.002117 | 0.003711 | 0.006097 | 0.004403 | 0.005333 | 0.004782 | 0.003628 | 0.0058 | 0.01068 | 0.006649 | 0.010608 |
| k__Bacteria;p__Bacteroidetes;c__Bacteroidia;o__Bacteroidales;f__NA;g__NA;s__sp12656 | 0.022811 | 0.057912 | 0.052892 | 0.082479 | 0.074737 | 0.041613 | 0.00673 | 0.007899 | 0.031745 | 0.043367 | 0.035965 | 0.057618 | 0.023012 | 0.016557 | 0.018323 | 0.012042 | 0.010521 | 0.019247 |
| k__Bacteria;p__Bacteroidetes;c__Bacteroidia;o__Bacteroidales;f__NA;g__NA;s__sp12666 | 0.018805 | 0.041654 | 0.030108 | 0.024654 | 0.011275 | 0.003472 | 0 | 0.030909 | 0.016658 | 0.019813 | 0.015865 | 0.007055 | 0.014878 | 0.006744 | 0.015047 | 0.013523 | 0.010465 | 0.019852 |
| k__Bacteria;p__Bacteroidetes;c__Bacteroidia;o__Bacteroidales;f__NA;g__NA;s__sp12757 | 0 | 0 | 0.002069 | 0 | 0.002261 | 0 | 0 | 0.001313 | 0.002221 | 0.002188 | 0.002423 | 0 | 0.01308 | 0.006627 | 0.006577 | 0.010601 | 0.004994 | 0.004115 |
| k__Bacteria;p__Bacteroidetes;c__Bacteroidia;o__Bacteroidales;f__NA;g__NA;s__sp12777 | 0.000289 | 0 | 0.02077 | 0 | 0.011689 | 0.042696 | 0.019189 | 0.023344 | 0.002817 | 0.007551 | 0.016961 | 0.033366 | 0.005845 | 0.006139 | 0.003786 | 0.006238 | 0.003423 | 0.006279 |
| k__Bacteria;p__Bacteroidetes;c__Bacteroidia;o__Bacteroidales;f__NA;g__NA;s__sp12778 | 0 | 0 | 0.001878 | 0 | 0.003391 | 0.000894 | 0 | 0 | 0.038354 | 0.032843 | 0.001243 | 0 | 0.106783 | 0.102874 | 0.07242 | 0.076062 | 0.052633 | 0.134926 |
| k__Bacteria;p__Bacteroidetes;c__Bacteroidia;o__Bacteroidales;f__NA;g__NA;s__sp12785 | 0 | 0.000886 | 0.004301 | 0 | 0.006947 | 0.004108 | 0.002122 | 0.003724 | 0.003332 | 0.003642 | 0.004656 | 0.002863 | 0.003924 | 0.001744 | 0 | 0.001737 | 0.001403 | 0.002379 |
| k__Bacteria;p__Bacteroidetes;c__Bacteroidia;o__Bacteroidales;f__NA;g__NA;s__sp12790 | 0 | 0 | 0.006098 | 0 | 0.009704 | 0.005071 | 0 | 0.015602 | 0.006338 | 0.008937 | 0.011672 | 0.056271 | 0.006724 | 0.008185 | 0.005218 | 0.006593 | 0.005639 | 0.009068 |
| k__Bacteria;p__Bacteroidetes;c__Bacteroidia;o__Bacteroidales;f__NA;g__NA;s__sp12802 | 0 | 0 | 0.004682 | 0.001364 | 0.021806 | 0.010502 | 0 | 0.004861 | 0.011241 | 0.016221 | 0.040094 | 0.04261 | 0.010464 | 0.006372 | 0.005242 | 0.007896 | 0.004545 | 0.007878 |
| k__Bacteria;p__Bacteroidetes;c__Bacteroidia;o__Bacteroidales;f__Porphyromonadaceae;g__Odoribacter;s__sp13184 | 0 | 0 | 0.011678 | 0.000447 | 0.012929 | 0.010107 | 0.003941 | 0.013995 | 0.014356 | 0.013414 | 0.012304 | 0.024496 | 0.015369 | 0.011929 | 0.013567 | 0.010482 | 0.008192 | 0.01209 |
| k__Bacteria;p__Bacteroidetes;c__Bacteroidia;o__Bacteroidales;f__Porphyromonadaceae;g__Parabacteroides;s__distasonis | 0 | 0 | 0 | 0 | 0 | 0 | 0 | 0 | 0 | 0 | 0 | 0 | 0 | 0 | 0.000291 | 0 | 0 | 0 |
| k__Bacteria;p__Bacteroidetes;c__Bacteroidia;o__Bacteroidales;f__Porphyromonadaceae;g__Parabacteroides;s__goldsteinii | 0 | 0 | 0 | 0 | 0 | 0 | 0 | 0 | 0 | 0.00147 | 0 | 0.000692 | 0.001615 | 0.000535 | 0.000291 | 0.000691 | 0.000533 | 0.000722 |
| k__Bacteria;p__Bacteroidetes;c__Bacteroidia;o__Bacteroidales;f__Prevotellaceae;g__NA;s__sp13885 | 0 | 0 | 0 | 0 | 0 | 0 | 0 | 0 | 0 | 0 | 0 | 0 | 0 | 0 | 0.00165 | 0 | 0 | 0 |
| k__Bacteria;p__Bacteroidetes;c__Bacteroidia;o__Bacteroidales;f__Prevotellaceae;g__NA;s__sp14210 | 0 | 0 | 0.00754 | 0 | 0.015355 | 0.006136 | 0 | 0.003312 | 0.042742 | 0.044486 | 0.002402 | 0 | 0.032576 | 0.034997 | 0.0324 | 0.089446 | 0.026317 | 0.03828 |
| k__Bacteria;p__Bacteroidetes;c__Bacteroidia;o__Bacteroidales;f__Prevotellaceae;g__Prevotella;s__copri | 0 | 0 | 0 | 0 | 0 | 0 | 0 | 0 | 0 | 0 | 0 | 0 | 0 | 0 | 0.016091 | 0 | 0 | 0 |
| k__Bacteria;p__Bacteroidetes;c__Bacteroidia;o__Bacteroidales;f__Prevotellaceae;g__Prevotella;s__copri-sp13942 | 0 | 0 | 0 | 0 | 0 | 0 | 0 | 0 | 0 | 0 | 0 | 0 | 0 | 0 | 0.015751 | 0 | 0 | 0 |
| k__Bacteria;p__Bacteroidetes;c__Bacteroidia;o__Bacteroidales;f__Rikenellaceae;g__Alistipes;s__putredinis | 0 | 0 | 0 | 0 | 0 | 0.004727 | 0 | 0 | 0.00149 | 0.000869 | 0 | 0 | 0.003638 | 0.003535 | 0.003204 | 0.004955 | 0.001768 | 0.006396 |
| k__Bacteria;p__Bacteroidetes;c__Bacteroidia;o__Bacteroidales;f__Rikenellaceae;g__Alistipes;s__sp14330 | 0.001222 | 0 | 0 | 0 | 0.000744 | 0.001409 | 0.003031 | 0 | 0.000948 | 0.001136 | 0.002613 | 0 | 0.005538 | 0.004627 | 0.004296 | 0.005705 | 0.003619 | 0.002964 |
| k__Bacteria;p__Bacteroidetes;c__Bacteroidia;o__Bacteroidales;f__Rikenellaceae;g__Alistipes;s__sp14336 | 0 | 0 | 0.014509 | 0 | 0.007471 | 0.013304 | 0.003486 | 0.020933 | 0.022265 | 0.021633 | 0.045952 | 0.018152 | 0.022562 | 0.018347 | 0.015387 | 0.011371 | 0.012036 | 0.014138 |
| k__Bacteria;p__Bacteroidetes;c__Bacteroidia;o__Bacteroidales;f__Rikenellaceae;g__Alistipes;s__sp14338 | 0 | 0 | 0.01195 | 0 | 0.026548 | 0.018151 | 0 | 0 | 0.006094 | 0.00862 | 0.019089 | 0.012519 | 0.014122 | 0.010836 | 0.008155 | 0.006534 | 0.00404 | 0.00743 |
| k__Bacteria;p__Bacteroidetes;c__Bacteroidia;o__Bacteroidales;f__Rikenellaceae;g__Alistipes;s__sp14343 | 0 | 0 | 0 | 0 | 0 | 0 | 0 | 0 | 0 | 0.000668 | 0 | 0 | 0.001206 | 0 | 0.000752 | 0.000612 | 0 | 0.001053 |
| k__Bacteria;p__Bacteroidetes;c__Bacteroidia;o__Bacteroidales;f__Rikenellaceae;g__Alistipes;s__sp14354 | 0 | 0 | 0.00343 | 0 | 0.001489 | 0.003128 | 0.056356 | 0.065347 | 0.031989 | 0.055161 | 0.013273 | 0.000524 | 0.054342 | 0.047089 | 0.041598 | 0.03887 | 0.035884 | 0.044949 |
| k__Bacteria;p__Bacteroidetes;c__Bacteroidia;o__Bacteroidales;f__Rikenellaceae;g__Alistipes;s__timonensis | 0.018568 | 0 | 0.002749 | 0.002799 | 0.006809 | 0.00385 | 0.002334 | 0.005527 | 0.002546 | 0.002857 | 0.012578 | 0.009619 | 0.004598 | 0.007929 | 0.003907 | 0.01297 | 0.005303 | 0.004349 |
| k__Bacteria;p__Bacteroidetes;c__Bacteroidia;o__Bacteroidales;f__Rikenellaceae;g__NA;s__sp14636 | 0 | 0 | 0.000544 | 0.102875 | 0.006892 | 0.014748 | 0 | 0.003802 | 0.008126 | 0.008754 | 0.012094 | 0.014297 | 0.011956 | 0.00572 | 0.004102 | 0.018991 | 0.005892 | 0.007683 |
| k__Bacteria;p__Cyanobacteria;c__Melainabacteria;o__Gastranaerophilales;f__NA;g__NA;s__sp23727 | 0 | 0 | 0 | 0 | 0.005238 | 0.002131 | 0 | 0 | 0.002979 | 0.000919 | 0.009081 | 0 | 0.000654 | 0 | 0.001966 | 0.000829 | 0.000701 | 0.00197 |
| k__Bacteria;p__Cyanobacteria;c__Melainabacteria;o__Gastranaerophilales;f__NA;g__NA;s__sp23729 | 0 | 0 | 0 | 0 | 0 | 0 | 0 | 0 | 0 | 0 | 0 | 0 | 0.001533 | 0.001535 | 0.001893 | 0.002724 | 0.001206 | 0.003296 |
| k__Bacteria;p__Deferribacteres;c__Deferribacteres;o__Deferribacterales;f__Deferribacteraceae;g__Mucispirillum;s__sp23890 | 0.001137 | 0 | 0.002913 | 0 | 0 | 0.006308 | 0.018886 | 0.001627 | 0 | 0.000384 | 0.000337 | 0.000655 | 0.000797 | 0.000442 | 0.000607 | 0.000257 | 0.00216 | 0.000468 |
| k__Bacteria;p__Firmicutes;c__Bacilli;o__Bacillales;f__Staphylococcaceae;g__Jeotgalicoccus;s__halophilus-halotolerans-nanhaiensis | 0 | 0 | 0 | 0 | 0 | 0 | 0 | 0 | 0 | 0 | 0 | 0 | 0.000511 | 0 | 0 | 0 | 0 | 0 |
| k__Bacteria;p__Firmicutes;c__Bacilli;o__Bacillales;f__Staphylococcaceae;g__Staphylococcus;s__saprophyticus-xylosus | 0 | 0 | 0 | 0 | 0 | 0 | 0 | 0 | 0 | 0 | 0 | 0 | 0.000245 | 0 | 0 | 0 | 0 | 0 |
| k__Bacteria;p__Firmicutes;c__Bacilli;o__Lactobacillales;f__Aerococcaceae;g__Aerococcus;s__urinaeequi-viridans | 0 | 0 | 0 | 0 | 0 | 0 | 0 | 0 | 0 | 0 | 0 | 0 | 0.000715 | 0 | 0 | 0 | 0 | 0 |
| k__Bacteria;p__Firmicutes;c__Bacilli;o__Lactobacillales;f__Enterococcaceae;g__Enterococcus;s__faecalis | 0 | 0 | 0 | 0.001364 | 0.001158 | 0.000241 | 0 | 0 | 0 | 0 | 0 | 0.000636 | 0 | 0 | 0 | 0 | 0 | 0 |
| k__Bacteria;p__Firmicutes;c__Bacilli;o__Lactobacillales;f__Lactobacillaceae;g__Lactobacillus;s__NA | 0.03498 | 0.004807 | 0.010263 | 0.003035 | 0.003253 | 0.12587 | 0.004669 | 0.095061 | 0.007828 | 0.013331 | 0.006679 | 0.018732 | 0.002759 | 0.000814 | 0.007718 | 0.005251 | 0.002974 | 0.028705 |
| k__Bacteria;p__Firmicutes;c__Bacilli;o__Lactobacillales;f__Lactobacillaceae;g__Lactobacillus;s__johnsonii | 0 | 0.003478 | 0.04508 | 0.086572 | 0.145559 | 0.046735 | 0 | 0 | 0.014193 | 0.012746 | 0.00335 | 0.000786 | 0 | 0 | 0 | 0 | 0 | 0 |
| k__Bacteria;p__Firmicutes;c__Bacilli;o__Lactobacillales;f__Lactobacillaceae;g__Lactobacillus;s__reuteri-vaginalis | 0.009997 | 0.00379 | 0.015979 | 0.021361 | 0.026603 | 0.028997 | 0.006821 | 0.011486 | 0.007801 | 0.003425 | 0.003961 | 0 | 0.004149 | 0.00107 | 0 | 0.012358 | 0.007154 | 0.002028 |
| k__Bacteria;p__Firmicutes;c__Bacilli;o__Lactobacillales;f__Lactobacillaceae;g__Lactobacillus;s__taiwanensis | 0.001782 | 0.001362 | 0.012495 | 0 | 0 | 0 | 0.020675 | 0.026637 | 0.009778 | 0.002539 | 0.00413 | 0 | 0.021357 | 0.006139 | 0.016455 | 0.02057 | 0.017759 | 0.005519 |
| k__Bacteria;p__Firmicutes;c__Clostridia;o__Clostridiales;f__Christensenellaceae;g__NA;s__sp30268 | 0 | 0 | 0 | 0 | 0 | 0 | 0 | 0 | 0 | 0 | 0 | 0 | 0.000552 | 0.000744 | 0.000607 | 0 | 0 | 0 |
| k__Bacteria;p__Firmicutes;c__Clostridia;o__Clostridiales;f__Family XIII;g__Anaerovorax;s__sp31505 | 0 | 0 | 0 | 0 | 0 | 0 | 0 | 0 | 0 | 0 | 0 | 0 | 0 | 0 | 0.00034 | 0 | 0 | 0 |
| k__Bacteria;p__Firmicutes;c__Clostridia;o__Clostridiales;f__Family XIII;g__NA;s__sp31521 | 0 | 0 | 0 | 0 | 0 | 0 | 0 | 0 | 0 | 0 | 0 | 0.000337 | 0 | 0 | 0 | 0 | 0 | 0 |
| k__Bacteria;p__Firmicutes;c__Clostridia;o__Clostridiales;f__Family XIII;g__NA;s__sp31651 | 0 | 0 | 0 | 0 | 0 | 0 | 0 | 0 | 0 | 0 | 0 | 0 | 0 | 0 | 0 | 0 | 0 | 0.000273 |
| k__Bacteria;p__Firmicutes;c__Clostridia;o__Clostridiales;f__Lachnospiraceae;g__Acetatifactor;s__sp31893 | 0 | 0 | 0 | 0 | 0 | 0.000773 | 0.001758 | 0.000529 | 0 | 0 | 0 | 0 | 0 | 0 | 0 | 0 | 0 | 0 |
| k__Bacteria;p__Firmicutes;c__Clostridia;o__Clostridiales;f__Lachnospiraceae;g__Acetitomaculum;s__sp31923 | 0 | 0 | 0 | 0 | 0 | 0 | 0 | 0 | 0 | 0 | 0 | 0 | 0 | 0 | 0 | 0.00075 | 0.002104 | 0 |
| k__Bacteria;p__Firmicutes;c__Clostridia;o__Clostridiales;f__Lachnospiraceae;g__Anaerostipes;s__hadrus | 0 | 0 | 0 | 0 | 0 | 0 | 0 | 0 | 0 | 0 | 0 | 0 | 0 | 0 | 0.002354 | 0 | 0 | 0 |
| k__Bacteria;p__Firmicutes;c__Clostridia;o__Clostridiales;f__Lachnospiraceae;g__Blautia;s__coccoides | 0 | 0.010434 | 0 | 0 | 0 | 0 | 0 | 0.000921 | 0 | 0 | 0 | 0 | 0 | 0 | 0 | 0.000948 | 0.01992 | 0.003452 |
| k__Bacteria;p__Firmicutes;c__Clostridia;o__Clostridiales;f__Lachnospiraceae;g__Blautia;s__hansenii-producta | 0 | 0 | 0 | 0 | 0 | 0 | 0 | 0 | 0 | 0 | 0 | 0 | 0 | 0 | 0.001553 | 0 | 0 | 0 |
| k__Bacteria;p__Firmicutes;c__Clostridia;o__Clostridiales;f__Lachnospiraceae;g__Blautia;s__sp32009 | 0 | 0 | 0 | 0 | 0 | 0 | 0 | 0 | 0 | 0 | 0 | 0 | 0 | 0 | 0.001189 | 0 | 0 | 0 |
| k__Bacteria;p__Firmicutes;c__Clostridia;o__Clostridiales;f__Lachnospiraceae;g__Blautia;s__sp32015 | 0.00387 | 0.009368 | 0.001089 | 0.000729 | 0.000414 | 0.00153 | 0.006154 | 0.003685 | 0.002465 | 0.000718 | 0.001096 | 0.000487 | 0.000817 | 0.001744 | 0.003301 | 0.001362 | 0.001291 | 0.000449 |

| | | | | | | | | | | | | | | | | | | |
|---|---|---|---|---|---|---|---|---|---|---|---|---|---|---|---|---|---|---|
| k__Bacteria;p__Firmicutes;c__Clostridia;o__Clostridiales;f__Lachnospiraceae;g__Blautia;s__sp32048 | 0 | 0 | 0 | 0 | 0 | 0 | 0 | 0 | 0 | 0 | 0 | 0 | 0 | 0 | 0.001165 | 0 | 0 | 0 |
| k__Bacteria;p__Firmicutes;c__Clostridia;o__Clostridiales;f__Lachnospiraceae;g__Blautia;s__sp32056 | 0 | 0 | 0 | 0 | 0 | 0 | 0 | 0 | 0 | 0 | 0 | 0 | 0 | 0 | 0.001869 | 0 | 0 | 0 |
| k__Bacteria;p__Firmicutes;c__Clostridia;o__Clostridiales;f__Lachnospiraceae;g__Blautia;s__wexlerae | 0 | 0 | 0 | 0 | 0 | 0 | 0 | 0 | 0 | 0 | 0 | 0 | 0 | 0 | 0.004004 | 0 | 0 | 0 |
| k__Bacteria;p__Firmicutes;c__Clostridia;o__Clostridiales;f__Lachnospiraceae;g__Coprococcus;s__comes-sp32193 | 0 | 0 | 0 | 0 | 0 | 0 | 0 | 0 | 0 | 0 | 0 | 0 | 0 | 0 | 0.000922 | 0 | 0 | 0 |
| k__Bacteria;p__Firmicutes;c__Clostridia;o__Clostridiales;f__Lachnospiraceae;g__Dorea;s__longicatena | 0 | 0 | 0 | 0 | 0 | 0 | 0 | 0 | 0 | 0 | 0 | 0 | 0 | 0 | 0.001529 | 0 | 0 | 0 |
| k__Bacteria;p__Firmicutes;c__Clostridia;o__Clostridiales;f__Lachnospiraceae;g__Eubacterium;s__hallii | 0 | 0 | 0 | 0 | 0 | 0 | 0 | 0 | 0 | 0 | 0 | 0 | 0 | 0 | 0.002014 | 0 | 0 | 0 |
| k__Bacteria;p__Firmicutes;c__Clostridia;o__Clostridiales;f__Lachnospiraceae;g__Eubacterium;s__rectale | 0 | 0 | 0 | 0 | 0 | 0 | 0 | 0 | 0 | 0 | 0 | 0 | 0 | 0 | 0.00381 | 0 | 0 | 0 |
| k__Bacteria;p__Firmicutes;c__Clostridia;o__Clostridiales;f__Lachnospiraceae;g__Fusicatenibacter;s__saccharivorans | 0 | 0 | 0 | 0 | 0 | 0 | 0 | 0 | 0 | 0 | 0 | 0 | 0 | 0 | 0.003398 | 0 | 0 | 0 |
| k__Bacteria;p__Firmicutes;c__Clostridia;o__Clostridiales;f__Lachnospiraceae;g__Lachnoclostridium;s__sp32351 | 0.001595 | 0 | 0.004846 | 0 | 0.00929 | 0.005449 | 0.003456 | 0 | 0.002086 | 0.000334 | 0 | 0 | 0 | 0 | 0.000752 | 0 | 0 | 0 |
| k__Bacteria;p__Firmicutes;c__Clostridia;o__Clostridiales;f__Lachnospiraceae;g__Lachnoclostridium;s__sp32362-sp33711 | 0 | 0 | 0 | 0 | 0 | 0 | 0.002334 | 0.005449 | 0 | 0 | 0 | 0 | 0 | 0 | 0 | 0.000612 | 0 | 0 |
| k__Bacteria;p__Firmicutes;c__Clostridia;o__Clostridiales;f__Lachnospiraceae;g__Lachnoclostridium;s__sp32364 | 0.000272 | 0.000262 | 0 | 0 | 0 | 0 | 0 | 0.000255 | 0.001002 | 0 | 0 | 0 | 0 | 0 | 0 | 0 | 0 | 0 |
| k__Bacteria;p__Firmicutes;c__Clostridia;o__Clostridiales;f__Lachnospiraceae;g__Lachnoclostridium;s__sp32366 | 0.000407 | 0.0021 | 0 | 0 | 0 | 0.000309 | 0.004093 | 0.00147 | 0.000704 | 0.000401 | 0.000464 | 0 | 0 | 0.000977 | 0.001578 | 0.001224 | 0.001319 | 0 |
| k__Bacteria;p__Firmicutes;c__Clostridia;o__Clostridiales;f__Lachnospiraceae;g__Lachnoclostridium;s__sp32371-sp33589 | 0 | 0 | 0 | 0 | 0 | 0 | 0 | 0 | 0 | 0 | 0 | 0 | 0 | 0.000419 | 0 | 0 | 0 | 0 |
| k__Bacteria;p__Firmicutes;c__Clostridia;o__Clostridiales;f__Lachnospiraceae;g__Lachnoclostridium;s__sp32380-sp32437 | 0 | 0 | 0 | 0 | 0 | 0 | 0 | 0 | 0 | 0 | 0 | 0 | 0 | 0 | 0.002378 | 0 | 0 | 0 |
| k__Bacteria;p__Firmicutes;c__Clostridia;o__Clostridiales;f__Lachnospiraceae;g__Lachnoclostridium;s__sp32387 | 0 | 0 | 0 | 0 | 0 | 0.000636 | 0.003456 | 0 | 0 | 0 | 0 | 0 | 0 | 0 | 0 | 0 | 0 | 0 |
| k__Bacteria;p__Firmicutes;c__Clostridia;o__Clostridiales;f__Lachnospiraceae;g__Lachnoclostridium;s__sp32400 | 0 | 0 | 0 | 0 | 0 | 0 | 0 | 0 | 0 | 0 | 0 | 0 | 0 | 0 | 0.000413 | 0 | 0 | 0 |
| k__Bacteria;p__Firmicutes;c__Clostridia;o__Clostridiales;f__Lachnospiraceae;g__Lachnoclostridium;s__sp32414 | 0 | 0 | 0 | 0 | 0 | 0 | 0 | 0 | 0 | 0 | 0 | 0 | 0 | 0.000512 | 0 | 0.000592 | 0.001178 | 0.000371 |
| k__Bacteria;p__Firmicutes;c__Clostridia;o__Clostridiales;f__Lachnospiraceae;g__Lachnoclostridium;s__sp32428 | 0 | 0 | 0 | 0 | 0 | 0 | 0 | 0 | 0 | 0 | 0 | 0 | 0 | 0 | 0.000582 | 0 | 0 | 0 |
| k__Bacteria;p__Firmicutes;c__Clostridia;o__Clostridiales;f__Lachnospiraceae;g__Lachnoclostridium;s__sp32442 | 0 | 0.001296 | 0 | 0 | 0 | 0 | 0 | 0 | 0 | 0 | 0 | 0 | 0 | 0.001488 | 0 | 0 | 0 | 0 |
| k__Bacteria;p__Firmicutes;c__Clostridia;o__Clostridiales;f__Lachnospiraceae;g__Lachnoclostridium-Roseburia;s__sp32368-sp33144 | 0 | 0 | 0 | 0 | 0 | 0 | 0 | 0 | 0 | 0 | 0 | 0 | 0 | 0 | 0.000558 | 0 | 0 | 0 |
| k__Bacteria;p__Firmicutes;c__Clostridia;o__Clostridiales;f__Lachnospiraceae;g__Marvinbryantia;s__sp32979 | 0.004413 | 0.010221 | 0.005172 | 0.001059 | 0 | 0 | 0.004032 | 0.007115 | 0.005878 | 0.003157 | 0.001369 | 0.002133 | 0.004047 | 0.002697 | 0 | 0.005251 | 0.000393 | 0.003822 |
| k__Bacteria;p__Firmicutes;c__Clostridia;o__Clostridiales;f__Lachnospiraceae;g__Marvinbryantia;s__sp32983 | 0.001273 | 0 | 0 | 0 | 0 | 0.000567 | 0 | 0 | 0 | 0 | 0 | 0.00146 | 0 | 0 | 0 | 0.001599 | 0 | 0 |
| k__Bacteria;p__Firmicutes;c__Clostridia;o__Clostridiales;f__Lachnospiraceae;g__NA;s__NA | 0 | 0 | 0 | 0 | 0 | 0 | 0 | 0 | 0 | 0.001019 | 0 | 0 | 0 | 0 | 0 | 0.001678 | 0 | 0 |
| k__Bacteria;p__Firmicutes;c__Clostridia;o__Clostridiales;f__Lachnospiraceae;g__NA;s__sp32146 | 0.002461 | 0.001641 | 0.001279 | 0 | 0 | 0.001272 | 0.002001 | 0.000647 | 0.001463 | 0.000484 | 0.000906 | 0.000674 | 0.001349 | 0.002488 | 0.001917 | 0 | 0 | 0 |
| k__Bacteria;p__Firmicutes;c__Clostridia;o__Clostridiales;f__Lachnospiraceae;g__NA;s__sp32147 | 0 | 0 | 0 | 0 | 0 | 0 | 0 | 0 | 0 | 0 | 0 | 0 | 0 | 0 | 0 | 0.000612 | 0.001487 | 0 |
| k__Bacteria;p__Firmicutes;c__Clostridia;o__Clostridiales;f__Lachnospiraceae;g__NA;s__sp32165 | 0 | 0 | 0 | 0 | 0 | 0.000619 | 0 | 0 | 0.000813 | 0 | 0.0004 | 0 | 0 | 0.000977 | 0.000437 | 0.000711 | 0 | 0 |
| k__Bacteria;p__Firmicutes;c__Clostridia;o__Clostridiales;f__Lachnospiraceae;g__NA;s__sp32165-sp32166 | 0 | 0 | 0 | 0 | 0 | 0 | 0 | 0 | 0 | 0 | 0 | 0 | 0 | 0 | 0 | 0 | 0.00101 | 0 |
| k__Bacteria;p__Firmicutes;c__Clostridia;o__Clostridiales;f__Lachnospiraceae;g__NA;s__sp32166 | 0.000475 | 0.000295 | 0 | 0 | 0 | 0 | 0 | 0 | 0 | 0 | 0 | 0 | 0 | 0.000581 | 0.000534 | 0 | 0 | 0 |
| k__Bacteria;p__Firmicutes;c__Clostridia;o__Clostridiales;f__Lachnospiraceae;g__NA;s__sp32254 | 0 | 0 | 0 | 0 | 0 | 0 | 0 | 0 | 0 | 0 | 0 | 0 | 0.000674 | 0.002139 | 0.00199 | 0.001875 | 0.003619 | 0.001131 |
| k__Bacteria;p__Firmicutes;c__Clostridia;o__Clostridiales;f__Lachnospiraceae;g__NA;s__sp32255 | 0 | 0 | 0 | 0 | 0 | 0.006067 | 0 | 0 | 0 | 0 | 0 | 0 | 0.002963 | 0.001791 | 0.000631 | 0.004205 | 0.002413 | 0.000488 |
| k__Bacteria;p__Firmicutes;c__Clostridia;o__Clostridiales;f__Lachnospiraceae;g__NA;s__sp32261 | 0.002223 | 0.004594 | 0.005444 | 0 | 0.004714 | 0.00373 | 0.009489 | 0.003312 | 0.006582 | 0.002489 | 0.002486 | 0.002882 | 0.000736 | 0.004348 | 0.001602 | 0.000829 | 0.001936 | 0 |
| k__Bacteria;p__Firmicutes;c__Clostridia;o__Clostridiales;f__Lachnospiraceae;g__NA;s__sp32263 | 0 | 0 | 0 | 0 | 0 | 0.000619 | 0 | 0 | 0 | 0 | 0 | 0 | 0 | 0 | 0 | 0 | 0.002609 | 0 |
| k__Bacteria;p__Firmicutes;c__Clostridia;o__Clostridiales;f__Lachnospiraceae;g__NA;s__sp32277 | 0 | 0 | 0 | 0 | 0 | 0 | 0 | 0 | 0 | 0 | 0 | 0 | 0 | 0 | 0.001116 | 0 | 0 | 0 |
| k__Bacteria;p__Firmicutes;c__Clostridia;o__Clostridiales;f__Lachnospiraceae;g__NA;s__sp32486 | 0 | 0 | 0 | 0 | 0 | 0 | 0 | 0 | 0 | 0 | 0 | 0 | 0 | 0.000419 | 0 | 0 | 0 | 0 |
| k__Bacteria;p__Firmicutes;c__Clostridia;o__Clostridiales;f__Lachnospiraceae;g__NA;s__sp32498 | 0 | 0 | 0.000299 | 0 | 0 | 0 | 0 | 0 | 0 | 0 | 0 | 0 | 0 | 0 | 0 | 0 | 0 | 0 |
| k__Bacteria;p__Firmicutes;c__Clostridia;o__Clostridiales;f__Lachnospiraceae;g__NA;s__sp32519 | 0 | 0 | 0 | 0 | 0 | 0 | 0 | 0 | 0 | 0 | 0 | 0 | 0 | 0 | 0.001772 | 0 | 0 | 0 |
| k__Bacteria;p__Firmicutes;c__Clostridia;o__Clostridiales;f__Lachnospiraceae;g__NA;s__sp32590 | 0 | 0 | 0 | 0 | 0 | 0 | 0 | 0 | 0 | 0 | 0 | 0 | 0 | 0 | 0 | 0.005192 | 0 | 0 |
| k__Bacteria;p__Firmicutes;c__Clostridia;o__Clostridiales;f__Lachnospiraceae;g__NA;s__sp32593 | 0 | 0 | 0.001007 | 0 | 0 | 0 | 0 | 0 | 0 | 0 | 0 | 0 | 0.003842 | 0.007929 | 0 | 0.000612 | 0 | 0 |
| k__Bacteria;p__Firmicutes;c__Clostridia;o__Clostridiales;f__Lachnospiraceae;g__NA;s__sp32593-sp32722 | 0 | 0 | 0 | 0 | 0 | 0 | 0 | 0 | 0 | 0 | 0 | 0 | 0 | 0 | 0.007548 | 0.006475 | 0.00996 | 0.002204 |
| k__Bacteria;p__Firmicutes;c__Clostridia;o__Clostridiales;f__Lachnospiraceae;g__NA;s__sp32594 | 0.00353 | 0.001624 | 0.001987 | 0.001411 | 0.002784 | 0.01516 | 0.010913 | 0.004802 | 0.001544 | 0.001336 | 0.002908 | 0.000543 | 0 | 0.000512 | 0 | 0.003652 | 0.003254 | 0.00119 |
| k__Bacteria;p__Firmicutes;c__Clostridia;o__Clostridiales;f__Lachnospiraceae;g__NA;s__sp32594-sp32647 | 0 | 0 | 0 | 0 | 0 | 0 | 0 | 0 | 0 | 0 | 0 | 0 | 0.001247 | 0 | 0.002427 | 0 | 0 | 0 |
| k__Bacteria;p__Firmicutes;c__Clostridia;o__Clostridiales;f__Lachnospiraceae;g__NA;s__sp32597 | 0 | 0 | 0 | 0 | 0 | 0 | 0 | 0 | 0 | 0.000936 | 0.000295 | 0 | 0 | 0.000419 | 0 | 0.000395 | 0 | 0 |
| k__Bacteria;p__Firmicutes;c__Clostridia;o__Clostridiales;f__Lachnospiraceae;g__NA;s__sp32604-sp32722 | 0 | 0 | 0 | 0 | 0 | 0 | 0 | 0 | 0 | 0 | 0 | 0 | 0.000388 | 0 | 0 | 0 | 0 | 0 |
| k__Bacteria;p__Firmicutes;c__Clostridia;o__Clostridiales;f__Lachnospiraceae;g__NA;s__sp32617-sp32782 | 0 | 0 | 0 | 0 | 0 | 0 | 0 | 0 | 0 | 0.003191 | 0.004551 | 0 | 0.000981 | 0.000837 | 0.004174 | 0.000948 | 0.001908 | 0.003374 |
| k__Bacteria;p__Firmicutes;c__Clostridia;o__Clostridiales;f__Lachnospiraceae;g__NA;s__sp32622 | 0.000611 | 0.001772 | 0.003321 | 0 | 0 | 0.001856 | 0.006245 | 0.001215 | 0.001219 | 0.000635 | 0.001348 | 0 | 0.001063 | 0.001698 | 0 | 0 | 0 | 0 |
| k__Bacteria;p__Firmicutes;c__Clostridia;o__Clostridiales;f__Lachnospiraceae;g__NA;s__sp32623 | 0 | 0 | 0 | 0 | 0 | 0.001427 | 0 | 0 | 0 | 0 | 0 | 0 | 0.000572 | 0 | 0 | 0 | 0 | 0 |
| k__Bacteria;p__Firmicutes;c__Clostridia;o__Clostridiales;f__Lachnospiraceae;g__NA;s__sp32630 | 0 | 0 | 0 | 0 | 0 | 0 | 0 | 0 | 0 | 0 | 0 | 0 | 0.000368 | 0 | 0.000291 | 0.000513 | 0.001178 | 0.000683 |
| k__Bacteria;p__Firmicutes;c__Clostridia;o__Clostridiales;f__Lachnospiraceae;g__NA;s__sp32648 | 0 | 0 | 0.000354 | 0 | 0.001489 | 0.006755 | 0.007548 | 0.001901 | 0.0013 | 0.000718 | 0 | 0 | 0.000266 | 0.00472 | 0 | 0.003928 | 0.014252 | 0 |
| k__Bacteria;p__Firmicutes;c__Clostridia;o__Clostridiales;f__Lachnospiraceae;g__NA;s__sp32649 | 0 | 0 | 0 | 0 | 0 | 0 | 0 | 0 | 0 | 0 | 0 | 0 | 0.00049 | 0.000535 | 0 | 0 | 0 | 0 |
| k__Bacteria;p__Firmicutes;c__Clostridia;o__Clostridiales;f__Lachnospiraceae;g__NA;s__sp32655 | 0.001765 | 0.001624 | 0.005907 | 0 | 0.004769 | 0.002544 | 0.021099 | 0.00247 | 0.007367 | 0.002322 | 0.000969 | 0.000356 | 0.002044 | 0.004697 | 0.003106 | 0 | 0.006621 | 0 |
| k__Bacteria;p__Firmicutes;c__Clostridia;o__Clostridiales;f__Lachnospiraceae;g__NA;s__sp32658 | 0 | 0 | 0.005009 | 0 | 0.006506 | 0.014026 | 0 | 0 | 0.010103 | 0.001721 | 0 | 0 | 0.006295 | 0.021138 | 0.011625 | 0.003731 | 0.010858 | 0.000858 |
| k__Bacteria;p__Firmicutes;c__Clostridia;o__Clostridiales;f__Lachnospiraceae;g__NA;s__sp32668 | 0.004328 | 0 | 0.03283 | 0.009692 | 0.008877 | 0.030784 | 0 | 0 | 0.011132 | 0.00299 | 0.002065 | 0.001048 | 0.005109 | 0.01644 | 0.002281 | 0.002902 | 0.007098 | 0.001424 |
| k__Bacteria;p__Firmicutes;c__Clostridia;o__Clostridiales;f__Lachnospiraceae;g__NA;s__sp32683-sp32693-sp32746 | 0 | 0 | 0 | 0 | 0 | 0.001083 | 0 | 0 | 0 | 0 | 0 | 0 | 0.001921 | 0.001139 | 0 | 0 | 0 | 0 |
| k__Bacteria;p__Firmicutes;c__Clostridia;o__Clostridiales;f__Lachnospiraceae;g__NA;s__sp32701 | 0 | 0 | 0 | 0 | 0.000717 | 0.003713 | 0 | 0.013485 | 0.005282 | 0.00157 | 0.004804 | 0.005708 | 0.005416 | 0.011743 | 0.004441 | 0.002033 | 0.031086 | 0.001248 |
| k__Bacteria;p__Firmicutes;c__Clostridia;o__Clostridiales;f__Lachnospiraceae;g__NA;s__sp32704 | 0.007332 | 0.000476 | 0 | 0 | 0 | 0 | 0.012581 | 0 | 0 | 0 | 0 | 0 | 0 | 0 | 0 | 0 | 0 | 0 |
| k__Bacteria;p__Firmicutes;c__Clostridia;o__Clostridiales;f__Lachnospiraceae;g__NA;s__sp32706-sp32766 | 0 | 0.002264 | 0.005363 | 0.001764 | 0 | 0.002802 | 0.004487 | 0.00145 | 0.001381 | 0.000735 | 0 | 0 | 0 | 0 | 0 | 0 | 0 | 0 |
| k__Bacteria;p__Firmicutes;c__Clostridia;o__Clostridiales;f__Lachnospiraceae;g__NA;s__sp32721 | 0 | 0 | 0 | 0 | 0 | 0 | 0 | 0 | 0 | 0 | 0 | 0 | 0.001328 | 0.000907 | 0.001286 | 0.0015 | 0.001796 | 0 |
| k__Bacteria;p__Firmicutes;c__Clostridia;o__Clostridiales;f__Lachnospiraceae;g__NA;s__sp32735 | 0 | 0 | 0.017014 | 0 | 0 | 0 | 0 | 0 | 0.013977 | 0.002873 | 0 | 0 | 0.006519 | 0.012301 | 0.00432 | 0.001125 | 0.047751 | 0.004095 |
| k__Bacteria;p__Firmicutes;c__Clostridia;o__Clostridiales;f__Lachnospiraceae;g__NA;s__sp32746-sp32777 | 0.001392 | 0.000459 | 0 | 0.001553 | 0.005844 | 0.017137 | 0.002183 | 0 | 0.011891 | 0.002506 | 0.003266 | 0.003724 | 0 | 0 | 0 | 0 | 0 | 0 |
| k__Bacteria;p__Firmicutes;c__Clostridia;o__Clostridiales;f__Lachnospiraceae;g__NA;s__sp32758 | 0.001731 | 0.004659 | 0 | 0.002847 | 0.001406 | 0.005277 | 0.007457 | 0 | 0.002113 | 0.00127 | 0.001664 | 0.000599 | 0 | 0 | 0 | 0.000494 | 0.001431 | 0 |
| k__Bacteria;p__Firmicutes;c__Clostridia;o__Clostridiales;f__Lachnospiraceae;g__NA;s__sp32758-sp32766 | 0 | 0 | 0.010671 | 0 | 0 | 0 | 0 | 0.00345 | 0 | 0 | 0 | 0 | 0 | 0 | 0 | 0 | 0 | 0 |
| k__Bacteria;p__Firmicutes;c__Clostridia;o__Clostridiales;f__Lachnospiraceae;g__NA;s__sp32761 | 0 | 0 | 0.060378 | 0.000541 | 0.027596 | 0.008869 | 0 | 0 | 0.003711 | 0.003324 | 0.00592 | 0.002208 | 0.000899 | 0.00572 | 0.013251 | 0.000553 | 0.005218 | 0 |
| k__Bacteria;p__Firmicutes;c__Clostridia;o__Clostridiales;f__Lachnospiraceae;g__NA;s__sp32763 | 0.002122 | 0.006021 | 0 | 0 | 0 | 0 | 0.103647 | 0.035025 | 0 | 0.003642 | 0 | 0 | 0.003842 | 0.002488 | 0.002136 | 0.004837 | 0.002609 | 0.004154 |
| k__Bacteria;p__Firmicutes;c__Clostridia;o__Clostridiales;f__Lachnospiraceae;g__NA;s__sp32766 | 0.0074 | 0.002231 | 0.005063 | 0.0016 | 0.000634 | 0.002458 | 0.013369 | 0.001588 | 0.000786 | 0.000535 | 0.001622 | 0.000374 | 0 | 0.000953 | 0.00051 | 0.000316 | 0 | 0 |
| k__Bacteria;p__Firmicutes;c__Clostridia;o__Clostridiales;f__Lachnospiraceae;g__NA;s__sp32778 | 0 | 0 | 0.013665 | 0 | 0 | 0 | 0 | 0 | 0 | 0.001888 | 0 | 0 | 0.00515 | 0.010441 | 0 | 0 | 0.037118 | 0 |
| k__Bacteria;p__Firmicutes;c__Clostridia;o__Clostridiales;f__Lachnospiraceae;g__NA;s__sp32791 | 0.006755 | 0.003626 | 0.00392 | 0.003411 | 0.004438 | 0.029203 | 0.022797 | 0.010368 | 0.00363 | 0.002172 | 0.00571 | 0.001553 | 0.000817 | 0.000907 | 0 | 0.007107 | 0.00721 | 0.002184 |
| k__Bacteria;p__Firmicutes;c__Clostridia;o__Clostridiales;f__Lachnospiraceae;g__NA;s__sp32806 | 0.000322 | 0 | 0 | 0 | 0 | 0 | 0.002243 | 0 | 0 | 0 | 0 | 0 | 0 | 0 | 0 | 0 | 0 | 0 |
| k__Bacteria;p__Firmicutes;c__Clostridia;o__Clostridiales;f__Lachnospiraceae;g__NA;s__sp32816 | 0 | 0 | 0 | 0 | 0 | 0.000602 | 0 | 0 | 0 | 0 | 0.000442 | 0.000281 | 0 | 0 | 0 | 0 | 0 | 0 |
| k__Bacteria;p__Firmicutes;c__Clostridia;o__Clostridiales;f__Lachnospiraceae;g__NA;s__sp32826 | 0 | 0 | 0 | 0 | 0 | 0.001324 | 0 | 0 | 0 | 0 | 0 | 0 | 0.000797 | 0 | 0.000291 | 0.001362 | 0.000449 | 0 |
| k__Bacteria;p__Firmicutes;c__Clostridia;o__Clostridiales;f__Lachnospiraceae;g__NA;s__sp32850 | 0 | 0 | 0 | 0 | 0 | 0 | 0 | 0 | 0.012595 | 0.035716 | 0 | 0 | 0.002289 | 0.002767 | 0.00165 | 0.000849 | 0 | 0.011876 |
| k__Bacteria;p__Firmicutes;c__Clostridia;o__Clostridiales;f__Lachnospiraceae;g__NA;s__sp32856-sp32859 | 0.000305 | 0 | 0 | 0 | 0 | 0 | 0 | 0 | 0 | 0 | 0 | 0 | 0 | 0 | 0 | 0.000454 | 0.001599 | 0 |
| k__Bacteria;p__Firmicutes;c__Clostridia;o__Clostridiales;f__Lachnospiraceae;g__NA;s__sp32859 | 0 | 0 | 0 | 0 | 0 | 0 | 0 | 0 | 0 | 0 | 0 | 0 | 0 | 0 | 0.000655 | 0 | 0 | 0 |
| k__Bacteria;p__Firmicutes;c__Clostridia;o__Clostridiales;f__Lachnospiraceae;g__NA;s__sp32862-sp32872-sp32880 | 0 | 0 | 0 | 0 | 0 | 0 | 0.000576 | 0 | 0 | 0 | 0 | 0 | 0 | 0 | 0 | 0 | 0 | 0 |
| k__Bacteria;p__Firmicutes;c__Clostridia;o__Clostridiales;f__Lachnospiraceae;g__NA;s__sp32872 | 0 | 0 | 0 | 0 | 0 | 0.00067 | 0 | 0 | 0 | 0 | 0 | 0 | 0 | 0.000465 | 0 | 0 | 0.000814 | 0 |
| k__Bacteria;p__Firmicutes;c__Clostridia;o__Clostridiales;f__Lachnospiraceae;g__NA;s__sp32910 | 0 | 0 | 0.006207 | 0.001694 | 0.009346 | 0.005792 | 0 | 0 | 0 | 0.001804 | 0.000885 | 0.000973 | 0 | 0 | 0.001311 | 0 | 0.003563 | 0.001755 |
| k__Bacteria;p__Firmicutes;c__Clostridia;o__Clostridiales;f__Lachnospiraceae;g__NA;s__sp33357 | 0 | 0 | 0 | 0 | 0 | 0 | 0 | 0 | 0 | 0 | 0 | 0 | 0 | 0 | 0.001408 | 0 | 0 | 0 |
| k__Bacteria;p__Firmicutes;c__Clostridia;o__Clostridiales;f__Lachnospiraceae;g__NA;s__sp33395 | 0 | 0 | 0 | 0 | 0 | 0 | 0 | 0 | 0 | 0 | 0 | 0 | 0 | 0 | 0.000752 | 0 | 0 | 0 |
| k__Bacteria;p__Firmicutes;c__Clostridia;o__Clostridiales;f__Lachnospiraceae;g__NA;s__sp33413 | 0 | 0 | 0 | 0 | 0 | 0 | 0 | 0 | 0 | 0 | 0 | 0 | 0 | 0 | 0 | 0 | 0.000701 | 0 |
| k__Bacteria;p__Firmicutes;c__Clostridia;o__Clostridiales;f__Lachnospiraceae;g__NA;s__sp33416-sp33419 | 0 | 0 | 0 | 0.002705 | 0 | 0 | 0 | 0 | 0 | 0 | 0 | 0.000299 | 0 | 0 | 0 | 0 | 0.002188 | 0 |
| k__Bacteria;p__Firmicutes;c__Clostridia;o__Clostridiales;f__Lachnospiraceae;g__NA;s__sp33417 | 0 | 0 | 0 | 0 | 0.001351 | 0 | 0 | 0 | 0 | 0 | 0.000737 | 0 | 0.003985 | 0.002023 | 0 | 0.000217 | 0 | 0 |
| k__Bacteria;p__Firmicutes;c__Clostridia;o__Clostridiales;f__Lachnospiraceae;g__NA;s__sp33419 | 0 | 0 | 0 | 0 | 0 | 0 | 0 | 0 | 0 | 0 | 0.000485 | 0 | 0 | 0 | 0 | 0 | 0 | 0.000429 |
| k__Bacteria;p__Firmicutes;c__Clostridia;o__Clostridiales;f__Lachnospiraceae;g__NA;s__sp33421-sp33679 | 0 | 0 | 0 | 0 | 0 | 0 | 0 | 0 | 0 | 0 | 0 | 0 | 0.001349 | 0 | 0.000728 | 0.001323 | 0.003283 | 0.000449 |
| k__Bacteria;p__Firmicutes;c__Clostridia;o__Clostridiales;f__Lachnospiraceae;g__NA;s__sp33426 | 0 | 0 | 0 | 0 | 0 | 0 | 0 | 0 | 0 | 0 | 0 | 0 | 0 | 0 | 0 | 0 | 0.000786 | 0 |

| | | | | | | | | | | | | | | | | | | |
|---|---|---|---|---|---|---|---|---|---|---|---|---|---|---|---|---|---|---|
| k__Bacteria;p__Firmicutes;c__Clostridia;o__Clostridiales;f__Lachnospiraceae;g__NA;s__sp33428 | 0 | 0 | 0 | 0 | 0 | 0 | 0 | 0 | 0 | 0 | 0 | 0 | 0.000368 | 0 | 0 | 0 | 0 | 0 |
| k__Bacteria;p__Firmicutes;c__Clostridia;o__Clostridiales;f__Lachnospiraceae;g__NA;s__sp33432 | 0 | 0 | 0 | 0 | 0.001048 | 0.004125 | 0 | 0 | 0.001761 | 0 | 0 | 0 | 0.001124 | 0.001767 | 0.001747 | 0.000513 | 0.001571 | 0.0008 |
| k__Bacteria;p__Firmicutes;c__Clostridia;o__Clostridiales;f__Lachnospiraceae;g__NA;s__sp33433 | 0 | 0 | 0.000762 | 0 | 0 | 0.00177 | 0.002456 | 0.000706 | 0.000406 | 0 | 0 | 0 | 0.00049 | 0.000791 | 0 | 0 | 0.00188 | 0.000293 |
| k__Bacteria;p__Firmicutes;c__Clostridia;o__Clostridiales;f__Lachnospiraceae;g__NA;s__sp33436 | 0.000373 | 0 | 0 | 0 | 0 | 0.000241 | 0 | 0 | 0 | 0 | 0 | 0 | 0 | 0 | 0 | 0 | 0 | 0 |
| k__Bacteria;p__Firmicutes;c__Clostridia;o__Clostridiales;f__Lachnospiraceae;g__NA;s__sp33440 | 0.000356 | 0 | 0.000953 | 0 | 0.000634 | 0.000997 | 0.001091 | 0 | 0 | 0.000267 | 0 | 0 | 0 | 0 | 0 | 0 | 0.000393 | 0 |
| k__Bacteria;p__Firmicutes;c__Clostridia;o__Clostridiales;f__Lachnospiraceae;g__NA;s__sp33451-sp33593 | 0.000289 | 0.000459 | 0.002123 | 0 | 0 | 0 | 0.00382 | 0 | 0.000352 | 0 | 0 | 0 | 0.000266 | 0.001023 | 0 | 0.000217 | 0.001403 | 0 |
| k__Bacteria;p__Firmicutes;c__Clostridia;o__Clostridiales;f__Lachnospiraceae;g__NA;s__sp33453 | 0.001544 | 0.002625 | 0.004573 | 0.000494 | 0.001571 | 0.003661 | 0.02116 | 0.001078 | 0.00195 | 0.001086 | 0.001559 | 0 | 0.000695 | 0.001372 | 0.000655 | 0.000829 | 0.004489 | 0 |
| k__Bacteria;p__Firmicutes;c__Clostridia;o__Clostridiales;f__Lachnospiraceae;g__NA;s__sp33456 | 0.000424 | 0 | 0 | 0 | 0 | 0 | 0.005275 | 0.000627 | 0 | 0 | 0 | 0 | 0 | 0.000814 | 0 | 0.000257 | 0.001206 | 0 |
| k__Bacteria;p__Firmicutes;c__Clostridia;o__Clostridiales;f__Lachnospiraceae;g__NA;s__sp33459 | 0.000645 | 0.00041 | 0.003049 | 0 | 0.000717 | 0.001341 | 0.006215 | 0.001156 | 0.001408 | 0.000468 | 0.0004 | 0 | 0.000593 | 0.002093 | 0.000777 | 0.000632 | 0.003591 | 0.000429 |
| k__Bacteria;p__Firmicutes;c__Clostridia;o__Clostridiales;f__Lachnospiraceae;g__NA;s__sp33476 | 0.001952 | 0 | 0 | 0 | 0 | 0 | 0 | 0 | 0 | 0 | 0 | 0 | 0 | 0 | 0 | 0 | 0 | 0 |
| k__Bacteria;p__Firmicutes;c__Clostridia;o__Clostridiales;f__Lachnospiraceae;g__NA;s__sp33478 | 0 | 0 | 0 | 0 | 0 | 0 | 0 | 0 | 0 | 0 | 0 | 0 | 0.00139 | 0.000814 | 0.002354 | 0 | 0.001178 | 0 |
| k__Bacteria;p__Firmicutes;c__Clostridia;o__Clostridiales;f__Lachnospiraceae;g__NA;s__sp33492 | 0 | 0 | 0 | 0 | 0.001103 | 0 | 0 | 0 | 0 | 0 | 0 | 0 | 0 | 0 | 0 | 0 | 0 | 0 |
| k__Bacteria;p__Firmicutes;c__Clostridia;o__Clostridiales;f__Lachnospiraceae;g__NA;s__sp33503 | 0 | 0 | 0.002559 | 0.000659 | 0.001902 | 0.003352 | 0.007215 | 0.001274 | 0.003575 | 0.001804 | 0.000569 | 0 | 0.001901 | 0.003488 | 0.002087 | 0.000553 | 0.003086 | 0 |
| k__Bacteria;p__Firmicutes;c__Clostridia;o__Clostridiales;f__Lachnospiraceae;g__NA;s__sp33503-sp33540 | 0 | 0 | 0 | 0 | 0 | 0 | 0 | 0 | 0 | 0 | 0 | 0 | 0 | 0 | 0 | 0 | 0.000533 | 0 |
| k__Bacteria;p__Firmicutes;c__Clostridia;o__Clostridiales;f__Lachnospiraceae;g__NA;s__sp33513 | 0.002987 | 0.000952 | 0.028365 | 0.010257 | 0.009125 | 0.017395 | 0.041228 | 0.00196 | 0.003196 | 0.003976 | 0.001327 | 0.000374 | 0 | 0.000767 | 0 | 0 | 0 | 0 |
| k__Bacteria;p__Firmicutes;c__Clostridia;o__Clostridiales;f__Lachnospiraceae;g__NA;s__sp33518 | 0.006687 | 0.005283 | 0 | 0 | 0 | 0.000584 | 0 | 0 | 0 | 0 | 0 | 0 | 0 | 0.000977 | 0 | 0.000987 | 0.001711 | 0.000429 |
| k__Bacteria;p__Firmicutes;c__Clostridia;o__Clostridiales;f__Lachnospiraceae;g__NA;s__sp33522 | 0 | 0 | 0 | 0 | 0 | 0.000653 | 0.002819 | 0 | 0.000569 | 0 | 0 | 0 | 0 | 0 | 0 | 0.000513 | 0 | 0 |
| k__Bacteria;p__Firmicutes;c__Clostridia;o__Clostridiales;f__Lachnospiraceae;g__NA;s__sp33525 | 0 | 0 | 0.000653 | 0 | 0 | 0.000275 | 0 | 0 | 0 | 0 | 0 | 0 | 0 | 0 | 0 | 0 | 0 | 0 |
| k__Bacteria;p__Firmicutes;c__Clostridia;o__Clostridiales;f__Lachnospiraceae;g__NA;s__sp33529 | 0 | 0 | 0 | 0 | 0 | 0 | 0 | 0 | 0 | 0 | 0 | 0 | 0 | 0 | 0 | 0 | 0.000786 | 0 |
| k__Bacteria;p__Firmicutes;c__Clostridia;o__Clostridiales;f__Lachnospiraceae;g__NA;s__sp33537 | 0 | 0 | 0 | 0 | 0.000855 | 0.003919 | 0 | 0 | 0 | 0 | 0.000674 | 0.000243 | 0 | 0.001698 | 0.003276 | 0 | 0.001796 | 0.000605 |
| k__Bacteria;p__Firmicutes;c__Clostridia;o__Clostridiales;f__Lachnospiraceae;g__NA;s__sp33539 | 0 | 0 | 0 | 0.003905 | 0.000414 | 0.001616 | 0 | 0 | 0.001408 | 0.001955 | 0.000274 | 0.000225 | 0.004271 | 0.003721 | 0 | 0.000612 | 0.001571 | 0 |
| k__Bacteria;p__Firmicutes;c__Clostridia;o__Clostridiales;f__Lachnospiraceae;g__NA;s__sp33540 | 0 | 0 | 0.004111 | 0 | 0.001792 | 0.004211 | 0 | 0 | 0.002709 | 0 | 0 | 0 | 0 | 0 | 0 | 0 | 0 | 0 |
| k__Bacteria;p__Firmicutes;c__Clostridia;o__Clostridiales;f__Lachnospiraceae;g__NA;s__sp33565 | 0.002138 | 0.000771 | 0.013665 | 0.011927 | 0.00521 | 0.004744 | 0.026647 | 0.002509 | 0.005228 | 0.002456 | 0.004172 | 0.002938 | 0.004128 | 0.00865 | 0.002306 | 0.002862 | 0.005667 | 0.001073 |
| k__Bacteria;p__Firmicutes;c__Clostridia;o__Clostridiales;f__Lachnospiraceae;g__NA;s__sp33573 | 0 | 0 | 0 | 0.000447 | 0 | 0 | 0 | 0 | 0 | 0 | 0 | 0 | 0 | 0 | 0 | 0 | 0 | 0 |
| k__Bacteria;p__Firmicutes;c__Clostridia;o__Clostridiales;f__Lachnospiraceae;g__NA;s__sp33577 | 0.000764 | 0 | 0.001089 | 0 | 0 | 0.000825 | 0.003092 | 0.000745 | 0.000921 | 0 | 0.0004 | 0 | 0 | 0.001163 | 0 | 0 | 0.000954 | 0 |
| k__Bacteria;p__Firmicutes;c__Clostridia;o__Clostridiales;f__Lachnospiraceae;g__NA;s__sp33582 | 0.00129 | 0 | 0 | 0.001411 | 0 | 0.000825 | 0.004396 | 0.000882 | 0.001354 | 0.000718 | 0.001538 | 0.000524 | 0.001676 | 0.005976 | 0.002742 | 0.002014 | 0.006649 | 0.000624 |
| k__Bacteria;p__Firmicutes;c__Clostridia;o__Clostridiales;f__Lachnospiraceae;g__NA;s__sp33589 | 0 | 0 | 0 | 0 | 0 | 0 | 0 | 0 | 0 | 0 | 0 | 0 | 0 | 0 | 0 | 0 | 0.000477 | 0 |
| k__Bacteria;p__Firmicutes;c__Clostridia;o__Clostridiales;f__Lachnospiraceae;g__NA;s__sp33597 | 0 | 0 | 0.002641 | 0 | 0 | 0.000602 | 0 | 0 | 0.001354 | 0.000334 | 0 | 0 | 0 | 0 | 0.001068 | 0.000908 | 0.001852 | 0 |
| k__Bacteria;p__Firmicutes;c__Clostridia;o__Clostridiales;f__Lachnospiraceae;g__NA;s__sp33616-sp33639 | 0 | 0 | 0 | 0 | 0 | 0 | 0 | 0.000235 | 0 | 0 | 0 | 0 | 0.00049 | 0.000395 | 0 | 0.000434 | 0.002329 | 0 |
| k__Bacteria;p__Firmicutes;c__Clostridia;o__Clostridiales;f__Lachnospiraceae;g__NA;s__sp33620 | 0 | 0 | 0 | 0 | 0 | 0 | 0 | 0 | 0 | 0 | 0 | 0 | 0.000838 | 0.000767 | 0 | 0.000395 | 0 | 0 |
| k__Bacteria;p__Firmicutes;c__Clostridia;o__Clostridiales;f__Lachnospiraceae;g__NA;s__sp33625 | 0 | 0 | 0 | 0 | 0 | 0 | 0 | 0 | 0 | 0 | 0 | 0 | 0 | 0 | 0 | 0 | 0.000589 | 0 |
| k__Bacteria;p__Firmicutes;c__Clostridia;o__Clostridiales;f__Lachnospiraceae;g__NA;s__sp33636 | 0.000747 | 0 | 0 | 0 | 0 | 0 | 0.012369 | 0.000784 | 0 | 0.001219 | 0 | 0.001366 | 0.00045 | 0.004348 | 0.00034 | 0.000632 | 0.001347 | 0.0008 |
| k__Bacteria;p__Firmicutes;c__Clostridia;o__Clostridiales;f__Lachnospiraceae;g__NA;s__sp33639 | 0 | 0 | 0.000354 | 0 | 0 | 0.000275 | 0.001334 | 0 | 0.000786 | 0 | 0.000421 | 0 | 0 | 0 | 0 | 0 | 0 | 0 |
| k__Bacteria;p__Firmicutes;c__Clostridia;o__Clostridiales;f__Lachnospiraceae;g__NA;s__sp33645-sp33744 | 0.001273 | 0 | 0 | 0.000776 | 0.001241 | 0.000498 | 0.010277 | 0 | 0 | 0.000752 | 0.001032 | 0 | 0.002085 | 0.004232 | 0.004805 | 0.000572 | 0.00404 | 0.000546 |
| k__Bacteria;p__Firmicutes;c__Clostridia;o__Clostridiales;f__Lachnospiraceae;g__NA;s__sp33658 | 0 | 0 | 0.002232 | 0 | 0.001847 | 0.000481 | 0.005275 | 0.000882 | 0 | 0 | 0 | 0 | 0 | 0.001721 | 0.001262 | 0.001224 | 0.004377 | 0.000663 |
| k__Bacteria;p__Firmicutes;c__Clostridia;o__Clostridiales;f__Lachnospiraceae;g__NA;s__sp33669 | 0 | 0 | 0 | 0 | 0 | 0 | 0 | 0 | 0 | 0 | 0 | 0 | 0.000429 | 0 | 0 | 0 | 0 | 0 |
| k__Bacteria;p__Firmicutes;c__Clostridia;o__Clostridiales;f__Lachnospiraceae;g__NA;s__sp33670 | 0.00095 | 0.000607 | 0 | 0.000329 | 0.001516 | 0.001581 | 0.003911 | 0.001352 | 0.002384 | 0.000952 | 0.000822 | 0.000636 | 0.001778 | 0.003953 | 0.001869 | 0.002487 | 0.003956 | 0.001053 |
| k__Bacteria;p__Firmicutes;c__Clostridia;o__Clostridiales;f__Lachnospiraceae;g__NA;s__sp33673 | 0 | 0.000459 | 0 | 0 | 0 | 0 | 0 | 0 | 0 | 0 | 0 | 0 | 0 | 0.001093 | 0.000874 | 0.000671 | 0.002413 | 0 |
| k__Bacteria;p__Firmicutes;c__Clostridia;o__Clostridiales;f__Lachnospiraceae;g__NA;s__sp33674 | 0 | 0 | 0 | 0 | 0 | 0 | 0 | 0 | 0 | 0 | 0 | 0 | 0 | 0.000326 | 0 | 0 | 0 | 0 |
| k__Bacteria;p__Firmicutes;c__Clostridia;o__Clostridiales;f__Lachnospiraceae;g__NA;s__sp33682 | 0 | 0 | 0.000953 | 0 | 0 | 0 | 0 | 0.000412 | 0 | 0 | 0 | 0 | 0 | 0 | 0 | 0.000395 | 0.001936 | 0 |
| k__Bacteria;p__Firmicutes;c__Clostridia;o__Clostridiales;f__Lachnospiraceae;g__NA;s__sp33711 | 0 | 0.00023 | 0 | 0 | 0 | 0 | 0 | 0 | 0 | 0 | 0 | 0 | 0 | 0 | 0 | 0 | 0.002693 | 0 |
| k__Bacteria;p__Firmicutes;c__Clostridia;o__Clostridiales;f__Lachnospiraceae;g__NA;s__sp33714 | 0.000441 | 0 | 0.000953 | 0 | 0 | 0.000327 | 0.00479 | 0 | 0.001219 | 0.000418 | 0 | 0 | 0 | 0.000767 | 0 | 0 | 0 | 0 |
| k__Bacteria;p__Firmicutes;c__Clostridia;o__Clostridiales;f__Lachnospiraceae;g__NA;s__sp33718 | 0 | 0 | 0 | 0 | 0 | 0 | 0 | 0 | 0 | 0 | 0 | 0 | 0 | 0.001349 | 0 | 0 | 0 | 0 |
| k__Bacteria;p__Firmicutes;c__Clostridia;o__Clostridiales;f__Lachnospiraceae;g__NA;s__sp33721 | 0.000373 | 0.000509 | 0 | 0 | 0 | 0.00043 | 0.001455 | 0 | 0 | 0 | 0 | 0 | 0 | 0 | 0 | 0 | 0 | 0 |
| k__Bacteria;p__Firmicutes;c__Clostridia;o__Clostridiales;f__Lachnospiraceae;g__NA;s__sp33725 | 0 | 0 | 0 | 0 | 0 | 0 | 0 | 0 | 0 | 0 | 0 | 0 | 0.002657 | 0.001814 | 0.001505 | 0.004422 | 0.008052 | 0 |
| k__Bacteria;p__Firmicutes;c__Clostridia;o__Clostridiales;f__Lachnospiraceae;g__NA;s__sp33731 | 0 | 0 | 0 | 0 | 0 | 0 | 0 | 0 | 0 | 0 | 0 | 0 | 0.000879 | 0.002116 | 0.001019 | 0.000632 | 0.001291 | 0.000371 |
| k__Bacteria;p__Firmicutes;c__Clostridia;o__Clostridiales;f__Lachnospiraceae;g__NA;s__sp33733 | 0 | 0 | 0 | 0 | 0 | 0.001203 | 0 | 0 | 0 | 0 | 0 | 0 | 0 | 0 | 0 | 0 | 0 | 0 |
| k__Bacteria;p__Firmicutes;c__Clostridia;o__Clostridiales;f__Lachnospiraceae;g__NA;s__sp33734 | 0 | 0 | 0.004464 | 0.00367 | 0 | 0.001684 | 0.008276 | 0 | 0.00149 | 0 | 0 | 0 | 0.001471 | 0.002418 | 0 | 0.000671 | 0 | 0 |
| k__Bacteria;p__Firmicutes;c__Clostridia;o__Clostridiales;f__Lachnospiraceae;g__NA;s__sp33738 | 0 | 0 | 0 | 0 | 0 | 0 | 0 | 0 | 0 | 0 | 0 | 0 | 0.000429 | 0.00086 | 0.00051 | 0.000276 | 0 | 0 |
| k__Bacteria;p__Firmicutes;c__Clostridia;o__Clostridiales;f__Lachnospiraceae;g__NA;s__sp33755 | 0.000407 | 0 | 0 | 0 | 0.000662 | 0.002939 | 0.016249 | 0.000745 | 0.00065 | 0.001069 | 0.000674 | 0 | 0.002064 | 0.003046 | 0.001578 | 0.000908 | 0.00376 | 0 |
| k__Bacteria;p__Firmicutes;c__Clostridia;o__Clostridiales;f__Lachnospiraceae;g__NA;s__sp33756 | 0 | 0 | 0 | 0 | 0 | 0 | 0 | 0 | 0 | 0 | 0 | 0 | 0 | 0.001418 | 0 | 0 | 0 | 0 |
| k__Bacteria;p__Firmicutes;c__Clostridia;o__Clostridiales;f__Lachnospiraceae;g__NA;s__sp33760 | 0.000577 | 0 | 0.002532 | 0.002917 | 0.001461 | 0.004916 | 0.007882 | 0.000725 | 0.000677 | 0.000551 | 0.000421 | 0.000281 | 0.001615 | 0.002976 | 0.000777 | 0.001224 | 0.003002 | 0.000663 |
| k__Bacteria;p__Firmicutes;c__Clostridia;o__Clostridiales;f__Lachnospiraceae;g__NA;s__sp33766 | 0.002393 | 0.001969 | 0.028011 | 0.006375 | 0.008822 | 0.007374 | 0.029345 | 0.001882 | 0.009832 | 0.001052 | 0.001917 | 0.001441 | 0.00282 | 0.002488 | 0.002233 | 0.001737 | 0.018096 | 0.000488 |
| k__Bacteria;p__Firmicutes;c__Clostridia;o__Clostridiales;f__Lachnospiraceae;g__NA;s__sp33767 | 0.00353 | 0.001132 | 0.002014 | 0 | 0 | 0.001599 | 0 | 0.001784 | 0 | 0 | 0.001306 | 0 | 0 | 0 | 0 | 0 | 0 | 0 |
| k__Bacteria;p__Firmicutes;c__Clostridia;o__Clostridiales;f__Lachnospiraceae;g__Roseburia;s__sp33112 | 0.000305 | 0.001181 | 0.000708 | 0.000259 | 0 | 0.000808 | 0.009792 | 0.001294 | 0 | 0 | 0 | 0.000599 | 0 | 0 | 0 | 0 | 0 | 0 |
| k__Bacteria;p__Firmicutes;c__Clostridia;o__Clostridiales;f__Lachnospiraceae;g__Roseburia;s__sp33112-sp33140 | 0 | 0 | 0 | 0 | 0 | 0 | 0 | 0 | 0 | 0 | 0.000358 | 0 | 0 | 0 | 0 | 0 | 0 | 0 |
| k__Bacteria;p__Firmicutes;c__Clostridia;o__Clostridiales;f__Lachnospiraceae;g__Roseburia;s__sp33136 | 0 | 0 | 0 | 0 | 0 | 0 | 0 | 0 | 0.000515 | 0 | 0 | 0 | 0 | 0 | 0 | 0 | 0 | 0 |
| k__Bacteria;p__Firmicutes;c__Clostridia;o__Clostridiales;f__Lachnospiraceae;g__Roseburia;s__sp33140 | 0 | 0 | 0 | 0 | 0 | 0.000842 | 0 | 0 | 0 | 0 | 0 | 0 | 0.000838 | 0.001023 | 0 | 0.000651 | 0.008866 | 0.000527 |
| k__Bacteria;p__Firmicutes;c__Clostridia;o__Clostridiales;f__Lachnospiraceae;g__Roseburia;s__sp33143 | 0.003344 | 0.010401 | 0 | 0 | 0 | 0.001375 | 0.005548 | 0.001744 | 0 | 0 | 0.00236 | 0.000505 | 0 | 0.000512 | 0.000704 | 0.000533 | 0.002132 | 0 |
| k__Bacteria;p__Firmicutes;c__Clostridia;o__Clostridiales;f__Lachnospiraceae;g__Roseburia;s__sp33163 | 0 | 0.000246 | 0 | 0 | 0 | 0 | 0 | 0 | 0 | 0 | 0 | 0 | 0 | 0 | 0 | 0 | 0 | 0 |
| k__Bacteria;p__Firmicutes;c__Clostridia;o__Clostridiales;f__Lachnospiraceae;g__Roseburia;s__sp33164-sp33173 | 0 | 0 | 0 | 0 | 0 | 0 | 0 | 0 | 0 | 0 | 0 | 0 | 0 | 0 | 0.00034 | 0 | 0.000505 | 0 |
| k__Bacteria;p__Firmicutes;c__Clostridia;o__Clostridiales;f__Lachnospiraceae;g__Roseburia;s__sp33177 | 0 | 0 | 0.002695 | 0.009504 | 0.002481 | 0.002681 | 0 | 0 | 0 | 0 | 0.00099 | 0.000692 | 0.00049 | 0 | 0 | 0 | 0.002244 | 0 |
| k__Bacteria;p__Firmicutes;c__Clostridia;o__Clostridiales;f__Lachnospiraceae;g__Tyzzerella;s__sp33258-sp33301 | 0 | 0 | 0 | 0 | 0 | 0.000361 | 0 | 0.000666 | 0.000406 | 0 | 0 | 0.000243 | 0.000593 | 0 | 0.000291 | 0.000296 | 0.00188 | 0.000332 |
| k__Bacteria;p__Firmicutes;c__Clostridia;o__Clostridiales;f__Lachnospiraceae;g__Tyzzerella;s__sp33281 | 0 | 0 | 0 | 0 | 0 | 0.000344 | 0 | 0 | 0 | 0 | 0 | 0 | 0 | 0 | 0 | 0.000355 | 0 | 0 |
| k__Bacteria;p__Firmicutes;c__Clostridia;o__Clostridiales;f__Lachnospiraceae;g__Tyzzerella;s__sp33287 | 0 | 0 | 0.000844 | 0 | 0 | 0.000859 | 0 | 0 | 0 | 0.000217 | 0.000253 | 0 | 0 | 0.000744 | 0 | 0 | 0 | 0 |
| k__Bacteria;p__Firmicutes;c__Clostridia;o__Clostridiales;f__NA;g__NA;s__sp31072 | 0 | 0 | 0.002695 | 0 | 0 | 0.000602 | 0 | 0 | 0.000433 | 0.000334 | 0 | 0 | 0 | 0 | 0 | 0 | 0.000365 | 0 |
| k__Bacteria;p__Firmicutes;c__Clostridia;o__Clostridiales;f__NA;g__NA;s__sp31075 | 0 | 0 | 0 | 0 | 0 | 0 | 0 | 0 | 0 | 0 | 0.000337 | 0 | 0 | 0.000372 | 0 | 0 | 0 | 0 |
| k__Bacteria;p__Firmicutes;c__Clostridia;o__Clostridiales;f__NA;g__NA;s__sp31082 | 0 | 0 | 0.000871 | 0 | 0 | 0.00055 | 0.000576 | 0 | 0 | 0.000601 | 0.001896 | 0.00043 | 0.001369 | 0.000419 | 0 | 0 | 0.007267 | 0.000468 |
| k__Bacteria;p__Firmicutes;c__Clostridia;o__Clostridiales;f__NA;g__NA;s__sp31094 | 0 | 0 | 0 | 0 | 0 | 0 | 0 | 0 | 0 | 0 | 0 | 0 | 0.000266 | 0 | 0 | 0 | 0 | 0 |
| k__Bacteria;p__Firmicutes;c__Clostridia;o__Clostridiales;f__NA;g__NA;s__sp31115 | 0 | 0 | 0 | 0 | 0 | 0 | 0 | 0 | 0.001679 | 0.001888 | 0.00217 | 0 | 0.003576 | 0.003256 | 0.004053 | 0.004185 | 0.00491 | 0.003354 |
| k__Bacteria;p__Firmicutes;c__Clostridia;o__Clostridiales;f__NA;g__NA;s__sp31116 | 0 | 0 | 0 | 0 | 0 | 0 | 0 | 0 | 0 | 0.000735 | 0 | 0 | 0 | 0 | 0 | 0 | 0 | 0 |
| k__Bacteria;p__Firmicutes;c__Clostridia;o__Clostridiales;f__NA;g__NA;s__sp31119 | 0 | 0 | 0 | 0 | 0.001213 | 0.000464 | 0 | 0 | 0.005038 | 0.006832 | 0.015359 | 0.008402 | 0.000286 | 0 | 0 | 0.001066 | 0.001403 | 0.001892 |
| k__Bacteria;p__Firmicutes;c__Clostridia;o__Clostridiales;f__Peptococcaceae;g__NA;s__sp34196 | 0 | 0 | 0 | 0 | 0 | 0 | 0 | 0.000274 | 0 | 0 | 0 | 0 | 0 | 0 | 0 | 0 | 0 | 0 |
| k__Bacteria;p__Firmicutes;c__Clostridia;o__Clostridiales;f__Peptococcaceae;g__NA;s__sp34246 | 0 | 0 | 0.000354 | 0 | 0.000303 | 0.000756 | 0 | 0 | 0 | 0.000217 | 0 | 0 | 0.00049 | 0.001023 | 0.000849 | 0.000849 | 0.001459 | 0 |
| k__Bacteria;p__Firmicutes;c__Clostridia;o__Clostridiales;f__Peptococcaceae;g__Peptococcus;s__sp34117 | 0 | 0 | 0.000517 | 0 | 0 | 0 | 0.000394 | 0 | 0 | 0 | 0.000253 | 0 | 0 | 0.000512 | 0 | 0 | 0.000337 | 0 |
| k__Bacteria;p__Firmicutes;c__Clostridia;o__Clostridiales;f__Peptostreptococcaceae;g__Peptoclostridium;s__difficile | 0.000187 | 0 | 0 | 0.029877 | 0 | 0 | 0 | 0 | 0 | 0 | 0 | 0 | 0 | 0 | 0 | 0 | 0 | 0 |
| k__Bacteria;p__Firmicutes;c__Clostridia;o__Clostridiales;f__Peptostreptococcaceae;g__Romboutsia;s__ilealis | 0 | 0 | 0 | 0 | 0 | 0 | 0 | 0 | 0 | 0 | 0 | 0 | 0 | 0 | 0.004466 | 0 | 0 | 0 |
| k__Bacteria;p__Firmicutes;c__Clostridia;o__Clostridiales;f__Ruminococcaceae;g__Anaerotruncus;s__colihominis | 0.000441 | 0 | 0.001633 | 0.000306 | 0 | 0.000928 | 0.001698 | 0 | 0 | 0 | 0.000295 | 0 | 0.000245 | 0.000721 | 0 | 0 | 0.000701 | 0 |
| k__Bacteria;p__Firmicutes;c__Clostridia;o__Clostridiales;f__Ruminococcaceae;g__Anaerotruncus;s__sp34438 | 0 | 0 | 0 | 0 | 0 | 0 | 0.001273 | 0 | 0 | 0 | 0 | 0 | 0.000266 | 0.000488 | 0 | 0 | 0 | 0 |
| k__Bacteria;p__Firmicutes;c__Clostridia;o__Clostridiales;f__Ruminococcaceae;g__Anaerotruncus;s__sp34443 | 0.000255 | 0 | 0 | 0 | 0 | 0 | 0.000394 | 0 | 0 | 0 | 0 | 0 | 0 | 0 | 0 | 0 | 0 | 0 |
| k__Bacteria;p__Firmicutes;c__Clostridia;o__Clostridiales;f__Ruminococcaceae;g__Anaerotruncus;s__sp34452-sp34475 | 0.000509 | 0 | 0 | 0 | 0 | 0 | 0.007245 | 0.002313 | 0 | 0 | 0 | 0 | 0 | 0 | 0 | 0 | 0 | 0 |
| k__Bacteria;p__Firmicutes;c__Clostridia;o__Clostridiales;f__Ruminococcaceae;g__Anaerotruncus;s__sp34453 | 0.001256 | 0.00233 | 0.006071 | 0.004305 | 0.000606 | 0.003283 | 0.010641 | 0.001117 | 0.001896 | 0.000585 | 0.001496 | 0.000823 | 0.002595 | 0.002442 | 0.001432 | 0.00075 | 0.005527 | 0.000819 |
| k__Bacteria;p__Firmicutes;c__Clostridia;o__Clostridiales;f__Ruminococcaceae;g__Anaerotruncus;s__sp34462 | 0 | 0 | 0 | 0 | 0 | 0 | 0 | 0 | 0 | 0 | 0.000442 | 0 | 0.001226 | 0.002883 | 0.00148 | 0.000829 | 0.001796 | 0.000254 |

| | | | | | | | | | | | | | | | | | | |
|---|---|---|---|---|---|---|---|---|---|---|---|---|---|---|---|---|---|---|
| k__Bacteria;p__Firmicutes;c__Clostridia;o__Clostridiales;f__Ruminococcaceae;g__Anaerotruncus;s__sp34462-sp35834 | 0 | 0 | 0 | 0 | 0 | 0 | 0 | 0 | 0 | 0 | 0 | 0 | 0 | 0 | 0.000388 | 0 | 0 | 0 |
| k__Bacteria;p__Firmicutes;c__Clostridia;o__Clostridiales;f__Ruminococcaceae;g__Anaerotruncus;s__sp34466 | 0 | 0 | 0 | 0 | 0 | 0 | 0 | 0 | 0 | 0 | 0 | 0 | 0 | 0 | 0.001335 | 0 | 0 | 0.001365 |
| k__Bacteria;p__Firmicutes;c__Clostridia;o__Clostridiales;f__Ruminococcaceae;g__Anaerotruncus;s__sp34471 | 0 | 0 | 0.002314 | 0 | 0 | 0.000602 | 0 | 0 | 0.00065 | 0 | 0.00059 | 0 | 0 | 0 | 0 | 0.000592 | 0.001627 | 0 |
| k__Bacteria;p__Firmicutes;c__Clostridia;o__Clostridiales;f__Ruminococcaceae;g__Anaerotruncus;s__sp34471-sp34483 | 0 | 0 | 0 | 0 | 0 | 0 | 0 | 0 | 0 | 0 | 0 | 0 | 0.000327 | 0 | 0 | 0 | 0 | 0 |
| k__Bacteria;p__Firmicutes;c__Clostridia;o__Clostridiales;f__Ruminococcaceae;g__Anaerotruncus;s__sp34475 | 0.000305 | 0 | 0.004519 | 0.005552 | 0.001323 | 0.001083 | 0.002395 | 0 | 0.002221 | 0.00142 | 0.001327 | 0.000206 | 0.001696 | 0.001395 | 0 | 0.001757 | 0.004601 | 0 |
| k__Bacteria;p__Firmicutes;c__Clostridia;o__Clostridiales;f__Ruminococcaceae;g__Anaerotruncus;s__sp34481 | 0 | 0 | 0 | 0 | 0 | 0 | 0.000697 | 0 | 0 | 0 | 0 | 0 | 0 | 0 | 0 | 0 | 0 | 0 |
| k__Bacteria;p__Firmicutes;c__Clostridia;o__Clostridiales;f__Ruminococcaceae;g__Anaerotruncus;s__sp34483 | 0 | 0 | 0 | 0 | 0.000634 | 0.001031 | 0.002941 | 0 | 0 | 0 | 0 | 0 | 0 | 0 | 0.00051 | 0 | 0 | 0 |
| k__Bacteria;p__Firmicutes;c__Clostridia;o__Clostridiales;f__Ruminococcaceae;g__Faecalibacterium;s__prausnitzii | 0 | 0 | 0 | 0 | 0 | 0 | 0 | 0 | 0 | 0 | 0 | 0 | 0 | 0 | 0.015387 | 0 | 0 | 0 |
| k__Bacteria;p__Firmicutes;c__Clostridia;o__Clostridiales;f__Ruminococcaceae;g__Faecalibacterium;s__sp34544 | 0 | 0 | 0 | 0 | 0 | 0 | 0 | 0 | 0 | 0 | 0 | 0 | 0 | 0 | 0.001505 | 0 | 0 | 0 |
| k__Bacteria;p__Firmicutes;c__Clostridia;o__Clostridiales;f__Ruminococcaceae;g__NA;s__sp33140 | 0 | 0 | 0.000544 | 0 | 0 | 0.00067 | 0 | 0 | 0.000731 | 0 | 0 | 0 | 0.000531 | 0.001674 | 0.001068 | 0.000355 | 0.001571 | 0.000312 |
| k__Bacteria;p__Firmicutes;c__Clostridia;o__Clostridiales;f__Ruminococcaceae;g__NA;s__sp34731 | 0.001069 | 0.000574 | 0 | 0 | 0 | 0 | 0 | 0.002411 | 0 | 0 | 0 | 0 | 0 | 0 | 0 | 0 | 0 | 0 |
| k__Bacteria;p__Firmicutes;c__Clostridia;o__Clostridiales;f__Ruminococcaceae;g__NA;s__sp34771 | 0 | 0 | 0 | 0 | 0.000469 | 0 | 0 | 0 | 0 | 0.000768 | 0 | 0 | 0 | 0 | 0 | 0 | 0 | 0 |
| k__Bacteria;p__Firmicutes;c__Clostridia;o__Clostridiales;f__Ruminococcaceae;g__NA;s__sp34820 | 0 | 0 | 0.003076 | 0 | 0 | 0.002836 | 0.00191 | 0 | 0.000433 | 0.000384 | 0.000548 | 0 | 0 | 0.000977 | 0.001189 | 0 | 0.001066 | 0 |
| k__Bacteria;p__Firmicutes;c__Clostridia;o__Clostridiales;f__Ruminococcaceae;g__NA;s__sp34822 | 0 | 0 | 0 | 0 | 0 | 0.000223 | 0 | 0 | 0 | 0 | 0 | 0 | 0 | 0 | 0 | 0 | 0 | 0 |
| k__Bacteria;p__Firmicutes;c__Clostridia;o__Clostridiales;f__Ruminococcaceae;g__NA;s__sp34826 | 0 | 0 | 0 | 0 | 0.000744 | 0 | 0 | 0 | 0 | 0 | 0 | 0 | 0.002882 | 0.018417 | 0.011989 | 0.008883 | 0.018545 | 0.008424 |
| k__Bacteria;p__Firmicutes;c__Clostridia;o__Clostridiales;f__Ruminococcaceae;g__NA;s__sp34838 | 0 | 0 | 0 | 0 | 0.002178 | 0 | 0 | 0 | 0.001598 | 0.001069 | 0.000758 | 0 | 0.001451 | 0.000907 | 0 | 0 | 0.000645 | 0 |
| k__Bacteria;p__Firmicutes;c__Clostridia;o__Clostridiales;f__Ruminococcaceae;g__NA;s__sp34858-sp34873 | 0 | 0.000377 | 0.002831 | 0.002541 | 0.001434 | 0.001839 | 0.003153 | 0 | 0 | 0 | 0 | 0 | 0.001962 | 0.003442 | 0.002621 | 0.001066 | 0.00592 | 0.001229 |
| k__Bacteria;p__Firmicutes;c__Clostridia;o__Clostridiales;f__Ruminococcaceae;g__NA;s__sp34863 | 0 | 0 | 0 | 0 | 0 | 0 | 0 | 0.000314 | 0 | 0 | 0 | 0 | 0.000838 | 0.000884 | 0 | 0.000632 | 0 | 0 |
| k__Bacteria;p__Firmicutes;c__Clostridia;o__Clostridiales;f__Ruminococcaceae;g__NA;s__sp34871 | 0 | 0 | 0 | 0 | 0 | 0.001358 | 0 | 0 | 0 | 0 | 0 | 0 | 0 | 0 | 0 | 0 | 0.001403 | 0 |
| k__Bacteria;p__Firmicutes;c__Clostridia;o__Clostridiales;f__Ruminococcaceae;g__NA;s__sp34875-sp34878 | 0 | 0 | 0 | 0 | 0 | 0 | 0 | 0 | 0.00149 | 0 | 0 | 0 | 0 | 0 | 0 | 0 | 0 | 0 |
| k__Bacteria;p__Firmicutes;c__Clostridia;o__Clostridiales;f__Ruminococcaceae;g__NA;s__sp34878 | 0.003581 | 0.001887 | 0.00539 | 0 | 0.001268 | 0.000602 | 0.005487 | 0.000921 | 0.001463 | 0.002406 | 0.00158 | 0.000711 | 0.001942 | 0.002581 | 0.003373 | 0.001777 | 0.002749 | 0.000488 |
| k__Bacteria;p__Firmicutes;c__Clostridia;o__Clostridiales;f__Ruminococcaceae;g__NA;s__sp34878-sp34883 | 0 | 0 | 0 | 0 | 0 | 0 | 0 | 0 | 0 | 0 | 0 | 0 | 0 | 0 | 0 | 0 | 0.001796 | 0 |
| k__Bacteria;p__Firmicutes;c__Clostridia;o__Clostridiales;f__Ruminococcaceae;g__NA;s__sp34879 | 0 | 0.000377 | 0 | 0 | 0.001406 | 0.001805 | 0.001334 | 0.00198 | 0 | 0.00162 | 0.002212 | 0.000299 | 0.002391 | 0.003442 | 0.002888 | 0.002882 | 0.001739 | 0.000917 |
| k__Bacteria;p__Firmicutes;c__Clostridia;o__Clostridiales;f__Ruminococcaceae;g__NA;s__sp34883 | 0 | 0 | 0 | 0 | 0 | 0 | 0 | 0 | 0 | 0 | 0 | 0 | 0 | 0.000628 | 0 | 0 | 0 | 0 |
| k__Bacteria;p__Firmicutes;c__Clostridia;o__Clostridiales;f__Ruminococcaceae;g__NA;s__sp34977 | 0 | 0 | 0 | 0 | 0 | 0 | 0 | 0 | 0 | 0.000384 | 0 | 0 | 0.00045 | 0.000744 | 0.000582 | 0.000651 | 0.000673 | 0.000663 |
| k__Bacteria;p__Firmicutes;c__Clostridia;o__Clostridiales;f__Ruminococcaceae;g__NA;s__sp34983 | 0 | 0 | 0 | 0 | 0 | 0 | 0 | 0 | 0 | 0 | 0 | 0 | 0 | 0.000349 | 0 | 0 | 0.000309 | 0 |
| k__Bacteria;p__Firmicutes;c__Clostridia;o__Clostridiales;f__Ruminococcaceae;g__NA;s__sp35181 | 0 | 0 | 0 | 0 | 0 | 0 | 0 | 0 | 0 | 0 | 0 | 0 | 0.000756 | 0.000605 | 0.000558 | 0 | 0.000505 | 0.000975 |
| k__Bacteria;p__Firmicutes;c__Clostridia;o__Clostridiales;f__Ruminococcaceae;g__NA;s__sp35360 | 0 | 0 | 0 | 0 | 0 | 0 | 0 | 0 | 0 | 0 | 0 | 0 | 0 | 0.000512 | 0 | 0.000533 | 0 | 0.000273 |
| k__Bacteria;p__Firmicutes;c__Clostridia;o__Clostridiales;f__Ruminococcaceae;g__NA;s__sp35382-sp35403 | 0 | 0 | 0 | 0 | 0 | 0 | 0 | 0 | 0.003413 | 0.000735 | 0.00276 | 0.001048 | 0 | 0.002651 | 0.000728 | 0 | 0.001487 | 0 |
| k__Bacteria;p__Firmicutes;c__Clostridia;o__Clostridiales;f__Ruminococcaceae;g__NA;s__sp35383 | 0 | 0 | 0 | 0 | 0 | 0 | 0 | 0 | 0 | 0 | 0 | 0 | 0 | 0.000302 | 0 | 0 | 0 | 0 |
| k__Bacteria;p__Firmicutes;c__Clostridia;o__Clostridiales;f__Ruminococcaceae;g__NA;s__sp35384-sp35410 | 0 | 0 | 0 | 0 | 0 | 0 | 0 | 0 | 0 | 0 | 0 | 0 | 0.000531 | 0 | 0 | 0 | 0 | 0 |
| k__Bacteria;p__Firmicutes;c__Clostridia;o__Clostridiales;f__Ruminococcaceae;g__NA;s__sp35388 | 0 | 0 | 0.001143 | 0 | 0 | 0 | 0 | 0 | 0 | 0.000785 | 0 | 0 | 0.000613 | 0 | 0 | 0.001283 | 0 | 0 |
| k__Bacteria;p__Firmicutes;c__Clostridia;o__Clostridiales;f__Ruminococcaceae;g__NA;s__sp35392 | 0 | 0 | 0 | 0 | 0 | 0 | 0 | 0 | 0 | 0 | 0 | 0 | 0 | 0 | 0 | 0.000336 | 0 | 0 |
| k__Bacteria;p__Firmicutes;c__Clostridia;o__Clostridiales;f__Ruminococcaceae;g__NA;s__sp35403 | 0 | 0 | 0 | 0 | 0 | 0 | 0 | 0 | 0 | 0 | 0 | 0 | 0.000777 | 0.001093 | 0 | 0.000908 | 0 | 0.001092 |
| k__Bacteria;p__Firmicutes;c__Clostridia;o__Clostridiales;f__Ruminococcaceae;g__NA;s__sp35419 | 0 | 0 | 0 | 0 | 0 | 0 | 0 | 0 | 0 | 0 | 0 | 0 | 0 | 0 | 0 | 0.001461 | 0 | 0.000254 |
| k__Bacteria;p__Firmicutes;c__Clostridia;o__Clostridiales;f__Ruminococcaceae;g__NA;s__sp35423 | 0 | 0 | 0 | 0 | 0.001654 | 0 | 0 | 0 | 0 | 0 | 0.002739 | 0 | 0 | 0 | 0 | 0.000355 | 0 | 0.000546 |
| k__Bacteria;p__Firmicutes;c__Clostridia;o__Clostridiales;f__Ruminococcaceae;g__NA;s__sp35487 | 0 | 0 | 0 | 0 | 0 | 0 | 0 | 0 | 0 | 0 | 0 | 0 | 0 | 0 | 0.000971 | 0 | 0 | 0 |
| k__Bacteria;p__Firmicutes;c__Clostridia;o__Clostridiales;f__Ruminococcaceae;g__NA;s__sp35494 | 0 | 0 | 0 | 0 | 0.000689 | 0 | 0 | 0 | 0 | 0 | 0 | 0 | 0.000838 | 0.002256 | 0.001772 | 0.004343 | 0.005218 | 0.008541 |
| k__Bacteria;p__Firmicutes;c__Clostridia;o__Clostridiales;f__Ruminococcaceae;g__NA;s__sp35598 | 0.002987 | 0.008432 | 0 | 0 | 0 | 0 | 0 | 0.002019 | 0 | 0 | 0 | 0 | 0.001737 | 0.002163 | 0.001505 | 0 | 0 | 0 |
| k__Bacteria;p__Firmicutes;c__Clostridia;o__Clostridiales;f__Ruminococcaceae;g__NA;s__sp35669 | 0 | 0 | 0 | 0 | 0 | 0 | 0 | 0 | 0 | 0 | 0 | 0 | 0 | 0 | 0.000461 | 0 | 0 | 0 |
| k__Bacteria;p__Firmicutes;c__Clostridia;o__Clostridiales;f__Ruminococcaceae;g__NA;s__sp35688-sp35748 | 0 | 0 | 0 | 0 | 0.00091 | 0.000773 | 0 | 0 | 0 | 0 | 0 | 0 | 0 | 0.001511 | 0.000922 | 0.000494 | 0.001375 | 0 |
| k__Bacteria;p__Firmicutes;c__Clostridia;o__Clostridiales;f__Ruminococcaceae;g__NA;s__sp35693 | 0 | 0 | 0.004519 | 0 | 0 | 0.006755 | 0.006487 | 0 | 0.004117 | 0.001086 | 0 | 0 | 0.001104 | 0.006325 | 0.004878 | 0.001994 | 0.009202 | 0.001365 |
| k__Bacteria;p__Firmicutes;c__Clostridia;o__Clostridiales;f__Ruminococcaceae;g__NA;s__sp35696 | 0 | 0 | 0.008275 | 0.001012 | 0.001213 | 0.003438 | 0 | 0 | 0 | 0 | 0 | 0 | 0 | 0.00107 | 0.000898 | 0.000809 | 0 | 0.001034 |
| k__Bacteria;p__Firmicutes;c__Clostridia;o__Clostridiales;f__Ruminococcaceae;g__NA;s__sp35730 | 0 | 0 | 0.001388 | 0 | 0.000303 | 0.000894 | 0.002152 | 0 | 0 | 0 | 0 | 0 | 0.00047 | 0.000907 | 0.000995 | 0.000217 | 0.002441 | 0.000293 |
| k__Bacteria;p__Firmicutes;c__Clostridia;o__Clostridiales;f__Ruminococcaceae;g__NA;s__sp35733 | 0.000883 | 0.002067 | 0.002886 | 0.000965 | 0.003777 | 0.006549 | 0.000909 | 0 | 0.002275 | 0.001387 | 0.00375 | 0.000487 | 0.000674 | 0.001605 | 0.000971 | 0.000454 | 0.001403 | 0.000585 |
| k__Bacteria;p__Firmicutes;c__Clostridia;o__Clostridiales;f__Ruminococcaceae;g__NA;s__sp35748 | 0 | 0 | 0.001443 | 0 | 0.001075 | 0.001599 | 0.003638 | 0 | 0.001977 | 0 | 0.00078 | 0 | 0.002289 | 0.001698 | 0.00165 | 0.001342 | 0.001824 | 0.000488 |
| k__Bacteria;p__Firmicutes;c__Clostridia;o__Clostridiales;f__Ruminococcaceae;g__NA;s__sp35799 | 0 | 0 | 0 | 0 | 0.001351 | 0.000739 | 0.003638 | 0 | 0 | 0 | 0 | 0 | 0.000368 | 0.001186 | 0 | 0.000355 | 0.001263 | 0 |
| k__Bacteria;p__Firmicutes;c__Clostridia;o__Clostridiales;f__Ruminococcaceae;g__NA;s__sp35832 | 0 | 0 | 0 | 0 | 0 | 0.000894 | 0 | 0 | 0 | 0 | 0 | 0 | 0 | 0 | 0 | 0 | 0.000926 | 0 |
| k__Bacteria;p__Firmicutes;c__Clostridia;o__Clostridiales;f__Ruminococcaceae;g__NA;s__sp35841 | 0.000679 | 0 | 0.003974 | 0.000518 | 0.00113 | 0.004675 | 0.013066 | 0.000902 | 0 | 0 | 0.0004 | 0 | 0.00188 | 0.001209 | 0.002209 | 0.000651 | 0.004433 | 0.000624 |
| k__Bacteria;p__Firmicutes;c__Clostridia;o__Clostridiales;f__Ruminococcaceae;g__NA;s__sp35849 | 0 | 0 | 0 | 0 | 0 | 0 | 0.001546 | 0.000255 | 0 | 0 | 0 | 0 | 0 | 0 | 0 | 0 | 0.000505 | 0 |
| k__Bacteria;p__Firmicutes;c__Clostridia;o__Clostridiales;f__Ruminococcaceae;g__NA;s__sp35869 | 0.001935 | 0 | 0.003729 | 0 | 0 | 0 | 0.007427 | 0.001725 | 0 | 0.000601 | 0 | 0 | 0 | 0.002 | 0.00182 | 0.000809 | 0.003058 | 0.000741 |
| k__Bacteria;p__Firmicutes;c__Clostridia;o__Clostridiales;f__Ruminococcaceae;g__NA;s__sp35873 | 0 | 0 | 0 | 0 | 0 | 0.000533 | 0 | 0 | 0 | 0 | 0 | 0 | 0 | 0 | 0 | 0 | 0 | 0 |
| k__Bacteria;p__Firmicutes;c__Clostridia;o__Clostridiales;f__Ruminococcaceae;g__NA;s__sp35877 | 0 | 0 | 0.000327 | 0 | 0 | 0 | 0.00388 | 0.00047 | 0 | 0 | 0 | 0 | 0.000777 | 0 | 0 | 0.000375 | 0.001459 | 0.000761 |
| k__Bacteria;p__Firmicutes;c__Clostridia;o__Clostridiales;f__Ruminococcaceae;g__NA;s__sp35882 | 0 | 0 | 0 | 0.000894 | 0 | 0.008766 | 0 | 0 | 0 | 0 | 0.002086 | 0.001254 | 0 | 0 | 0.000558 | 0.000276 | 0.001543 | 0.000215 |
| k__Bacteria;p__Firmicutes;c__Clostridia;o__Clostridiales;f__Ruminococcaceae;g__Oscillibacter;s__sp34634 | 0 | 0.000902 | 0.005417 | 0.000659 | 0.00306 | 0.002939 | 0.014127 | 0.001235 | 0.001029 | 0.000986 | 0.001011 | 0 | 0.004312 | 0.010139 | 0.004344 | 0.002112 | 0.011194 | 0.001287 |
| k__Bacteria;p__Firmicutes;c__Clostridia;o__Clostridiales;f__Ruminococcaceae;g__Oscillibacter;s__sp34638 | 0.000356 | 0 | 0 | 0 | 0 | 0 | 0 | 0.000431 | 0 | 0 | 0 | 0 | 0 | 0 | 0 | 0 | 0.000898 | 0 |
| k__Bacteria;p__Firmicutes;c__Clostridia;o__Clostridiales;f__Ruminococcaceae;g__Oscillibacter;s__sp34640 | 0.002749 | 0.000738 | 0.002069 | 0 | 0 | 0.003025 | 0 | 0.000666 | 0.003007 | 0.001503 | 0.000632 | 0 | 0.000654 | 0 | 0.001213 | 0 | 0.002693 | 0 |
| k__Bacteria;p__Firmicutes;c__Clostridia;o__Clostridiales;f__Ruminococcaceae;g__Oscillibacter;s__sp34640-sp34643 | 0 | 0 | 0 | 0 | 0 | 0 | 0 | 0 | 0 | 0 | 0 | 0 | 0 | 0.00107 | 0 | 0.000948 | 0 | 0 |
| k__Bacteria;p__Firmicutes;c__Clostridia;o__Clostridiales;f__Ruminococcaceae;g__Oscillibacter;s__sp34642 | 0.003208 | 0.00064 | 0 | 0 | 0 | 0 | 0.00291 | 0 | 0 | 0 | 0 | 0 | 0 | 0 | 0 | 0 | 0.003142 | 0.000702 |
| k__Bacteria;p__Firmicutes;c__Clostridia;o__Clostridiales;f__Ruminococcaceae;g__Oscillibacter;s__sp34643 | 0 | 0 | 0 | 0 | 0.000882 | 0 | 0.007912 | 0 | 0 | 0.001554 | 0 | 0 | 0.00047 | 0.001791 | 0 | 0 | 0 | 0 |
| k__Bacteria;p__Firmicutes;c__Clostridia;o__Clostridiales;f__Ruminococcaceae;g__Oscillibacter;s__sp34645 | 0.000204 | 0.000738 | 0 | 0 | 0 | 0.000241 | 0.001122 | 0.000431 | 0 | 0 | 0 | 0 | 0 | 0 | 0 | 0 | 0.000561 | 0 |
| k__Bacteria;p__Firmicutes;c__Clostridia;o__Clostridiales;f__Ruminococcaceae;g__Oscillibacter;s__sp34646 | 0 | 0 | 0 | 0 | 0 | 0 | 0 | 0 | 0 | 0 | 0 | 0 | 0 | 0 | 0 | 0 | 0.001066 | 0 |
| k__Bacteria;p__Firmicutes;c__Clostridia;o__Clostridiales;f__Ruminococcaceae;g__Oscillibacter;s__sp34648 | 0 | 0 | 0.001443 | 0.000306 | 0 | 0.000963 | 0 | 0 | 0 | 0 | 0 | 0 | 0 | 0 | 0 | 0 | 0 | 0 |
| k__Bacteria;p__Firmicutes;c__Clostridia;o__Clostridiales;f__Ruminococcaceae;g__Oscillibacter-Oscillospira;s__sp34648-sp34660 | 0 | 0 | 0 | 0 | 0 | 0.00067 | 0 | 0 | 0 | 0 | 0 | 0 | 0.001124 | 0.001837 | 0.000947 | 0.000474 | 0.002946 | 0.000605 |
| k__Bacteria;p__Firmicutes;c__Clostridia;o__Clostridiales;f__Ruminococcaceae;g__Ruminiclostridium;s__sp34857-sp34950 | 0.002342 | 0 | 0 | 0 | 0 | 0 | 0.012156 | 0.00198 | 0 | 0 | 0 | 0 | 0 | 0 | 0 | 0 | 0 | 0 |
| k__Bacteria;p__Firmicutes;c__Clostridia;o__Clostridiales;f__Ruminococcaceae;g__Ruminiclostridium;s__sp34858-sp34873-sp34950 | 0 | 0 | 0.00196 | 0 | 0 | 0 | 0 | 0 | 0 | 0 | 0.001011 | 0 | 0 | 0.000628 | 0 | 0 | 0 | 0 |
| k__Bacteria;p__Firmicutes;c__Clostridia;o__Clostridiales;f__Ruminococcaceae;g__Ruminiclostridium;s__sp34901 | 0.00572 | 0 | 0 | 0 | 0 | 0 | 0 | 0 | 0 | 0 | 0 | 0 | 0 | 0.001279 | 0 | 0 | 0 | 0 |
| k__Bacteria;p__Firmicutes;c__Clostridia;o__Clostridiales;f__Ruminococcaceae;g__Ruminiclostridium;s__sp34925 | 0 | 0 | 0.005853 | 0.000706 | 0.001571 | 0.005896 | 0.012035 | 0.001215 | 0.000948 | 0.000317 | 0.00059 | 0.000318 | 0.001574 | 0.00193 | 0.001141 | 0.000671 | 0.009539 | 0 |
| k__Bacteria;p__Firmicutes;c__Clostridia;o__Clostridiales;f__Ruminococcaceae;g__Ruminococcus;s__bromii | 0 | 0 | 0 | 0 | 0 | 0 | 0 | 0 | 0 | 0 | 0 | 0 | 0 | 0 | 0.003665 | 0 | 0 | 0 |
| k__Bacteria;p__Firmicutes;c__Clostridia;o__Clostridiales;f__Ruminococcaceae;g__Subdoligranulum;s__sp35580-sp35585 | 0 | 0 | 0 | 0 | 0 | 0 | 0 | 0 | 0 | 0 | 0 | 0 | 0 | 0 | 0.001942 | 0 | 0 | 0 |
| k__Bacteria;p__Firmicutes;c__Clostridia;o__Clostridiales;f__Ruminococcaceae;g__Subdoligranulum;s__sp35582 | 0 | 0 | 0 | 0 | 0 | 0 | 0 | 0 | 0 | 0 | 0 | 0 | 0 | 0 | 0.002014 | 0 | 0 | 0 |
| k__Bacteria;p__Firmicutes;c__Clostridia;o__Clostridiales;f__Ruminococcaceae;g__Subdoligranulum;s__sp35590 | 0 | 0 | 0 | 0 | 0 | 0 | 0 | 0 | 0 | 0 | 0 | 0 | 0.000572 | 0.001791 | 0.000534 | 0.000869 | 0.003928 | 0.000995 |
| k__Bacteria;p__Firmicutes;c__Erysipelotrichia;o__Erysipelotrichales;f__Erysipelotrichaceae;g__Allobaculum;s__sp36555 | 0.000984 | 0.003445 | 0 | 0 | 0 | 0 | 0 | 0.001254 | 0 | 0 | 0 | 0 | 0 | 0 | 0 | 0 | 0 | 0.003218 |
| k__Bacteria;p__Firmicutes;c__Erysipelotrichia;o__Erysipelotrichales;f__Erysipelotrichaceae;g__NA;s__sp36686-sp36689 | 0 | 0 | 0 | 0 | 0 | 0 | 0 | 0 | 0 | 0 | 0 | 0 | 0 | 0 | 0.005194 | 0 | 0 | 0 |
| k__Bacteria;p__Firmicutes;c__Erysipelotrichia;o__Erysipelotrichales;f__Erysipelotrichaceae;g__NA;s__sp36777 | 0 | 0 | 0 | 0.000541 | 0 | 0 | 0 | 0.000314 | 0 | 0.0002 | 0.000632 | 0.000823 | 0 | 0.000767 | 0 | 0 | 0 | 0.000605 |
| k__Bacteria;p__Firmicutes;c__Erysipelotrichia;o__Erysipelotrichales;f__Erysipelotrichaceae;g__NA;s__sp36787 | 0 | 0 | 0 | 0 | 0 | 0 | 0 | 0 | 0 | 0.000468 | 0 | 0 | 0.00047 | 0 | 0.00034 | 0.000711 | 0 | 0 |
| k__Bacteria;p__Firmicutes;c__Erysipelotrichia;o__Erysipelotrichales;f__Erysipelotrichaceae;g__Turicibacter;s__sanguinis | 0 | 0 | 0.007241 | 0.007552 | 0.011468 | 0.000361 | 0 | 0 | 0.051247 | 0.034029 | 0.039336 | 0.027434 | 0.010362 | 0.018208 | 0.019828 | 0.04343 | 0.026597 | 0.02227 |
| k__Bacteria;p__Firmicutes;c__Negativicutes;o__Selenomonadales;f__Acidaminococcaceae;g__Phascolarctobacterium;s__sp36830 | 0 | 0 | 0 | 0 | 0 | 0 | 0 | 0 | 0 | 0 | 0 | 0 | 0 | 0 | 0.001917 | 0 | 0 | 0 |
| k__Bacteria;p__Proteobacteria;c__Alphaproteobacteria;o__Rhodospirillales;f__Rhodospirillaceae;g__Thalassospira;s__sp46235 | 0.004684 | 0.001821 | 0 | 0 | 0 | 0 | 0 | 0 | 0.001706 | 0.000752 | 0.006658 | 0 | 0.000797 | 0.001418 | 0.001359 | 0.000809 | 0.000449 | 0.002184 |
| k__Bacteria;p__Proteobacteria;c__Alphaproteobacteria;o__Sphingomonadales;f__Sphingomonadaceae;g__Sphingopyxis;s__alaskensis-bauzanensis-chilensis | 0.000238 | 0.000738 | 0 | 0 | 0 | 0 | 0 | 0 | 0 | 0 | 0 | 0 | 0 | 0 | 0 | 0 | 0 | 0 |
| k__Bacteria;p__Proteobacteria;c__Betaproteobacteria;o__Burkholderiales;f__Alcaligenaceae;g__Parasutterella;s__sp48235 | 0.00538 | 0.003855 | 0.001824 | 0.056625 | 0 | 0 | 0 | 0.001431 | 0 | 0.003608 | 0.002992 | 0.001553 | 0.007071 | 0.005674 | 0.003859 | 0.004817 | 0.001627 | 0.011408 |
| k__Bacteria;p__Proteobacteria;c__Betaproteobacteria;o__Burkholderiales;f__Alcaligenaceae;g__Sutterella;s__wadsworthensis | 0 | 0 | 0 | 0 | 0 | 0 | 0 | 0 | 0 | 0 | 0 | 0 | 0 | 0 | 0.00034 | 0 | 0 | 0 |
| k__Bacteria;p__Proteobacteria;c__Deltaproteobacteria;o__Desulfovibrionales;f__Desulfovibrionaceae;g__Bilophila;s__sp52475 | 0.000543 | 0.000705 | 0.002613 | 0 | 0.000469 | 0.000963 | 0.005063 | 0.000392 | 0.002546 | 0.000368 | 0.001601 | 0 | 0.00045 | 0.009487 | 0.002111 | 0.000908 | 0.00721 | 0.000936 |

| | | | | | | | | | | | | | | | | | | |
|---|---|---|---|---|---|---|---|---|---|---|---|---|---|---|---|---|---|---|
| k__Bacteria;p__Proteobacteria;c__Gammaproteobacteria;o__Enterobacteriales;f__Enterobacteriaceae;g__Enterobacter;s__NA | 0 | 0 | 0 | 0 | 0 | 0 | 0 | 0 | 0 | 0 | 0 | 0.004791 | 0 | 0 | 0 | 0 | 0 | 0 |
| k__Bacteria;p__Proteobacteria;c__Gammaproteobacteria;o__Enterobacteriales;f__Enterobacteriaceae;g__Enterobacter;s__cloacae-ludwigii | 0 | 0 | 0 | 0 | 0 | 0 | 0 | 0 | 0 | 0 | 0 | 0.002994 | 0 | 0 | 0 | 0 | 0 | 0 |
| k__Bacteria;p__Proteobacteria;c__Gammaproteobacteria;o__Enterobacteriales;f__Enterobacteriaceae;g__Escherichia-Shigella;s__coli | 0.000407 | 0.000525 | 0 | 0.017855 | 0.005707 | 0.004641 | 0 | 0.000804 | 0 | 0.000301 | 0 | 0.000243 | 0 | 0 | 0.007038 | 0.000296 | 0 | 0 |
| k__Bacteria;p__Proteobacteria;c__Gammaproteobacteria;o__Enterobacteriales;f__Enterobacteriaceae;g__Klebsiella;s__pneumoniae-quasipneumoniae | 0 | 0 | 0 | 0.000776 | 0 | 0 | 0 | 0 | 0 | 0 | 0 | 0 | 0 | 0 | 0 | 0 | 0 | 0 |
| k__Bacteria;p__Proteobacteria;c__Gammaproteobacteria;o__Pseudomonadales;f__Moraxellaceae;g__Acinetobacter;s__bouvetii | 0 | 0.000771 | 0 | 0 | 0 | 0 | 0 | 0 | 0 | 0 | 0 | 0 | 0 | 0 | 0 | 0 | 0 | 0 |
| k__Bacteria;p__Saccharibacteria;c__NA;o__NA;f__NA;g__*Saccharimonas;s__sp65946 | 0 | 0.000246 | 0.002123 | 0 | 0.002205 | 0 | 0 | 0 | 0 | 0.000184 | 0.000421 | 0 | 0.000674 | 0.000953 | 0 | 0 | 0.000309 | 0 |
| k__Bacteria;p__Tenericutes;c__Mollicutes;o__Anaeroplasmatales;f__Anaeroplasmataceae;g__Anaeroplasma;s__sp67615 | 0 | 0 | 0.021641 | 0.001482 | 0.01508 | 0.000206 | 0 | 0 | 0.006311 | 0.004611 | 0.011462 | 0.005053 | 0.002289 | 0.001442 | 0.002742 | 0.000869 | 0.004124 | 0.001053 |
| k__Bacteria;p__Tenericutes;c__Mollicutes;o__NA;f__NA;g__NA;s__sp67943 | 0 | 0 | 0 | 0 | 0 | 0 | 0 | 0 | 0 | 0 | 0 | 0 | 0 | 0 | 0.000291 | 0 | 0 | 0 |
| k__Bacteria;p__Verrucomicrobia;c__Verrucomicrobiae;o__Verrucomicrobiales;f__Verrucomicrobiaceae;g__Akkermansia;s__muciniphila | 0 | 0 | 0 | 0 | 0 | 0 | 0 | 0 | 0 | 0 | 0 | 0 | 0 | 0 | 0.00051 | 0 | 0 | 0 |
