## Supplementary material for "The gut microbiota of environmentally enriched mice regulates visual cortical plasticity": Suppl. Table 3

| | | | | | | |
|---|---|---|---|---|---|---|
| **Distance from soma (um)** | **Tukey's multiple comparisons test** | **Predicted (LS) mean diff.** | **95.00% CI of diff.** | **Significant?** | **Summary** | **Adjusted P Value** |
| 0 | Row 1 | | | | | |
| | STANDARD vs. EE | 7.105E-15 | -1.914 to 1.914 | No | ns | >0.9999 |
| | STANDARD vs. EEabx | 0 | -1.968 to 1.968 | No | ns | >0.9999 |
| | EE vs. EEabx | -7.105E-15 | -1.997 to 1.997 | No | ns | >0.9999 |
| 1 | Row 2 | | | | | |
| | STANDARD vs. EE | -0.04583 | -1.959 to 1.868 | No | ns | 0.9983 |
| | STANDARD vs. EEabx | 0.3912 | -1.576 to 2.359 | No | ns | 0.8872 |
| | EE vs. EEabx | 0.437 | -1.560 to 2.435 | No | ns | 0.865 |
| 2 | Row 3 | | | | | |
| | STANDARD vs. EE | -0.1521 | -2.066 to 1.762 | No | ns | 0.981 |
| | STANDARD vs. EEabx | 0.5775 | -1.390 to 2.545 | No | ns | 0.7705 |
| | EE vs. EEabx | 0.7296 | -1.268 to 2.727 | No | ns | 0.6679 |
| 3 | Row 4 | | | | | |
| | STANDARD vs. EE | -0.1687 | -2.082 to 1.745 | No | ns | 0.9767 |
| | STANDARD vs. EEabx | 0.1979 | -1.770 to 2.166 | No | ns | 0.9698 |
| | EE vs. EEabx | 0.3667 | -1.631 to 2.364 | No | ns | 0.903 |
| 4 | Row 5 | | | | | |
| | STANDARD vs. EE | 0.2104 | -1.703 to 2.124 | No | ns | 0.964 |
| | STANDARD vs. EEabx | 0.64 | -1.328 to 2.608 | No | ns | 0.726 |
| | EE vs. EEabx | 0.4296 | -1.568 to 2.427 | No | ns | 0.8693 |
| 5 | Row 6 | | | | | |
| | STANDARD vs. EE | -0.01667 | -1.930 to 1.897 | No | ns | 0.9998 |
| | STANDARD vs. EEabx | 1.009 | -0.9584 to 2.977 | No | ns | 0.4517 |
| | EE vs. EEabx | 1.026 | -0.9716 to 3.023 | No | ns | 0.4508 |
| 6 | Row 7 | | | | | |
| | STANDARD vs. EE | 0.6229 | -1.291 to 2.537 | No | ns | 0.7257 |
| | STANDARD vs. EEabx | 1.582 | -0.3855 to 3.550 | No | ns | 0.1431 |
| | EE vs. EEabx | 0.9593 | -1.038 to 2.957 | No | ns | 0.4981 |
| 7 | Row 8 | | | | | |
| | STANDARD vs. EE | 1.042 | -0.8719 to 2.955 | No | ns | 0.4087 |
| | STANDARD vs. EEabx | 2.301 | 0.3332 to 4.269 | Yes | * | 0.0169 |
| | EE vs. EEabx | 1.259 | -0.7382 to 3.257 | No | ns | 0.3016 |
| 8 | Row 9 | | | | | |
| | STANDARD vs. EE | 0.7396 | -1.174 to 2.653 | No | ns | 0.6365 |
| | STANDARD vs. EEabx | 2.48 | 0.5126 to 4.448 | Yes | ** | 0.0088 |
| | EE vs. EEabx | 1.741 | -0.2567 to 3.738 | No | ns | 0.1022 |
| 9 | Row 10 | | | | | |
| | STANDARD vs. EE | 0.8354 | -1.078 to 2.749 | No | ns | 0.562 |
| | STANDARD vs. EEabx | 1.728 | -0.2397 to 3.696 | No | ns | 0.0987 |
| | EE vs. EEabx | 0.8926 | -1.105 to 2.890 | No | ns | 0.5468 |
| 10 | Row 11 | | | | | |

| | | | | | | |
|---|---|---|---|---|---|---|
| | STANDARD vs. EE | 0.5271 | -1.387 to 2.441 | No | ns | 0.7948 |
| | STANDARD vs. EEabx | 2.205 | 0.2372 to 4.173 | Yes | * | 0.0235 |
| | EE vs. EEabx | 1.678 | -0.3197 to 3.675 | No | ns | 0.12 |
| 11 | Row 12 | | | | | |
| | STANDARD vs. EE | -0.7854 | -2.699 to 1.128 | No | ns | 0.6008 |
| | STANDARD vs. EEabx | 1.337 | -0.6309 to 3.304 | No | ns | 0.2488 |
| | EE vs. EEabx | 2.122 | 0.1247 to 4.120 | Yes | * | 0.0341 |
| 12 | Row 13 | | | | | |
| | STANDARD vs. EE | -1.075 | -2.989 to 0.8386 | No | ns | 0.3857 |
| | STANDARD vs. EEabx | 0.4213 | -1.546 to 2.389 | No | ns | 0.8704 |
| | EE vs. EEabx | 1.496 | -0.5012 to 3.494 | No | ns | 0.1847 |
| 13 | Row 14 | | | | | |
| | STANDARD vs. EE | -1.944 | -3.857 to -0.03016 | Yes | * | 0.0455 |
| | STANDARD vs. EEabx | 0.04514 | -1.923 to 2.013 | No | ns | 0.9984 |
| | EE vs. EEabx | 1.989 | -0.008593 to 3.986 | No | ns | 0.0513 |
| 14 | Row 15 | | | | | |
| | STANDARD vs. EE | -2.408 | -4.322 to -0.4947 | Yes | ** | 0.0089 |
| | STANDARD vs. EEabx | 0.4954 | -1.472 to 2.463 | No | ns | 0.8254 |
| | EE vs. EEabx | 2.904 | 0.9062 to 4.901 | Yes | ** | 0.0019 |
| 15 | Row 16 | | | | | |
| | STANDARD vs. EE | -3.754 | -5.668 to -1.841 | Yes | **** | <0.0001 |
| | STANDARD vs. EEabx | -0.4838 | -2.451 to 1.484 | No | ns | 0.8327 |
| | EE vs. EEabx | 3.27 | 1.273 to 5.268 | Yes | *** | 0.0004 |
| 16 | Row 17 | | | | | |
| | STANDARD vs. EE | -3.219 | -5.132 to -1.305 | Yes | *** | 0.0002 |
| | STANDARD vs. EEabx | 0.04051 | -1.927 to 2.008 | No | ns | 0.9987 |
| | EE vs. EEabx | 3.259 | 1.262 to 5.257 | Yes | *** | 0.0004 |
| 17 | Row 18 | | | | | |
| | STANDARD vs. EE | -3.577 | -5.491 to -1.663 | Yes | **** | <0.0001 |
| | STANDARD vs. EEabx | 0.7674 | -1.200 to 2.735 | No | ns | 0.6313 |
| | EE vs. EEabx | 4.344 | 2.347 to 6.342 | Yes | **** | <0.0001 |
| 18 | Row 19 | | | | | |
| | STANDARD vs. EE | -3.59 | -5.503 to -1.676 | Yes | **** | <0.0001 |
| | STANDARD vs. EEabx | 1.492 | -0.4758 to 3.460 | No | ns | 0.1773 |
| | EE vs. EEabx | 5.081 | 3.084 to 7.079 | Yes | **** | <0.0001 |
| 19 | Row 20 | | | | | |
| | STANDARD vs. EE | -4.065 | -5.978 to -2.151 | Yes | **** | <0.0001 |
| | STANDARD vs. EEabx | 0.9132 | -1.054 to 2.881 | No | ns | 0.5216 |
| | EE vs. EEabx | 4.978 | 2.980 to 6.975 | Yes | **** | <0.0001 |
| 20 | Row 21 | | | | | |
| | STANDARD vs. EE | -3.354 | -5.268 to -1.441 | Yes | *** | 0.0001 |
| | STANDARD vs. EEabx | 0.831 | -1.137 to 2.799 | No | ns | 0.5832 |
| | EE vs. EEabx | 4.185 | 2.188 to 6.183 | Yes | **** | <0.0001 |
| 21 | Row 22 | | | | | |

| | | | | | | |
|---|---|---|---|---|---|---|
| | STANDARD vs. EE | -4 | -5.914 to -2.086 | Yes | **** | <0.0001 |
| | STANDARD vs. EEabx | 0.4815 | -1.486 to 2.449 | No | ns | 0.8342 |
| | EE vs. EEabx | 4.481 | 2.484 to 6.479 | Yes | **** | <0.0001 |
| 22 | Row 23 | | | | | |
| | STANDARD vs. EE | -4.104 | -6.018 to -2.191 | Yes | **** | <0.0001 |
| | STANDARD vs. EEabx | 1.025 | -0.9422 to 2.993 | No | ns | 0.4403 |
| | EE vs. EEabx | 5.13 | 3.132 to 7.127 | Yes | **** | <0.0001 |
| 23 | Row 24 | | | | | |
| | STANDARD vs. EE | -5.563 | -7.476 to -3.649 | Yes | **** | <0.0001 |
| | STANDARD vs. EEabx | 0.2338 | -1.734 to 2.201 | No | ns | 0.9581 |
| | EE vs. EEabx | 5.796 | 3.799 to 7.794 | Yes | **** | <0.0001 |
| 24 | Row 25 | | | | | |
| | STANDARD vs. EE | -6.802 | -8.716 to -4.888 | Yes | **** | <0.0001 |
| | STANDARD vs. EEabx | 0.235 | -1.733 to 2.203 | No | ns | 0.9577 |
| | EE vs. EEabx | 7.037 | 5.040 to 9.035 | Yes | **** | <0.0001 |
| 25 | Row 26 | | | | | |
| | STANDARD vs. EE | -6.892 | -8.805 to -4.978 | Yes | **** | <0.0001 |
| | STANDARD vs. EEabx | 1.171 | -0.7964 to 3.139 | No | ns | 0.3433 |
| | EE vs. EEabx | 8.063 | 6.065 to 10.06 | Yes | **** | <0.0001 |
| 26 | Row 27 | | | | | |
| | STANDARD vs. EE | -7.006 | -8.920 to -5.093 | Yes | **** | <0.0001 |
| | STANDARD vs. EEabx | -0.4248 | -2.392 to 1.543 | No | ns | 0.8684 |
| | EE vs. EEabx | 6.581 | 4.584 to 8.579 | Yes | **** | <0.0001 |
| 27 | Row 28 | | | | | |
| | STANDARD vs. EE | -6.573 | -8.487 to -4.659 | Yes | **** | <0.0001 |
| | STANDARD vs. EEabx | -0.184 | -2.152 to 1.784 | No | ns | 0.9738 |
| | EE vs. EEabx | 6.389 | 4.391 to 8.386 | Yes | **** | <0.0001 |
| 28 | Row 29 | | | | | |
| | STANDARD vs. EE | -5.652 | -7.566 to -3.738 | Yes | **** | <0.0001 |
| | STANDARD vs. EEabx | 0.1146 | -1.853 to 2.082 | No | ns | 0.9898 |
| | EE vs. EEabx | 5.767 | 3.769 to 7.764 | Yes | **** | <0.0001 |
| 29 | Row 30 | | | | | |
| | STANDARD vs. EE | -5.192 | -7.105 to -3.278 | Yes | **** | <0.0001 |
| | STANDARD vs. EEabx | -0.01389 | -1.982 to 1.954 | No | ns | 0.9998 |
| | EE vs. EEabx | 5.178 | 3.180 to 7.175 | Yes | **** | <0.0001 |
| 30 | Row 31 | | | | | |
| | STANDARD vs. EE | -5.35 | -7.264 to -3.436 | Yes | **** | <0.0001 |
| | STANDARD vs. EEabx | -0.06481 | -2.033 to 1.903 | No | ns | 0.9967 |
| | EE vs. EEabx | 5.285 | 3.288 to 7.283 | Yes | **** | <0.0001 |
| 31 | Row 32 | | | | | |
| | STANDARD vs. EE | -4.348 | -6.262 to -2.434 | Yes | **** | <0.0001 |
| | STANDARD vs. EEabx | 0.2373 | -1.730 to 2.205 | No | ns | 0.9569 |
| | EE vs. EEabx | 4.585 | 2.588 to 6.583 | Yes | **** | <0.0001 |
| 32 | Row 33 | | | | | |

| | | | | | | |
|---|---|---|---|---|---|---|
| | STANDARD vs. EE | -4.692 | -6.605 to -2.778 | Yes | **** | <0.0001 |
| | STANDARD vs. EEabx | -0.06944 | -2.037 to 1.898 | No | ns | 0.9962 |
| | EE vs. EEabx | 4.622 | 2.625 to 6.620 | Yes | **** | <0.0001 |
| 33 | Row 34 | | | | | |
| | STANDARD vs. EE | -4.425 | -6.339 to -2.511 | Yes | **** | <0.0001 |
| | STANDARD vs. EEabx | 0.3935 | -1.574 to 2.361 | No | ns | 0.8859 |
| | EE vs. EEabx | 4.819 | 2.821 to 6.816 | Yes | **** | <0.0001 |
| 34 | Row 35 | | | | | |
| | STANDARD vs. EE | -4.127 | -6.041 to -2.213 | Yes | **** | <0.0001 |
| | STANDARD vs. EEabx | 0.6285 | -1.339 to 2.596 | No | ns | 0.7344 |
| | EE vs. EEabx | 4.756 | 2.758 to 6.753 | Yes | **** | <0.0001 |
| 35 | Row 36 | | | | | |
| | STANDARD vs. EE | -3.258 | -5.172 to -1.345 | Yes | *** | 0.0002 |
| | STANDARD vs. EEabx | 1.06 | -0.9075 to 3.028 | No | ns | 0.4162 |
| | EE vs. EEabx | 4.319 | 2.321 to 6.316 | Yes | **** | <0.0001 |
| 36 | Row 37 | | | | | |
| | STANDARD vs. EE | -3.075 | -4.989 to -1.161 | Yes | *** | 0.0005 |
| | STANDARD vs. EEabx | 0.7546 | -1.213 to 2.722 | No | ns | 0.6409 |
| | EE vs. EEabx | 3.83 | 1.832 to 5.827 | Yes | **** | <0.0001 |
| 37 | Row 38 | | | | | |
| | STANDARD vs. EE | -2.875 | -4.789 to -0.9614 | Yes | ** | 0.0013 |
| | STANDARD vs. EEabx | 0.5139 | -1.454 to 2.482 | No | ns | 0.8134 |
| | EE vs. EEabx | 3.389 | 1.391 to 5.386 | Yes | *** | 0.0002 |
| 38 | Row 39 | | | | | |
| | STANDARD vs. EE | -2.585 | -4.499 to -0.6718 | Yes | ** | 0.0044 |
| | STANDARD vs. EEabx | 0.4664 | -1.501 to 2.434 | No | ns | 0.8435 |
| | EE vs. EEabx | 3.052 | 1.054 to 5.049 | Yes | ** | 0.001 |
| 39 | Row 40 | | | | | |
| | STANDARD vs. EE | -2.556 | -4.470 to -0.6427 | Yes | ** | 0.005 |
| | STANDARD vs. EEabx | 0.2141 | -1.754 to 2.182 | No | ns | 0.9648 |
| | EE vs. EEabx | 2.77 | 0.7729 to 4.768 | Yes | ** | 0.0033 |
| 40 | Row 41 | | | | | |
| | STANDARD vs. EE | -2.388 | -4.301 to -0.4739 | Yes | ** | 0.0097 |
| | STANDARD vs. EEabx | 0.09028 | -1.877 to 2.058 | No | ns | 0.9936 |
| | EE vs. EEabx | 2.478 | 0.4803 to 4.475 | Yes | * | 0.0102 |
| 41 | Row 42 | | | | | |
| | STANDARD vs. EE | -2.294 | -4.207 to -0.3802 | Yes | * | 0.0138 |
| | STANDARD vs. EEabx | 0.0544 | -1.913 to 2.022 | No | ns | 0.9977 |
| | EE vs. EEabx | 2.348 | 0.3507 to 4.346 | Yes | * | 0.0162 |
| 42 | Row 43 | | | | | |
| | STANDARD vs. EE | -1.929 | -3.843 to -0.01558 | Yes | * | 0.0476 |
| | STANDARD vs. EEabx | 0.3079 | -1.660 to 2.276 | No | ns | 0.9285 |
| | EE vs. EEabx | 2.237 | 0.2396 to 4.235 | Yes | * | 0.0236 |
| 43 | Row 44 | | | | | |

| | | | | | | |
|---|---|---|---|---|---|---|
| | STANDARD vs. EE | -1.548 | -3.462 to 0.3657 | No | ns | 0.1398 |
| | STANDARD vs. EEabx | 0.2373 | -1.730 to 2.205 | No | ns | 0.9569 |
| | EE vs. EEabx | 1.785 | -0.2123 to 3.783 | No | ns | 0.0909 |
| 44 | Row 45 | | | | | |
| | STANDARD vs. EE | -1.204 | -3.118 to 0.7094 | No | ns | 0.3029 |
| | STANDARD vs. EEabx | 0.08102 | -1.887 to 2.049 | No | ns | 0.9949 |
| | EE vs. EEabx | 1.285 | -0.7123 to 3.283 | No | ns | 0.287 |
| 45 | Row 46 | | | | | |
| | STANDARD vs. EE | -1.158 | -3.072 to 0.7553 | No | ns | 0.3311 |
| | STANDARD vs. EEabx | -0.1065 | -2.074 to 1.861 | No | ns | 0.9912 |
| | EE vs. EEabx | 1.052 | -0.9456 to 3.049 | No | ns | 0.4328 |
| 46 | Row 47 | | | | | |
| | STANDARD vs. EE | -1.315 | -3.228 to 0.5990 | No | ns | 0.2412 |
| | STANDARD vs. EEabx | -0.1516 | -2.119 to 1.816 | No | ns | 0.9822 |
| | EE vs. EEabx | 1.163 | -0.8345 to 3.160 | No | ns | 0.3595 |
| 47 | Row 48 | | | | | |
| | STANDARD vs. EE | -1.106 | -3.020 to 0.8073 | No | ns | 0.3647 |
| | STANDARD vs. EEabx | -0.2766 | -2.244 to 1.691 | No | ns | 0.9419 |
| | EE vs. EEabx | 0.8296 | -1.168 to 2.827 | No | ns | 0.5936 |
| 48 | Row 49 | | | | | |
| | STANDARD vs. EE | -1.006 | -2.920 to 0.9073 | No | ns | 0.4338 |
| | STANDARD vs. EEabx | -0.1285 | -2.096 to 1.839 | No | ns | 0.9872 |
| | EE vs. EEabx | 0.8778 | -1.120 to 2.875 | No | ns | 0.5578 |
| 49 | Row 50 | | | | | |
| | STANDARD vs. EE | -0.7354 | -2.649 to 1.178 | No | ns | 0.6397 |
| | STANDARD vs. EEabx | -0.07986 | -2.048 to 1.888 | No | ns | 0.995 |
| | EE vs. EEabx | 0.6556 | -1.342 to 2.653 | No | ns | 0.7218 |
| 50 | Row 51 | | | | | |
| | STANDARD vs. EE | -0.6021 | -2.516 to 1.312 | No | ns | 0.7411 |
| | STANDARD vs. EEabx | -0.07986 | -2.048 to 1.888 | No | ns | 0.995 |
| | EE vs. EEabx | 0.5222 | -1.475 to 2.520 | No | ns | 0.8131 |
| 51 | Row 52 | | | | | |
| | STANDARD vs. EE | -0.4021 | -2.316 to 1.512 | No | ns | 0.8748 |
| | STANDARD vs. EEabx | -0.1169 | -2.085 to 1.851 | No | ns | 0.9894 |
| | EE vs. EEabx | 0.2852 | -1.712 to 2.283 | No | ns | 0.9401 |
| 52 | Row 53 | | | | | |
| | STANDARD vs. EE | -0.2021 | -2.116 to 1.712 | No | ns | 0.9668 |
| | STANDARD vs. EEabx | -0.1169 | -2.085 to 1.851 | No | ns | 0.9894 |
| | EE vs. EEabx | 0.08519 | -1.912 to 2.083 | No | ns | 0.9945 |
| 53 | Row 54 | | | | | |
| | STANDARD vs. EE | -0.1 | -2.014 to 1.814 | No | ns | 0.9918 |
| | STANDARD vs. EEabx | -0.1111 | -2.079 to 1.857 | No | ns | 0.9904 |
| | EE vs. EEabx | -0.01111 | -2.009 to 1.986 | No | ns | >0.9999 |
| 54 | Row 55 | | | | | |

| | | | | | | |
|---|---|---|---|---|---|---|
| | STANDARD vs. EE | -0.06667 | -1.980 to 1.847 | No | ns | 0.9963 |
| | STANDARD vs. EEabx | -0.1111 | -2.079 to 1.857 | No | ns | 0.9904 |
| | EE vs. EEabx | -0.04444 | -2.042 to 1.953 | No | ns | 0.9985 |
| 55 | Row 56 | | | | | |
| | STANDARD vs. EE | -0.06667 | -1.980 to 1.847 | No | ns | 0.9963 |
| | STANDARD vs. EEabx | -0.1111 | -2.079 to 1.857 | No | ns | 0.9904 |
| | EE vs. EEabx | -0.04444 | -2.042 to 1.953 | No | ns | 0.9985 |
| 56 | Row 57 | | | | | |
| | STANDARD vs. EE | -0.06667 | -1.980 to 1.847 | No | ns | 0.9963 |
| | STANDARD vs. EEabx | -0.1111 | -2.079 to 1.857 | No | ns | 0.9904 |
| | EE vs. EEabx | -0.04444 | -2.042 to 1.953 | No | ns | 0.9985 |
| 57 | Row 58 | | | | | |
| | STANDARD vs. EE | -0.03333 | -1.947 to 1.880 | No | ns | 0.9991 |
| | STANDARD vs. EEabx | -0.1111 | -2.079 to 1.857 | No | ns | 0.9904 |
| | EE vs. EEabx | -0.07778 | -2.075 to 1.920 | No | ns | 0.9954 |
| 58 | Row 59 | | | | | |
| | STANDARD vs. EE | 3.553E-15 | -1.914 to 1.914 | No | ns | >0.9999 |
| | STANDARD vs. EEabx | -0.1111 | -2.079 to 1.857 | No | ns | 0.9904 |
| | EE vs. EEabx | -0.1111 | -2.109 to 1.886 | No | ns | 0.9907 |
| 59 | Row 60 | | | | | |
| | STANDARD vs. EE | 3.553E-15 | -1.914 to 1.914 | No | ns | >0.9999 |
| | STANDARD vs. EEabx | -0.07407 | -2.042 to 1.894 | No | ns | 0.9957 |
| | EE vs. EEabx | -0.07407 | -2.072 to 1.923 | No | ns | 0.9958 |
| 60 | Row 61 | | | | | |
| | STANDARD vs. EE | 1.776E-15 | -1.914 to 1.914 | No | ns | >0.9999 |
| | STANDARD vs. EEabx | -0.07407 | -2.042 to 1.894 | No | ns | 0.9957 |
| | EE vs. EEabx | -0.07407 | -2.072 to 1.923 | No | ns | 0.9958 |
| 61 | Row 62 | | | | | |
| | STANDARD vs. EE | 5.329E-15 | -1.914 to 1.914 | No | ns | >0.9999 |
| | STANDARD vs. EEabx | -0.03704 | -2.005 to 1.931 | No | ns | 0.9989 |
| | EE vs. EEabx | -0.03704 | -2.035 to 1.960 | No | ns | 0.999 |
| 62 | Row 63 | | | | | |
| | STANDARD vs. EE | 0 | -1.914 to 1.914 | No | ns | >0.9999 |
| | STANDARD vs. EEabx | 0 | -1.968 to 1.968 | No | ns | >0.9999 |
| | EE vs. EEabx | 0 | -1.997 to 1.997 | No | ns | >0.9999 |
