## Supplementary material for "The gut microbiota of environmentally enriched mice regulates visual cortical plasticity": Suppl. Table 4

| STANDARD - SCFA | | | | | | |
|---|---|---|---|---|---|---|
| **Sidak's multiple comparisons test** | **Distance from soma (um)** | **Predicted (LS) mean diff.** | **95.00% CI of diff.** | **Significant?** | **Summary** | **Adjusted P Value** |
| Row 1 | 0 | 0 | -2.736 to 2.736 | No | ns | >0.9999 |
| Row 2 | 1 | -0.0725 | -2.808 to 2.663 | No | ns | >0.9999 |
| Row 3 | 2 | -0.3588 | -3.095 to 2.377 | No | ns | >0.9999 |
| Row 4 | 3 | -0.9488 | -3.685 to 1.787 | No | ns | >0.9999 |
| Row 5 | 4 | -0.7363 | -3.472 to 2.000 | No | ns | >0.9999 |
| Row 6 | 5 | -1.53 | -4.266 to 1.206 | No | ns | 0.9811 |
| Row 7 | 6 | -1.944 | -4.680 to 0.7922 | No | ns | 0.6677 |
| Row 8 | 7 | -2.105 | -4.841 to 0.6310 | No | ns | 0.4682 |
| Row 9 | 8 | -2.074 | -4.810 to 0.6622 | No | ns | 0.506 |
| Row 10 | 9 | -3.051 | -5.787 to -0.3153 | Yes | * | 0.0118 |
| Row 11 | 10 | -2.786 | -5.522 to -0.05028 | Yes | * | 0.0402 |
| Row 12 | 11 | -4.099 | -6.835 to -1.363 | Yes | **** | <0.0001 |
| Row 13 | 12 | -4.355 | -7.091 to -1.619 | Yes | **** | <0.0001 |
| Row 14 | 13 | -3.604 | -6.340 to -0.8678 | Yes | *** | 0.0007 |
| Row 15 | 14 | -2.795 | -5.531 to -0.05903 | Yes | * | 0.0387 |
| Row 16 | 15 | -2.708 | -5.443 to 0.02847 | No | ns | 0.0565 |
| Row 17 | 16 | -1.099 | -3.835 to 1.637 | No | ns | >0.9999 |
| Row 18 | 17 | -0.6638 | -3.400 to 2.072 | No | ns | >0.9999 |
| Row 19 | 18 | -1.096 | -3.832 to 1.640 | No | ns | >0.9999 |
| Row 20 | 19 | -1.131 | -3.867 to 1.605 | No | ns | >0.9999 |
| Row 21 | 20 | -1.288 | -4.023 to 1.448 | No | ns | 0.9995 |
| Row 22 | 21 | -1 | -3.736 to 1.736 | No | ns | >0.9999 |
| Row 23 | 22 | -1.658 | -4.393 to 1.078 | No | ns | 0.9348 |
| Row 24 | 23 | -1.543 | -4.278 to 1.193 | No | ns | 0.9782 |
| Row 25 | 24 | -1.349 | -4.085 to 1.387 | No | ns | 0.9986 |
| Row 26 | 25 | -1.645 | -4.381 to 1.091 | No | ns | 0.9413 |
| Row 27 | 26 | -2.386 | -5.122 to 0.3497 | No | ns | 0.1978 |
| Row 28 | 27 | -1.966 | -4.702 to 0.7697 | No | ns | 0.6399 |
| Row 29 | 28 | -1.779 | -4.515 to 0.9572 | No | ns | 0.8477 |
| Row 30 | 29 | -1.285 | -4.021 to 1.451 | No | ns | 0.9996 |
| Row 31 | 30 | -0.85 | -3.586 to 1.886 | No | ns | >0.9999 |
| Row 32 | 31 | -0.9213 | -3.657 to 1.815 | No | ns | >0.9999 |
| Row 33 | 32 | -0.945 | -3.681 to 1.791 | No | ns | >0.9999 |
| Row 34 | 33 | -0.845 | -3.581 to 1.891 | No | ns | >0.9999 |
| Row 35 | 34 | -0.8738 | -3.610 to 1.862 | No | ns | >0.9999 |
| Row 36 | 35 | -0.885 | -3.621 to 1.851 | No | ns | >0.9999 |
| Row 37 | 36 | -1.195 | -3.931 to 1.541 | No | ns | >0.9999 |
| Row 38 | 37 | -1.055 | -3.791 to 1.681 | No | ns | >0.9999 |
| Row 39 | 38 | -0.9588 | -3.695 to 1.777 | No | ns | >0.9999 |
| Row 40 | 39 | -1.036 | -3.772 to 1.700 | No | ns | >0.9999 |
| Row 41 | 40 | -0.9675 | -3.703 to 1.768 | No | ns | >0.9999 |
| Row 42 | 41 | -0.8538 | -3.590 to 1.882 | No | ns | >0.9999 |
| Row 43 | 42 | -0.3425 | -3.078 to 2.393 | No | ns | >0.9999 |

| | | | | | | |
|---|---|---|---|---|---|---|
| Row 44 | 43 | -0.4813 | -3.217 to 2.255 | No | ns | >0.9999 |
| Row 45 | 44 | -0.4375 | -3.173 to 2.298 | No | ns | >0.9999 |
| Row 46 | 45 | -0.385 | -3.121 to 2.351 | No | ns | >0.9999 |
| Row 47 | 46 | -0.4213 | -3.157 to 2.315 | No | ns | >0.9999 |
| Row 48 | 47 | -0.5463 | -3.282 to 2.190 | No | ns | >0.9999 |
| Row 49 | 48 | -0.4263 | -3.162 to 2.310 | No | ns | >0.9999 |
| Row 50 | 49 | -0.4487 | -3.185 to 2.287 | No | ns | >0.9999 |
| Row 51 | 50 | -0.3287 | -3.065 to 2.407 | No | ns | >0.9999 |
| Row 52 | 51 | -0.2087 | -2.945 to 2.527 | No | ns | >0.9999 |
| Row 53 | 52 | -0.08875 | -2.825 to 2.647 | No | ns | >0.9999 |
| Row 54 | 53 | -0.08 | -2.816 to 2.656 | No | ns | >0.9999 |
| Row 55 | 54 | -0.04 | -2.776 to 2.696 | No | ns | >0.9999 |
| Row 56 | 55 | -0.04 | -2.776 to 2.696 | No | ns | >0.9999 |
| Row 57 | 56 | -0.04 | -2.776 to 2.696 | No | ns | >0.9999 |
| Row 58 | 57 | 3.553E-15 | -2.736 to 2.736 | No | ns | >0.9999 |
| Row 59 | 58 | 0 | -2.736 to 2.736 | No | ns | >0.9999 |
| Row 60 | 59 | 3.553E-15 | -2.736 to 2.736 | No | ns | >0.9999 |
| Row 61 | 60 | 0 | -2.736 to 2.736 | No | ns | >0.9999 |
| Row 62 | 61 | 0 | -2.736 to 2.736 | No | ns | >0.9999 |
| Row 63 | 62 | 0 | -2.736 to 2.736 | No | ns | >0.9999 |
