## Supplementary material for "The gut microbiota of environmentally enriched mice regulates visual cortical plasticity": Suppl. Table 5

| SampleID | EE3 | EE5 | EE1 | EE4 | EE2 | EE6 | EE7 | EE8 | postFT.1 | postFT.8 | postFT.10 | postFT.11 | postFT.18 | postFT.20 | postFT.22 | postFT.23 |
|---|---|---|---|---|---|---|---|---|---|---|---|---|---|---|---|---|
| None;Other;Other;Other;Other;Other;Other | 0 | 0.000335 | 0 | 0.003322 | 0 | 0.000297 | 0.000728 | 0 | 0 | 0 | 0 | 0 | 0 | 0 | 0.000481 | 0 |
| k__Bacteria;p__Actinobacteria;c__Actinobacteria;o__Bifidobacteriales;f__Bifidobacteriaceae;g__Bifidobacterium;s__pseudolongum | 0 | 0.000512 | 0 | 0 | 0 | 0 | 0 | 0 | 0.00162 | 0.00139 | 0 | 0.000771 | 0.000805 | 0 | 0.002259 | 0 |
| k__Bacteria;p__Actinobacteria;c__Actinobacteria;o__Propionibacteriales;f__Propionibacteriaceae;g__Propionibacterium;s__acnes | 0.000564 | 0 | 0 | 0 | 0 | 0 | 0 | 0 | 0 | 0 | 0 | 0 | 0 | 0 | 0 | 0 |
| k__Bacteria;p__Actinobacteria;c__Coriobacteriia;o__Coriobacteriales;f__Coriobacteriaceae;g__Enterorhabdus;s__mucosicola | 0 | 0 | 0 | 0 | 0 | 0 | 0 | 0 | 0 | 0 | 0 | 0 | 0.001568 | 0 | 0 | 0 |
| k__Bacteria;p__Actinobacteria;c__Coriobacteriia;o__Coriobacteriales;f__Coriobacteriaceae;g__Olsenella;s__profusa | 0 | 0.000846 | 0.001593 | 0 | 0 | 0 | 0 | 0 | 0 | 0 | 0 | 0 | 0 | 0 | 0 | 0 |
| k__Bacteria;p__Bacteroidetes;c__Bacteroidia;o__Bacteroidales;f__Bacteroidaceae;g__Bacteroides;s__acidifaciens | 0.007358 | 0.009328 | 0.022336 | 0.013186 | 0.022155 | 0.004774 | 0.009388 | 0.010622 | 0.004703 | 0.005375 | 0.005956 | 0.002191 | 0.004344 | 0.005444 | 0.002163 | 0.00909 |
| k__Bacteria;p__Bacteroidetes;c__Bacteroidia;o__Bacteroidales;f__Bacteroidaceae;g__Bacteroides;s__intestinalis | 0 | 0 | 0 | 0 | 0 | 0 | 0 | 0 | 0 | 0 | 0 | 0 | 0 | 0 | 0 | 0.000168 |
| k__Bacteria;p__Bacteroidetes;c__Bacteroidia;o__Bacteroidales;f__Bacteroidaceae;g__Bacteroides;s__oleiciplenus-rodentium | 0.016331 | 0.023496 | 0.034287 | 0.040389 | 0.046017 | 0.02907 | 0.025253 | 0.041495 | 0.006075 | 0.01374 | 0.010999 | 0.004329 | 0.000996 | 0.009646 | 0.002716 | 0.007031 |
| k__Bacteria;p__Bacteroidetes;c__Bacteroidia;o__Bacteroidales;f__Bacteroidaceae;g__Bacteroides;s__stercorirosoris | 0 | 0 | 0.000495 | 0 | 0.001597 | 0 | 0.001064 | 0.001092 | 0 | 0 | 0 | 0 | 0 | 0 | 0.000361 | 0 |
| k__Bacteria;p__Bacteroidetes;c__Bacteroidia;o__Bacteroidales;f__NA;g__NA;s__sp12473-sp12526 | 0.000718 | 0 | 0 | 0 | 0.006343 | 0 | 0.003098 | 0 | 0.003015 | 0.003568 | 0.006614 | 0.003768 | 0.002416 | 0.004625 | 0.001034 | 0.006574 |
| k__Bacteria;p__Bacteroidetes;c__Bacteroidia;o__Bacteroidales;f__NA;g__NA;s__sp12475-sp12557 | 0.020946 | 0.021233 | 0.043078 | 0.032622 | 0.02628 | 0.010737 | 0.008194 | 0.00438 | 0.02806 | 0.018837 | 0.026534 | 0.025168 | 0.016358 | 0.041675 | 0.02144 | 0.057362 |
| k__Bacteria;p__Bacteroidetes;c__Bacteroidia;o__Bacteroidales;f__NA;g__NA;s__sp12494 | 0.010691 | 0.014877 | 0.028352 | 0.017048 | 0.007274 | 0.008509 | 0.010676 | 0.009251 | 0.012556 | 0.020968 | 0.019514 | 0.01339 | 0.011718 | 0.012104 | 0.012691 | 0.019172 |
| k__Bacteria;p__Bacteroidetes;c__Bacteroidia;o__Bacteroidales;f__NA;g__NA;s__sp12497 | 0 | 0.006199 | 0 | 0.03094 | 0.001419 | 0.006281 | 0.005226 | 0.000826 | 0.011858 | 0.0095 | 0.019362 | 0.017719 | 0.001843 | 0.018129 | 0.001514 | 0.042171 |
| k__Bacteria;p__Bacteroidetes;c__Bacteroidia;o__Bacteroidales;f__NA;g__NA;s__sp12499 | 0.001743 | 0.003444 | 0.007033 | 0.013913 | 0.008072 | 0.002419 | 0.003733 | 0.003317 | 0.003488 | 0.006951 | 0.007476 | 0.003593 | 0.004111 | 0.009276 | 0.009254 | 0.0124 |
| k__Bacteria;p__Bacteroidetes;c__Bacteroidia;o__Bacteroidales;f__NA;g__NA;s__sp12502 | 0.003307 | 0.002893 | 0.004011 | 0.003821 | 0 | 0.002483 | 0 | 0.003261 | 0 | 0 | 0 | 0 | 0.002458 | 0 | 0.001827 | 0 |
| k__Bacteria;p__Bacteroidetes;c__Bacteroidia;o__Bacteroidales;f__NA;g__NA;s__sp12503 | 0 | 0.003168 | 0 | 0.011504 | 0 | 0.002398 | 0.00181 | 0.000392 | 0.004748 | 0.002757 | 0.005956 | 0.006064 | 0.000445 | 0.005629 | 0.000505 | 0.013391 |
| k__Bacteria;p__Bacteroidetes;c__Bacteroidia;o__Bacteroidales;f__NA;g__NA;s__sp12505 | 0.041123 | 0.039259 | 0.11319 | 0.163984 | 0.152732 | 0.01479 | 0.127328 | 0.066028 | 0.012286 | 0.028267 | 0.04179 | 0.014109 | 0.009895 | 0.021723 | 0.026199 | 0.002623 |
| k__Bacteria;p__Bacteroidetes;c__Bacteroidia;o__Bacteroidales;f__NA;g__NA;s__sp12520 | 0.004076 | 0.006258 | 0.00555 | 0.012252 | 0.004812 | 0.003544 | 0.003584 | 0.001022 | 0.019351 | 0.021409 | 0.016346 | 0.013022 | 0.027249 | 0.022067 | 0.04074 | 0.023747 |
| k__Bacteria;p__Bacteroidetes;c__Bacteroidia;o__Bacteroidales;f__NA;g__NA;s__sp12532 | 0.036816 | 0.002716 | 0.007116 | 0.050003 | 0.020691 | 0.004583 | 0.045989 | 0.003625 | 0.018699 | 0.038068 | 0.022479 | 0.008693 | 0.022355 | 0.025053 | 0.026944 | 0.043757 |
| k__Bacteria;p__Bacteroidetes;c__Bacteroidia;o__Bacteroidales;f__NA;g__NA;s__sp12535 | 0.005179 | 0.003955 | 0.008242 | 0.012418 | 0.004191 | 0.001761 | 0.006141 | 0.00063 | 0.001125 | 0.001552 | 0.001039 | 0.000438 | 0.001356 | 0.002061 | 0.00274 | 0.002425 |
| k__Bacteria;p__Bacteroidetes;c__Bacteroidia;o__Bacteroidales;f__NA;g__NA;s__sp12536-sp12753 | 0.011152 | 0.005195 | 0.005989 | 0.006105 | 0.01253 | 0.006769 | 0.015361 | 0.004254 | 0.00423 | 0.003846 | 0.006133 | 0.004154 | 0.001949 | 0.01324 | 0.00709 | 0.006467 |
| k__Bacteria;p__Bacteroidetes;c__Bacteroidia;o__Bacteroidales;f__NA;g__NA;s__sp12539 | 0.003256 | 0.00488 | 0.005962 | 0.012812 | 0.012441 | 0.002398 | 0.008847 | 0.001008 | 0.006773 | 0.009314 | 0.009808 | 0.007326 | 0.008115 | 0.010122 | 0.017738 | 0.012567 |
| k__Bacteria;p__Bacteroidetes;c__Bacteroidia;o__Bacteroidales;f__NA;g__NA;s__sp12549 | 0.007871 | 0.000531 | 0.001209 | 0.007206 | 0.00275 | 0.000382 | 0.00293 | 0.001903 | 0.001373 | 0 | 0.001318 | 0.000859 | 0.001865 | 0.001903 | 0.001346 | 0.002151 |
| k__Bacteria;p__Bacteroidetes;c__Bacteroidia;o__Bacteroidales;f__NA;g__NA;s__sp12554 | 0 | 0 | 0 | 0.005025 | 0.001441 | 0 | 0.001829 | 0 | 0.003173 | 0.005978 | 0.005347 | 0.002839 | 0.003645 | 0.008906 | 0.006514 | 0.007565 |
| k__Bacteria;p__Bacteroidetes;c__Bacteroidia;o__Bacteroidales;f__NA;g__NA;s__sp12565 | 0.021433 | 0.015192 | 0.039864 | 0.004921 | 0.017187 | 0.007066 | 0.006383 | 0.025233 | 0.004748 | 0.004657 | 0.019413 | 0.006204 | 0.000954 | 0.014693 | 0.005047 | 0.011866 |
| k__Bacteria;p__Bacteroidetes;c__Bacteroidia;o__Bacteroidales;f__NA;g__NA;s__sp12572-sp12578-sp12693 | 0 | 0 | 0 | 0.014743 | 0 | 0 | 0 | 0 | 0 | 0 | 0 | 0 | 0 | 0 | 0 | 0.065293 |
| k__Bacteria;p__Bacteroidetes;c__Bacteroidia;o__Bacteroidales;f__NA;g__NA;s__sp12589 | 0.022818 | 0.007655 | 0.01022 | 0.027618 | 0.025925 | 0.00783 | 0.025664 | 0.013645 | 0.006751 | 0.012303 | 0.00996 | 0.007834 | 0.000911 | 0.013742 | 0.005865 | 0.015237 |
| k__Bacteria;p__Bacteroidetes;c__Bacteroidia;o__Bacteroidales;f__NA;g__NA;s__sp12590 | 0.001513 | 0.001043 | 0.002363 | 0.002824 | 0.001774 | 0.000828 | 0.001792 | 0.00126 | 0.001148 | 0.002108 | 0.002458 | 0.000929 | 0.000742 | 0.000687 | 0.002428 | 0.004042 |
| k__Bacteria;p__Bacteroidetes;c__Bacteroidia;o__Bacteroidales;f__NA;g__NA;s__sp12595 | 0.003153 | 0.006494 | 0.019094 | 0.006915 | 0.005079 | 0.001422 | 0.003378 | 0.005276 | 0.003015 | 0.004333 | 0.004106 | 0.002611 | 0.004132 | 0.008298 | 0.004158 | 0.008175 |
| k__Bacteria;p__Bacteroidetes;c__Bacteroidia;o__Bacteroidales;f__NA;g__NA;s__sp12595-sp12654 | 0.002051 | 0.004585 | 0.005907 | 0.006396 | 0.001841 | 0.00174 | 0.0014 | 0.000826 | 0.00549 | 0.007113 | 0.005221 | 0.005258 | 0.008264 | 0.001506 | 0.017017 | 0.002974 |
| k__Bacteria;p__Bacteroidetes;c__Bacteroidia;o__Bacteroidales;f__NA;g__NA;s__sp12597 | 0.003999 | 0.001968 | 0.00206 | 0.018066 | 0.005677 | 0.002101 | 0.005394 | 0 | 0.00639 | 0.00702 | 0.003573 | 0.007396 | 0.000445 | 0.005021 | 0.035044 | 0.009273 |
| k__Bacteria;p__Bacteroidetes;c__Bacteroidia;o__Bacteroidales;f__NA;g__NA;s__sp12629 | 0.000333 | 0 | 0 | 0.000644 | 0 | 0 | 0 | 0 | 0 | 0.000394 | 0.000279 | 0 | 0.00072 | 0 | 0.002548 | 0.001235 |
| k__Bacteria;p__Bacteroidetes;c__Bacteroidia;o__Bacteroidales;f__NA;g__NA;s__sp12637 | 0.000769 | 0.000866 | 0.003901 | 0.001786 | 0.000776 | 0.000318 | 0 | 0.000252 | 0.004928 | 0.003499 | 0.007552 | 0.001052 | 0.00642 | 0.009989 | 0.008821 | 0.00816 |
| k__Bacteria;p__Bacteroidetes;c__Bacteroidia;o__Bacteroidales;f__NA;g__NA;s__sp12641 | 0.012357 | 0.00553 | 0.006594 | 0.005586 | 0.006343 | 0.007151 | 0.007298 | 0.004282 | 0.004275 | 0.003638 | 0.007349 | 0.004592 | 0.002246 | 0.013134 | 0.007235 | 0.006726 |
| k__Bacteria;p__Bacteroidetes;c__Bacteroidia;o__Bacteroidales;f__NA;g__NA;s__sp12647 | 0 | 0 | 0 | 0.007164 | 0 | 0 | 0 | 0 | 0.020207 | 0.034059 | 0.020654 | 0.012742 | 0.010425 | 0.002801 | 0.027377 | 0.016716 |
| k__Bacteria;p__Bacteroidetes;c__Bacteroidia;o__Bacteroidales;f__NA;g__NA;s__sp12654 | 0.002846 | 0.00366 | 0.009121 | 0.004361 | 0.002462 | 0.001273 | 0.001885 | 0.002519 | 0.004478 | 0.005538 | 0.0037 | 0.003628 | 0.00089 | 0.00111 | 0.007643 | 0.003111 |
| k__Bacteria;p__Bacteroidetes;c__Bacteroidia;o__Bacteroidales;f__NA;g__NA;s__sp12656 | 0.016665 | 0.014306 | 0.042556 | 0.013207 | 0.03331 | 0.019267 | 0.017115 | 0.013099 | 0.003983 | 0.015246 | 0.02106 | 0.013408 | 0.004132 | 0.029863 | 0.001394 | 0.032075 |
| k__Bacteria;p__Bacteroidetes;c__Bacteroidia;o__Bacteroidales;f__NA;g__NA;s__sp12666 | 0.005435 | 0.002322 | 0.005055 | 0.009074 | 0.007429 | 0.001464 | 0.003098 | 0.002743 | 0.002993 | 0.003892 | 0.00626 | 0.002822 | 0.006314 | 0.004017 | 0.002812 | 0.005887 |
| k__Bacteria;p__Bacteroidetes;c__Bacteroidia;o__Bacteroidales;f__NA;g__NA;s__sp12757 | 0.00441 | 0.002617 | 0.005714 | 0.01275 | 0.004746 | 0.002546 | 0.011703 | 0.002785 | 0.004028 | 0.014388 | 0.006716 | 0.006362 | 0.011548 | 0.009223 | 0.006033 | 0.019568 |
| k__Bacteria;p__Bacteroidetes;c__Bacteroidia;o__Bacteroidales;f__NA;g__NA;s__sp12777 | 0.003615 | 0.005294 | 0.009231 | 0.022032 | 0.009536 | 0.005135 | 0.008567 | 0.007012 | 0.005333 | 0.013693 | 0.013939 | 0.007221 | 0.007183 | 0.017891 | 0.014181 | 0.021825 |
| k__Bacteria;p__Bacteroidetes;c__Bacteroidia;o__Bacteroidales;f__NA;g__NA;s__sp12778 | 0.003179 | 0.021135 | 0.046677 | 0.003177 | 0.002484 | 0.008254 | 0 | 0.011518 | 0.014221 | 0.013647 | 0.016752 | 0.009289 | 0.017439 | 0.023705 | 0.018507 | 0.026492 |
| k__Bacteria;p__Bacteroidetes;c__Bacteroidia;o__Bacteroidales;f__NA;g__NA;s__sp12778-sp12792 | 0 | 0 | 0 | 0 | 0 | 0 | 0 | 0 | 0 | 0 | 0 | 0 | 0.004979 | 0 | 0 | 0 |
| k__Bacteria;p__Bacteroidetes;c__Bacteroidia;o__Bacteroidales;f__NA;g__NA;s__sp12785 | 0.005256 | 0.008285 | 0.014011 | 0.005773 | 0.011909 | 0.005793 | 0.005487 | 0.006228 | 0.004388 | 0.005792 | 0.006336 | 0.003873 | 0.003136 | 0.009355 | 0.007042 | 0.009959 |
| k__Bacteria;p__Bacteroidetes;c__Bacteroidia;o__Bacteroidales;f__NA;g__NA;s__sp12790 | 0.004076 | 0.002106 | 0.002473 | 0.01003 | 0.001397 | 0.00157 | 0.000821 | 0.001358 | 0.003285 | 0.004055 | 0.004714 | 0.002559 | 0.004789 | 0.007003 | 0.00298 | 0.00453 |
| k__Bacteria;p__Bacteroidetes;c__Bacteroidia;o__Bacteroidales;f__NA;g__NA;s__sp12792 | 0.035688 | 0.061475 | 0.145718 | 0.009137 | 0.008095 | 0.025357 | 0.017601 | 0.041033 | 0.052069 | 0.050116 | 0.062394 | 0.036052 | 0.05524 | 0.087711 | 0.072443 | 0.098908 |
| k__Bacteria;p__Bacteroidetes;c__Bacteroidia;o__Bacteroidales;f__NA;g__NA;s__sp12802 | 0.007435 | 0.003778 | 0.006017 | 0.030193 | 0.004613 | 0.002355 | 0.002202 | 0.001469 | 0.004703 | 0.00336 | 0.004511 | 0.00163 | 0.003157 | 0.01028 | 0.011249 | 0.013086 |
| k__Bacteria;p__Bacteroidetes;c__Bacteroidia;o__Bacteroidales;f__NA;g__NA;s__sp12804 | 0 | 0 | 0 | 0.001391 | 0.005034 | 0 | 0.004591 | 0 | 0 | 0.006094 | 0.002053 | 0.003313 | 0.001801 | 0.002299 | 0.005985 | 0.005552 |
| k__Bacteria;p__Bacteroidetes;c__Bacteroidia;o__Bacteroidales;f__NA;g__NA;s__sp12805 | 0 | 0 | 0 | 0 | 0 | 0 | 0 | 0 | 0.016021 | 0.024861 | 0.018728 | 0.018175 | 0.002543 | 0.052246 | 0.047278 | 0.041698 |
| k__Bacteria;p__Bacteroidetes;c__Bacteroidia;o__Bacteroidales;f__Porphyromonadaceae;g__Odoribacter;s__sp13184 | 0.017716 | 0.017061 | 0.022968 | 0.009344 | 0.00703 | 0.013835 | 0.005618 | 0.028158 | 0.018024 | 0.017563 | 0.016752 | 0.014477 | 0.007713 | 0.011919 | 0.007932 | 0.01725 |
| k__Bacteria;p__Bacteroidetes;c__Bacteroidia;o__Bacteroidales;f__Porphyromonadaceae;g__Parabacteroides;s__distasonis | 0 | 0 | 0 | 0 | 0 | 0 | 0 | 0 | 0 | 0 | 0 | 0 | 0 | 0 | 0 | 0.001068 |
| k__Bacteria;p__Bacteroidetes;c__Bacteroidia;o__Bacteroidales;f__Porphyromonadaceae;g__Parabacteroides;s__goldsteinii | 0 | 0.000492 | 0.000824 | 0.001412 | 0.000687 | 0 | 0.000803 | 0.001078 | 0.00081 | 0 | 0.000456 | 0.000456 | 0.001208 | 0.000449 | 0.001106 | 0.000961 |
| k__Bacteria;p__Bacteroidetes;c__Bacteroidia;o__Bacteroidales;f__Prevotellaceae;g__NA;s__sp14210 | 0.006743 | 0.010922 | 0.014478 | 0.039205 | 0.021467 | 0.004944 | 0.021016 | 0.006508 | 0.008438 | 0.015176 | 0.008794 | 0.013443 | 0.015065 | 0.013584 | 0.015575 | 0.018394 |
| k__Bacteria;p__Bacteroidetes;c__Bacteroidia;o__Bacteroidales;f__Rikenellaceae;g__Alistipes;s__putredinis | 0.002948 | 0.000748 | 0.001181 | 0.002907 | 0.000399 | 0.000658 | 0.00224 | 0.001246 | 0.00117 | 0.001854 | 0 | 0.000929 | 0.001144 | 0.001268 | 0.001009 | 0.002684 |
| k__Bacteria;p__Bacteroidetes;c__Bacteroidia;o__Bacteroidales;f__Rikenellaceae;g__Alistipes;s__sp14330 | 0.003051 | 0.010626 | 0.008901 | 0.021866 | 0.003615 | 0.004753 | 0.005749 | 0.005024 | 0.004388 | 0.001645 | 0.001394 | 0.001437 | 0.007755 | 0.001956 | 0.007139 | 0.003035 |
| k__Bacteria;p__Bacteroidetes;c__Bacteroidia;o__Bacteroidales;f__Rikenellaceae;g__Alistipes;s__sp14336 | 0.015793 | 0.025582 | 0.025056 | 0.047885 | 0.04034 | 0.022471 | 0.057841 | 0.041859 | 0.034495 | 0.053058 | 0.042779 | 0.044658 | 0.023838 | 0.023573 | 0.024492 | 0.018653 |
| k__Bacteria;p__Bacteroidetes;c__Bacteroidia;o__Bacteroidales;f__Rikenellaceae;g__Alistipes;s__sp14338 | 0.005128 | 0.008599 | 0.01772 | 0.008493 | 0.008893 | 0.005793 | 0.009855 | 0.008439 | 0.001215 | 0.001923 | 0.003649 | 0.001069 | 0.000805 | 0.004413 | 0.001755 | 0.00543 |
| k__Bacteria;p__Bacteroidetes;c__Bacteroidia;o__Bacteroidales;f__Rikenellaceae;g__Alistipes;s__sp14343 | 0.000487 | 0.000807 | 0.001786 | 0.001578 | 0.001264 | 0.000573 | 0.001064 | 0.00056 | 0.001328 | 0.001135 | 0.001875 | 0.000824 | 0.00089 | 0.002114 | 0.000673 | 0.001388 |
| k__Bacteria;p__Bacteroidetes;c__Bacteroidia;o__Bacteroidales;f__Rikenellaceae;g__Alistipes;s__sp14354 | 0.053378 | 0.049176 | 0.04632 | 0.036443 | 0.034042 | 0.051054 | 0.057692 | 0.043021 | 0.026035 | 0.042609 | 0.038039 | 0.021418 | 0.011929 | 0.022965 | 0.02406 | 0.02515 |
| k__Bacteria;p__Bacteroidetes;c__Bacteroidia;o__Bacteroidales;f__Rikenellaceae;g__Alistipes;s__timonensis | 0 | 0.00793 | 0.009066 | 0.007725 | 0.001109 | 0.001804 | 0.000952 | 0 | 0.0036 | 0.004356 | 0.004384 | 0.003838 | 0 | 0.001744 | 0.003028 | 0.001296 |
| k__Bacteria;p__Bacteroidetes;c__Bacteroidia;o__Bacteroidales;f__Rikenellaceae;g__NA;s__sp14636 | 0.02251 | 0.011768 | 0.026127 | 0.022136 | 0.007962 | 0.009931 | 0.008679 | 0.019537 | 0.002363 | 0.003916 | 0.002103 | 0.000981 | 0.006187 | 0.00222 | 0.00286 | 0.003127 |
| k__Bacteria;p__Cyanobacteria;c__Melainabacteria;o__Gastranaerophilales;f__NA;g__NA;s__sp23727 | 0.001128 | 0.001555 | 0.000824 | 0.002077 | 0.001752 | 0.001443 | 0.004237 | 0.002589 | 0.001778 | 0.003869 | 0.00441 | 0.001928 | 0.003539 | 0.001295 | 0.0025 | 0.001312 |
| k__Bacteria;p__Cyanobacteria;c__Melainabacteria;o__Gastranaerophilales;f__NA;g__NA;s__sp23729 | 0 | 0.002912 | 0.006539 | 0.00191 | 0 | 0.000891 | 0 | 0.001176 | 0.00117 | 0.002525 | 0.001571 | 0.001612 | 0.00464 | 0.001057 | 0.000961 | 0.001068 |
| k__Bacteria;p__Deferribacteres;c__Deferribacteres;o__Deferribacterales;f__Deferribacteraceae;g__Mucispirillum;s__sp23890 | 0.00223 | 0.003837 | 0 | 0.000768 | 0.001331 | 0.010015 | 0.002408 | 0.003387 | 0.00324 | 0.000394 | 0.000558 | 0.006362 | 0.002712 | 0.003013 | 0.002404 | 0.000702 |
| k__Bacteria;p__Firmicutes;c__Bacilli;o__Lactobacillales;f__Lactobacillaceae;g__Lactobacillus;s__NA | 0.000282 | 0.00059 | 0.000907 | 0.000831 | 0.002573 | 0 | 0 | 0 | 0.000383 | 0.000533 | 0 | 0 | 0 | 0.00222 | 0 | 0.001678 |
| k__Bacteria;p__Firmicutes;c__Bacilli;o__Lactobacillales;f__Lactobacillaceae;g__Lactobacillus;s__reuteri-vaginalis | 0.001615 | 0 | 0 | 0 | 0 | 0 | 0 | 0 | 0 | 0.002595 | 0.001242 | 0 | 0 | 0.005312 | 0 | 0.008892 |
| k__Bacteria;p__Firmicutes;c__Bacilli;o__Lactobacillales;f__Lactobacillaceae;g__Lactobacillus;s__taiwanensis | 0.00282 | 0 | 0 | 0 | 0 | 0 | 0 | 0 | 0 | 0.00241 | 0.002256 | 0 | 0 | 0.003938 | 0 | 0.01057 |
| k__Bacteria;p__Firmicutes;c__Clostridia;o__Clostridiales;f__Christensenellaceae;g__NA;s__sp30444 | 0 | 0 | 0 | 0.000581 | 0 | 0 | 0 | 0 | 0.000293 | 0.000394 | 0 | 0.000263 | 0.001187 | 0 | 0.000336 | 0.000366 |
| k__Bacteria;p__Firmicutes;c__Clostridia;o__Clostridiales;f__Family XIII;g__Anaerovorax;s__sp31505 | 0 | 0.000315 | 0 | 0 | 0 | 0 | 0 | 0 | 0 | 0 | 0 | 0 | 0 | 0 | 0 | 0 |
| k__Bacteria;p__Firmicutes;c__Clostridia;o__Clostridiales;f__Family XIII;g__NA;s__sp31521 | 0 | 0 | 0 | 0 | 0 | 0 | 0 | 0 | 0 | 0 | 0 | 0 | 0.000466 | 0 | 0 | 0 |
| k__Bacteria;p__Firmicutes;c__Clostridia;o__Clostridiales;f__Family XIII;g__NA;s__sp31531 | 0 | 0 | 0 | 0 | 0 | 0 | 0 | 0 | 0 | 0 | 0 | 0.00021 | 0 | 0 | 0 | 0 |
| k__Bacteria;p__Firmicutes;c__Clostridia;o__Clostridiales;f__Family XIII;g__NA;s__sp31564 | 0 | 0.000492 | 0 | 0 | 0 | 0.000255 | 0 | 0 | 0 | 0 | 0 | 0.000298 | 0 | 0 | 0 | 0 |
| k__Bacteria;p__Firmicutes;c__Clostridia;o__Clostridiales;f__Family XIII;g__NA;s__sp31651 | 0 | 0 | 0 | 0.000291 | 0.000421 | 0 | 0 | 0 | 0 | 0 | 0 | 0 | 0 | 0 | 0 | 0 |
| k__Bacteria;p__Firmicutes;c__Clostridia;o__Clostridiales;f__Lachnospiraceae;g__Acetatifactor;s__sp31893 | 0.000615 | 0.000689 | 0 | 0 | 0.000821 | 0.000594 | 0.000859 | 0 | 0 | 0 | 0 | 0 | 0 | 0 | 0 | 0 |
| k__Bacteria;p__Firmicutes;c__Clostridia;o__Clostridiales;f__Lachnospiraceae;g__Acetitomaculum;s__sp31923 | 0 | 0.000669 | 0 | 0 | 0 | 0.000594 | 0 | 0 | 0.00081 | 0.001738 | 0.00109 | 0.000859 | 0.002606 | 0.002273 | 0.000817 | 0.000793 |
| k__Bacteria;p__Firmicutes;c__Clostridia;o__Clostridiales;f__Lachnospiraceae;g__Blautia;s__coccoides | 0.001128 | 0 | 0 | 0.002263 | 0.005877 | 0.001231 | 0.005487 | 0 | 0.00099 | 0 | 0 | 0 | 0 | 0.012077 | 0 | 0.001998 |
| k__Bacteria;p__Firmicutes;c__Clostridia;o__Clostridiales;f__Lachnospiraceae;g__Blautia;s__coccoides-sp32015 | 0 | 0 | 0 | 0 | 0 | 0 | 0 | 0 | 0 | 0 | 0 | 0 | 0 | 0.004281 | 0 | 0 |

| | | | | | | | | | | | | | | | | |
|---|---|---|---|---|---|---|---|---|---|---|---|---|---|---|---|---|
| k__Bacteria;p__Firmicutes;c__Clostridia;o__Clostridiales;f__Lachnospiraceae;g__Blautia;s__coccoides-sp32038 | 0 | 0 | 0 | 0 | 0 | 0 | 0 | 0 | 0 | 0 | 0.000431 | 0 | 0 | 0 | 0 | 0 |
| k__Bacteria;p__Firmicutes;c__Clostridia;o__Clostridiales;f__Lachnospiraceae;g__Blautia;s__sp32015 | 0.001128 | 0.000748 | 0 | 0 | 0.001353 | 0.001019 | 0.001269 | 0.000518 | 0.000495 | 0 | 0 | 0 | 0.001187 | 0.011998 | 0.000817 | 0.002852 |
| k__Bacteria;p__Firmicutes;c__Clostridia;o__Clostridiales;f__Lachnospiraceae;g__Blautia;s__sp32038 | 0 | 0 | 0 | 0.000519 | 0 | 0 | 0 | 0 | 0 | 0 | 0.000659 | 0.00028 | 0 | 0 | 0 | 0 |
| k__Bacteria;p__Firmicutes;c__Clostridia;o__Clostridiales;f__Lachnospiraceae;g__Butyrivibrio;s__sp32115 | 0 | 0 | 0 | 0 | 0 | 0 | 0 | 0 | 0 | 0 | 0 | 0 | 0.025236 | 0 | 0 | 0 |
| k__Bacteria;p__Firmicutes;c__Clostridia;o__Clostridiales;f__Lachnospiraceae;g__Lachnoclostridium;s__NA | 0 | 0 | 0 | 0 | 0 | 0.002037 | 0 | 0 | 0 | 0 | 0 | 0 | 0 | 0 | 0 | 0.002425 |
| k__Bacteria;p__Firmicutes;c__Clostridia;o__Clostridiales;f__Lachnospiraceae;g__Lachnoclostridium;s__sp32344 | 0.000308 | 0 | 0 | 0 | 0 | 0.000297 | 0 | 0.000406 | 0.00036 | 0 | 0 | 0 | 0.000445 | 0 | 0 | 0 |
| k__Bacteria;p__Firmicutes;c__Clostridia;o__Clostridiales;f__Lachnospiraceae;g__Lachnoclostridium;s__sp32350-sp33756 | 0 | 0 | 0.000522 | 0 | 0 | 0 | 0 | 0 | 0 | 0 | 0 | 0 | 0 | 0 | 0 | 0 |
| k__Bacteria;p__Firmicutes;c__Clostridia;o__Clostridiales;f__Lachnospiraceae;g__Lachnoclostridium;s__sp32351 | 0 | 0 | 0 | 0 | 0.000288 | 0 | 0 | 0 | 0 | 0 | 0 | 0 | 0.0025 | 0.001427 | 0.000697 | 0 |
| k__Bacteria;p__Firmicutes;c__Clostridia;o__Clostridiales;f__Lachnospiraceae;g__Lachnoclostridium;s__sp32366 | 0 | 0 | 0 | 0 | 0 | 0 | 0 | 0.000966 | 0 | 0.000394 | 0 | 0 | 0.000509 | 0 | 0 | 0 |
| k__Bacteria;p__Firmicutes;c__Clostridia;o__Clostridiales;f__Lachnospiraceae;g__Lachnoclostridium;s__sp32387 | 0 | 0.000787 | 0 | 0 | 0 | 0.000382 | 0.000915 | 0 | 0 | 0 | 0 | 0 | 0 | 0 | 0 | 0 |
| k__Bacteria;p__Firmicutes;c__Clostridia;o__Clostridiales;f__Lachnospiraceae;g__Lachnoclostridium;s__sp32402 | 0 | 0 | 0 | 0 | 0 | 0 | 0 | 0 | 0 | 0 | 0 | 0 | 0.004852 | 0 | 0 | 0.000412 |
| k__Bacteria;p__Firmicutes;c__Clostridia;o__Clostridiales;f__Lachnospiraceae;g__Lachnoclostridium;s__sp32414 | 0.001666 | 0.013795 | 0 | 0 | 0.001331 | 0.006684 | 0.001474 | 0.002197 | 0.003173 | 0.005445 | 0.00484 | 0.015213 | 0.006759 | 0.004096 | 0.002259 | 0.001678 |
| k__Bacteria;p__Firmicutes;c__Clostridia;o__Clostridiales;f__Lachnospiraceae;g__Lachnoclostridium;s__sp32442 | 0.001308 | 0.005018 | 0 | 0 | 0 | 0.021601 | 0.000317 | 0.007851 | 0.011138 | 0.003429 | 0.005626 | 0.003383 | 0 | 0 | 0.002355 | 0 |
| k__Bacteria;p__Firmicutes;c__Clostridia;o__Clostridiales;f__Lachnospiraceae;g__Marvinbryantia;s__sp32979 | 0.010486 | 0.003109 | 0.007885 | 0.005274 | 0.012752 | 0.008063 | 0.004218 | 0.004856 | 0.003195 | 0 | 0.002382 | 0.002471 | 0.000614 | 0 | 0.002331 | 0.000702 |
| k__Bacteria;p__Firmicutes;c__Clostridia;o__Clostridiales;f__Lachnospiraceae;g__Marvinbryantia;s__sp32983 | 0 | 0 | 0 | 0 | 0 | 0 | 0 | 0.001232 | 0 | 0 | 0 | 0 | 0 | 0 | 0 | 0 |
| k__Bacteria;p__Firmicutes;c__Clostridia;o__Clostridiales;f__Lachnospiraceae;g__Mobilitalea;s__sp32997 | 0 | 0.000453 | 0 | 0 | 0 | 0 | 0 | 0 | 0 | 0 | 0.000431 | 0.000473 | 0 | 0 | 0 | 0 |
| k__Bacteria;p__Firmicutes;c__Clostridia;o__Clostridiales;f__Lachnospiraceae;g__NA;s__NA | 0 | 0.007222 | 0 | 0 | 0.000333 | 0.00348 | 0 | 0.002029 | 0 | 0 | 0.001622 | 0.000666 | 0 | 0 | 0.000312 | 0 |
| k__Bacteria;p__Firmicutes;c__Clostridia;o__Clostridiales;f__Lachnospiraceae;g__NA;s__sp32146 | 0.001308 | 0.002027 | 0 | 0 | 0.001198 | 0.002949 | 0.001307 | 0.000924 | 0.001508 | 0.00095 | 0.000735 | 0.000684 | 0 | 0.001427 | 0.001322 | 0 |
| k__Bacteria;p__Firmicutes;c__Clostridia;o__Clostridiales;f__Lachnospiraceae;g__NA;s__sp32147 | 0 | 0 | 0 | 0 | 0 | 0.00104 | 0 | 0.000434 | 0 | 0 | 0 | 0 | 0.000932 | 0 | 0 | 0 |
| k__Bacteria;p__Firmicutes;c__Clostridia;o__Clostridiales;f__Lachnospiraceae;g__NA;s__sp32156 | 0 | 0.000275 | 0 | 0 | 0 | 0 | 0 | 0.00021 | 0 | 0 | 0 | 0 | 0.000678 | 0.000344 | 0 | 0.000168 |
| k__Bacteria;p__Firmicutes;c__Clostridia;o__Clostridiales;f__Lachnospiraceae;g__NA;s__sp32161 | 0 | 0 | 0 | 0 | 0 | 0 | 0 | 0 | 0 | 0.000255 | 0 | 0 | 0 | 0 | 0 | 0 |
| k__Bacteria;p__Firmicutes;c__Clostridia;o__Clostridiales;f__Lachnospiraceae;g__NA;s__sp32165 | 0.002307 | 0.001555 | 0 | 0 | 0 | 0.000934 | 0 | 0.000952 | 0.002385 | 0.001182 | 0.000786 | 0.001297 | 0 | 0 | 0 | 0 |
| k__Bacteria;p__Firmicutes;c__Clostridia;o__Clostridiales;f__Lachnospiraceae;g__NA;s__sp32166 | 0.001026 | 0.00059 | 0 | 0 | 0 | 0 | 0 | 0.000784 | 0 | 0 | 0 | 0 | 0 | 0 | 0 | 0 |
| k__Bacteria;p__Firmicutes;c__Clostridia;o__Clostridiales;f__Lachnospiraceae;g__NA;s__sp32180 | 0 | 0 | 0 | 0 | 0 | 0.000467 | 0 | 0 | 0.002048 | 0.002665 | 0.001673 | 0.003751 | 0.009726 | 0 | 0.000937 | 0 |
| k__Bacteria;p__Firmicutes;c__Clostridia;o__Clostridiales;f__Lachnospiraceae;g__NA;s__sp32254 | 0.007717 | 0 | 0 | 0 | 0 | 0 | 0 | 0 | 0 | 0 | 0 | 0.000824 | 0.000614 | 0 | 0.001514 | 0.000336 |
| k__Bacteria;p__Firmicutes;c__Clostridia;o__Clostridiales;f__Lachnospiraceae;g__NA;s__sp32255 | 0 | 0.002519 | 0 | 0 | 0 | 0.001867 | 0 | 0.000322 | 0 | 0.003267 | 0.001825 | 0.009552 | 0.002755 | 0 | 0.001466 | 0 |
| k__Bacteria;p__Firmicutes;c__Clostridia;o__Clostridiales;f__Lachnospiraceae;g__NA;s__sp32257 | 0 | 0.007438 | 0.002912 | 0.002284 | 0.002595 | 0.005135 | 0.001754 | 0.006774 | 0.006931 | 0.004032 | 0.005069 | 0.005679 | 0.009599 | 0.00629 | 0.001226 | 0.002379 |
| k__Bacteria;p__Firmicutes;c__Clostridia;o__Clostridiales;f__Lachnospiraceae;g__NA;s__sp32261-sp32270 | 0 | 0 | 0 | 0 | 0 | 0.000424 | 0 | 0 | 0 | 0 | 0 | 0 | 0 | 0.000793 | 0 | 0.000427 |
| k__Bacteria;p__Firmicutes;c__Clostridia;o__Clostridiales;f__Lachnospiraceae;g__NA;s__sp32263 | 0 | 0 | 0 | 0 | 0 | 0 | 0 | 0 | 0.001395 | 0 | 0 | 0 | 0 | 0 | 0 | 0.000381 |
| k__Bacteria;p__Firmicutes;c__Clostridia;o__Clostridiales;f__Lachnospiraceae;g__NA;s__sp32264-sp32276 | 0 | 0 | 0 | 0 | 0 | 0 | 0 | 0 | 0.002318 | 0 | 0 | 0 | 0.013942 | 0 | 0 | 0 |
| k__Bacteria;p__Firmicutes;c__Clostridia;o__Clostridiales;f__Lachnospiraceae;g__NA;s__sp32270 | 0 | 0 | 0 | 0 | 0.002329 | 0 | 0.000989 | 0.001036 | 0 | 0 | 0 | 0 | 0 | 0 | 0 | 0 |
| k__Bacteria;p__Firmicutes;c__Clostridia;o__Clostridiales;f__Lachnospiraceae;g__NA;s__sp32486 | 0 | 0 | 0 | 0 | 0 | 0.001549 | 0 | 0 | 0 | 0 | 0.000735 | 0.001034 | 0.000975 | 0 | 0.000913 | 0 |
| k__Bacteria;p__Firmicutes;c__Clostridia;o__Clostridiales;f__Lachnospiraceae;g__NA;s__sp32498 | 0.000872 | 0.001102 | 0 | 0 | 0.000687 | 0 | 0.000915 | 0.001246 | 0.001058 | 0.001437 | 0 | 0 | 0.000636 | 0.00148 | 0 | 0 |
| k__Bacteria;p__Firmicutes;c__Clostridia;o__Clostridiales;f__Lachnospiraceae;g__NA;s__sp32510 | 0 | 0 | 0 | 0 | 0 | 0 | 0 | 0.000322 | 0 | 0 | 0 | 0 | 0 | 0 | 0 | 0 |
| k__Bacteria;p__Firmicutes;c__Clostridia;o__Clostridiales;f__Lachnospiraceae;g__NA;s__sp32590 | 0 | 0 | 0 | 0 | 0 | 0 | 0 | 0 | 0.005445 | 0 | 0.005651 | 0.006941 | 0.021719 | 0.001031 | 0.007355 | 0.001464 |
| k__Bacteria;p__Firmicutes;c__Clostridia;o__Clostridiales;f__Lachnospiraceae;g__NA;s__sp32593 | 0.001948 | 0 | 0 | 0 | 0 | 0.001294 | 0 | 0.000266 | 0.001485 | 0 | 0.00071 | 0.000491 | 0.001865 | 0 | 0 | 0 |
| k__Bacteria;p__Firmicutes;c__Clostridia;o__Clostridiales;f__Lachnospiraceae;g__NA;s__sp32593-sp32722 | 0 | 0 | 0 | 0 | 0 | 0 | 0 | 0 | 0 | 0.000834 | 0 | 0 | 0.002077 | 0 | 0 | 0 |
| k__Bacteria;p__Firmicutes;c__Clostridia;o__Clostridiales;f__Lachnospiraceae;g__NA;s__sp32594 | 0.003666 | 0 | 0.000907 | 0.000415 | 0 | 0.002652 | 0.001456 | 0.002113 | 0.005963 | 0 | 0.003523 | 0.002156 | 0 | 0.000951 | 0 | 0 |
| k__Bacteria;p__Firmicutes;c__Clostridia;o__Clostridiales;f__Lachnospiraceae;g__NA;s__sp32594-sp32647 | 0 | 0.001732 | 0 | 0 | 0.00153 | 0 | 0.000597 | 0.001651 | 0.000675 | 0.001228 | 0 | 0.000824 | 0 | 0 | 0.001658 | 0.001251 |
| k__Bacteria;p__Firmicutes;c__Clostridia;o__Clostridiales;f__Lachnospiraceae;g__NA;s__sp32596 | 0.003794 | 0.000905 | 0 | 0.004132 | 0.0165 | 0.002483 | 0.003434 | 0.006228 | 0.008978 | 0.00614 | 0.007679 | 0.014635 | 0.003729 | 0 | 0.002476 | 0 |
| k__Bacteria;p__Firmicutes;c__Clostridia;o__Clostridiales;f__Lachnospiraceae;g__NA;s__sp32597 | 0 | 0.004388 | 0 | 0 | 0 | 0.001782 | 0 | 0.001036 | 0 | 0 | 0 | 0 | 0 | 0 | 0 | 0 |
| k__Bacteria;p__Firmicutes;c__Clostridia;o__Clostridiales;f__Lachnospiraceae;g__NA;s__sp32617-sp32782 | 0.011665 | 0.006415 | 0 | 0.002222 | 0.001597 | 0.006811 | 0.003416 | 0.004101 | 0.005085 | 0.00146 | 0.001369 | 0.004031 | 0.005085 | 0.004149 | 0.007643 | 0.002105 |
| k__Bacteria;p__Firmicutes;c__Clostridia;o__Clostridiales;f__Lachnospiraceae;g__NA;s__sp32622 | 0.001436 | 0.003719 | 0 | 0 | 0 | 0.003225 | 0.00196 | 0.002085 | 0.00243 | 0.001089 | 0.001571 | 0.003015 | 0.001377 | 0.002273 | 0.001394 | 0 |
| k__Bacteria;p__Firmicutes;c__Clostridia;o__Clostridiales;f__Lachnospiraceae;g__NA;s__sp32623 | 0.002307 | 0.000531 | 0 | 0 | 0.000399 | 0.000637 | 0 | 0.000378 | 0 | 0.000765 | 0 | 0.00312 | 0 | 0.002088 | 0.001779 | 0.003203 |
| k__Bacteria;p__Firmicutes;c__Clostridia;o__Clostridiales;f__Lachnospiraceae;g__NA;s__sp32628-sp32767 | 0.016126 | 0.001692 | 0 | 0.000872 | 0.002284 | 0.007002 | 0.00978 | 0.003191 | 0.00072 | 0.000626 | 0.001014 | 0.001981 | 0.003496 | 0 | 0 | 0.00029 |
| k__Bacteria;p__Firmicutes;c__Clostridia;o__Clostridiales;f__Lachnospiraceae;g__NA;s__sp32630 | 0.00323 | 0 | 0 | 0 | 0 | 0.000509 | 0.000523 | 0 | 0 | 0 | 0.000963 | 0.002471 | 0 | 0.001744 | 0 | 0 |
| k__Bacteria;p__Firmicutes;c__Clostridia;o__Clostridiales;f__Lachnospiraceae;g__NA;s__sp32635-sp32668 | 0 | 0.00063 | 0 | 0 | 0 | 0 | 0 | 0 | 0 | 0 | 0 | 0 | 0 | 0 | 0 | 0 |
| k__Bacteria;p__Firmicutes;c__Clostridia;o__Clostridiales;f__Lachnospiraceae;g__NA;s__sp32638 | 0 | 0 | 0 | 0 | 0 | 0 | 0 | 0 | 0 | 0 | 0 | 0 | 0 | 0 | 0.002788 | 0 |
| k__Bacteria;p__Firmicutes;c__Clostridia;o__Clostridiales;f__Lachnospiraceae;g__NA;s__sp32638-sp32701 | 0 | 0 | 0 | 0 | 0 | 0 | 0 | 0 | 0 | 0 | 0 | 0 | 0.000424 | 0 | 0 | 0 |
| k__Bacteria;p__Firmicutes;c__Clostridia;o__Clostridiales;f__Lachnospiraceae;g__NA;s__sp32648 | 0 | 0 | 0.000824 | 0 | 0.001708 | 0.001591 | 0.010527 | 0.003149 | 0.000878 | 0.000626 | 0.001064 | 0.000841 | 0 | 0.000766 | 0.000745 | 0.000885 |
| k__Bacteria;p__Firmicutes;c__Clostridia;o__Clostridiales;f__Lachnospiraceae;g__NA;s__sp32649 | 0.000538 | 0.001102 | 0 | 0 | 0.000754 | 0.000679 | 0.000317 | 0 | 0 | 0 | 0 | 0.000631 | 0.000424 | 0.001031 | 0 | 0 |
| k__Bacteria;p__Firmicutes;c__Clostridia;o__Clostridiales;f__Lachnospiraceae;g__NA;s__sp32650 | 0 | 0.002243 | 0 | 0 | 0 | 0.001273 | 0 | 0 | 0.002903 | 0.009754 | 0.013026 | 0.005714 | 0.005276 | 0 | 0.012835 | 0 |
| k__Bacteria;p__Firmicutes;c__Clostridia;o__Clostridiales;f__Lachnospiraceae;g__NA;s__sp32655 | 0.007537 | 0.00244 | 0.000989 | 0.000602 | 0.002772 | 0.007215 | 0.003322 | 0.003821 | 0.000878 | 0.002132 | 0.001445 | 0.002647 | 0.00428 | 0.003251 | 0.001731 | 0.001312 |
| k__Bacteria;p__Firmicutes;c__Clostridia;o__Clostridiales;f__Lachnospiraceae;g__NA;s__sp32658 | 0.005256 | 0.001555 | 0.000824 | 0 | 0 | 0.002652 | 0 | 0.00655 | 0.007561 | 0.002155 | 0.003928 | 0.00461 | 0.008772 | 0.003039 | 0.015791 | 0.000885 |
| k__Bacteria;p__Firmicutes;c__Clostridia;o__Clostridiales;f__Lachnospiraceae;g__NA;s__sp32668 | 0.003282 | 0 | 0.001044 | 0 | 0 | 0.001698 | 0.000709 | 0.00452 | 0.00396 | 0.003777 | 0.002281 | 0.002804 | 0.003009 | 0.002854 | 0.005672 | 0.011149 |
| k__Bacteria;p__Firmicutes;c__Clostridia;o__Clostridiales;f__Lachnospiraceae;g__NA;s__sp32683-sp32693-sp32746 | 0 | 0 | 0 | 0 | 0 | 0.003904 | 0.002949 | 0 | 0 | 0 | 0 | 0 | 0 | 0 | 0 | 0 |
| k__Bacteria;p__Firmicutes;c__Clostridia;o__Clostridiales;f__Lachnospiraceae;g__NA;s__sp32685 | 0 | 0 | 0 | 0 | 0 | 0 | 0 | 0 | 0.013906 | 0 | 0 | 0 | 0 | 0 | 0 | 0 |
| k__Bacteria;p__Firmicutes;c__Clostridia;o__Clostridiales;f__Lachnospiraceae;g__NA;s__sp32693 | 0 | 0.001909 | 0 | 0 | 0 | 0.001188 | 0 | 0 | 0 | 0 | 0 | 0 | 0 | 0 | 0 | 0 |
| k__Bacteria;p__Firmicutes;c__Clostridia;o__Clostridiales;f__Lachnospiraceae;g__NA;s__sp32701 | 0.008076 | 0.012988 | 0 | 0.003592 | 0.015524 | 0.01061 | 0.01579 | 0.012931 | 0 | 0.006279 | 0.010644 | 0.009149 | 0.003645 | 0 | 0 | 0.000259 |
| k__Bacteria;p__Firmicutes;c__Clostridia;o__Clostridiales;f__Lachnospiraceae;g__NA;s__sp32704 | 0 | 0 | 0 | 0.010819 | 0.078174 | 0 | 0.077457 | 0.028732 | 0 | 0 | 0 | 0 | 0.000275 | 0 | 0.011417 | 0.000214 |
| k__Bacteria;p__Firmicutes;c__Clostridia;o__Clostridiales;f__Lachnospiraceae;g__NA;s__sp32711 | 0 | 0.000315 | 0 | 0 | 0 | 0 | 0 | 0 | 0 | 0 | 0 | 0 | 0 | 0 | 0 | 0 |
| k__Bacteria;p__Firmicutes;c__Clostridia;o__Clostridiales;f__Lachnospiraceae;g__NA;s__sp32719-sp32755 | 0 | 0.00675 | 0.001209 | 0 | 0 | 0.013029 | 0 | 0 | 0.008933 | 0.006951 | 0.006995 | 0.007519 | 0 | 0.00592 | 0.006129 | 0.001556 |
| k__Bacteria;p__Firmicutes;c__Clostridia;o__Clostridiales;f__Lachnospiraceae;g__NA;s__sp32721 | 0.000974 | 0.000689 | 0 | 0.001329 | 0.004014 | 0.000509 | 0.002408 | 0.002071 | 0.000473 | 0.001089 | 0.000786 | 0.000859 | 0.003433 | 0.000396 | 0.000457 | 0 |
| k__Bacteria;p__Firmicutes;c__Clostridia;o__Clostridiales;f__Lachnospiraceae;g__NA;s__sp32722-sp32777-sp33764 | 0 | 0 | 0 | 0 | 0 | 0 | 0 | 0.000714 | 0 | 0 | 0 | 0 | 0.001356 | 0 | 0 | 0 |
| k__Bacteria;p__Firmicutes;c__Clostridia;o__Clostridiales;f__Lachnospiraceae;g__NA;s__sp32727 | 0 | 0.067438 | 0.000907 | 0 | 0 | 0.061196 | 0 | 0.018124 | 0.058009 | 0.039203 | 0.039459 | 0.067022 | 0.051087 | 0.016993 | 0.007042 | 0.003066 |
| k__Bacteria;p__Firmicutes;c__Clostridia;o__Clostridiales;f__Lachnospiraceae;g__NA;s__sp32735 | 0.008178 | 0.002952 | 0.003352 | 0 | 0 | 0.036794 | 0 | 0 | 0.00234 | 0 | 0 | 0 | 0 | 0.030893 | 0.007643 | 0.007107 |
| k__Bacteria;p__Firmicutes;c__Clostridia;o__Clostridiales;f__Lachnospiraceae;g__NA;s__sp32741-sp33513 | 0 | 0 | 0 | 0 | 0 | 0 | 0 | 0 | 0.00135 | 0.00234 | 0.001369 | 0.002524 | 0 | 0 | 0.002788 | 0 |
| k__Bacteria;p__Firmicutes;c__Clostridia;o__Clostridiales;f__Lachnospiraceae;g__NA;s__sp32746 | 0.018382 | 0.057579 | 0.006758 | 0 | 0 | 0.00418 | 0 | 0.060654 | 0.077383 | 0.01126 | 0.039205 | 0.025606 | 0.049816 | 0.005946 | 0.015263 | 0.004728 |
| k__Bacteria;p__Firmicutes;c__Clostridia;o__Clostridiales;f__Lachnospiraceae;g__NA;s__sp32746-sp32777 | 0.001333 | 0 | 0 | 0.001267 | 0.008072 | 0 | 0.00489 | 0.001931 | 0 | 0 | 0 | 0 | 0 | 0.000423 | 0 | 0 |
| k__Bacteria;p__Firmicutes;c__Clostridia;o__Clostridiales;f__Lachnospiraceae;g__NA;s__sp32747 | 0.000615 | 0 | 0 | 0 | 0 | 0 | 0 | 0.000546 | 0 | 0 | 0 | 0 | 0 | 0 | 0 | 0 |
| k__Bacteria;p__Firmicutes;c__Clostridia;o__Clostridiales;f__Lachnospiraceae;g__NA;s__sp32753 | 0 | 0.005628 | 0.006181 | 0.000498 | 0 | 0 | 0 | 0.016822 | 0.014446 | 0.038624 | 0.01148 | 0.032337 | 0 | 0.009514 | 0.023867 | 0.008907 |
| k__Bacteria;p__Firmicutes;c__Clostridia;o__Clostridiales;f__Lachnospiraceae;g__NA;s__sp32758 | 0 | 0 | 0 | 0 | 0 | 0.000255 | 0 | 0 | 0 | 0 | 0 | 0 | 0 | 0 | 0 | 0 |
| k__Bacteria;p__Firmicutes;c__Clostridia;o__Clostridiales;f__Lachnospiraceae;g__NA;s__sp32758-sp32766 | 0.001461 | 0 | 0.001566 | 0.001184 | 0 | 0 | 0 | 0.000672 | 0 | 0.000718 | 0 | 0 | 0 | 0.00185 | 0 | 0 |
| k__Bacteria;p__Firmicutes;c__Clostridia;o__Clostridiales;f__Lachnospiraceae;g__NA;s__sp32761 | 0.010024 | 0 | 0 | 0 | 0 | 0.000955 | 0 | 0.001036 | 0.000473 | 0 | 0 | 0 | 0 | 0.000344 | 0 | 0.001357 |
| k__Bacteria;p__Firmicutes;c__Clostridia;o__Clostridiales;f__Lachnospiraceae;g__NA;s__sp32762 | 0 | 0 | 0 | 0 | 0 | 0 | 0 | 0 | 0 | 0 | 0 | 0 | 0 | 0 | 0.000481 | 0 |
| k__Bacteria;p__Firmicutes;c__Clostridia;o__Clostridiales;f__Lachnospiraceae;g__NA;s__sp32763 | 0.034278 | 0.002243 | 0 | 0.000727 | 0 | 0.00539 | 0.001437 | 0.00501 | 0.003105 | 0.00183 | 0.002433 | 0.005083 | 0.00089 | 0.001189 | 0 | 0.00029 |
| k__Bacteria;p__Firmicutes;c__Clostridia;o__Clostridiales;f__Lachnospiraceae;g__NA;s__sp32766 | 0.001795 | 0.000433 | 0 | 0.010798 | 0.038876 | 0 | 0.03843 | 0.014485 | 0.000518 | 0 | 0.001064 | 0 | 0 | 0.003832 | 0.005312 | 0.001159 |

| | | | | | | | | | | | | | | | | |
|---|---|---|---|---|---|---|---|---|---|---|---|---|---|---|---|---|
| k__Bacteria;p__Firmicutes;c__Clostridia;o__Clostridiales;f__Lachnospiraceae;g__NA;s__sp32778 | 0.007691 | 0 | 0.003159 | 0 | 0 | 0.031192 | 0 | 0.000196 | 0 | 0 | 0 | 0 | 0 | 0.024471 | 0.006273 | 0.006223 |
| k__Bacteria;p__Firmicutes;c__Clostridia;o__Clostridiales;f__Lachnospiraceae;g__NA;s__sp32791 | 0.007563 | 0.004683 | 0.000934 | 0.001121 | 0.000643 | 0.005666 | 0.002426 | 0.005052 | 0.010958 | 0.001228 | 0.00669 | 0.004539 | 0.000869 | 0.001903 | 0.001346 | 0.000656 |
| k__Bacteria;p__Firmicutes;c__Clostridia;o__Clostridiales;f__Lachnospiraceae;g__NA;s__sp32798 | 0 | 0 | 0 | 0 | 0 | 0 | 0 | 0 | 0 | 0 | 0 | 0.001052 | 0.002649 | 0 | 0 | 0 |
| k__Bacteria;p__Firmicutes;c__Clostridia;o__Clostridiales;f__Lachnospiraceae;g__NA;s__sp32799 | 0 | 0 | 0 | 0 | 0 | 0 | 0 | 0.003457 | 0.001845 | 0.001043 | 0.002636 | 0 | 0.001759 | 0 | 0 | 0 |
| k__Bacteria;p__Firmicutes;c__Clostridia;o__Clostridiales;f__Lachnospiraceae;g__NA;s__sp32815 | 0.000897 | 0 | 0 | 0 | 0 | 0.00034 | 0 | 0.000602 | 0 | 0.000278 | 0.000938 | 0.002261 | 0 | 0 | 0.000961 | 0 |
| k__Bacteria;p__Firmicutes;c__Clostridia;o__Clostridiales;f__Lachnospiraceae;g__NA;s__sp32816 | 0 | 0.002106 | 0 | 0.000685 | 0.001774 | 0.003883 | 0.00153 | 0.001651 | 0.001328 | 0 | 0.000836 | 0.001209 | 0.000826 | 0 | 0 | 0 |
| k__Bacteria;p__Firmicutes;c__Clostridia;o__Clostridiales;f__Lachnospiraceae;g__NA;s__sp32826 | 0 | 0.000826 | 0 | 0 | 0 | 0.000743 | 0 | 0 | 0 | 0.000626 | 0.000329 | 0.002314 | 0.000614 | 0 | 0.000577 | 0 |
| k__Bacteria;p__Firmicutes;c__Clostridia;o__Clostridiales;f__Lachnospiraceae;g__NA;s__sp32850 | 0 | 0.00061 | 0 | 0.006479 | 0 | 0.000488 | 0.005021 | 0.002295 | 0.00324 | 0 | 0.002965 | 0.002033 | 0.003856 | 0.000766 | 0.000889 | 0.002822 |
| k__Bacteria;p__Firmicutes;c__Clostridia;o__Clostridiales;f__Lachnospiraceae;g__NA;s__sp32856-sp32859 | 0.000949 | 0.001102 | 0 | 0 | 0 | 0 | 0 | 0.00105 | 0 | 0 | 0 | 0 | 0 | 0 | 0 | 0 |
| k__Bacteria;p__Firmicutes;c__Clostridia;o__Clostridiales;f__Lachnospiraceae;g__NA;s__sp32859 | 0 | 0.002676 | 0.001099 | 0.00081 | 0.004435 | 0.001698 | 0.001139 | 0 | 0.00063 | 0.002062 | 0 | 0.001087 | 0 | 0.001242 | 0.001514 | 0 |
| k__Bacteria;p__Firmicutes;c__Clostridia;o__Clostridiales;f__Lachnospiraceae;g__NA;s__sp32862 | 0 | 0.000472 | 0.000412 | 0 | 0 | 0.000509 | 0 | 0.000364 | 0 | 0 | 0 | 0 | 0.000699 | 0 | 0 | 0 |
| k__Bacteria;p__Firmicutes;c__Clostridia;o__Clostridiales;f__Lachnospiraceae;g__NA;s__sp32862-sp32872-sp32880 | 0 | 0 | 0 | 0 | 0 | 0 | 0 | 0 | 0.003263 | 0 | 0 | 0 | 0 | 0.003171 | 0 | 0 |
| k__Bacteria;p__Firmicutes;c__Clostridia;o__Clostridiales;f__Lachnospiraceae;g__NA;s__sp32872 | 0 | 0.002735 | 0 | 0 | 0 | 0.000679 | 0 | 0.000672 | 0.002295 | 0.00234 | 0.002661 | 0.002156 | 0.006632 | 0 | 0.002452 | 0.001693 |
| k__Bacteria;p__Firmicutes;c__Clostridia;o__Clostridiales;f__Lachnospiraceae;g__NA;s__sp32880 | 0 | 0 | 0 | 0 | 0 | 0 | 0 | 0 | 0 | 0 | 0 | 0 | 0.002246 | 0 | 0 | 0 |
| k__Bacteria;p__Firmicutes;c__Clostridia;o__Clostridiales;f__Lachnospiraceae;g__NA;s__sp32883 | 0 | 0.000453 | 0 | 0 | 0 | 0.000403 | 0 | 0.000574 | 0 | 0 | 0 | 0 | 0 | 0 | 0 | 0 |
| k__Bacteria;p__Firmicutes;c__Clostridia;o__Clostridiales;f__Lachnospiraceae;g__NA;s__sp32885 | 0 | 0 | 0 | 0 | 0 | 0 | 0 | 0 | 0 | 0 | 0 | 0 | 0.00142 | 0.000714 | 0 | 0.000336 |
| k__Bacteria;p__Firmicutes;c__Clostridia;o__Clostridiales;f__Lachnospiraceae;g__NA;s__sp32910 | 0.000923 | 0 | 0 | 0 | 0 | 0 | 0 | 0.001246 | 0 | 0 | 0.000279 | 0.000596 | 0 | 0 | 0 | 0 |
| k__Bacteria;p__Firmicutes;c__Clostridia;o__Clostridiales;f__Lachnospiraceae;g__NA;s__sp33413 | 0 | 0.001673 | 0 | 0 | 0.000421 | 0.001103 | 0.000728 | 0.000686 | 0.000518 | 0.001297 | 0.00223 | 0.001297 | 0.008815 | 0.001163 | 0.000913 | 0.000564 |
| k__Bacteria;p__Firmicutes;c__Clostridia;o__Clostridiales;f__Lachnospiraceae;g__NA;s__sp33416-sp33419 | 0.004846 | 0.005746 | 0 | 0.001018 | 0 | 0.005835 | 0 | 0.004604 | 0.003443 | 0.002873 | 0 | 0.000999 | 0.001314 | 0.005893 | 0 | 0.002654 |
| k__Bacteria;p__Firmicutes;c__Clostridia;o__Clostridiales;f__Lachnospiraceae;g__NA;s__sp33417 | 0.001615 | 0.001063 | 0 | 0.000831 | 0.001508 | 0.001358 | 0.001325 | 0.00035 | 0.001103 | 0.000695 | 0.001521 | 0.002804 | 0 | 0.003436 | 0.002115 | 0.001418 |
| k__Bacteria;p__Firmicutes;c__Clostridia;o__Clostridiales;f__Lachnospiraceae;g__NA;s__sp33418 | 0 | 0 | 0 | 0 | 0 | 0 | 0 | 0 | 0 | 0 | 0 | 0 | 0.022588 | 0 | 0 | 0 |
| k__Bacteria;p__Firmicutes;c__Clostridia;o__Clostridiales;f__Lachnospiraceae;g__NA;s__sp33419 | 0 | 0 | 0.001319 | 0 | 0 | 0.002419 | 0 | 0.001763 | 0 | 0 | 0 | 0 | 0 | 0.002273 | 0 | 0 |
| k__Bacteria;p__Firmicutes;c__Clostridia;o__Clostridiales;f__Lachnospiraceae;g__NA;s__sp33421-sp33679 | 0.002359 | 0.002243 | 0.00033 | 0.000415 | 0.001375 | 0.002949 | 0.000392 | 0.003121 | 0.00135 | 0.001923 | 0.001191 | 0.002629 | 0.000593 | 0.003118 | 0.000961 | 0.001556 |
| k__Bacteria;p__Firmicutes;c__Clostridia;o__Clostridiales;f__Lachnospiraceae;g__NA;s__sp33426 | 0.002743 | 0.001063 | 0 | 0 | 0.001087 | 0.001952 | 0 | 0.002421 | 0.000563 | 0 | 0 | 0.002243 | 0 | 0 | 0 | 0 |
| k__Bacteria;p__Firmicutes;c__Clostridia;o__Clostridiales;f__Lachnospiraceae;g__NA;s__sp33426-sp33492 | 0 | 0 | 0 | 0 | 0 | 0 | 0 | 0 | 0 | 0 | 0 | 0 | 0 | 0.000502 | 0 | 0 |
| k__Bacteria;p__Firmicutes;c__Clostridia;o__Clostridiales;f__Lachnospiraceae;g__NA;s__sp33428 | 0 | 0 | 0 | 0 | 0 | 0 | 0.002184 | 0.001134 | 0 | 0 | 0 | 0.00298 | 0.001928 | 0 | 0 | 0 |
| k__Bacteria;p__Firmicutes;c__Clostridia;o__Clostridiales;f__Lachnospiraceae;g__NA;s__sp33431-sp33645 | 0 | 0 | 0 | 0 | 0 | 0 | 0 | 0.000252 | 0.00045 | 0 | 0 | 0 | 0 | 0 | 0 | 0 |
| k__Bacteria;p__Firmicutes;c__Clostridia;o__Clostridiales;f__Lachnospiraceae;g__NA;s__sp33432 | 0.002179 | 0.000453 | 0 | 0 | 0 | 0.002737 | 0 | 0.000224 | 0 | 0 | 0 | 0.000561 | 0 | 0 | 0 | 0 |
| k__Bacteria;p__Firmicutes;c__Clostridia;o__Clostridiales;f__Lachnospiraceae;g__NA;s__sp33432-sp33629 | 0.003179 | 0 | 0 | 0 | 0 | 0 | 0 | 0 | 0 | 0 | 0 | 0 | 0 | 0 | 0 | 0 |
| k__Bacteria;p__Firmicutes;c__Clostridia;o__Clostridiales;f__Lachnospiraceae;g__NA;s__sp33433 | 0.000615 | 0.000669 | 0 | 0 | 0 | 0.001082 | 0 | 0.001148 | 0.00036 | 0.000394 | 0 | 0.000999 | 0.00053 | 0 | 0 | 0.000229 |
| k__Bacteria;p__Firmicutes;c__Clostridia;o__Clostridiales;f__Lachnospiraceae;g__NA;s__sp33436 | 0 | 0 | 0 | 0 | 0.00031 | 0 | 0 | 0 | 0 | 0 | 0 | 0 | 0 | 0 | 0 | 0 |
| k__Bacteria;p__Firmicutes;c__Clostridia;o__Clostridiales;f__Lachnospiraceae;g__NA;s__sp33436-sp33645 | 0 | 0 | 0 | 0 | 0 | 0 | 0 | 0 | 0 | 0 | 0 | 0 | 0 | 0.000793 | 0 | 0 |
| k__Bacteria;p__Firmicutes;c__Clostridia;o__Clostridiales;f__Lachnospiraceae;g__NA;s__sp33440 | 0 | 0.000767 | 0 | 0 | 0 | 0.000891 | 0 | 0 | 0 | 0 | 0 | 0.000543 | 0.001144 | 0.00111 | 0 | 0.000229 |
| k__Bacteria;p__Firmicutes;c__Clostridia;o__Clostridiales;f__Lachnospiraceae;g__NA;s__sp33451-sp33593 | 0 | 0 | 0 | 0 | 0.000377 | 0 | 0 | 0 | 0 | 0 | 0.000431 | 0.003137 | 0.001059 | 0.003436 | 0.000625 | 0.001373 |
| k__Bacteria;p__Firmicutes;c__Clostridia;o__Clostridiales;f__Lachnospiraceae;g__NA;s__sp33453 | 0.007332 | 0.00309 | 0 | 0.002118 | 0.002772 | 0.010758 | 0.003826 | 0.005276 | 0.003105 | 0.002479 | 0.001597 | 0.004224 | 0.000509 | 0.005391 | 0.00137 | 0.000763 |
| k__Bacteria;p__Firmicutes;c__Clostridia;o__Clostridiales;f__Lachnospiraceae;g__NA;s__sp33456 | 0 | 0 | 0 | 0 | 0 | 0.000446 | 0.000448 | 0.000378 | 0 | 0 | 0 | 0.000491 | 0.000233 | 0.000476 | 0 | 0 |
| k__Bacteria;p__Firmicutes;c__Clostridia;o__Clostridiales;f__Lachnospiraceae;g__NA;s__sp33459 | 0.002564 | 0.002322 | 0 | 0 | 0.001685 | 0.00278 | 0.002202 | 0.003303 | 0.001058 | 0.000695 | 0.001191 | 0.002314 | 0.001568 | 0.001771 | 0 | 0 |
| k__Bacteria;p__Firmicutes;c__Clostridia;o__Clostridiales;f__Lachnospiraceae;g__NA;s__sp33470 | 0 | 0.000689 | 0 | 0 | 0 | 0 | 0 | 0 | 0 | 0 | 0 | 0 | 0 | 0.001876 | 0.001202 | 0 |
| k__Bacteria;p__Firmicutes;c__Clostridia;o__Clostridiales;f__Lachnospiraceae;g__NA;s__sp33477 | 0.045584 | 0.015074 | 0.016896 | 0.000997 | 0 | 0.026715 | 0.001866 | 0 | 0.045071 | 0.038044 | 0.035074 | 0.019665 | 0.021337 | 0.015803 | 0.030693 | 0.012339 |
| k__Bacteria;p__Firmicutes;c__Clostridia;o__Clostridiales;f__Lachnospiraceae;g__NA;s__sp33478 | 0.001436 | 0.000748 | 0 | 0.000685 | 0.013683 | 0.000658 | 0.010695 | 0.001218 | 0.000338 | 0 | 0 | 0.001735 | 0.000297 | 0 | 0 | 0 |
| k__Bacteria;p__Firmicutes;c__Clostridia;o__Clostridiales;f__Lachnospiraceae;g__NA;s__sp33489 | 0 | 0 | 0 | 0 | 0 | 0 | 0 | 0.000854 | 0 | 0 | 0 | 0.001823 | 0.000869 | 0 | 0 | 0 |
| k__Bacteria;p__Firmicutes;c__Clostridia;o__Clostridiales;f__Lachnospiraceae;g__NA;s__sp33492 | 0 | 0.002893 | 0.001071 | 0 | 0 | 0.003777 | 0.001605 | 0.00473 | 0.003825 | 0.002456 | 0.002002 | 0.006082 | 0.010128 | 0.000872 | 0.014253 | 0.00029 |
| k__Bacteria;p__Firmicutes;c__Clostridia;o__Clostridiales;f__Lachnospiraceae;g__NA;s__sp33503 | 0 | 0.002401 | 0 | 0 | 0.001863 | 0.002037 | 0.002277 | 0.002379 | 0.00054 | 0 | 0 | 0.000894 | 0.001144 | 0.000766 | 0.000409 | 0 |
| k__Bacteria;p__Firmicutes;c__Clostridia;o__Clostridiales;f__Lachnospiraceae;g__NA;s__sp33503-sp33540 | 0 | 0.00181 | 0.000577 | 0 | 0 | 0.00227 | 0.001904 | 0.001148 | 0 | 0 | 0 | 0 | 0.000805 | 0 | 0 | 0 |
| k__Bacteria;p__Firmicutes;c__Clostridia;o__Clostridiales;f__Lachnospiraceae;g__NA;s__sp33513 | 0 | 0.000669 | 0 | 0.000727 | 0.006564 | 0.001019 | 0.00209 | 0.003681 | 0.00027 | 0 | 0 | 0.000526 | 0 | 0 | 0.000577 | 0 |
| k__Bacteria;p__Firmicutes;c__Clostridia;o__Clostridiales;f__Lachnospiraceae;g__NA;s__sp33518 | 0 | 0.000728 | 0 | 0 | 0.002905 | 0 | 0.003322 | 0 | 0 | 0 | 0 | 0 | 0 | 0 | 0 | 0.00061 |
| k__Bacteria;p__Firmicutes;c__Clostridia;o__Clostridiales;f__Lachnospiraceae;g__NA;s__sp33518-sp33694-sp33762 | 0 | 0 | 0.001374 | 0 | 0 | 0 | 0 | 0 | 0 | 0 | 0 | 0 | 0 | 0 | 0 | 0 |
| k__Bacteria;p__Firmicutes;c__Clostridia;o__Clostridiales;f__Lachnospiraceae;g__NA;s__sp33518-sp33762 | 0.003051 | 0 | 0 | 0.00189 | 0.000754 | 0.000488 | 0.001157 | 0.000686 | 0 | 0 | 0 | 0 | 0 | 0.001242 | 0 | 0 |
| k__Bacteria;p__Firmicutes;c__Clostridia;o__Clostridiales;f__Lachnospiraceae;g__NA;s__sp33522 | 0.002179 | 0.000925 | 0 | 0 | 0.000665 | 0.002355 | 0.000485 | 0.001036 | 0 | 0.000487 | 0 | 0.000543 | 0.000763 | 0.000555 | 0.000433 | 0 |
| k__Bacteria;p__Firmicutes;c__Clostridia;o__Clostridiales;f__Lachnospiraceae;g__NA;s__sp33524 | 0.019921 | 0.025621 | 0.002665 | 0 | 0.000732 | 0.041781 | 0.006253 | 0.021692 | 0.025539 | 0.014759 | 0.006969 | 0.042081 | 0 | 0.005153 | 0.002548 | 0.001296 |
| k__Bacteria;p__Firmicutes;c__Clostridia;o__Clostridiales;f__Lachnospiraceae;g__NA;s__sp33524-sp33714 | 0 | 0 | 0 | 0 | 0.000754 | 0 | 0 | 0 | 0 | 0 | 0 | 0 | 0.000805 | 0 | 0 | 0 |
| k__Bacteria;p__Firmicutes;c__Clostridia;o__Clostridiales;f__Lachnospiraceae;g__NA;s__sp33525 | 0.00082 | 0.000925 | 0 | 0 | 0.000621 | 0.001719 | 0.000821 | 0.000658 | 0.00045 | 0.000348 | 0 | 0.00163 | 0.001398 | 0 | 0.000721 | 0 |
| k__Bacteria;p__Firmicutes;c__Clostridia;o__Clostridiales;f__Lachnospiraceae;g__NA;s__sp33529 | 0.000667 | 0 | 0 | 0 | 0.001397 | 0.00191 | 0.001176 | 0.001022 | 0 | 0.000487 | 0.000482 | 0 | 0.000975 | 0.001216 | 0 | 0.000885 |
| k__Bacteria;p__Firmicutes;c__Clostridia;o__Clostridiales;f__Lachnospiraceae;g__NA;s__sp33529-sp33614 | 0 | 0 | 0 | 0 | 0 | 0 | 0 | 0 | 0.000248 | 0 | 0 | 0.00021 | 0 | 0 | 0 | 0 |
| k__Bacteria;p__Firmicutes;c__Clostridia;o__Clostridiales;f__Lachnospiraceae;g__NA;s__sp33531 | 0 | 0.003286 | 0 | 0 | 0 | 0.004414 | 0 | 0.00035 | 0.00198 | 0.001297 | 0.000355 | 0 | 0 | 0 | 0 | 0 |
| k__Bacteria;p__Firmicutes;c__Clostridia;o__Clostridiales;f__Lachnospiraceae;g__NA;s__sp33537 | 0 | 0.000807 | 0 | 0 | 0 | 0.00053 | 0 | 0.000518 | 0.00099 | 0.000579 | 0.00038 | 0.000859 | 0.00125 | 0.003356 | 0.003461 | 0.002395 |
| k__Bacteria;p__Firmicutes;c__Clostridia;o__Clostridiales;f__Lachnospiraceae;g__NA;s__sp33539 | 0 | 0.005077 | 0.001648 | 0 | 0 | 0.007172 | 0 | 0.001204 | 0 | 0.002943 | 0.002357 | 0.000946 | 0 | 0 | 0 | 0 |
| k__Bacteria;p__Firmicutes;c__Clostridia;o__Clostridiales;f__Lachnospiraceae;g__NA;s__sp33542 | 0 | 0 | 0 | 0 | 0 | 0 | 0 | 0 | 0 | 0 | 0 | 0 | 0 | 0 | 0 | 0.000198 |
| k__Bacteria;p__Firmicutes;c__Clostridia;o__Clostridiales;f__Lachnospiraceae;g__NA;s__sp33558 | 0 | 0 | 0 | 0 | 0 | 0 | 0 | 0 | 0 | 0.000487 | 0 | 0 | 0 | 0 | 0 | 0 |
| k__Bacteria;p__Firmicutes;c__Clostridia;o__Clostridiales;f__Lachnospiraceae;g__NA;s__sp33565 | 0.008717 | 0 | 0 | 0.000789 | 0.004923 | 0.000552 | 0.00252 | 0.001232 | 0 | 0 | 0 | 0 | 0.001441 | 0 | 0 | 0 |
| k__Bacteria;p__Firmicutes;c__Clostridia;o__Clostridiales;f__Lachnospiraceae;g__NA;s__sp33577 | 0 | 0 | 0 | 0 | 0.001397 | 0 | 0 | 0.001148 | 0 | 0 | 0 | 0.001805 | 0.001123 | 0.00111 | 0.000817 | 0 |
| k__Bacteria;p__Firmicutes;c__Clostridia;o__Clostridiales;f__Lachnospiraceae;g__NA;s__sp33579 | 0 | 0.000472 | 0 | 0 | 0 | 0.000424 | 0.000261 | 0.000854 | 0.000495 | 0 | 0 | 0 | 0 | 0 | 0 | 0 |
| k__Bacteria;p__Firmicutes;c__Clostridia;o__Clostridiales;f__Lachnospiraceae;g__NA;s__sp33582 | 0.008332 | 0.002676 | 0.002033 | 0.00081 | 0 | 0.002822 | 0 | 0.010342 | 0 | 0.002085 | 0 | 0.002541 | 0.001822 | 0.002484 | 0.004495 | 0 |
| k__Bacteria;p__Firmicutes;c__Clostridia;o__Clostridiales;f__Lachnospiraceae;g__NA;s__sp33589 | 0 | 0 | 0 | 0 | 0 | 0 | 0 | 0.000308 | 0 | 0 | 0 | 0 | 0 | 0.000872 | 0 | 0 |
| k__Bacteria;p__Firmicutes;c__Clostridia;o__Clostridiales;f__Lachnospiraceae;g__NA;s__sp33597 | 0.003256 | 0.006828 | 0.001456 | 0.000561 | 0.000998 | 0.004095 | 0.00084 | 0.001903 | 0 | 0.001437 | 0.001115 | 0.001683 | 0.003645 | 0.001401 | 0.002572 | 0 |
| k__Bacteria;p__Firmicutes;c__Clostridia;o__Clostridiales;f__Lachnospiraceae;g__NA;s__sp33614 | 0 | 0.00183 | 0 | 0 | 0 | 0 | 0 | 0 | 0 | 0 | 0 | 0 | 0 | 0 | 0 | 0 |
| k__Bacteria;p__Firmicutes;c__Clostridia;o__Clostridiales;f__Lachnospiraceae;g__NA;s__sp33616-sp33639 | 0.000718 | 0.001377 | 0 | 0 | 0 | 0.000828 | 0.000541 | 0.000882 | 0 | 0 | 0.000279 | 0.001788 | 0 | 0 | 0 | 0 |
| k__Bacteria;p__Firmicutes;c__Clostridia;o__Clostridiales;f__Lachnospiraceae;g__NA;s__sp33620 | 0 | 0.000374 | 0 | 0 | 0.000266 | 0.000637 | 0.000411 | 0 | 0 | 0.000602 | 0 | 0.000999 | 0.001144 | 0.001136 | 0.001418 | 0 |
| k__Bacteria;p__Firmicutes;c__Clostridia;o__Clostridiales;f__Lachnospiraceae;g__NA;s__sp33625 | 0.001256 | 0.001043 | 0 | 0 | 0.000732 | 0.001358 | 0.000877 | 0.0007 | 0.000653 | 0.00051 | 0.000405 | 0.000876 | 0 | 0.000317 | 0.000577 | 0 |
| k__Bacteria;p__Firmicutes;c__Clostridia;o__Clostridiales;f__Lachnospiraceae;g__NA;s__sp33636 | 0.002615 | 0.001318 | 0.000495 | 0.003032 | 0.007141 | 0.002143 | 0.004871 | 0.004688 | 0 | 0.002966 | 0.001571 | 0.001665 | 0 | 0.000872 | 0.001971 | 0.000259 |
| k__Bacteria;p__Firmicutes;c__Clostridia;o__Clostridiales;f__Lachnospiraceae;g__NA;s__sp33639 | 0.001205 | 0.00061 | 0 | 0 | 0 | 0.000997 | 0 | 0.00049 | 0 | 0 | 0 | 0 | 0 | 0 | 0 | 0 |
| k__Bacteria;p__Firmicutes;c__Clostridia;o__Clostridiales;f__Lachnospiraceae;g__NA;s__sp33645 | 0 | 0 | 0 | 0 | 0 | 0 | 0 | 0 | 0 | 0.000857 | 0 | 0 | 0 | 0 | 0 | 0.000229 |
| k__Bacteria;p__Firmicutes;c__Clostridia;o__Clostridiales;f__Lachnospiraceae;g__NA;s__sp33645-sp33744 | 0.002359 | 0.00183 | 0 | 0.000395 | 0.001464 | 0.00575 | 0.003509 | 0.00459 | 0 | 0 | 0 | 0 | 0 | 0.001321 | 0.002283 | 0.000503 |
| k__Bacteria;p__Firmicutes;c__Clostridia;o__Clostridiales;f__Lachnospiraceae;g__NA;s__sp33658 | 0.001461 | 0.006632 | 0.001071 | 0.001765 | 0.003482 | 0.012159 | 0.008978 | 0.008257 | 0.002453 | 0.001112 | 0.001951 | 0.00326 | 0.005128 | 0.003092 | 0.004182 | 0 |
| k__Bacteria;p__Firmicutes;c__Clostridia;o__Clostridiales;f__Lachnospiraceae;g__NA;s__sp33669 | 0 | 0 | 0 | 0 | 0.001087 | 0 | 0.003472 | 0.002323 | 0 | 0 | 0.00038 | 0.001612 | 0.001335 | 0 | 0 | 0 |
| k__Bacteria;p__Firmicutes;c__Clostridia;o__Clostridiales;f__Lachnospiraceae;g__NA;s__sp33670 | 0.004307 | 0.002539 | 0.000934 | 0.001807 | 0.007562 | 0.008339 | 0.009631 | 0.003597 | 0.00279 | 0.002016 | 0.001622 | 0.003313 | 0.001632 | 0.003198 | 0.002572 | 0.000946 |
| k__Bacteria;p__Firmicutes;c__Clostridia;o__Clostridiales;f__Lachnospiraceae;g__NA;s__sp33673 | 0.004358 | 0.002539 | 0 | 0 | 0.000488 | 0.002016 | 0.001325 | 0.004282 | 0 | 0.000417 | 0.00076 | 0.000771 | 0.000996 | 0.000925 | 0 | 0 |

| | | | | | | | | | | | | | | | | |
|---|---|---|---|---|---|---|---|---|---|---|---|---|---|---|---|---|
| k__Bacteria;p__Firmicutes;c__Clostridia;o__Clostridiales;f__Lachnospiraceae;g__NA;s__sp33674 | 0 | 0 | 0 | 0.000768 | 0.00153 | 0 | 0.004666 | 0.005556 | 0 | 0 | 0 | 0 | 0 | 0 | 0 | 0 |
| k__Bacteria;p__Firmicutes;c__Clostridia;o__Clostridiales;f__Lachnospiraceae;g__NA;s__sp33682 | 0 | 0.000767 | 0 | 0 | 0.000488 | 0.000637 | 0 | 0.00112 | 0 | 0.000695 | 0 | 0.00163 | 0.000826 | 0.000925 | 0.000745 | 0 |
| k__Bacteria;p__Firmicutes;c__Clostridia;o__Clostridiales;f__Lachnospiraceae;g__NA;s__sp33690 | 0 | 0 | 0.000577 | 0.000415 | 0 | 0 | 0 | 0.000448 | 0 | 0.000417 | 0 | 0 | 0.000593 | 0.000846 | 0 | 0 |
| k__Bacteria;p__Firmicutes;c__Clostridia;o__Clostridiales;f__Lachnospiraceae;g__NA;s__sp33694 | 0 | 0 | 0 | 0 | 0 | 0 | 0 | 0 | 0 | 0 | 0 | 0 | 0 | 0 | 0.000553 | 0 |
| k__Bacteria;p__Firmicutes;c__Clostridia;o__Clostridiales;f__Lachnospiraceae;g__NA;s__sp33711 | 0.001026 | 0 | 0 | 0 | 0 | 0 | 0 | 0 | 0 | 0 | 0 | 0 | 0 | 0 | 0 | 0 |
| k__Bacteria;p__Firmicutes;c__Clostridia;o__Clostridiales;f__Lachnospiraceae;g__NA;s__sp33718 | 0.009691 | 0.002853 | 0 | 0.001516 | 0.002062 | 0.003289 | 0.002576 | 0.003121 | 0.001103 | 0.001483 | 0.001445 | 0.002927 | 0 | 0.000951 | 0.000793 | 0.000168 |
| k__Bacteria;p__Firmicutes;c__Clostridia;o__Clostridiales;f__Lachnospiraceae;g__NA;s__sp33721 | 0 | 0.000433 | 0 | 0 | 0 | 0 | 0.000317 | 0.00028 | 0 | 0 | 0 | 0 | 0 | 0 | 0 | 0.000198 |
| k__Bacteria;p__Firmicutes;c__Clostridia;o__Clostridiales;f__Lachnospiraceae;g__NA;s__sp33725 | 0 | 0.00059 | 0 | 0 | 0 | 0 | 0 | 0 | 0 | 0 | 0 | 0 | 0 | 0 | 0 | 0 |
| k__Bacteria;p__Firmicutes;c__Clostridia;o__Clostridiales;f__Lachnospiraceae;g__NA;s__sp33731 | 0.00082 | 0.000728 | 0 | 0.000602 | 0.000909 | 0.000594 | 0.001549 | 0.001861 | 0 | 0.00051 | 0.00071 | 0 | 0.001038 | 0.001559 | 0.000913 | 0.00029 |
| k__Bacteria;p__Firmicutes;c__Clostridia;o__Clostridiales;f__Lachnospiraceae;g__NA;s__sp33733 | 0.00041 | 0.000354 | 0 | 0 | 0.000377 | 0.000955 | 0.000485 | 0.000266 | 0.000495 | 0.000417 | 0.000279 | 0.000561 | 0.000403 | 0 | 0 | 0 |
| k__Bacteria;p__Firmicutes;c__Clostridia;o__Clostridiales;f__Lachnospiraceae;g__NA;s__sp33734 | 0.002512 | 0 | 0 | 0 | 0 | 0 | 0 | 0 | 0.000293 | 0 | 0 | 0 | 0 | 0 | 0 | 0.000351 |
| k__Bacteria;p__Firmicutes;c__Clostridia;o__Clostridiales;f__Lachnospiraceae;g__NA;s__sp33738 | 0 | 0.000275 | 0 | 0.00027 | 0 | 0.000615 | 0 | 0 | 0 | 0 | 0 | 0 | 0 | 0.000344 | 0 | 0.000259 |
| k__Bacteria;p__Firmicutes;c__Clostridia;o__Clostridiales;f__Lachnospiraceae;g__NA;s__sp33744 | 0 | 0.000866 | 0 | 0 | 0 | 0.001188 | 0 | 0.000756 | 0 | 0 | 0 | 0.000351 | 0.00267 | 0.002299 | 0.001851 | 0.000671 |
| k__Bacteria;p__Firmicutes;c__Clostridia;o__Clostridiales;f__Lachnospiraceae;g__NA;s__sp33755 | 0.008435 | 0.00551 | 0.001291 | 0.000748 | 0.002173 | 0.005305 | 0.00349 | 0.002099 | 0.001755 | 0.002108 | 0.003016 | 0.004522 | 0 | 0.002933 | 0 | 0 |
| k__Bacteria;p__Firmicutes;c__Clostridia;o__Clostridiales;f__Lachnospiraceae;g__NA;s__sp33756 | 0 | 0.002283 | 0 | 0 | 0 | 0.001698 | 0 | 0.003429 | 0.002565 | 0.00139 | 0 | 0.002576 | 0.002055 | 0.002114 | 0.000697 | 0.000458 |
| k__Bacteria;p__Firmicutes;c__Clostridia;o__Clostridiales;f__Lachnospiraceae;g__NA;s__sp33760 | 0.001692 | 0.000708 | 0 | 0 | 0.00122 | 0.000912 | 0.000709 | 0.000784 | 0.00099 | 0.000718 | 0 | 0.001595 | 0.001144 | 0 | 0 | 0 |
| k__Bacteria;p__Firmicutes;c__Clostridia;o__Clostridiales;f__Lachnospiraceae;g__NA;s__sp33766 | 0.009409 | 0.005038 | 0.001539 | 0.001474 | 0.00377 | 0.010397 | 0.007298 | 0.006732 | 0.004793 | 0.003707 | 0.005297 | 0.006169 | 0.001483 | 0.002537 | 0.010335 | 0.001601 |
| k__Bacteria;p__Firmicutes;c__Clostridia;o__Clostridiales;f__Lachnospiraceae;g__NA;s__sp33767 | 0 | 0.002224 | 0.000302 | 0.000706 | 0.004613 | 0.001082 | 0.001512 | 0.001959 | 0 | 0 | 0 | 0.000631 | 0.000678 | 0 | 0.00286 | 0 |
| k__Bacteria;p__Firmicutes;c__Clostridia;o__Clostridiales;f__Lachnospiraceae;g__Roseburia;s__sp33112 | 0.000974 | 0 | 0 | 0 | 0 | 0 | 0 | 0.000308 | 0 | 0 | 0.000532 | 0.002401 | 0 | 0 | 0 | 0.000244 |
| k__Bacteria;p__Firmicutes;c__Clostridia;o__Clostridiales;f__Lachnospiraceae;g__Roseburia;s__sp33112-sp33140 | 0 | 0.000492 | 0 | 0 | 0 | 0 | 0 | 0 | 0 | 0 | 0 | 0 | 0 | 0 | 0 | 0 |
| k__Bacteria;p__Firmicutes;c__Clostridia;o__Clostridiales;f__Lachnospiraceae;g__Roseburia;s__sp33139 | 0 | 0 | 0 | 0 | 0 | 0 | 0 | 0.00021 | 0 | 0 | 0 | 0 | 0 | 0 | 0 | 0 |
| k__Bacteria;p__Firmicutes;c__Clostridia;o__Clostridiales;f__Lachnospiraceae;g__Roseburia;s__sp33140 | 0 | 0.00124 | 0 | 0 | 0 | 0 | 0 | 0 | 0 | 0 | 0 | 0 | 0 | 0 | 0 | 0 |
| k__Bacteria;p__Firmicutes;c__Clostridia;o__Clostridiales;f__Lachnospiraceae;g__Roseburia;s__sp33143 | 0 | 0 | 0 | 0 | 0 | 0.000403 | 0 | 0 | 0 | 0 | 0 | 0 | 0 | 0 | 0 | 0 |
| k__Bacteria;p__Firmicutes;c__Clostridia;o__Clostridiales;f__Lachnospiraceae;g__Roseburia;s__sp33163 | 0 | 0 | 0 | 0 | 0 | 0.000488 | 0.000355 | 0.00021 | 0 | 0 | 0 | 0 | 0.000636 | 0.000819 | 0.000312 | 0 |
| k__Bacteria;p__Firmicutes;c__Clostridia;o__Clostridiales;f__Lachnospiraceae;g__Roseburia;s__sp33164-sp33173 | 0.000461 | 0 | 0 | 0 | 0 | 0.001061 | 0 | 0 | 0 | 0 | 0 | 0 | 0 | 0 | 0 | 0.00032 |
| k__Bacteria;p__Firmicutes;c__Clostridia;o__Clostridiales;f__Lachnospiraceae;g__Roseburia;s__sp33174 | 0 | 0.000216 | 0 | 0 | 0 | 0 | 0 | 0 | 0 | 0 | 0.000329 | 0 | 0 | 0 | 0.000625 | 0 |
| k__Bacteria;p__Firmicutes;c__Clostridia;o__Clostridiales;f__Lachnospiraceae;g__Roseburia;s__sp33177 | 0.001641 | 0.000846 | 0 | 0.000581 | 0.001929 | 0.0014 | 0.000747 | 0.00133 | 0 | 0.000417 | 0.000836 | 0.000754 | 0 | 0 | 0 | 0 |
| k__Bacteria;p__Firmicutes;c__Clostridia;o__Clostridiales;f__Lachnospiraceae;g__Roseburia;s__sp33183 | 0 | 0 | 0 | 0 | 0 | 0 | 0 | 0 | 0 | 0.000834 | 0.000482 | 0 | 0 | 0 | 0 | 0 |
| k__Bacteria;p__Firmicutes;c__Clostridia;o__Clostridiales;f__Lachnospiraceae;g__Tyzzerella;s__sp33258-sp33301 | 0.002589 | 0.000315 | 0 | 0 | 0 | 0.001209 | 0.001008 | 0.00133 | 0 | 0.000718 | 0.001521 | 0.002401 | 0 | 0.001453 | 0 | 0.000397 |
| k__Bacteria;p__Firmicutes;c__Clostridia;o__Clostridiales;f__Lachnospiraceae;g__Tyzzerella;s__sp33281 | 0 | 0 | 0 | 0 | 0 | 0.000361 | 0.000261 | 0 | 0.00045 | 0 | 0 | 0 | 0 | 0 | 0 | 0 |
| k__Bacteria;p__Firmicutes;c__Clostridia;o__Clostridiales;f__Lachnospiraceae;g__Tyzzerella;s__sp33287 | 0.001 | 0.000807 | 0 | 0.000395 | 0.000399 | 0.000594 | 0.000859 | 0.000826 | 0.0009 | 0 | 0.000456 | 0.000508 | 0.001271 | 0.00037 | 0.001274 | 0 |
| k__Bacteria;p__Firmicutes;c__Clostridia;o__Clostridiales;f__NA;g__NA;s__sp31072 | 0.001487 | 0.000492 | 0 | 0 | 0.000599 | 0 | 0.000317 | 0.000196 | 0.001148 | 0.001344 | 0.000634 | 0.001595 | 0 | 0 | 0 | 0 |
| k__Bacteria;p__Firmicutes;c__Clostridia;o__Clostridiales;f__NA;g__NA;s__sp31075 | 0 | 0 | 0 | 0 | 0 | 0.000276 | 0 | 0.000462 | 0 | 0 | 0 | 0 | 0 | 0 | 0 | 0 |
| k__Bacteria;p__Firmicutes;c__Clostridia;o__Clostridiales;f__NA;g__NA;s__sp31082 | 0.001026 | 0 | 0 | 0.002388 | 0.007341 | 0.001125 | 0.000933 | 0.004758 | 0 | 0.001483 | 0 | 0 | 0.000805 | 0 | 0.002428 | 0 |
| k__Bacteria;p__Firmicutes;c__Clostridia;o__Clostridiales;f__NA;g__NA;s__sp31103 | 0.000308 | 0.000551 | 0 | 0.000332 | 0 | 0.000467 | 0.000205 | 0.000252 | 0.000653 | 0 | 0 | 0.000228 | 0 | 0 | 0 | 0 |
| k__Bacteria;p__Firmicutes;c__Clostridia;o__Clostridiales;f__NA;g__NA;s__sp31106 | 0 | 0 | 0 | 0 | 0 | 0 | 0 | 0 | 0.000338 | 0 | 0 | 0.000315 | 0 | 0 | 0 | 0 |
| k__Bacteria;p__Firmicutes;c__Clostridia;o__Clostridiales;f__NA;g__NA;s__sp31115 | 0.010588 | 0.005549 | 0.007445 | 0 | 0 | 0.007787 | 0.005749 | 0.004856 | 0.002295 | 0 | 0.0037 | 0.0017 | 0 | 0.001665 | 0 | 0.003264 |
| k__Bacteria;p__Firmicutes;c__Clostridia;o__Clostridiales;f__NA;g__NA;s__sp31115-sp31116 | 0 | 0 | 0 | 0 | 0 | 0 | 0.000205 | 0 | 0 | 0 | 0 | 0 | 0 | 0 | 0 | 0 |
| k__Bacteria;p__Firmicutes;c__Clostridia;o__Clostridiales;f__NA;g__NA;s__sp31116 | 0.00059 | 0.000551 | 0 | 0 | 0 | 0.000276 | 0 | 0 | 0 | 0.000394 | 0 | 0.000491 | 0 | 0 | 0 | 0 |
| k__Bacteria;p__Firmicutes;c__Clostridia;o__Clostridiales;f__NA;g__NA;s__sp31119 | 0 | 0 | 0 | 0 | 0 | 0 | 0 | 0 | 0.001215 | 0 | 0.000431 | 0.00021 | 0 | 0 | 0 | 0 |
| k__Bacteria;p__Firmicutes;c__Clostridia;o__Clostridiales;f__NA;g__NA;s__sp31125 | 0 | 0.000571 | 0.000385 | 0 | 0 | 0 | 0 | 0.000322 | 0.002475 | 0 | 0.000456 | 0.000578 | 0 | 0 | 0 | 0.000519 |
| k__Bacteria;p__Firmicutes;c__Clostridia;o__Clostridiales;f__Peptococcaceae;g__NA;s__sp34196 | 0 | 0 | 0 | 0.00027 | 0 | 0 | 0 | 0 | 0 | 0 | 0 | 0 | 0 | 0 | 0 | 0.000229 |
| k__Bacteria;p__Firmicutes;c__Clostridia;o__Clostridiales;f__Peptococcaceae;g__NA;s__sp34246 | 0.001538 | 0.001928 | 0.000385 | 0 | 0.000532 | 0.001443 | 0.001698 | 0.002267 | 0.001913 | 0.001344 | 0.00114 | 0.001209 | 0.001695 | 0.000978 | 0.001322 | 0.000458 |
| k__Bacteria;p__Firmicutes;c__Clostridia;o__Clostridiales;f__Peptococcaceae;g__Peptococcus;s__sp34117 | 0.000615 | 0.000374 | 0 | 0 | 0 | 0.000764 | 0 | 0.000532 | 0.00054 | 0 | 0 | 0.000438 | 0.000318 | 0.00037 | 0 | 0 |
| k__Bacteria;p__Firmicutes;c__Clostridia;o__Clostridiales;f__Peptostreptococcaceae;g__Romboutsia;s__ilealis | 0 | 0.000531 | 0.00239 | 0 | 0 | 0 | 0 | 0 | 0 | 0 | 0 | 0 | 0 | 0.001136 | 0 | 0 |
| k__Bacteria;p__Firmicutes;c__Clostridia;o__Clostridiales;f__Ruminococcaceae;g__Anaerotruncus;s__colihominis | 0 | 0 | 0 | 0 | 0.00051 | 0 | 0 | 0 | 0 | 0 | 0 | 0 | 0 | 0 | 0 | 0 |
| k__Bacteria;p__Firmicutes;c__Clostridia;o__Clostridiales;f__Ruminococcaceae;g__Anaerotruncus;s__sp34437 | 0.002128 | 0.00061 | 0.001429 | 0.000789 | 0 | 0.000637 | 0 | 0.000476 | 0.001035 | 0.000649 | 0 | 0.000894 | 0.00053 | 0.000476 | 0.000745 | 0.000686 |
| k__Bacteria;p__Firmicutes;c__Clostridia;o__Clostridiales;f__Ruminococcaceae;g__Anaerotruncus;s__sp34438 | 0.000615 | 0.000571 | 0 | 0 | 0 | 0.001464 | 0.000485 | 0.00126 | 0.000585 | 0.000324 | 0.000482 | 0.000719 | 0 | 0 | 0.000312 | 0 |
| k__Bacteria;p__Firmicutes;c__Clostridia;o__Clostridiales;f__Ruminococcaceae;g__Anaerotruncus;s__sp34443 | 0 | 0 | 0 | 0 | 0 | 0 | 0 | 0.000182 | 0 | 0 | 0 | 0 | 0.000275 | 0 | 0 | 0 |
| k__Bacteria;p__Firmicutes;c__Clostridia;o__Clostridiales;f__Ruminococcaceae;g__Anaerotruncus;s__sp34450 | 0 | 0.000512 | 0 | 0 | 0.000266 | 0.000721 | 0 | 0.001134 | 0.00135 | 0 | 0.000583 | 0.000613 | 0 | 0 | 0.000433 | 0 |
| k__Bacteria;p__Firmicutes;c__Clostridia;o__Clostridiales;f__Ruminococcaceae;g__Anaerotruncus;s__sp34452-sp34475 | 0.003359 | 0.001909 | 0 | 0.001433 | 0.001464 | 0.002928 | 0.002781 | 0.002141 | 0.000833 | 0.000857 | 0 | 0.002384 | 0 | 0.001057 | 0.003413 | 0.000336 |
| k__Bacteria;p__Firmicutes;c__Clostridia;o__Clostridiales;f__Ruminococcaceae;g__Anaerotruncus;s__sp34453 | 0.007358 | 0.008127 | 0.000412 | 0.000872 | 0.005522 | 0.009336 | 0.007858 | 0.00578 | 0.002835 | 0.002919 | 0.003903 | 0.014722 | 0.001992 | 0.00629 | 0.004807 | 0.001662 |
| k__Bacteria;p__Firmicutes;c__Clostridia;o__Clostridiales;f__Ruminococcaceae;g__Anaerotruncus;s__sp34455 | 0 | 0.000394 | 0 | 0 | 0 | 0.001019 | 0.00028 | 0.000826 | 0 | 0 | 0 | 0.000648 | 0 | 0 | 0 | 0 |
| k__Bacteria;p__Firmicutes;c__Clostridia;o__Clostridiales;f__Ruminococcaceae;g__Anaerotruncus;s__sp34457 | 0.000282 | 0 | 0.000385 | 0 | 0 | 0.000509 | 0.000429 | 0.00035 | 0.000585 | 0.000417 | 0 | 0.000263 | 0.000297 | 0 | 0 | 0 |
| k__Bacteria;p__Firmicutes;c__Clostridia;o__Clostridiales;f__Ruminococcaceae;g__Anaerotruncus;s__sp34462 | 0.003897 | 0.000846 | 0.000495 | 0.000664 | 0.002151 | 0.001209 | 0.002669 | 0.000574 | 0 | 0.000579 | 0 | 0.000789 | 0.00072 | 0 | 0 | 0.00029 |
| k__Bacteria;p__Firmicutes;c__Clostridia;o__Clostridiales;f__Ruminococcaceae;g__Anaerotruncus;s__sp34462-sp34474 | 0 | 0 | 0 | 0 | 0 | 0 | 0 | 0.000182 | 0 | 0 | 0 | 0 | 0 | 0 | 0 | 0 |
| k__Bacteria;p__Firmicutes;c__Clostridia;o__Clostridiales;f__Ruminococcaceae;g__Anaerotruncus;s__sp34462-sp35834 | 0 | 0 | 0 | 0 | 0.000288 | 0.000382 | 0 | 0.000238 | 0.000518 | 0.000695 | 0.000431 | 0.000578 | 0 | 0.00037 | 0 | 0 |
| k__Bacteria;p__Firmicutes;c__Clostridia;o__Clostridiales;f__Ruminococcaceae;g__Anaerotruncus;s__sp34466 | 0 | 0.001909 | 0 | 0 | 0 | 0.000997 | 0 | 0 | 0 | 0 | 0 | 0 | 0 | 0 | 0 | 0 |
| k__Bacteria;p__Firmicutes;c__Clostridia;o__Clostridiales;f__Ruminococcaceae;g__Anaerotruncus;s__sp34466-sp34475 | 0 | 0 | 0 | 0 | 0 | 0 | 0 | 0 | 0 | 0 | 0 | 0 | 0 | 0.000687 | 0 | 0 |
| k__Bacteria;p__Firmicutes;c__Clostridia;o__Clostridiales;f__Ruminococcaceae;g__Anaerotruncus;s__sp34471 | 0.00282 | 0.001574 | 0.00044 | 0.000374 | 0.000776 | 0.001252 | 0 | 0.001134 | 0.001665 | 0.001529 | 0.000836 | 0 | 0.000254 | 0 | 0 | 0 |
| k__Bacteria;p__Firmicutes;c__Clostridia;o__Clostridiales;f__Ruminococcaceae;g__Anaerotruncus;s__sp34471-sp34483 | 0 | 0 | 0 | 0 | 0 | 0 | 0.000672 | 0.000588 | 0 | 0 | 0 | 0.002769 | 0 | 0.000846 | 0.00161 | 0.000397 |
| k__Bacteria;p__Firmicutes;c__Clostridia;o__Clostridiales;f__Ruminococcaceae;g__Anaerotruncus;s__sp34475 | 0.002769 | 0.001456 | 0 | 0 | 0.000954 | 0.002949 | 0 | 0.003163 | 0.00387 | 0.001715 | 0.00261 | 0.004136 | 0.003623 | 0.000978 | 0 | 0.000503 |
| k__Bacteria;p__Firmicutes;c__Clostridia;o__Clostridiales;f__Ruminococcaceae;g__Anaerotruncus;s__sp34481 | 0 | 0.000295 | 0 | 0 | 0 | 0 | 0 | 0 | 0 | 0 | 0 | 0.000333 | 0 | 0 | 0 | 0 |
| k__Bacteria;p__Firmicutes;c__Clostridia;o__Clostridiales;f__Ruminococcaceae;g__Anaerotruncus-Ruminiclostridium;s__sp34457-sp34934 | 0 | 0 | 0 | 0 | 0 | 0 | 0 | 0 | 0 | 0 | 0 | 0 | 0.000403 | 0 | 0 | 0 |
| k__Bacteria;p__Firmicutes;c__Clostridia;o__Clostridiales;f__Ruminococcaceae;g__NA;s__sp33140 | 0.002333 | 0.001574 | 0.00044 | 0.000353 | 0.001441 | 0.003268 | 0.002968 | 0.003051 | 0.00225 | 0.001437 | 0.00114 | 0.002033 | 0.001483 | 0.001242 | 0.000961 | 0.000336 |
| k__Bacteria;p__Firmicutes;c__Clostridia;o__Clostridiales;f__Ruminococcaceae;g__NA;s__sp34731 | 0.000795 | 0.000531 | 0 | 0 | 0.00051 | 0.000594 | 0 | 0.000588 | 0 | 0 | 0 | 0.000438 | 0 | 0.001638 | 0 | 0 |
| k__Bacteria;p__Firmicutes;c__Clostridia;o__Clostridiales;f__Ruminococcaceae;g__NA;s__sp34731-sp34771-sp34820 | 0 | 0 | 0 | 0 | 0 | 0 | 0 | 0.000182 | 0 | 0 | 0 | 0 | 0 | 0 | 0 | 0 |
| k__Bacteria;p__Firmicutes;c__Clostridia;o__Clostridiales;f__Ruminococcaceae;g__NA;s__sp34764 | 0 | 0 | 0 | 0 | 0 | 0 | 0 | 0 | 0 | 0 | 0 | 0 | 0.00036 | 0 | 0 | 0 |
| k__Bacteria;p__Firmicutes;c__Clostridia;o__Clostridiales;f__Ruminococcaceae;g__NA;s__sp34764-sp34771 | 0 | 0 | 0 | 0 | 0 | 0 | 0 | 0 | 0 | 0 | 0 | 0 | 0.000869 | 0 | 0 | 0 |
| k__Bacteria;p__Firmicutes;c__Clostridia;o__Clostridiales;f__Ruminococcaceae;g__NA;s__sp34771 | 0 | 0 | 0.00044 | 0 | 0 | 0 | 0 | 0 | 0 | 0 | 0 | 0 | 0 | 0 | 0 | 0 |
| k__Bacteria;p__Firmicutes;c__Clostridia;o__Clostridiales;f__Ruminococcaceae;g__NA;s__sp34771-sp34795-sp34820 | 0 | 0 | 0 | 0 | 0.000843 | 0 | 0 | 0 | 0 | 0 | 0 | 0 | 0 | 0 | 0 | 0 |
| k__Bacteria;p__Firmicutes;c__Clostridia;o__Clostridiales;f__Ruminococcaceae;g__NA;s__sp34771-sp34820 | 0 | 0 | 0 | 0 | 0 | 0 | 0 | 0 | 0 | 0 | 0 | 0 | 0 | 0.000344 | 0 | 0 |
| k__Bacteria;p__Firmicutes;c__Clostridia;o__Clostridiales;f__Ruminococcaceae;g__NA;s__sp34795 | 0 | 0.00059 | 0 | 0 | 0.000377 | 0 | 0.000616 | 0.000462 | 0.002183 | 0.000927 | 0 | 0.00028 | 0 | 0 | 0 | 0 |
| k__Bacteria;p__Firmicutes;c__Clostridia;o__Clostridiales;f__Ruminococcaceae;g__NA;s__sp34795-sp34820 | 0 | 0.000708 | 0 | 0.00108 | 0 | 0 | 0.000485 | 0 | 0 | 0.000927 | 0.000558 | 0 | 0.000551 | 0 | 0 | 0 |
| k__Bacteria;p__Firmicutes;c__Clostridia;o__Clostridiales;f__Ruminococcaceae;g__NA;s__sp34820 | 0.002948 | 0.000374 | 0 | 0 | 0.000998 | 0.001846 | 0 | 0.001176 | 0 | 0.002618 | 0.00109 | 0.001963 | 0 | 0 | 0.001827 | 0.000564 |
| k__Bacteria;p__Firmicutes;c__Clostridia;o__Clostridiales;f__Ruminococcaceae;g__NA;s__sp34822 | 0 | 0.000295 | 0 | 0 | 0 | 0 | 0 | 0.00021 | 0 | 0 | 0 | 0.000315 | 0.000381 | 0.000317 | 0 | 0 |
| k__Bacteria;p__Firmicutes;c__Clostridia;o__Clostridiales;f__Ruminococcaceae;g__NA;s__sp34826 | 0 | 0 | 0 | 0.000374 | 0 | 0 | 0.00922 | 0.000168 | 0 | 0 | 0 | 0 | 0 | 0 | 0.000312 | 0 |

| | | | | | | | | | | | | | | | | |
|---|---|---|---|---|---|---|---|---|---|---|---|---|---|---|---|---|
| k__Bacteria;p__Firmicutes;c__Clostridia;o__Clostridiales;f__Ruminococcaceae;g__NA;s__sp34838 | 0.008076 | 0 | 0 | 0.000748 | 0.002661 | 0.001995 | 0 | 0 | 0 | 0 | 0 | 0 | 0 | 0 | 0 | 0.000214 |
| k__Bacteria;p__Firmicutes;c__Clostridia;o__Clostridiales;f__Ruminococcaceae;g__NA;s__sp34849-sp34874 | 0 | 0 | 0 | 0 | 0 | 0.000509 | 0 | 0 | 0 | 0 | 0 | 0 | 0 | 0 | 0 | 0 |
| k__Bacteria;p__Firmicutes;c__Clostridia;o__Clostridiales;f__Ruminococcaceae;g__NA;s__sp34857 | 0 | 0 | 0 | 0 | 0 | 0 | 0 | 0.000882 | 0.001755 | 0 | 0 | 0 | 0.001653 | 0 | 0 | 0 |
| k__Bacteria;p__Firmicutes;c__Clostridia;o__Clostridiales;f__Ruminococcaceae;g__NA;s__sp34858-sp34873 | 0.004769 | 0.002578 | 0 | 0.00162 | 0.003903 | 0.005666 | 0.002669 | 0.003205 | 0.002318 | 0.001182 | 0.002154 | 0.005591 | 0.002394 | 0.002167 | 0.003581 | 0.001007 |
| k__Bacteria;p__Firmicutes;c__Clostridia;o__Clostridiales;f__Ruminococcaceae;g__NA;s__sp34863 | 0.000487 | 0.000728 | 0 | 0.000851 | 0 | 0 | 0.000485 | 0.001288 | 0 | 0.001576 | 0 | 0.000526 | 0.001356 | 0 | 0 | 0.000427 |
| k__Bacteria;p__Firmicutes;c__Clostridia;o__Clostridiales;f__Ruminococcaceae;g__NA;s__sp34867 | 0 | 0 | 0 | 0 | 0 | 0 | 0.000299 | 0.000196 | 0 | 0 | 0 | 0.000245 | 0.000403 | 0 | 0 | 0 |
| k__Bacteria;p__Firmicutes;c__Clostridia;o__Clostridiales;f__Ruminococcaceae;g__NA;s__sp34871 | 0 | 0.001377 | 0 | 0 | 0 | 0.006578 | 0.000597 | 0.002057 | 0.001845 | 0.001158 | 0.000634 | 0.001963 | 0.003496 | 0.000687 | 0.00274 | 0 |
| k__Bacteria;p__Firmicutes;c__Clostridia;o__Clostridiales;f__Ruminococcaceae;g__NA;s__sp34878 | 0.002923 | 0.003227 | 0.000879 | 0.00216 | 0.001619 | 0.004138 | 0.003714 | 0.007249 | 0.001733 | 0.001946 | 0.003624 | 0.006923 | 0 | 0.004308 | 0.002788 | 0.000854 |
| k__Bacteria;p__Firmicutes;c__Clostridia;o__Clostridiales;f__Ruminococcaceae;g__NA;s__sp34878-sp34883 | 0.00041 | 0.002735 | 0 | 0 | 0 | 0.002164 | 0 | 0 | 0.001778 | 0.001506 | 0.001622 | 0 | 0.001229 | 0.001559 | 0.002115 | 0 |
| k__Bacteria;p__Firmicutes;c__Clostridia;o__Clostridiales;f__Ruminococcaceae;g__NA;s__sp34879 | 0.002615 | 0.002184 | 0.002363 | 0.001516 | 0.001264 | 0.002037 | 0.001866 | 0.004478 | 0.00306 | 0.002804 | 0.002332 | 0.001945 | 0.001398 | 0.001401 | 0.001346 | 0.00241 |
| k__Bacteria;p__Firmicutes;c__Clostridia;o__Clostridiales;f__Ruminococcaceae;g__NA;s__sp34883 | 0 | 0 | 0 | 0 | 0 | 0.001591 | 0 | 0.000588 | 0 | 0 | 0 | 0 | 0.001547 | 0 | 0 | 0 |
| k__Bacteria;p__Firmicutes;c__Clostridia;o__Clostridiales;f__Ruminococcaceae;g__NA;s__sp34884 | 0.000487 | 0 | 0 | 0 | 0 | 0 | 0 | 0 | 0 | 0 | 0 | 0 | 0 | 0 | 0 | 0 |
| k__Bacteria;p__Firmicutes;c__Clostridia;o__Clostridiales;f__Ruminococcaceae;g__NA;s__sp34977 | 0 | 0.000433 | 0.000495 | 0 | 0.000244 | 0.000297 | 0 | 0.000546 | 0.000338 | 0.000348 | 0 | 0.000386 | 0 | 0 | 0 | 0.000549 |
| k__Bacteria;p__Firmicutes;c__Clostridia;o__Clostridiales;f__Ruminococcaceae;g__NA;s__sp34983 | 0.00041 | 0.000295 | 0 | 0 | 0 | 0.000488 | 0.000411 | 0.000602 | 0.000585 | 0 | 0 | 0.000526 | 0 | 0 | 0 | 0 |
| k__Bacteria;p__Firmicutes;c__Clostridia;o__Clostridiales;f__Ruminococcaceae;g__NA;s__sp35074 | 0 | 0.000295 | 0 | 0 | 0 | 0.000658 | 0 | 0 | 0.00027 | 0 | 0 | 0.000876 | 0 | 0 | 0 | 0 |
| k__Bacteria;p__Firmicutes;c__Clostridia;o__Clostridiales;f__Ruminococcaceae;g__NA;s__sp35077 | 0 | 0 | 0 | 0 | 0 | 0 | 0 | 0 | 0 | 0.000602 | 0 | 0 | 0 | 0 | 0 | 0 |
| k__Bacteria;p__Firmicutes;c__Clostridia;o__Clostridiales;f__Ruminococcaceae;g__NA;s__sp35077-sp35083 | 0 | 0 | 0 | 0 | 0 | 0 | 0 | 0 | 0 | 0 | 0.000405 | 0 | 0 | 0 | 0 | 0 |
| k__Bacteria;p__Firmicutes;c__Clostridia;o__Clostridiales;f__Ruminococcaceae;g__NA;s__sp35181 | 0.000513 | 0.000374 | 0.000467 | 0.001682 | 0.00031 | 0.000382 | 0.000373 | 0.000322 | 0.00126 | 0.000672 | 0.000811 | 0.000456 | 0.00072 | 0.001136 | 0.000745 | 0.00151 |
| k__Bacteria;p__Firmicutes;c__Clostridia;o__Clostridiales;f__Ruminococcaceae;g__NA;s__sp35200 | 0 | 0 | 0 | 0 | 0 | 0 | 0 | 0 | 0 | 0 | 0 | 0.00021 | 0 | 0 | 0 | 0 |
| k__Bacteria;p__Firmicutes;c__Clostridia;o__Clostridiales;f__Ruminococcaceae;g__NA;s__sp35215 | 0 | 0 | 0 | 0 | 0 | 0 | 0 | 0 | 0.000248 | 0 | 0 | 0 | 0 | 0 | 0 | 0.000229 |
| k__Bacteria;p__Firmicutes;c__Clostridia;o__Clostridiales;f__Ruminococcaceae;g__NA;s__sp35297 | 0 | 0 | 0 | 0 | 0 | 0 | 0.000299 | 0 | 0 | 0 | 0 | 0 | 0 | 0 | 0 | 0 |
| k__Bacteria;p__Firmicutes;c__Clostridia;o__Clostridiales;f__Ruminococcaceae;g__NA;s__sp35302 | 0.000359 | 0 | 0 | 0.000436 | 0 | 0 | 0.000411 | 0.000714 | 0 | 0 | 0 | 0 | 0 | 0 | 0 | 0 |
| k__Bacteria;p__Firmicutes;c__Clostridia;o__Clostridiales;f__Ruminococcaceae;g__NA;s__sp35310 | 0 | 0 | 0 | 0 | 0 | 0 | 0 | 0 | 0 | 0 | 0 | 0 | 0 | 0 | 0 | 0.00029 |
| k__Bacteria;p__Firmicutes;c__Clostridia;o__Clostridiales;f__Ruminococcaceae;g__NA;s__sp35326 | 0.000615 | 0 | 0 | 0 | 0 | 0 | 0 | 0 | 0 | 0 | 0 | 0 | 0 | 0 | 0 | 0 |
| k__Bacteria;p__Firmicutes;c__Clostridia;o__Clostridiales;f__Ruminococcaceae;g__NA;s__sp35337 | 0.001051 | 0.001988 | 0.002143 | 0.001163 | 0.002639 | 0.002143 | 0.001251 | 0.001525 | 0.001598 | 0.001691 | 0.003092 | 0.001507 | 0.003411 | 0.002986 | 0.000985 | 0.003142 |
| k__Bacteria;p__Firmicutes;c__Clostridia;o__Clostridiales;f__Ruminococcaceae;g__NA;s__sp35341 | 0 | 0 | 0 | 0 | 0 | 0 | 0 | 0 | 0.000518 | 0 | 0 | 0 | 0.000911 | 0 | 0 | 0 |
| k__Bacteria;p__Firmicutes;c__Clostridia;o__Clostridiales;f__Ruminococcaceae;g__NA;s__sp35345-sp35384 | 0 | 0 | 0 | 0 | 0 | 0.000403 | 0 | 0 | 0 | 0 | 0 | 0 | 0 | 0 | 0 | 0 |
| k__Bacteria;p__Firmicutes;c__Clostridia;o__Clostridiales;f__Ruminococcaceae;g__NA;s__sp35360 | 0 | 0.001869 | 0.00272 | 0.001952 | 0.001819 | 0.00053 | 0.001045 | 0.00077 | 0.001643 | 0.003082 | 0.00147 | 0.001052 | 0.001144 | 0.00222 | 0.001634 | 0.002349 |
| k__Bacteria;p__Firmicutes;c__Clostridia;o__Clostridiales;f__Ruminococcaceae;g__NA;s__sp35374-sp35384-sp35416 | 0 | 0 | 0 | 0 | 0 | 0 | 0 | 0 | 0 | 0.000788 | 0 | 0 | 0.000487 | 0 | 0 | 0 |
| k__Bacteria;p__Firmicutes;c__Clostridia;o__Clostridiales;f__Ruminococcaceae;g__NA;s__sp35377 | 0 | 0.000295 | 0 | 0 | 0 | 0 | 0 | 0 | 0 | 0 | 0 | 0 | 0 | 0 | 0 | 0 |
| k__Bacteria;p__Firmicutes;c__Clostridia;o__Clostridiales;f__Ruminococcaceae;g__NA;s__sp35382 | 0 | 0 | 0.001126 | 0 | 0 | 0 | 0 | 0 | 0 | 0 | 0 | 0 | 0 | 0 | 0 | 0.00154 |
| k__Bacteria;p__Firmicutes;c__Clostridia;o__Clostridiales;f__Ruminococcaceae;g__NA;s__sp35382-sp35388-sp35422 | 0 | 0 | 0 | 0 | 0 | 0 | 0 | 0 | 0 | 0.000324 | 0 | 0 | 0 | 0 | 0 | 0 |
| k__Bacteria;p__Firmicutes;c__Clostridia;o__Clostridiales;f__Ruminococcaceae;g__NA;s__sp35382-sp35403 | 0 | 0 | 0 | 0 | 0 | 0 | 0 | 0.000434 | 0 | 0 | 0 | 0 | 0 | 0 | 0 | 0 |
| k__Bacteria;p__Firmicutes;c__Clostridia;o__Clostridiales;f__Ruminococcaceae;g__NA;s__sp35383 | 0 | 0.000315 | 0 | 0.000353 | 0 | 0.001337 | 0 | 0.000476 | 0 | 0 | 0 | 0 | 0.000445 | 0 | 0 | 0.000183 |
| k__Bacteria;p__Firmicutes;c__Clostridia;o__Clostridiales;f__Ruminococcaceae;g__NA;s__sp35384 | 0 | 0.001869 | 0.000385 | 0 | 0 | 0.000679 | 0 | 0.000546 | 0.001013 | 0 | 0 | 0 | 0 | 0.000449 | 0 | 0.000366 |
| k__Bacteria;p__Firmicutes;c__Clostridia;o__Clostridiales;f__Ruminococcaceae;g__NA;s__sp35387 | 0 | 0 | 0 | 0 | 0 | 0 | 0 | 0 | 0 | 0.009639 | 0.009833 | 0 | 0.041891 | 0 | 0 | 0 |
| k__Bacteria;p__Firmicutes;c__Clostridia;o__Clostridiales;f__Ruminococcaceae;g__NA;s__sp35388 | 0 | 0 | 0.000659 | 0.000478 | 0 | 0 | 0 | 0.000308 | 0 | 0.000556 | 0 | 0 | 0.00053 | 0 | 0 | 0.000854 |
| k__Bacteria;p__Firmicutes;c__Clostridia;o__Clostridiales;f__Ruminococcaceae;g__NA;s__sp35388-sp35422-sp35423 | 0 | 0 | 0 | 0 | 0 | 0 | 0 | 0.000406 | 0 | 0 | 0.000532 | 0 | 0 | 0.000581 | 0 | 0.000427 |
| k__Bacteria;p__Firmicutes;c__Clostridia;o__Clostridiales;f__Ruminococcaceae;g__NA;s__sp35400 | 0 | 0 | 0.000385 | 0 | 0 | 0 | 0 | 0 | 0 | 0 | 0 | 0 | 0 | 0 | 0 | 0 |
| k__Bacteria;p__Firmicutes;c__Clostridia;o__Clostridiales;f__Ruminococcaceae;g__NA;s__sp35403 | 0 | 0.001397 | 0 | 0.002762 | 0.00051 | 0.000721 | 0.000448 | 0 | 0.000473 | 0.001274 | 0 | 0.000315 | 0.000742 | 0.001348 | 0.000769 | 0 |
| k__Bacteria;p__Firmicutes;c__Clostridia;o__Clostridiales;f__Ruminococcaceae;g__NA;s__sp35413 | 0 | 0 | 0 | 0 | 0 | 0 | 0 | 0 | 0 | 0 | 0 | 0 | 0 | 0.000793 | 0 | 0.000336 |
| k__Bacteria;p__Firmicutes;c__Clostridia;o__Clostridiales;f__Ruminococcaceae;g__NA;s__sp35419 | 0 | 0 | 0 | 0 | 0 | 0 | 0 | 0 | 0 | 0.000301 | 0 | 0 | 0 | 0 | 0 | 0 |
| k__Bacteria;p__Firmicutes;c__Clostridia;o__Clostridiales;f__Ruminococcaceae;g__NA;s__sp35423 | 0 | 0 | 0 | 0 | 0.000466 | 0 | 0.00084 | 0 | 0 | 0 | 0 | 0 | 0 | 0 | 0 | 0 |
| k__Bacteria;p__Firmicutes;c__Clostridia;o__Clostridiales;f__Ruminococcaceae;g__NA;s__sp35431 | 0 | 0 | 0 | 0 | 0 | 0 | 0.000336 | 0 | 0.000428 | 0 | 0 | 0.000456 | 0 | 0 | 0 | 0 |
| k__Bacteria;p__Firmicutes;c__Clostridia;o__Clostridiales;f__Ruminococcaceae;g__NA;s__sp35432 | 0 | 0 | 0 | 0.000623 | 0 | 0 | 0 | 0 | 0 | 0 | 0 | 0 | 0 | 0 | 0 | 0 |
| k__Bacteria;p__Firmicutes;c__Clostridia;o__Clostridiales;f__Ruminococcaceae;g__NA;s__sp35494 | 0 | 0.001673 | 0 | 0.003426 | 0 | 0.000276 | 0 | 0 | 0.006526 | 0.002224 | 0.002129 | 0.005135 | 0.011485 | 0 | 0.003077 | 0.000702 |
| k__Bacteria;p__Firmicutes;c__Clostridia;o__Clostridiales;f__Ruminococcaceae;g__NA;s__sp35498 | 0 | 0 | 0 | 0 | 0 | 0 | 0 | 0 | 0 | 0 | 0 | 0 | 0 | 0 | 0.00125 | 0 |
| k__Bacteria;p__Firmicutes;c__Clostridia;o__Clostridiales;f__Ruminococcaceae;g__NA;s__sp35499 | 0 | 0.004743 | 0 | 0 | 0 | 0.005411 | 0 | 0 | 0.024864 | 0.011886 | 0.012063 | 0.015283 | 0.011845 | 0.004334 | 0.019084 | 0.002684 |
| k__Bacteria;p__Firmicutes;c__Clostridia;o__Clostridiales;f__Ruminococcaceae;g__NA;s__sp35598 | 0.000923 | 0.001515 | 0.002637 | 0.004132 | 0.004458 | 0.000997 | 0.0014 | 0.001148 | 0.000968 | 0.001089 | 0.003041 | 0.000964 | 0.003327 | 0.002669 | 0.001514 | 0.00482 |
| k__Bacteria;p__Firmicutes;c__Clostridia;o__Clostridiales;f__Ruminococcaceae;g__NA;s__sp35687-sp35748 | 0.000923 | 0 | 0 | 0 | 0 | 0 | 0 | 0.000336 | 0 | 0 | 0.000786 | 0.000754 | 0 | 0 | 0 | 0 |
| k__Bacteria;p__Firmicutes;c__Clostridia;o__Clostridiales;f__Ruminococcaceae;g__NA;s__sp35688-sp35748 | 0 | 0 | 0 | 0.000519 | 0 | 0.000573 | 0.001064 | 0.000784 | 0.000855 | 0.000718 | 0 | 0 | 0.000848 | 0 | 0 | 0 |
| k__Bacteria;p__Firmicutes;c__Clostridia;o__Clostridiales;f__Ruminococcaceae;g__NA;s__sp35693 | 0.006281 | 0.005372 | 0.000632 | 0.002803 | 0.006941 | 0.014111 | 0.00293 | 0.008733 | 0.00612 | 0.002804 | 0.002737 | 0.003277 | 0.001949 | 0.003832 | 0.002524 | 0.001327 |
| k__Bacteria;p__Firmicutes;c__Clostridia;o__Clostridiales;f__Ruminococcaceae;g__NA;s__sp35696 | 0.006256 | 0.002165 | 0 | 0 | 0.00173 | 0.011374 | 0.001232 | 0.00049 | 0.006435 | 0.005584 | 0.002179 | 0.000631 | 0 | 0 | 0.000361 | 0.000503 |
| k__Bacteria;p__Firmicutes;c__Clostridia;o__Clostridiales;f__Ruminococcaceae;g__NA;s__sp35697 | 0 | 0 | 0 | 0 | 0 | 0.00053 | 0 | 0 | 0 | 0 | 0 | 0 | 0 | 0 | 0 | 0 |
| k__Bacteria;p__Firmicutes;c__Clostridia;o__Clostridiales;f__Ruminococcaceae;g__NA;s__sp35701 | 0 | 0.000256 | 0 | 0 | 0 | 0 | 0 | 0 | 0 | 0 | 0 | 0 | 0 | 0 | 0 | 0 |
| k__Bacteria;p__Firmicutes;c__Clostridia;o__Clostridiales;f__Ruminococcaceae;g__NA;s__sp35723 | 0 | 0 | 0 | 0 | 0 | 0 | 0 | 0 | 0 | 0 | 0 | 0 | 0.003645 | 0 | 0 | 0 |
| k__Bacteria;p__Firmicutes;c__Clostridia;o__Clostridiales;f__Ruminococcaceae;g__NA;s__sp35730 | 0.00141 | 0.007419 | 0 | 0.000602 | 0.000887 | 0.008382 | 0.002856 | 0.002323 | 0.004455 | 0.00139 | 0.002509 | 0.013285 | 0 | 0 | 0 | 0 |
| k__Bacteria;p__Firmicutes;c__Clostridia;o__Clostridiales;f__Ruminococcaceae;g__NA;s__sp35733 | 0.002743 | 0.000925 | 0.001374 | 0.000768 | 0.001087 | 0.002886 | 0.003714 | 0.003289 | 0.003443 | 0.001205 | 0.00185 | 0.003961 | 0 | 0 | 0 | 0 |
| k__Bacteria;p__Firmicutes;c__Clostridia;o__Clostridiales;f__Ruminococcaceae;g__NA;s__sp35736 | 0.000897 | 0.000728 | 0 | 0 | 0.001042 | 0 | 0.001195 | 0 | 0 | 0 | 0 | 0 | 0.000593 | 0.000396 | 0 | 0 |
| k__Bacteria;p__Firmicutes;c__Clostridia;o__Clostridiales;f__Ruminococcaceae;g__NA;s__sp35748 | 0.002307 | 0.001377 | 0 | 0 | 0.00071 | 0.001443 | 0.001139 | 0.000504 | 0.00072 | 0.001367 | 0 | 0.001683 | 0.004174 | 0.002299 | 0.002764 | 0.00061 |
| k__Bacteria;p__Firmicutes;c__Clostridia;o__Clostridiales;f__Ruminococcaceae;g__NA;s__sp35799 | 0.001256 | 0.000512 | 0 | 0 | 0 | 0.002058 | 0.000653 | 0.001176 | 0 | 0.000695 | 0 | 0 | 0.000763 | 0.001665 | 0.001731 | 0.000427 |
| k__Bacteria;p__Firmicutes;c__Clostridia;o__Clostridiales;f__Ruminococcaceae;g__NA;s__sp35832 | 0 | 0.000669 | 0 | 0.000415 | 0.001064 | 0.000976 | 0 | 0.001973 | 0 | 0.000741 | 0.001597 | 0.002156 | 0.001547 | 0 | 0.001322 | 0.000473 |
| k__Bacteria;p__Firmicutes;c__Clostridia;o__Clostridiales;f__Ruminococcaceae;g__NA;s__sp35837 | 0 | 0 | 0 | 0 | 0 | 0 | 0 | 0 | 0 | 0.000788 | 0 | 0 | 0 | 0 | 0 | 0 |
| k__Bacteria;p__Firmicutes;c__Clostridia;o__Clostridiales;f__Ruminococcaceae;g__NA;s__sp35841 | 0.004615 | 0.002912 | 0.000714 | 0.000561 | 0.000488 | 0.002759 | 0.001717 | 0.004324 | 0.00099 | 0.001506 | 0.001926 | 0.00312 | 0.000784 | 0.000766 | 0.000961 | 0.00058 |
| k__Bacteria;p__Firmicutes;c__Clostridia;o__Clostridiales;f__Ruminococcaceae;g__NA;s__sp35849 | 0.000667 | 0.000275 | 0 | 0 | 0 | 0.000446 | 0 | 0.000392 | 0.00117 | 0 | 0.000279 | 0.001104 | 0 | 0 | 0 | 0 |
| k__Bacteria;p__Firmicutes;c__Clostridia;o__Clostridiales;f__Ruminococcaceae;g__NA;s__sp35869 | 0 | 0 | 0 | 0.00081 | 0 | 0 | 0 | 0 | 0 | 0 | 0 | 0 | 0 | 0 | 0 | 0 |
| k__Bacteria;p__Firmicutes;c__Clostridia;o__Clostridiales;f__Ruminococcaceae;g__NA;s__sp35873 | 0 | 0 | 0 | 0 | 0 | 0.001719 | 0 | 0.001106 | 0.001508 | 0.000371 | 0.001014 | 0.000333 | 0 | 0 | 0.000697 | 0 |
| k__Bacteria;p__Firmicutes;c__Clostridia;o__Clostridiales;f__Ruminococcaceae;g__NA;s__sp35877 | 0.000513 | 0.000335 | 0 | 0 | 0 | 0.000679 | 0.000448 | 0.000714 | 0 | 0 | 0 | 0.000386 | 0 | 0 | 0 | 0 |
| k__Bacteria;p__Firmicutes;c__Clostridia;o__Clostridiales;f__Ruminococcaceae;g__NA;s__sp35882 | 0 | 0.003424 | 0 | 0.001059 | 0 | 0.000509 | 0.000541 | 0.000868 | 0.003825 | 0 | 0 | 0.001788 | 0 | 0 | 0 | 0 |
| k__Bacteria;p__Firmicutes;c__Clostridia;o__Clostridiales;f__Ruminococcaceae;g__Oscillibacter;s__sp34634 | 0.031355 | 0.008954 | 0.001951 | 0.00353 | 0.007629 | 0.01061 | 0.00545 | 0.015199 | 0.004163 | 0.002271 | 0.002686 | 0.004504 | 0.008433 | 0.003858 | 0.00399 | 0.002379 |
| k__Bacteria;p__Firmicutes;c__Clostridia;o__Clostridiales;f__Ruminococcaceae;g__Oscillibacter;s__sp34638 | 0.001282 | 0.001358 | 0 | 0 | 0.000754 | 0.001443 | 0.000709 | 0.001092 | 0.00207 | 0.000695 | 0.000912 | 0.001577 | 0.001123 | 0.000581 | 0.000697 | 0 |
| k__Bacteria;p__Firmicutes;c__Clostridia;o__Clostridiales;f__Ruminococcaceae;g__Oscillibacter;s__sp34638-sp34646 | 0 | 0.000335 | 0 | 0 | 0 | 0 | 0 | 0 | 0 | 0 | 0.000684 | 0 | 0 | 0 | 0 | 0 |
| k__Bacteria;p__Firmicutes;c__Clostridia;o__Clostridiales;f__Ruminococcaceae;g__Oscillibacter;s__sp34640 | 0 | 0 | 0 | 0 | 0 | 0.000509 | 0 | 0 | 0 | 0.001321 | 0 | 0.003698 | 0 | 0 | 0 | 0 |
| k__Bacteria;p__Firmicutes;c__Clostridia;o__Clostridiales;f__Ruminococcaceae;g__Oscillibacter;s__sp34642 | 0.003743 | 0.005116 | 0 | 0 | 0.004147 | 0.005602 | 0.005245 | 0.002799 | 0.00216 | 0.002317 | 0.001723 | 0.001525 | 0.001674 | 0.00111 | 0.001442 | 0.00058 |
| k__Bacteria;p__Firmicutes;c__Clostridia;o__Clostridiales;f__Ruminococcaceae;g__Oscillibacter;s__sp34643 | 0.002359 | 0.002342 | 0 | 0.00162 | 0.001486 | 0.004201 | 0.001288 | 0.001483 | 0.000743 | 0.00088 | 0 | 0.002787 | 0.00036 | 0.000608 | 0.001923 | 0 |
| k__Bacteria;p__Firmicutes;c__Clostridia;o__Clostridiales;f__Ruminococcaceae;g__Oscillibacter;s__sp34645 | 0.000359 | 0 | 0 | 0 | 0 | 0.001273 | 0.001101 | 0.000672 | 0.000338 | 0.000556 | 0 | 0.002471 | 0.000445 | 0 | 0.00387 | 0 |
| k__Bacteria;p__Firmicutes;c__Clostridia;o__Clostridiales;f__Ruminococcaceae;g__Oscillibacter;s__sp34646 | 0 | 0 | 0 | 0 | 0 | 0.003395 | 0 | 0 | 0 | 0 | 0 | 0.000561 | 0 | 0 | 0.001923 | 0.000442 |

| | | | | | | | | | | | | | | | | |
|---|---|---|---|---|---|---|---|---|---|---|---|---|---|---|---|---|
| k__Bacteria;p__Firmicutes;c__Clostridia;o__Clostridiales;f__Ruminococcaceae;g__Oscillibacter;s__sp34648 | 0 | 0 | 0 | 0 | 0.001198 | 0.00157 | 0.000765 | 0.00084 | 0 | 0 | 0 | 0.000438 | 0.000487 | 0 | 0.000985 | 0.000381 |
| k__Bacteria;p__Firmicutes;c__Clostridia;o__Clostridiales;f__Ruminococcaceae;g__Oscillibacter;s__sp34650 | 0 | 0 | 0 | 0 | 0 | 0 | 0 | 0 | 0 | 0 | 0 | 0 | 0 | 0.00037 | 0 | 0 |
| k__Bacteria;p__Firmicutes;c__Clostridia;o__Clostridiales;f__Ruminococcaceae;g__Oscillibacter;s__sp34650-sp34654 | 0.004743 | 0 | 0.001154 | 0 | 0 | 0 | 0 | 0.00056 | 0.00243 | 0 | 0 | 0 | 0.005933 | 0 | 0 | 0 |
| k__Bacteria;p__Firmicutes;c__Clostridia;o__Clostridiales;f__Ruminococcaceae;g__Oscillibacter-Oscillospira;s__sp34648-sp34660 | 0 | 0.004998 | 0 | 0.000602 | 0 | 0.006111 | 0.000411 | 0.003513 | 0.003938 | 0.002363 | 0.003573 | 0.00461 | 0.005615 | 0.001321 | 0.003677 | 0.001083 |
| k__Bacteria;p__Firmicutes;c__Clostridia;o__Clostridiales;f__Ruminococcaceae;g__Ruminiclostridium;s__sp34857-sp34950 | 0 | 0 | 0 | 0 | 0 | 0 | 0 | 0.001511 | 0 | 0 | 0 | 0 | 0 | 0.001004 | 0 | 0 |
| k__Bacteria;p__Firmicutes;c__Clostridia;o__Clostridiales;f__Ruminococcaceae;g__Ruminiclostridium;s__sp34901 | 0 | 0.001456 | 0 | 0.000789 | 0 | 0 | 0.002688 | 0 | 0 | 0 | 0 | 0 | 0 | 0.004915 | 0 | 0 |
| k__Bacteria;p__Firmicutes;c__Clostridia;o__Clostridiales;f__Ruminococcaceae;g__Ruminiclostridium;s__sp34916 | 0 | 0 | 0.000467 | 0 | 0 | 0 | 0 | 0.008425 | 0.007111 | 0.003661 | 0.005145 | 0.013723 | 0.012396 | 0 | 0.008124 | 0 |
| k__Bacteria;p__Firmicutes;c__Clostridia;o__Clostridiales;f__Ruminococcaceae;g__Ruminiclostridium;s__sp34925 | 0.003615 | 0.001043 | 0 | 0.000457 | 0.000821 | 0.002207 | 0.000467 | 0 | 0 | 0 | 0 | 0 | 0 | 0.000529 | 0 | 0.000717 |
| k__Bacteria;p__Firmicutes;c__Clostridia;o__Clostridiales;f__Ruminococcaceae;g__Ruminiclostridium;s__sp34944 | 0 | 0 | 0 | 0 | 0 | 0 | 0 | 0 | 0 | 0 | 0 | 0.000438 | 0 | 0 | 0 | 0 |
| k__Bacteria;p__Firmicutes;c__Clostridia;o__Clostridiales;f__Ruminococcaceae;g__Ruminiclostridium;s__sp34950 | 0.001231 | 0.001122 | 0.001099 | 0 | 0 | 0 | 0.000411 | 0 | 0 | 0.000765 | 0 | 0 | 0 | 0 | 0.000625 | 0 |
| k__Bacteria;p__Firmicutes;c__Clostridia;o__Clostridiales;f__Ruminococcaceae;g__Subdoligranulum;s__sp35590 | 0.000538 | 0.001633 | 0.000412 | 0.00027 | 0 | 0.001931 | 0.000373 | 0.000574 | 0.00243 | 0.00139 | 0.002788 | 0.005188 | 0.002288 | 0.000634 | 0.002043 | 0.000259 |
| k__Bacteria;p__Firmicutes;c__Erysipelotrichia;o__Erysipelotrichales;f__Erysipelotrichaceae;g__Allobaculum;s__sp36555 | 0.002154 | 0.001712 | 0.004148 | 0 | 0 | 0 | 0 | 0 | 0 | 0 | 0.000938 | 0.001069 | 0.014027 | 0.002854 | 0.01358 | 0.002806 |
| k__Bacteria;p__Firmicutes;c__Erysipelotrichia;o__Erysipelotrichales;f__Erysipelotrichaceae;g__Erysipelatoclostridium;s__sp36617 | 0 | 0 | 0 | 0 | 0 | 0 | 0 | 0 | 0 | 0 | 0 | 0 | 0.000297 | 0 | 0 | 0.000625 |
| k__Bacteria;p__Firmicutes;c__Erysipelotrichia;o__Erysipelotrichales;f__Erysipelotrichaceae;g__NA;s__sp36773 | 0 | 0 | 0 | 0 | 0 | 0 | 0 | 0 | 0 | 0 | 0 | 0 | 0 | 0 | 0 | 0.000519 |
| k__Bacteria;p__Firmicutes;c__Erysipelotrichia;o__Erysipelotrichales;f__Erysipelotrichaceae;g__NA;s__sp36777 | 0 | 0.000748 | 0.001181 | 0.001557 | 0.000754 | 0 | 0.000485 | 0.00112 | 0.000428 | 0 | 0.000811 | 0 | 0 | 0.000951 | 0 | 0.001617 |
| k__Bacteria;p__Firmicutes;c__Erysipelotrichia;o__Erysipelotrichales;f__Erysipelotrichaceae;g__NA;s__sp36783 | 0 | 0.00061 | 0 | 0.000311 | 0.001575 | 0 | 0 | 0 | 0 | 0.000278 | 0 | 0.000403 | 0 | 0 | 0.000264 | 0 |
| k__Bacteria;p__Firmicutes;c__Erysipelotrichia;o__Erysipelotrichales;f__Erysipelotrichaceae;g__NA;s__sp36787 | 0 | 0 | 0 | 0.000519 | 0.000266 | 0.000573 | 0 | 0.000252 | 0 | 0 | 0.000329 | 0.000228 | 0.000403 | 0 | 0 | 0.000686 |
| k__Bacteria;p__Firmicutes;c__Erysipelotrichia;o__Erysipelotrichales;f__Erysipelotrichaceae;g__Turicibacter;s__sanguinis | 0 | 0.004959 | 0.014341 | 0 | 0 | 0.000424 | 0 | 0.00021 | 0.014289 | 0.019972 | 0.018576 | 0.00822 | 0.00161 | 0.000661 | 0.000553 | 0.00032 |
| k__Bacteria;p__Proteobacteria;c__Alphaproteobacteria;o__Rhodospirillales;f__Rhodospirillaceae;g__Thalassospira;s__sp46235 | 0 | 0.001082 | 0.001071 | 0.001827 | 0 | 0.000361 | 0.000299 | 0.000532 | 0.004298 | 0.003892 | 0.002332 | 0.00149 | 0.000954 | 0 | 0.000312 | 0.000824 |
| k__Bacteria;p__Proteobacteria;c__Betaproteobacteria;o__Burkholderiales;f__Alcaligenaceae;g__Parasutterella;s__excrementihominis | 0 | 0 | 0 | 0 | 0 | 0 | 0 | 0 | 0.000495 | 0.000996 | 0.000938 | 0.000368 | 0.000975 | 0.00037 | 0.00125 | 0.000686 |
| k__Bacteria;p__Proteobacteria;c__Betaproteobacteria;o__Burkholderiales;f__Alcaligenaceae;g__Parasutterella;s__sp48235 | 0.003948 | 0.005274 | 0.015522 | 0.003385 | 0.004502 | 0.001634 | 0.004349 | 0.0048 | 0.00135 | 0.001761 | 0.000608 | 0.000543 | 0.005679 | 0.004149 | 0.003125 | 0.006009 |
| k__Bacteria;p__Proteobacteria;c__Deltaproteobacteria;o__Desulfovibrionales;f__Desulfovibrionaceae;g__Bilophila;s__sp52475 | 0.007307 | 0.004979 | 0.001374 | 0 | 0.000444 | 0.00505 | 0.001064 | 0.005654 | 0.00081 | 0.000672 | 0.000304 | 0.001525 | 0.001038 | 0 | 0.000457 | 0 |
| k__Bacteria;p__Proteobacteria;c__Deltaproteobacteria;o__Desulfovibrionales;f__Desulfovibrionaceae;g__Desulfovibrio;s__sp52643 | 0 | 0 | 0 | 0.000498 | 0.000976 | 0 | 0.002837 | 0 | 0.014401 | 0.010079 | 0.008718 | 0.004382 | 0.012735 | 0.007849 | 0.007715 | 0.00392 |
| k__Bacteria;p__Proteobacteria;c__Gammaproteobacteria;o__Enterobacteriales;f__Enterobacteriaceae;g__Escherichia-Shigella;s__coli | 0.000564 | 0 | 0 | 0 | 0.000266 | 0 | 0 | 0 | 0 | 0 | 0 | 0 | 0 | 0 | 0 | 0 |
| k__Bacteria;p__Saccharibacteria;c__NA;o__NA;f__NA;g__*Saccharimonas;s__sp65946 | 0 | 0.001279 | 0 | 0.000311 | 0 | 0.000361 | 0 | 0.00077 | 0.000743 | 0.004055 | 0.003168 | 0.000298 | 0.001229 | 0.001163 | 0.000793 | 0.001266 |
| k__Bacteria;p__Tenericutes;c__Mollicutes;o__Anaeroplasmatales;f__Anaeroplasmataceae;g__Anaeroplasma;s__sp67615 | 0.001718 | 0.002263 | 0.000879 | 0.000644 | 0 | 0.001698 | 0 | 0.003023 | 0.00189 | 0.003058 | 0.001521 | 0.002033 | 0 | 0.000555 | 0.002644 | 0.001601 |
| k__Bacteria;p__Tenericutes;c__Mollicutes;o__NA;f__NA;g__NA;s__sp67838 | 0 | 0.001122 | 0 | 0 | 0 | 0.00034 | 0 | 0 | 0 | 0 | 0 | 0 | 0 | 0 | 0 | 0 |
| k__Bacteria;p__Tenericutes;c__Mollicutes;o__NA;f__NA;g__NA;s__sp67844 | 0 | 0.000394 | 0 | 0 | 0 | 0 | 0 | 0.001232 | 0.000383 | 0.000857 | 0.001318 | 0.000876 | 0.001271 | 0 | 0 | 0 |
| k__Bacteria;p__Tenericutes;c__Mollicutes;o__NA;f__NA;g__NA;s__sp67901 | 0 | 0 | 0 | 0 | 0.000421 | 0 | 0 | 0 | 0 | 0 | 0 | 0 | 0 | 0 | 0 | 0 |
| k__Bacteria;p__Tenericutes;c__Mollicutes;o__NA;f__NA;g__NA;s__sp67923 | 0 | 0 | 0 | 0 | 0 | 0 | 0 | 0.000182 | 0 | 0 | 0 | 0 | 0 | 0 | 0 | 0 |
| k__Bacteria;p__Tenericutes;c__Mollicutes;o__NA;f__NA;g__NA;s__sp67941 | 0 | 0 | 0 | 0.000311 | 0 | 0 | 0 | 0 | 0 | 0 | 0 | 0 | 0 | 0 | 0 | 0 |
| k__Bacteria;p__Verrucomicrobia;c__Verrucomicrobiae;o__Verrucomicrobiales;f__Verrucomicrobiaceae;g__Akkermansia;s__muciniphila | 0 | 0 | 0 | 0 | 0 | 0 | 0 | 0 | 0 | 0 | 0 | 0 | 0.041488 | 0 | 0 | 0 |
